# *J-AST*: a web-based analysis platform for antimicrobial susceptibility testing

**DOI:** 10.64898/2026.07.22.739984

**Authors:** Ruman Gerst, Austin Mottola, Dhruv Khatri, Judith Berman, Marc Thilo Figge

## Abstract

Antimicrobial resistance and tolerance pose escalating global health threats, necessitating reproducible and accessible tools for antimicrobial susceptibility testing (AST). While disk diffusion assays (DDAs) and Epsilometer tests (Etests) are widely used, there are limited open-source tools to analyze them. We present *J-AST*, a free, open-source, web-based platform for analyzing both DDAs and Etests. It provides automated and interactive annotation of regions of interest and metadata management, and quantifies microbial resistance and tolerance. *J-AST* outputs correlate strongly with those of existing tools, and equivalent DDA and Etest results correlate strongly with each other. The automated MIC detection achieved >90% agreement with manual readouts. *J-AST* is deployable both as desktop software and cloud service, unifies automated analysis with interactive review, and advances both fundamental research and clinical AST workflows.

## Introduction

Mortality due to infectious diseases impose a serious global burden. In 2021 alone, sepsis was associated with 21.4 million deaths worldwide, of which, 4.71 million of which were linked to antibacterial drug resistance ^1^. The population-level phenotype of drug resistance is characterized by the ability of microbes to grow well at drug concentrations above the minimum inhibitory concentration (MIC) established for representative strains. While most work has focused on bacterial infections, an increasing number of deaths is also associated with fungal infections. Increasing proportions of these infections are caused by intrinsically resistant fungal species ^2^. Furthermore, antifungal drug tolerance, heteroresistance, and other alternative mechanisms are used by bacteria and fungi to grow despite drug concentrations above the MIC. These drug responses include mechanistically distinct pathways that are less well understood than resistance, and usually involve subpopulations of cells that grow and/or survive in drug concentrations above the standard MIC for a given species ^3,4^. This escalating threat highlights the urgent necessity for reproducible, cost-efficient, and openly accessible approaches for antimicrobial susceptibility tests (ASTs), complemented by rigorous computational analyses, to support both fundamental research and clinical decision-making.

Clinically relevant ASTs standardized by CLSI and EUCAST^5^ fall into two broad categories: broth dilution assays (BMDAs) that are considered the gold standard and involve testing microbial growth in liquid medium containing serial drug dilutions in microtiter plate assays, and agar-based diffusion assay. BMDAs are laborious and costly, while diffusion assay can be assembled in-house or performed with commercially available kits (e.g., Bruker MICRONAUT-AM or Thermo Scientific Sensititre YeastOne AST Plate).

Diffusion-based assays use agar plates and a drug source that diffuses into the medium under a lawn of cells. The two main diffusion assay modalities are Disk Diffusion Assays (DDAs) and Epsilometer tests (Etests). DDAs provide the antimicrobial agent in a small disk-shaped filter paper that is positioned on an agar plate; such disks can be produced in-house or purchased commercially. The disks generally contain the amount of drug specified by CLSI and/or EUCAST for testing a given drug with a particular species. The degree of antimicrobial susceptibility is a measurement of the radius or diameter of the circular zone of inhibition (ZOI), the region where microbial growth is inhibited. MIC determinations are made by comparing ZOI parameters of test strains relative to susceptible and resistant strains used as standards.

Etest strips are available commercially (e.g. from bioMeriéux, Meizheng, etc.) and contain a gradient of the relevant antimicrobial that is placed on the lawn of cells. Alignment of the ZOI with pre-defined graduations on the strip facilitates direct measurement of MIC by visual inspection of the minimum drug concentration inhibiting growth of the isolate. Currently, no commonly used open source tools are available for digitally extracting the MIC from Etests; Scientists are limited to commercial options provided with imaging systems, for example BIOMIC V3 (Giles Scientific Inc.) or VITEK (bioMérieux SA).

*diskImageR* ^6^ is a freely available open source R-based image analysis pipeline built on *ImageJ* ^7,8^ and R (https://www.r-project.org/) that determines growth-related measurements from individual images of DDAs^9,10^. It collects pixel intensity measurements across 72 radii running from the drug disk in the center of the DDA to points near the agar plate edge, averages them as a function of distance from the disk, and plots the resulting curves. The generally sigmoidal curves provide parameters for maximum growth, using radii corresponding to 20, 50 and 80% inhibition of growth relative to the maximum growth to quantify resistance. It also measures tolerance as the fraction of growth (FoG) inside the ZOI, obtained by calculating the pixel intensity within the ZOI starting from the radial position determined by the RAD_20_ or RAD_50_ relative to full growth outside of the ZOI. Ideally, tolerance (FoG) measurements require a comparison of growth within the ZOI at different time points, for example, using the RAD_20_ position at 24h of growth to determine the FoG_20_ at 48h. However, this needs to be done manually and most studies use the RAD_20_ position at 24h, based on the assumption that RAD values do not change considerably between 24h and 48h ^11^. Unfortunately, this assumption does not hold for all strains, and thus a computational tool that can compare 24h and 48h images of growth on the same plate would improve tolerance measurements. *diskImageR* also lacks interactive review of intermediate results, enforces a strict naming scheme for data input, and is limited to DDAs.

Here, we describe the JIPipe Antimicrobial Susceptibility Test (*J-AST*) platform for analyzing drug susceptibility and antifungal drug tolerance from both DDAs and Etests. *J-AST* quantifies resistance (RAD) and tolerance (FoG) via timeline analysis across two time points, provides automated MIC extraction from Etests, facilitates metadata-driven analysis, and integrates automated annotation with interactive review. *J-AST* is deployable as desktop software and cloud service and contributes towards the Findability Accessibility Interoperability, and Reproducibility (FAIR) ^12^ of diffusion assay analysis.

## Results

### *J-AST* enables flexible analysis of disk diffusion and Etest assays

The *J-AST* platform provides an all-in-one solution for managing AST analysis projects in one modern user interface (**Fig. 1**). *J-AST* provides users with full flexibility to choose from a set of powerful automated and interactive tools for organizing, annotating, and analyzing datasets with single or multiple time points. A key feature is the tight integration between automated and interactive tools, which combines the benefits of both workflows. Users can freely combine interactive and automated tooling for image and metadata processing. *J-AST* is built on modern open-source web-based technology that is platform-independent and can be easily modified and expanded. It provides the same user experience both as a cloud-based web application, and as an offline desktop software. A guest access with limits on project size and available data storage, as well as documentation and tutorials are available at https://asb.hki-jena.de/j-ast.

**Fig. 1:**
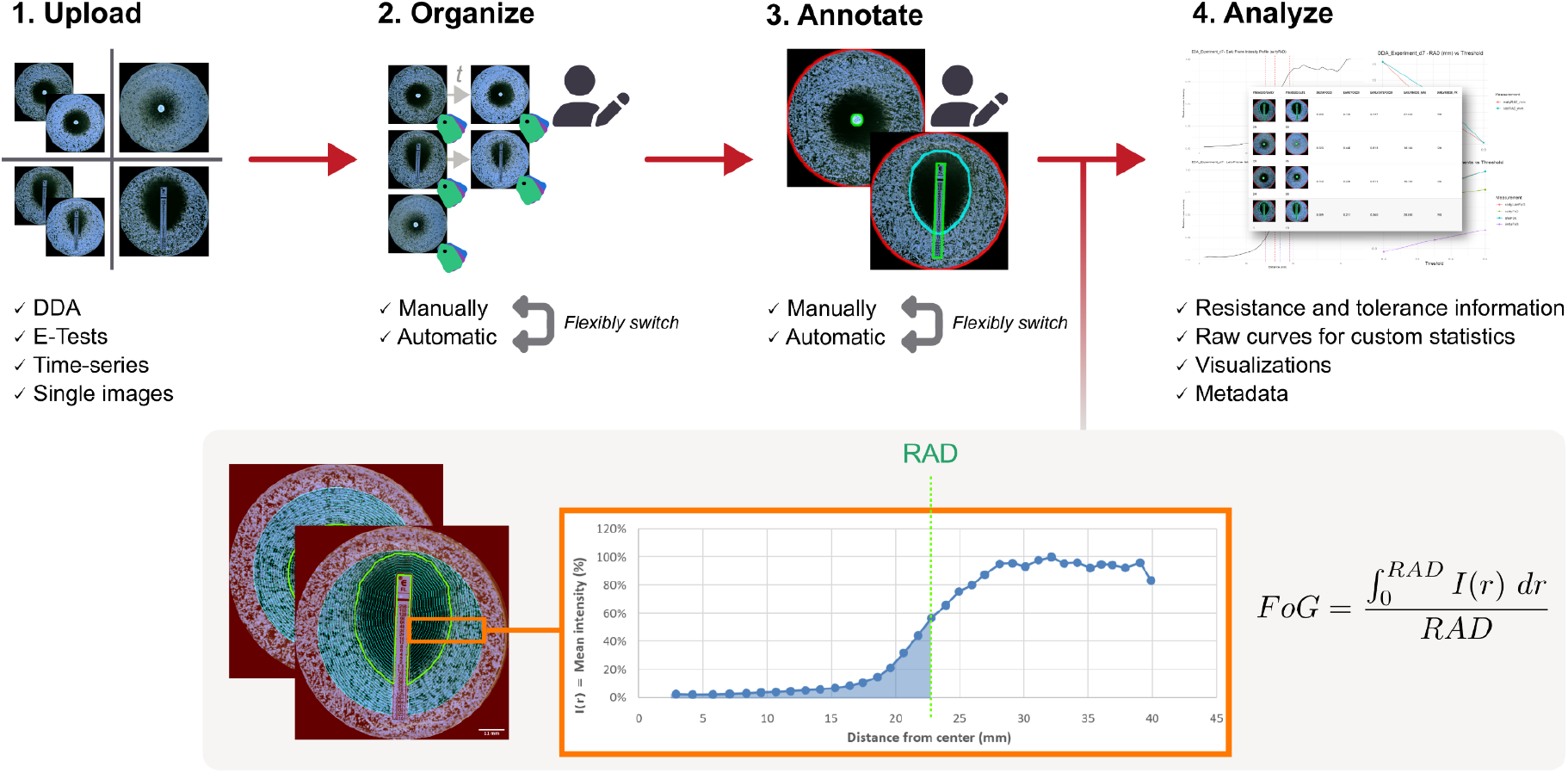
Analyzing data with *J-AST*. The process begins with the upload of the images. Users have the option to organize the data manually through the user interface, or to use the automated operations. Then, relevant ROIs are annotated through automated algorithms and manual inputs, with users again having complete flexibility to combine tools of their choice. The default analysis algorithm extracts resistance and tolerance measurements by comparing intensity measurements over the distance to the ZOI center.

### *J-AST* provides metadata-driven image management and annotation workflows

*J-AST* users can manage multiple projects for the analysis of independent images, or for organizing and processing a time series (**Fig. 2**). The necessary metadata fields, including the “Experiment”, “Sample”, and “Assay type”, as well as the timeline order can be filled out manually or through automated methods.

**Fig. 2:**
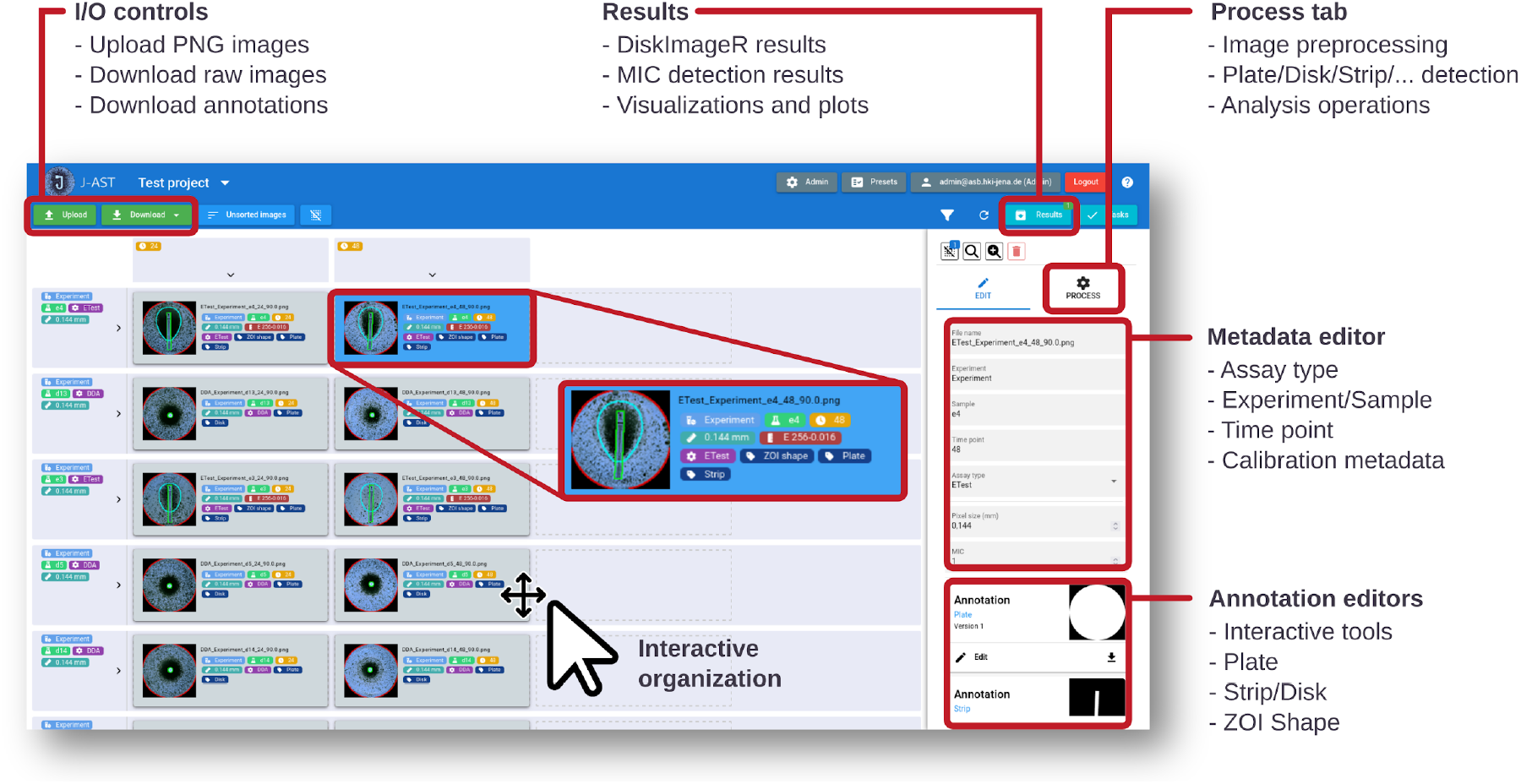
*J-AST* interface overview. At the top left, the web interface contains operations that allow image uploading and downloading. If at least one image is selected, a variety of fully automated image analysis and processing operations are offered through a “Process” tab. At any point, metadata and ROI-based annotations can be edited interactively via a panel on the right-hand side. Images are either organized into timelines through interactive or automated methods, or handled independently. All generated results are organized through a dedicated page.

A core concept of *J-AST* is the dynamic management of per-image regions of interest (ROIs), such as the plate, DDA disk, and Etest strip (**Supplementary Figures 3a** and **3b**). The standard annotation workflow comprises plate detection (**Supplementary Section 1.2.1**), followed by calibration of the pixel size (**Supplementary Section 1.2.2**), and then bifurcates depending on the assay type. DDA images require an outline of the disk region (**Supplementary Section 1.2.3**); Etests require a ROI that highlights the strip and the ZOI shape (**Supplementary Sections 1.2.4 and 1.2.5**). *J-AST* provides a comprehensive and extensible set of fully automated algorithms that detect all of these ROIs within the uploaded images that includes two plate detection algorithms (**Supplementary Section 2.1.1 and 2.1.2**), physical pixel size calibration (**Supplementary Section 2.2**), three DDA disk segmentation tools (**Supplementary Sections 2.3.1-2.3.4**), a workflow for Etest strip detection (**Supplementary Section 2.4**), and outlining of the ZOI shape (**Supplementary Section 2.5**). This set of algorithms is complemented by powerful interactive tools (**Supplementary Figure 3c** and **Supplementary Section 1.2.6**) for manual segmentation.

### *J-AST* computation demonstrates quantitative agreement with *diskImageR*

We developed *J-AST* using the visual programming platform *JIPipe*^13^; the graphical programming and semantic versioning for both JIPipe and associated third-party software makes the workflow easier to maintain and run. *J-AST* allows users to quantify the colony growth metrics by first calculating the radius of inhibition (*RAD*_*n*_) for customizable inhibition thresholds n, by default 20%, 50% and 80%. It then quantifies growth within the radius given by *RAD*_*n*_ , producing a measurement termed *fraction of growth* (*FOG*_*n*_) for the threshold n. Unlike *diskImageR*, which samples 72 radial line profiles, *J-AST* generates ring-shaped measurement regions (default width 1 mm) around the disk, excluding a safety distance (default 10 mm) from the plate border (**Fig. 3a**). The average colony intensities are used to derive the *RAD*_*n*_ , *FOG*_*n*_ , and other metrics including, *FOG*_*early, n*_, *FOG* _*late, n*_ , and Δ*FOG*_*n*_ = *FOG* _*late, n*_ − *FOG* _*early, n*_ via normalized intensity curves (**Supplementary Section 2.6**). A single-image mode is also included (**Supplementary Section 2.7**).

**Fig. 3:**
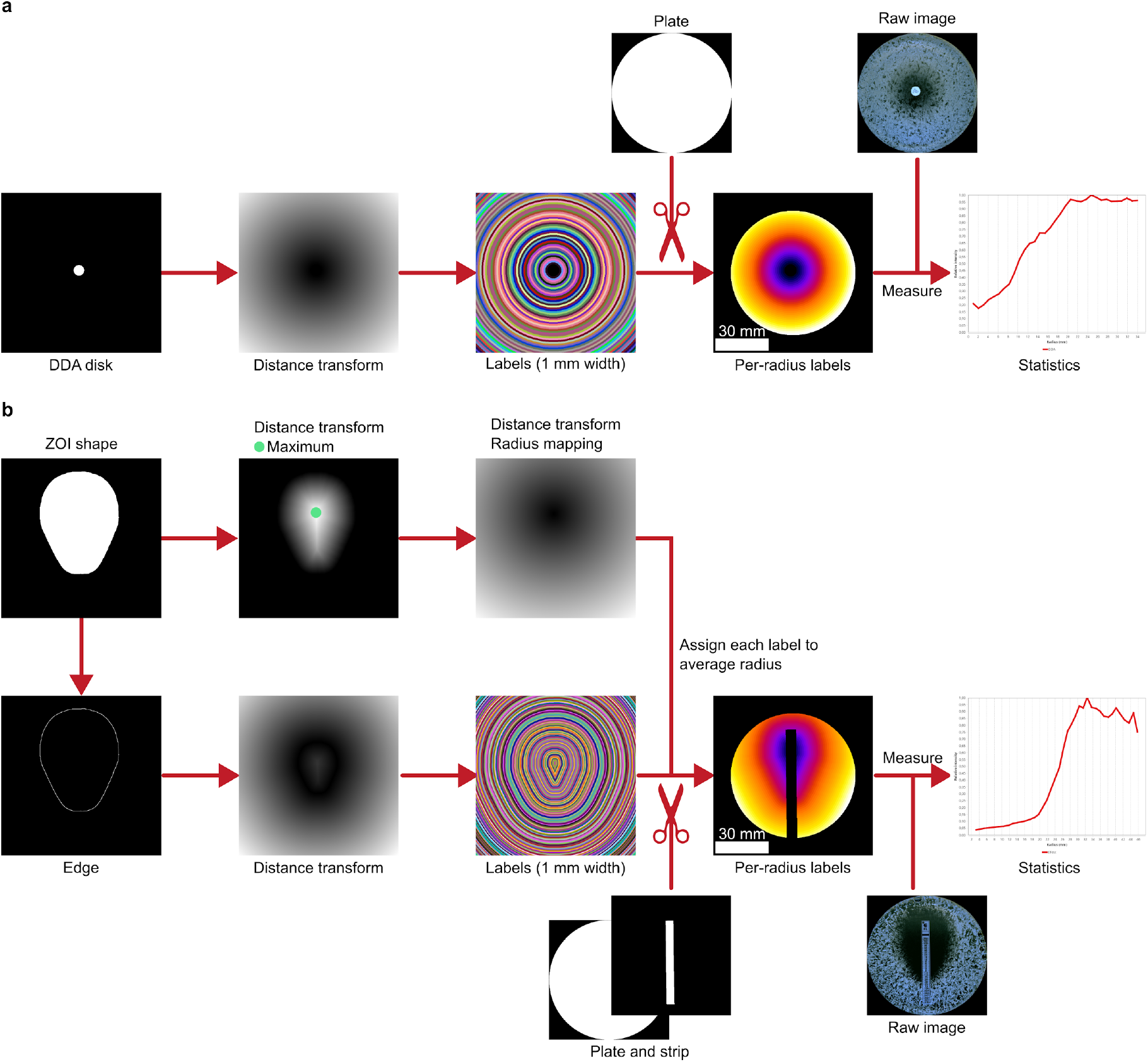
*J-AST* implementation of disk diffusion assay and Etest analyses. **(a)** For DDAs, the algorithm starts with the disk annotation that is processed via a distance transform. The resulting per-pixel distances are converted into labels that are used to measure the average colony intensity per label. **(b)** The processing of Etests begins with the ZOI shape. First, the center is detected by finding the maximum of the distance transform (green dot). The resulting point is then used to create a radius mapping. To generate the labels, the edge of the ZOI shape is extracted and a distance transform is applied. The pixel values are then rounded, mapped to a radius, and restricted to areas outside the plate and disk (scissors). Finally, the average colony intensity per label is calculated. Label images are visualized via the JIPipe 5.3.0 node “Labels to RGB”, where the Fire look up table (LUT) is used for per-radius labels, and the “Glasbey” LUT is used for all other labels.

### Automated Etest analysis with *J-AST*

*J-AST* allowed us to extend the principles behind *diskImageR*, which is limited to DDA analysis, to the quantitative analysis of Etests, for which the generation of the ring-shaped measurement labels needed to be adapted to follow the non-circular ZOI shape (**Fig. 3b**). Due to the nontrivial shape of the ZOI, the analysis of Etests requires an additional ROI-based annotation that highlights the approximate ZOI shape. This outline can be generated automatically or using interactive tools. As the ZOI may not be visible anymore in later time points, we additionally implemented an algorithm that registers the ZOI shape from the first time point to all others.

For Etests, the algorithm detects the ZOI centroid via distance-transform maxima, then generates measurement labels following the ZOI shape by applying a second distance transform to the ZOI outline, mapping each label to a radius from the centroid (**Fig. 3b**; **Supplementary Section 2.6**).

To validate Etest-based tolerance quantification, we analyzed a set of 19 clinical *C. albicans* isolates on two different growth media (YPD and SDC), using both DDAs and Etests in the timeline analysis mode. The FoG_50_ correlated strongly between the assay types (Pearson *r* ≈ 0. 837; **Fig. 4**) , demonstrating that *J-AST* enables tolerance quantification across both platforms.

**Fig. 4:**
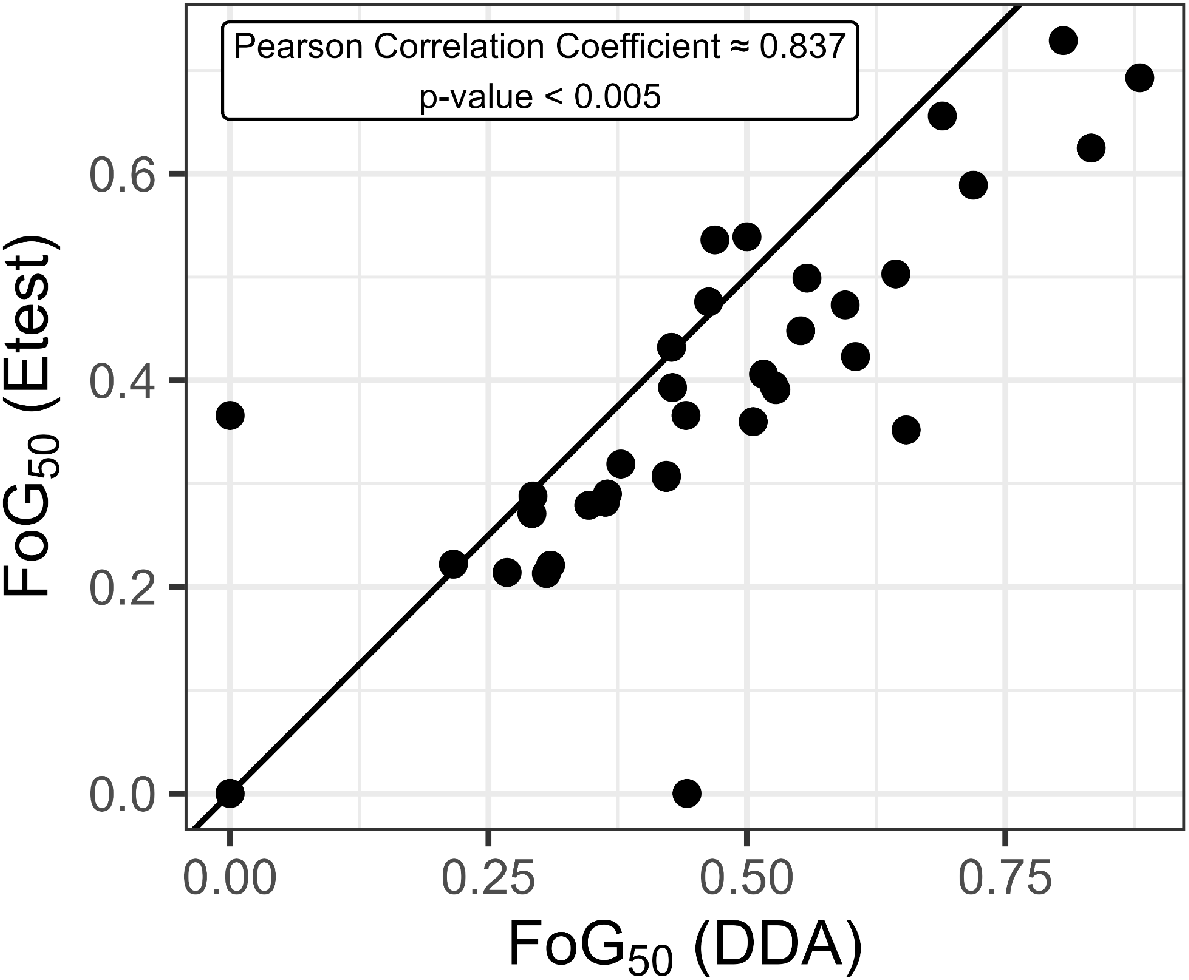
Comparison of tolerance quantification between DDAs and Etests. *J-AST* was used to quantify 38 ASTs performed in parallel with both DDAs and Etests in parallel, and quantified using the timeline mode. The tolerance (FoG_50_) values for corresponding DDAs and Etests show a strong linear correlation. The diagonal line is where x=y.

### Quantitative comparison between *J-AST* and *diskImageR*

We compared the RAD and FoG values produced by *diskImageR* and *J-AST* using both 20% and 50% thresholds for DDAs with 37 *C. albicans* strains grown on two different media (**Fig. 5a** and **5b**). Values correlate strongly, with the exception of late time point FoG at the 50% inhibition threshold (**Fig. 5b**). Due to extensive growth inside the ZOI, *J-AST* here is assigned a RAD value of 0 (hence, a FoG value of 0) whereas *diskImageR* detects small RAD values with a high FoG. Timeline analysis allows the carryover of RAD from an early time point to calculate FoG for a later time point, reflecting the temporal distinctions between resistance and tolerance when using a microbistatic drug ^4^. A brief comparison of features between *diskImageR* and *J-AST* is provided in **Supplementary Table 1**.

**Fig. 5:**
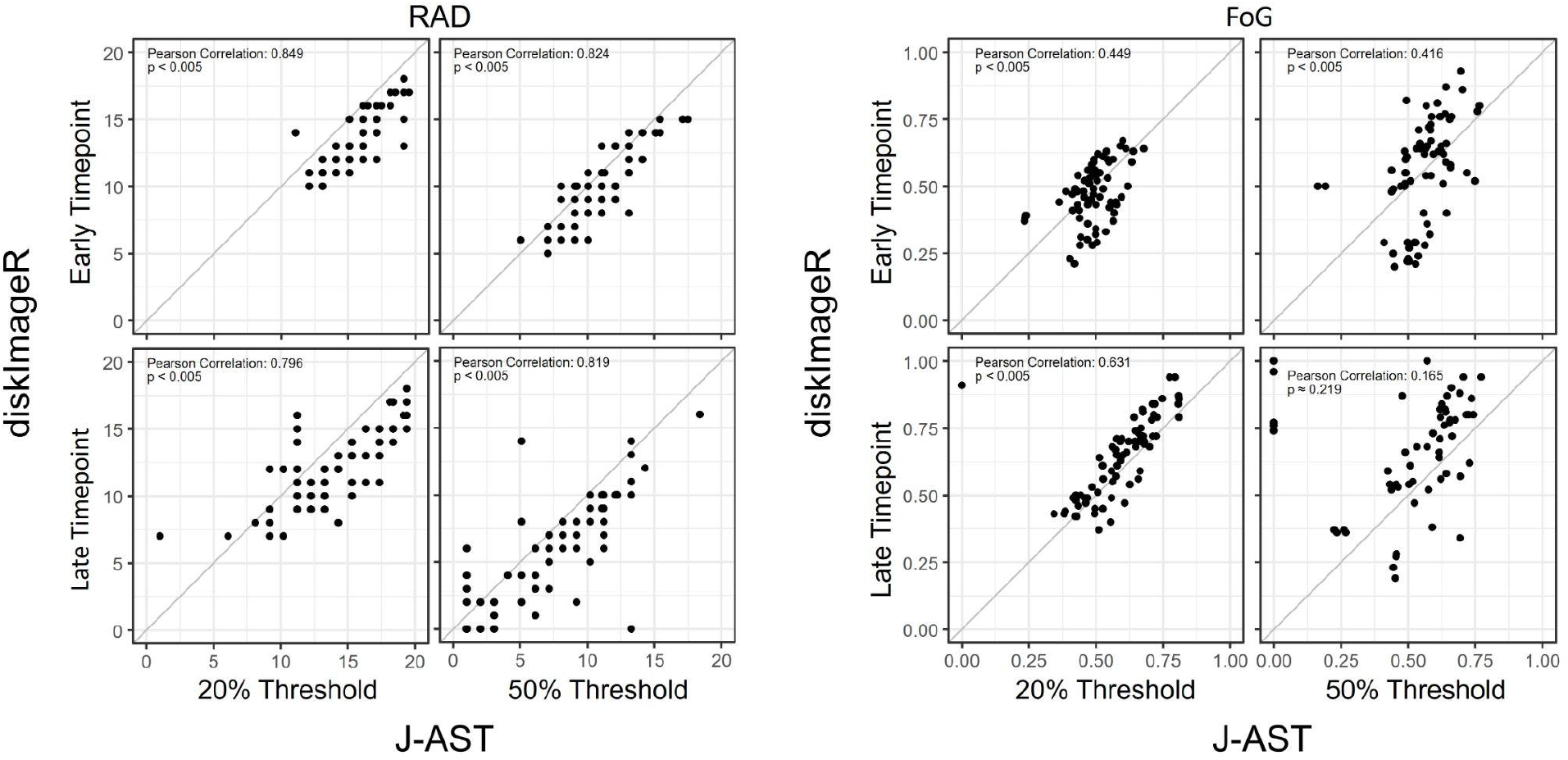
Comparison of *J-AST* and *diskImageR* quantification of **(a)** RAD and **(b)** FoG. n=74 Note that all quantifications largely correlate, with the exception of late time point FoG calculations using a 50% threshold. The diagonal line is where x=y.

### *J-AST* implements a fully automated algorithm for MIC detection

A common use of Etests is the readout of the minimum inhibitory concentration (MIC), which is usually done manually and thus subject to researcher bias. *J-AST* provides a fully automated algorithm that extracts the MIC given the physical pixel size, plate mask, strip mask, and strip layout.

The *J-AST* MIC detector applies an optical character recognition (OCR) workflow using *Tesseract* (https://github.com/tesseract-ocr/tesseract) to extract the exact tick positions (**Fig. 6a**). The vicinity around the strip is then segmented into concentration-associated cells for intensity extraction (**Fig. 6b**). The MIC is the highest concentration where the normalized growth passes a configurable threshold (default 25.5%) , with an optional module for detecting assay lacking a valid MIC. A full description of the algorithm is provided in **Supplementary Section 3**.

**Fig. 6:**
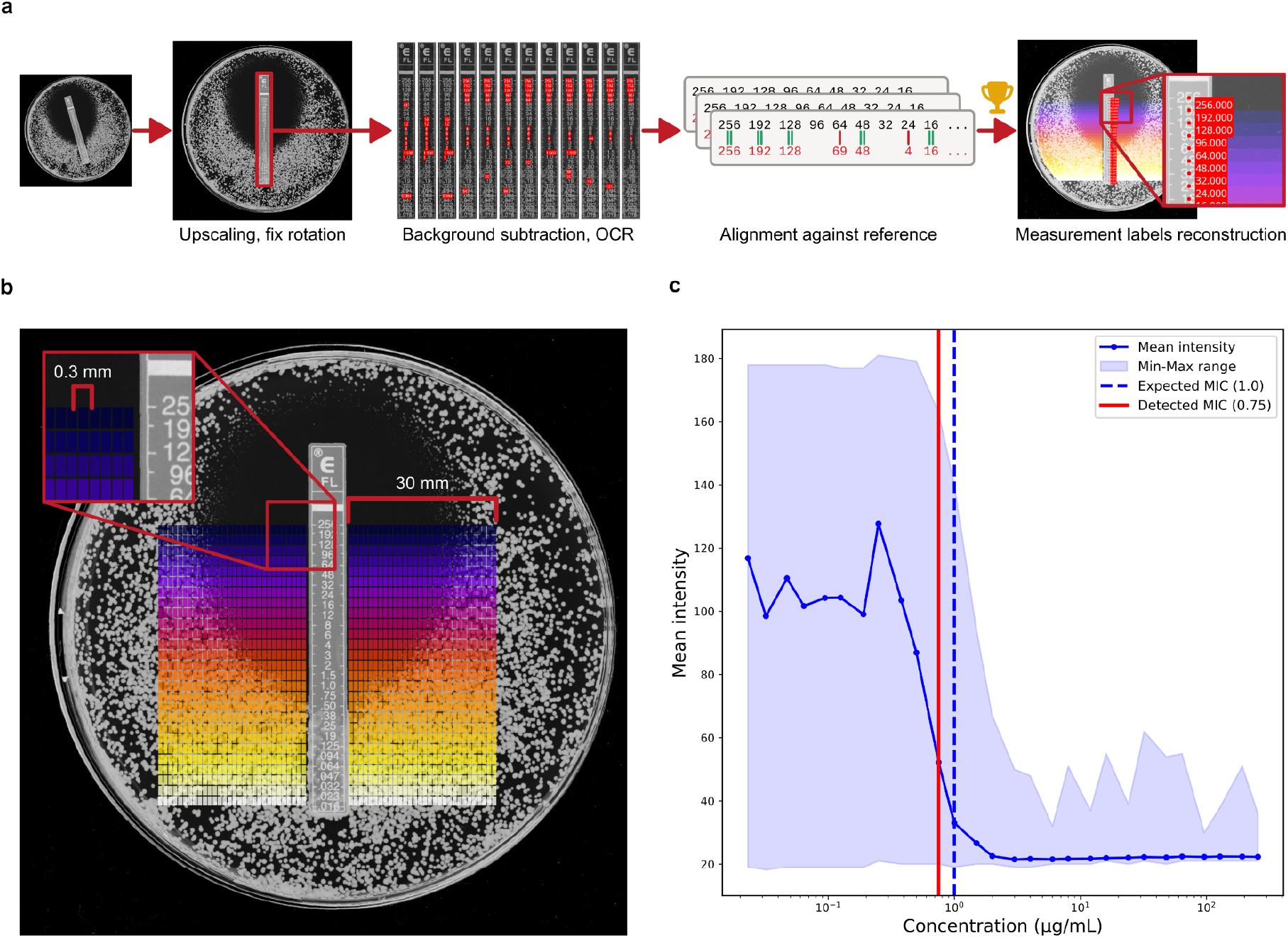
MIC detection algorithm implemented in *J-AST*. **(a)** The image is scaled up to 600 DPI and rotated to ensure that the strip is vertical. Then OCR is applied on different parameter sets for background correction. The best candidate is used to reconstruct the locations of all ticks, yielding measurement regions for each concentration. **(b)** Visualization of how the MIC detection algorithm segments the area around the strip into cells (colored squares) for the extraction of different measurements. **(c)** Extracted measurement curve of an image with the mean, minimum, and maximum intensity within the concentration-associated cells that are at most 3 mm away from the strip edge. The detected and expected MIC are marked as vertical lines.

The MIC detection algorithm achieved an essential agreement (EA) of 94.5%. Of 55 evaluable images 23 had exact matches and 29 assessments differed from manual evaluation by only one tick up or down (**Supplementary Figures 45** and **46**). The N/A MIC detection algorithm performed with a precision of 96.7%, a recall of 96.7%, specificity of 98.2%, F1 of 96.7%, balanced accuracy of 97.4%, and Cohen’s Kappa of 94.9% (**Supplementary Figure 47**). One image caused the OCR step to fail, as no candidate passed the Smith-Waterman alignment filter. The pipeline defaulted to reporting a MIC of N/A. Notably, this fallback result was consistent with the manual evaluation, which also classified the image as having no detectable MIC.

## Discussion

Clinical microbiology laboratories have a critical need^2,14–16^ to rapidly quantify antimicrobial susceptibility reliably and accurately. Basic research laboratories also need facile tools to study antimicrobial susceptibility, and enable the development of novel treatment strategies. We developed *J-AST*, an open-source platform, to provide seamless data integration with FAIR metadata management, automated analysis algorithms, and interactive data annotation. Importantly, *J-AST* is a modern web application, deployable as both desktop software and on servers; this minimizes the usage of third-party dependencies, and ensures easy maintenance of the software.

Assays on agar plates vary in image composition, quality, and differences in experimental setups, challenging the design of a fully automated analysis workflow. *J-AST* addresses this through configurable automated algorithms for preprocessing, ROI detection, and quantification, combined with interactive annotation tools available at any stage. Users can process images independently or as timelines and extend functionality via JIPipe plugins.

*J-AST* improves on *diskImageR* in two ways: first, the *JIPipe* integration eliminates manual installation of *R, ImageJ*, and dependencies, improving portability. Second, ring-shaped measurement labels segment the entire plate (excluding disk and plate edge) rather than 72 radial profiles, trading slightly lower radial resolution (1 mm) for more robust intensity estimation across all available pixels. This label-based approach also enabled Etest adaptation, though the non-circular, drug-dependent ZOI shapes required the introduction of a user-friendly shape-based solution (**Supplementary Section 2.5**). Here, the algorithm then applies a distance transform to estimate labels that approximately follow the underlying concentration gradient.

RAD and FoG values from *J-AST* agree with *diskImageR*. The one exception was at the extreme margins and later time point where the tools differed in interpretation of the RAD (**Fig. 5**). This is circumvented by timeline analysis using early-timepoint RAD, provided at least one image has a visible ZOI.

By quantifying tolerance within the ZOI of Etest experiments, *J-AST* expands the utility of *diskImageR* functionality. FoG quantification between DDAs and Etests have a strong linear correlation (**Fig. 6**). Users comparing DDA and Etest assays regularly should consider using a calibration curve comparing the two. Nonetheless, *J-AST* is fully capable of quantifying FoG on Etest assays and fills an unfulfilled niche in open-access AST analysis.

Additional benefits of the implementation of the *J-AST* platform in *JIPipe* include the generation of measurements (e.g., per-image FoG, and their difference in given a time series) not captured by other tools, and a single image RAD and FoG quantification that may be of interest for future studies. *J-AST* also includes a new algorithm for MIC detection, which performs well on Etests where the MIC can be easily identified manually, and was designed to provide easily explainable parameters based on physical size expectations, image contrast, and percentage thresholds. Given that the exact choice of the MIC can be highly subjective, manual readouts are prone to intra-rater variability, where the same researcher may assign different MIC values to the same image upon re-evaluation. This is highlighted in the discrepancy between detected and expected MIC in **Fig. 6c** where a manual re-assessment may agree with the automated algorithm. *J-AST* thus also contributes towards reducing researcher bias in the field by ensuring consistent application of objective criteria.

While *J-AST* is focused on providing a semi-automatic analysis workflow, the underlying algorithms can be interfaced with workflow managers like *Galaxy* ^17^, *Nextflow* ^18^, or *Snakemake* ^19^. New algorithms can be added via the *Java* API or plugin system, and the interactive platform is adaptable to other image analysis tasks combining automated and manual annotation.

In summary, *J-AST* is a state-of-the-art tool with broad applications in infection biology, capable of quantifying multiple facets of two widely used susceptibility assays. Thus, *J-AST* has the potential to improve both the basic research of drug susceptibility as well as the interpretation of clinical tests.

## Methods

### Software architecture and design

*J-AST* is split into two components (**Supplementary Figure 43**): the Java-based backend that is responsible for image, metadata, and results storage, as well as for managing users and executing image analysis workflows. The frontend is specialized on displaying information obtained from the backend through a Representational State Transfer (REST) ^20^ interface, and sending requests to the backend, for example, uploading images, updating metadata, or starting an image analysis task.

The project itself uses the *Maven* build system (Apache Software Foundation) and has two submodules, the backend built on *Spring Boot* (Broadcom Inc.), and the frontend built on *Node*.*JS* (OpenJS Foundation), allowing to build both components easily using one project management system.

#### Backend

We implemented the backend using *Spring Boot* 3.3.4 (Broadcom Inc.) on *Java* 17 (Oracle Corporation). Our tool ships with *JIPipe* 5.3.0, which is managed through *JobRunr* 7.4.0 (Rosoco BV) and its associated *Spring Boot* library. The list of all dependencies can be found in **Supplementary Table 2**. Our tool is designed to be deployed with *MariaDB* (MariaDB Foundation) for usage on a server, but uses a local self-contained *H2* database (www.h2database.com) when packaged as a desktop application. Due to known limitations of database servers regarding the storage of large binary objects (BLOBs), i.e., images, annotations, and results, we implemented a custom Universally Unique Identifier (UUID) –based Binary Large Object (BLOB) storage service. This design allows to limit the number of external dependencies to a minimum, ensuring that *J-AST* can be packaged and installed even on *Windows* PCs without administrative privileges.

The backend features a powerful component-based API to organize and deploy computationally expensive image processing tasks: to add a new task, developers just have to inherit from a dedicated “Task type” class that encapsulates all required metadata and workloads, including the unique ID, name, description, supported assay types, parameters and their default values, and if a task is executed on a single-image or timeline-level. The backend automatically collects task types and offers REST API endpoints that communicate all relevant information for the frontend.

As an additional option for extending *J-AST*, we also provide a plugin API that automatically analyzes *JIPipe* workflows that are placed inside a specific directory. Given the required metadata is included within the pipeline using *JIPipe*, our tool then automatically handles all required underlying operations without the need to change any code and recompile the *J-AST* backend.

The backend offers a variety of REST API endpoints responsible for handling user authentication and registration, project management, image upload and organization, metadata management, access to results, as well as the scheduling and monitoring of tasks.

#### Frontend

The frontend component is implemented in *Typescript* (Microsoft Corporation) using *Vue*.*js* 3.4.18 (https://vuejs.org/) and the *Quasar* 2.16.0 (PULSARDEV SRL) user interface library. Packages are managed using NPM. The frontend features an interactive image mask editor implemented using *Vue-Konva* 3.1.2 (https://konvajs.org/) and *Q-Floodfill* 1.3.1 (https://github.com/pavelkukov/q-floodfill). The desktop application uses the *Electron* (https://www.electronjs.org/) integration provided by *Quasar*. The list of all dependencies can be found in **Supplementary Table 3**.

The frontend makes a distinction between two image processing task types. Lightweight frontend tasks are executed exclusively on the frontend and are used for organizational purposes, including the automated extraction of text metadata, and image sorting. Computationally intensive tasks, including image processing and analysis, are handled by the backend. During the startup of the single page application (SPA), the frontend queries a dedicated REST API endpoint and processes the resulting information to automatically generate the necessary user interfaces, and apply basic preliminary checks prior to sending tasks to the backend via a payload object.

#### Image analysis

All image analysis operations were implemented in *JIPipe* 5.3.0 using the official *ImageJ* distribution that is shipped with *JIPipe*. Our *diskImageR*-reimplementation uses the *JIPipe*-provided *R* environment 4.4.0.1002 to execute *R* code.

### Benchmarking setup and evaluation datasets

#### Strains and growth conditions

The *C. albicans* strains utilized in this study are listed in **Supplementary Table 4** of the supplemental material. Strains were stored in 15% glycerol at -80°C and were subcultured on YPD agar (1% yeast extract, 2% peptone, 2% glucose, 2% agar) at 30°C.

#### Drug Diffusion Assays

Both DDAs and Etests were used to assay antifungal susceptibility. To this end, several colonies which had previously grown on YPD agar at 30°C for 48 hours were suspended in sterile water. Samples were diluted to an OD_600_ of 0.03 and 100µL was spread on either YPD or SDC (2% glucose, 6.7g/L Yeast Nitrogen Base (Difco), and 2g/L complete amino acid supplement (MP Biomedicals)) using glass beads. For DDAs, a single filter paper disk containing 25µg fluconazole (Oxoid, UK) was placed in the center of the plate using forceps. For Etests, a strip containing a gradient of drug (bioMérieux) was placed in the middle of the plate using forceps. Plates were incubated at either 30 or 37°C for 48 hours, and photographs were taken at 24 and 48 hours using a Phenobooth Plus (Singer Instruments, UK). For comparison, DDAs were quantified with both *J-AST* and *diskImageR* version 1.0.0 ^6^. Outputs were compared using Pearson correlation.

#### MIC detection

The MIC detection algorithm was tested on a set of 86 Etest images where the MIC was evaluated manually. A subset of 30 images was identified to have no MIC by an expert. We executed the *J-AST* workflow on the full Etest image data set and used a *Python* script to compare the performance to the human assessment. For the detection of images without a MIC we calculated the precision, recall, specificity, F1, balanced accuracy, and Cohen’s Kappa. If both our algorithm and the manual assessment detected a MIC we compared the difference in ticks of the MIC scale. Differences of up to ± 1 ticks of the MIC scale were considered to be acceptable. Based on this evaluation, we calculated the essential agreement 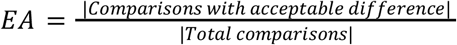. In cases where the optical character recognition (OCR) fails the algorithm assumes that there is no MIC.

### Installation and deployment options

*J-AST* was designed to be used both as a web application, and as desktop software. This allows the deployment of our tool as a cloud service for multiple users, and as an offline and local desktop application.

#### Web-based installation

To deploy *J-AST* as a web application on a server, we provide pre-made *Docker Compose* (Docker Inc.) configurations. Due to the low number of third-party service dependencies, server administrators can also easily deploy our tool without *Docker*. Here, only an existing installation of *MariaDB* is required.

#### Electron desktop application

For the installation on a desktop computer, we provide pre-packaged software packages for *Windows, Linux*, and *macOS*. These bundles include all necessary dependencies and do not require the installation of any additional components, thus making the software easily usable without administrator privileges.

## Data availability

Supplementary data has been uploaded to https://doi.org/10.5281/zenodo.20627415.

## Code availability

*J-AST* is open source and licensed under the MIT license. All code can be accessed through our GitHub repository: https://github.com/applied-systems-biology/j-ast/.

## Acknowledgements

R.G. is funded by the Deutsche Forschungsgemeinschaft (DFG, German Research Foundation) under the National Research Data Infrastructure NFDI4Bioimage – NFDI 46/1 – 50186459. D.K. is funded by the Deutsche Forschungsgemeinschaft (DFG, German Research Foundation) within Germany’s Excellence Strategy – EXC 2051 – Project-ID 390713860 and within CRC 1768 “VirusREvolution: Decoding tools for virus research” – Project number 553944903. A.M. and J.B. received funding from the European Research Council (ERC) under the European Union’s Horizon 2020 research and innovation program (grant agreement No 951475).

## Author contributions

R.G. developed the *J-AST* software and all analysis algorithms, implemented the MIC detection pipeline, performed the formal analyses comparing *J-AST* with *diskImageR*, and wrote the initial manuscript draft. A.M. performed all experiments, conducted the DDA and Etest comparisons, analyzed the tolerance correlation data, and contributed to writing the experimental sections. D.K. contributed to the development of the OCR algorithm used for MIC detection. J.B. conceptualized the biological study, supervised the experimental work, acquired funding, and edited the manuscript. M.T.F. conceptualized the software study, supervised the project, acquired funding, and edited the manuscript. All authors reviewed and approved the final manuscript.

## Competing interests

No competing interests are declared.

## Notes

https://doi.org/10.5281/zenodo.20627415

https://github.com/applied-systems-biology/j-ast/

### Competing Interest Statement

The authors have declared no competing interest.

## References

1. Naghavi, M. et al. Global burden of bacterial antimicrobial resistance 1990–2021: a systematic analysis with forecasts to 2050. The Lancet 404, 1199–1226 (2024).

2. Pfaller, M. A., Diekema, D. J., Turnidge, J. D., Castanheira, M. & Jones, R. N. Twenty Years of the SENTRY Antifungal Surveillance Program: Results for Candida Species From 1997–2016. Open Forum Infect. Dis. 6, S79–S94 (2019).

3. Brauner, A., Fridman, O., Gefen, O. & Balaban, N. Q. Distinguishing between resistance, tolerance and persistence to antibiotic treatment. Nat. Rev. Microbiol. 14, 320–330 (2016).

4. Berman, J. & Krysan, D. J. Drug resistance and tolerance in fungi. Nat. Rev. Microbiol. 18, 319–331 (2020).

5. Pfaller, M. A. et al. Comparison of European Committee on Antimicrobial Susceptibility Testing (EUCAST) and Etest Methods with the CLSI Broth Microdilution Method for Echinocandin Susceptibility Testing of Candida Species. J. Clin. Microbiol. 48, 1592–1599 (2010).

6. Gerstein, A. C., Rosenberg, A., Hecht, I. & Berman, J. diskImageR: quantification of resistance and tolerance to antimicrobial drugs using disk diffusion assays. Microbiology 162, 1059–1068 (2016).

7. Rueden, C. T. et al. ImageJ2: ImageJ for the next generation of scientific image data. BMC Bioinformatics 18, 529 (2017).

8. Schindelin, J. et al. Fiji: an open-source platform for biological-image analysis. Nat. Methods 9, 676–682 (2012).

9. Mackey, A. I. et al. Aneuploidy confers a unique transcriptional and phenotypic profile to Candida albicans. Nat. Commun. 16, 3287 (2025).

10. Alabi, P. E. et al. Small molecules restore azole activity against drug-tolerant and drug-resistant Candida isolates. mBio 14, e00479–23 (2023).

11. Ostrosky-Zeichner, L. et al. Rationale for Reading Fluconazole MICs at 24 Hours Rather than 48 Hours When Testing Candida spp. by the CLSI M27-A2 Standard Method. Antimicrob. Agents Chemother. 52, 4175–4177 (2008).

12. Wilkinson, M. D. et al. The FAIR Guiding Principles for scientific data management and stewardship. Sci. Data 3, 160018 (2016).

13. Gerst, R., Cseresnyés, Z. & Figge, M. T. JIPipe: visual batch processing for ImageJ. Nat. Methods 20, 168–169 (2023).

14. Rosenberg, A. et al. Antifungal tolerance is a subpopulation effect distinct from resistance and is associated with persistent candidemia. Nat. Commun. 9, 2470 (2018).

15. Levinson, T. et al. Impact of tolerance to fluconazole on treatment response in Candida albicans bloodstream infection. Mycoses 64, 78–85 (2021).

16. Gomez, J. E. & McKinney, J. D. M. tuberculosis persistence, latency, and drug tolerance. Tuberculosis 84, 29–44 (2004).

17. Afgan, E. et al. The Galaxy platform for accessible, reproducible and collaborative biomedical analyses: 2016 update. Nucleic Acids Res. 44, W3–W10 (2016).

18. Di Tommaso, P. et al. Nextflow enables reproducible computational workflows. Nat. Biotechnol. 35, 316–319 (2017).

19. Mölder, F. et al. Sustainable data analysis with Snakemake. F1000Research 10, 33 (2025).

20. Fielding, R. T. Architectural Styles and the Design of Network-Based Software Architectures. (University of California, Irvine, 2000).

