## Supplemental information for "*J-AST*: a web-based analysis platform for antimicrobial susceptibility testing"

1 J-AST user interface 3
- 1.1 Project creation and data organization 3
- 1.2 Annotating regions of interest 5
  - 1.2.1 Plate detection 6
  - 1.2.2 Pixel size calibration 6
  - 1.2.3 DDA disk detection 6
  - 1.2.4 Etest strip detection 7
  - 1.2.5 Etest ZOI shape detection 7
  - 1.2.6 Manual image annotation 7
- 1.3 Browsing results 8

2 Image analysis algorithms 9
- 2.1 Plate detection 9
  - 2.1.1 Slow algorithm 9
  - 2.1.2 Fast algorithm 14
- 2.2 Pixel size calibration 18
- 2.3 DDA disk detection 21
  - 2.3.1 Variant 1 (Slow) 21
  - 2.3.2 Variant 1 (Fast) 27
  - 2.3.3 Variant 2 33
  - 2.3.4 Variant 3 39
- 2.4 Etest strip detection 47
- 2.5 Etest ZOI shape detection 60
- 2.6 DiskImageR re-implementation 69
    - 2.6.1.1 Compartment C1 "Data handling" 70
    - 2.6.1.2 Compartment C2 "Align ZOI Shape" 75
    - 2.6.1.3 Compartment C3 "Generate labels" 79
    - 2.6.1.4 Compartment C4 "Raw statistics" 85

2.6.1.5 Compartment C5 "Statistics" 90
2.6.1.6 Compartment C6 "Save Visualizations" 100
2.6.1.7 Compartment C7 "Save tables" 107
2.7 DiskImageR re-implementation (single image) 113
2.7.1.1 Compartment C1 "Data handling" 113
2.7.1.2 Compartment C2 "Align ZOI Shape" 116
2.7.1.3 Compartment C3 "Generate labels" 117
2.7.1.4 Compartment C4 "Raw statistics" 122
2.7.1.5 Compartment C5 "Statistics" 126
2.7.1.6 Compartment C6 "Save Visualizations" 134
2.7.1.7 Compartment C7 "Save tables" 140
3 MIC detection algorithm 146
Project-wide parameters 147
Compartment C1 “Data handling” 148
Compartment C2 “Correct rotation” 151
Compartment C3 “OCR” 156
Compartment C4 “Labels” 166
Compartment C5 “Visualize strip” 168
Compartment C6 “Measure” 171
Compartment C7 “Visualize measurements” 212
4 J-AST application 217
4.1 Backend 218
4.2 Frontend 219
5 Benchmarking setup and evaluation datasets 221
6 MIC detection results 222

### 1 J-AST user interface

#### 1.1 Project creation and data organization

After logging into the cloud-based service or starting the local desktop application, users are presented with the option to open or create a new project (see **Supplementary Figure 1**). On creating a project, the name and default organization method can be selected: “Single images” for projects where each image should be analyzed independently; or “Timeline” for analyses where multiple time points need to be arranged (e.g., 24 h and 48 h images for determining RAD and FoG). At any time after the project has been created, the project editor can be used to alter parameters (e.g., timing of images).

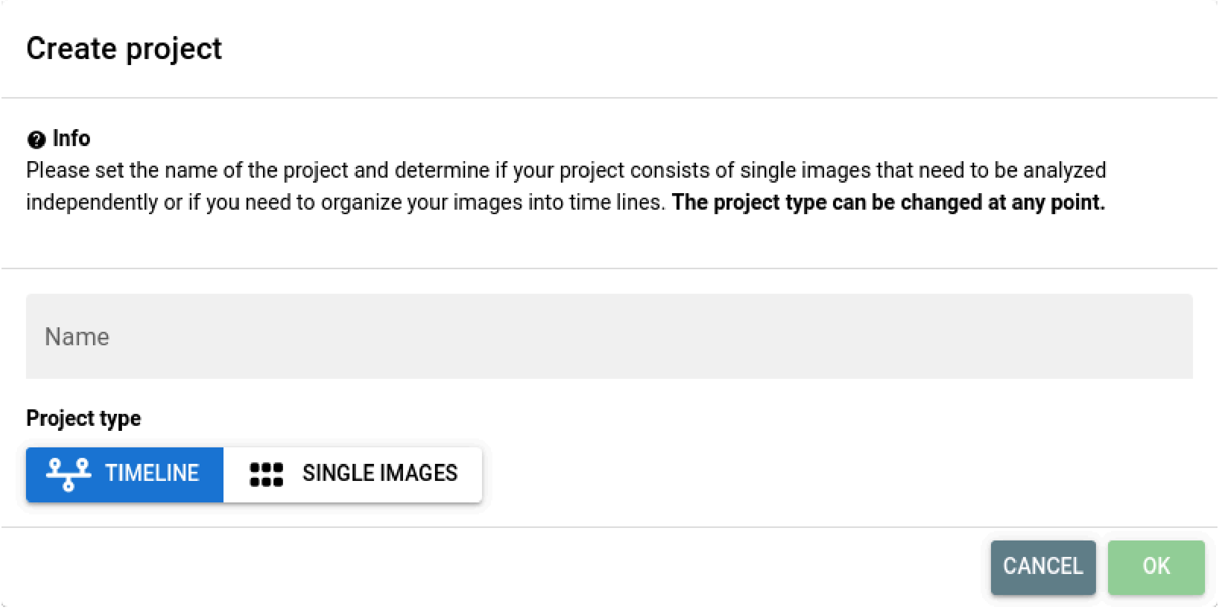

**Supplementary Figure 1**: Project creation dialog. Users can set the name and the project type. A timeline project allows organizing the image into a time series, while the “single images” type enables the independent per-image analysis.

The “Upload” tool is used to upload PNG images into J-AST either in an empty list or a table of images. Images also can be uploaded multiple times, as the data management system uses unique database identifiers to handle images. In the case of a “Timeline” project, images are first placed within a dedicated “Unsorted images” list that gives users the option to modify and review the image metadata prior to sorting them into timelines. No additional sorting is required if the project type is “Single images”, i.e. the mode for single-image analyses.

Independent of the project type, users are expected to fill out the required metadata fields “Experiment”, “Sample”, and “Assay Type”. For timeline projects, additional “Time point” information is required. While all metadata can be filled in manually by selecting the image and typing the information, J-AST also provides convenient tools to automatically extract information from the file name. This feature is accessed by selecting one or multiple images, navigating to the “Process” tab, and choosing the “Auto-fill metadata” option. Users are provided with an intuitive interface that determines how file names are split, and which components are mapped to which metadata field. The tool also allows setting a static value for a field if the information is not part of the file name. If all metadata is filled out, it can be used for organizing, analyzing, and searching through data.

For organizing images into timelines, J-AST again provides both interactive and automated solutions that can be freely combined. To create a timeline manually, users merely need to drag images from the “Unsorted” list into a table where columns represent time points and rows represent timelines. Here the interface further aids researchers by highlighting the relevant metadata for each row and column. Another use of the metadata is the fully-automated sorting process that can be triggered by selecting the unsorted images, and navigating to the “Process” tab and its “Auto-sort by metadata” item. The tool analyzes the available time points and queries users to input the order if needed. Then all selected images are arranged fully automatically. Again, users can apply manual changes interactively.

J-AST simplifies both the manual and automated annotation of Etests with their MIC. To aid with the interactive process, projects can be conveniently switched into a “Browser” mode (see **Supplementary Figure 2**) where only one image is displayed in full screen together with controls for editing metadata and navigating through the project’s images. Users then can go through each individual image and manually review and modify the MIC and other metadata.

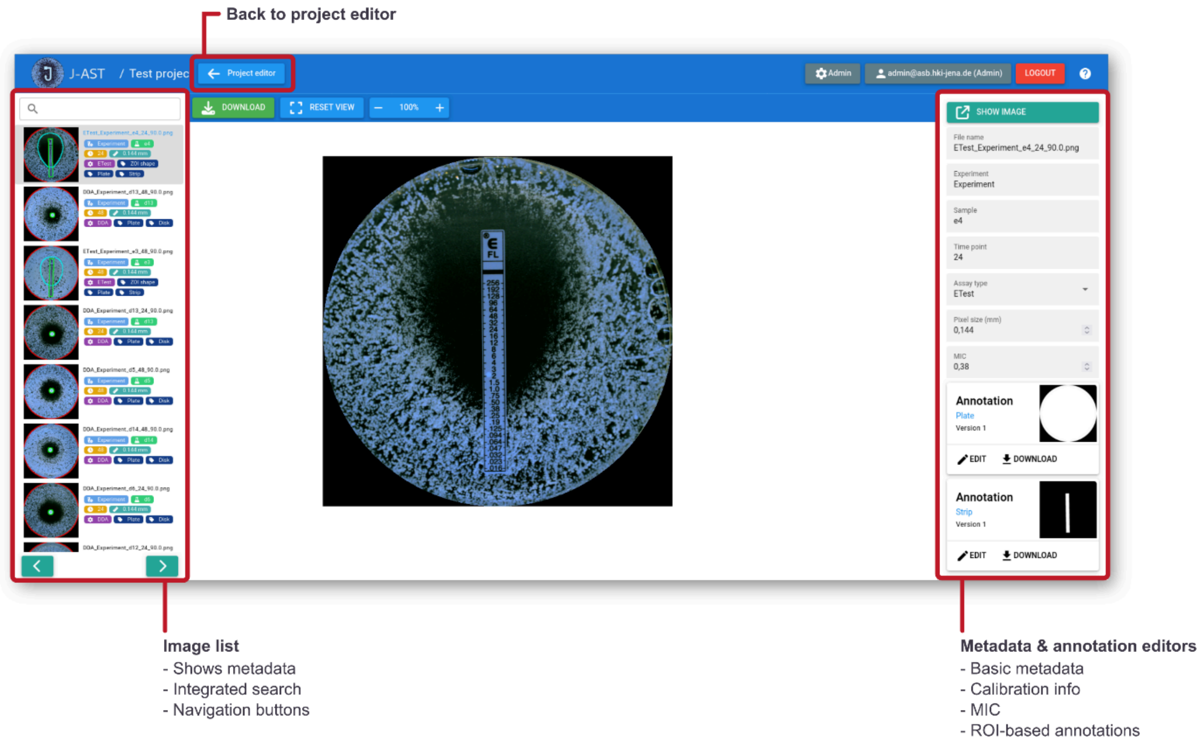

**Supplementary Figure 2:** Browser mode. J-AST features a dedicated view that easily allows one-by-one review and annotation of the uploaded images. On the left-hand side there is a list of all images with their associated metadata. The image list can be filtered via a search bar and conveniently navigated through by clicking navigation buttons that go to the next or previous image. The mode can be at any point exited by clicking the “Project editor” button.

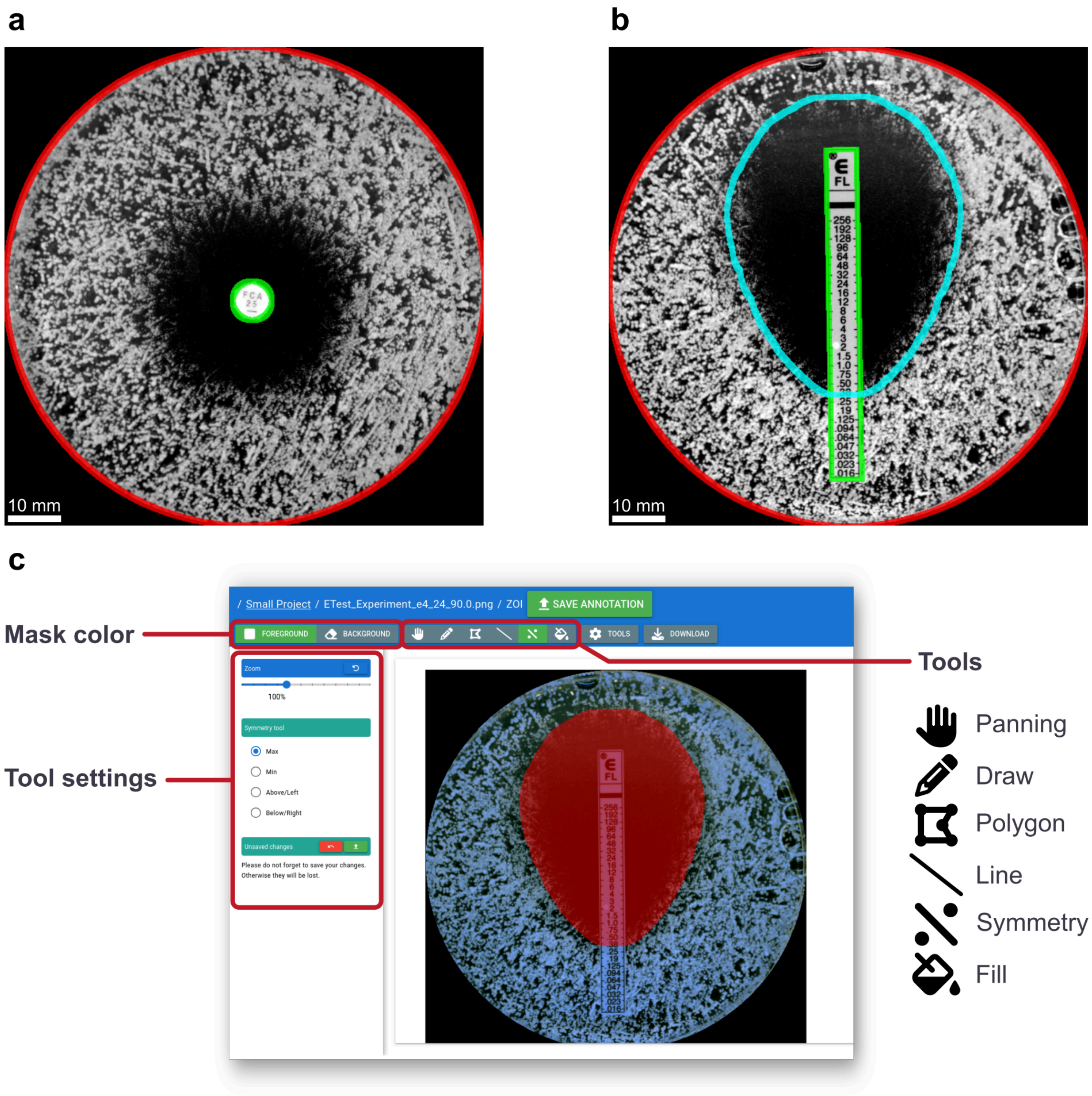

**Supplementary Figure 3**: Example of ROI-based annotations and the interactive editor implemented in *J-AST*. **(a)** Example of a fully annotated DDA. The plate is outlined in red and the disk is highlighted in green. **(b)** Example of a fully annotated Etest where the plate and the strip are highlighted in red and green respectively, and the ZOI shape is shown in blue. **(c)** Screenshot of the built-in interactive ROI-based annotation editor. Several available tools allow users to manually highlight the annotation region, including free-hand drawing, polygon, and line operations. These tools can be configured through controls for changing the drawn color and tool-specific settings.

#### 1.2 Annotating regions of interest

Depending on the algorithm, specific image regions need to be annotated before analysis. J-AST automatically applies validation checks before running an operation and informs users about missing metadata and annotations through a graphical interface. J-AST supports the following region of interest (ROI) annotations, depending on the assay type: the plate area and its associated pixel size metadata, the drug disk in DDAs, the drug strip in Etests, and the ZOI of an Etest (see **Supplementary figures. 3a** and **3b**).

J-AST provides both interactive and fully automated methods for creating, reviewing, and modifying ROI annotations. Automatically generated ROIs can be edited at any time through the integrated interactive editor, enabling a workflow where the majority of annotation tasks can be automated, while manual modifications can be used to correct mistakes such as misidentification of plate boundaries, disk location, or strip shape. As J-AST provides multiple automated algorithms that each approach the analysis from different image features and involve different processing steps, this greatly enhances the flexibility of our software and makes it usable for a wider audience.

##### 1.2.1 Plate detection

The fully automated plate detection algorithm is applied by selecting the images, and clicking the “Process” menu. Here, the “Auto-detect plate” operation can be found within the “Plate” category. Currently, there are two algorithms that are available through the same command: the slow algorithm tests all known ImageJ auto thresholding methods and selects the most circular object that covers by default at least 40% of the image. A faster algorithm that avoids testing all thresholding methods uses a different approach where the contrast between the inside of the plate and the background is first enhanced by applying a local maximum operation, which is followed by an Otsu thresholding operation. An ellipse is fit to the resulting mask to obtain a smooth plate border. Independent of the algorithm implementation, the generated ROI mask is automatically stored inside a dedicated “Plate” annotation field that is associated with the image. The frontend then updates its visualization to allow users to see if the operation succeeded. A detailed description of both algorithms can be found in **Supplementary section 2.1**. All default parameters can be changed within the J-AST interface.

##### 1.2.2 Pixel size calibration

Many AST images are stored in common image storage formats like JPEG or PNG that lack information about the physical pixel size. This introduces a difficulty in calibrating the parameters of the image processing and analysis steps. J-AST solves this problem by requiring an additional calibration step that uses the known size of plates – usually 90 mm – to estimate the physical pixel size, given an existing plate annotation, and is applied automatically after the plate detection algorithm. The resulting pixel size in millimeters is then written into the associated image metadata field and is utilized by J-AST operations to calibrate the parameters of the underlying image processing algorithms.
The calibration step also can be executed alone by selecting the images, clicking the “Process” button, and running the “Calibrate pixel size by plate” operation within the “Plate” category. Users can also manually input the pixel size via the metadata editor. A detailed description of the automated plate calibration algorithm can be found in **Supplementary section 2.2**.

##### 1.2.3 DDA disk detection

Automated detection of the DDA disk is affected by different lighting and image contrast conditions. To cover a broad range of images, J-AST currently provides three different algorithm variants that can be accessed by selecting the images, clicking “Process”, and navigating to the “DDA” category.
The first algorithm variant assumes that the disk is bright compared to the colonies. It starts by removing the area outside the plate, and then testing a range of ImageJ auto-thresholding operations. For each thresholding method, the detected objects are filtered to ensure that with the default settings the disk candidates have a diameter ranging from 2 mm to 12 mm, and a minimum circularity of 50%. From all candidates, the most circular object is returned. For the first variant, there is an optional setting that accelerates the operation by assuming that the disk is located near the plate’s center. Here, the plot ROI is first downscaled to 25% of its original size and used to crop the image prior to evaluating the thresholding methods. A detailed description can be found in **Supplementary sections 2.3.1 and 2.3.2**.

The second disk detection algorithm assumes that the ZOI has a low intensity variance. It starts by applying a local variance filter with the radius set to 16th of the expected disk diameter of by default 6 mm. The resulting image is binarized using ImageJ’s “Default” auto-thresholding method, yielding a mask where the disk is separated from the colonies by a dark ring. To select the disk, the minimum of the local variance image is detected and used as a key point for picking the central disk region. Finally, the resulting object is then further processed by a circle fitting algorithm. A detailed description can be found in **Supplementary section 2.3.3**.

The third algorithm variant combines concepts of the two other options and applies both an intensity-based thresholding and a detection of the disk using local variance. Finally, the candidate that has a diameter closest to the expected value of by default 8 mm is selected. A detailed description of all algorithms can be found in the **Supplementary section 2.3.4**. All default parameters can be changed within the J-AST interface.

##### 1.2.4 Etest strip detection

J-AST detects Etest strips using an algorithm designed for assay variants where the whole strip is surrounded by a black and high-contrast line. The algorithm starts with a morphological gradient operation to enhance such lines. An ImageJ ridge detector is then used to find the line segments. Next, the two major sides of the strip are detected by applying Hough line detection, combined into one object, and fitted with a rotated rectangle operation, thereby yielding the vertical strip. To find the two minor sides, the gradient image is thresholded with respect to statistics obtained from within the vertical strip. This is followed by a thresholding and object detection, as well as a per-object convex hull operator. The algorithm then produces multiple candidate strips by iteratively assembling the top N largest parts into one ROI and applying a rotated rectangle fitting. From all candidates, the one closest to the expected aspect ratio of 11 by default is selected, as this is the ratio of the Etests in our sample data. All details of the algorithm are described in **Supplementary section 2.4**.

##### 1.2.5 Etest ZOI shape detection

For Etest strips with different antimicrobial drugs, the ZOI shape differs, likely as a function of their diffusion rates and concentration gradients. First, both the strip and the background are erased, then all available ImageJ thresholding methods are tested. The resulting masks are stacked into a 3D stack and Z-projected using a median filter to obtain a majority-vote thresholding. The JIPipe filament toolkit creates two sets of key points: the first set is calculated by finding the edges around the mask, and converting them into a graph. All vertices with degree zero are deleted before removing all edges, yielding the colony vertex set. The second set is generated by converting the Etest strip mask into a ROI and converting it into a graph. JIPipe is then instructed to split edges until the maximum distance between two vertices is at most 10 pixels. The edges are deleted, yielding the strip vertex set. The workflow then connects each strip vertex to the closest colony vertex while ensuring that only one connection is created. Then all vertices except colony vertices with at least one connection are deleted, and converted into ImageJ point ROIs. Finally, the workflow calculates the concave hull over all collected points and applies a mirror operation to make the shape symmetric. A detailed description of the algorithm can be found in **Supplementary section 2.5**.

##### 1.2.6 Manual image annotation

The interactive editor in J-AST supports the creation, review, and modification of ROI-based annotations (see **Supplementary Figure 3c**). The tool can be accessed at any point by selecting the image and clicking the “Edit” button associated with the annotation. This will open a full-screen dialog where users have access to drawing tools used for marking pixels as background or foreground. The set of interactive tools include a paint brush, polygon filling tool, line tool, flood-filling operation, and a mirror tool designed specifically for creating symmetric shapes by mirroring the mask over a hand-drawn axis. At any point, the user can reset the ROI to its original state, download the currently displayed mask as a PNG, and clear the mask.

When the user is finished drawing the ROI, the changes can be uploaded by clicking the “Save Annotation” button. This will close the dialog, returning the user to the list of images.

#### 1.3 Browsing results

J-AST features a convenient results browser (see **Supplementary Figure 4**) that supports complex hierarchies of measurements, plots, visualizations, and other outputs. Projects can store multiple results packages that are annotated with a name that is set during the setup of an analysis. The generated folders are then placed within a dedicated view that can be accessed through a “Results” button. Our interface allows users to browse through results just as a file system on a local computer or cloud drive, view images and tables without downloading them, and download the entire results dataset or specific files or directories.

Currently, J-AST provides four algorithms that utilize the results system: (1) a visualization algorithm that draws outlines of the available ROI annotations, i.e. for the plate, DDA disk, Etest strip, and ZOI shape, on the raw image, (2) our re-implementation of DiskImageR that exports multiple result tables, the measurement labels, plots, visualizations, and (3) the single-image variant of the DiskImageR algorithm that generates similar results to the timeline variant.

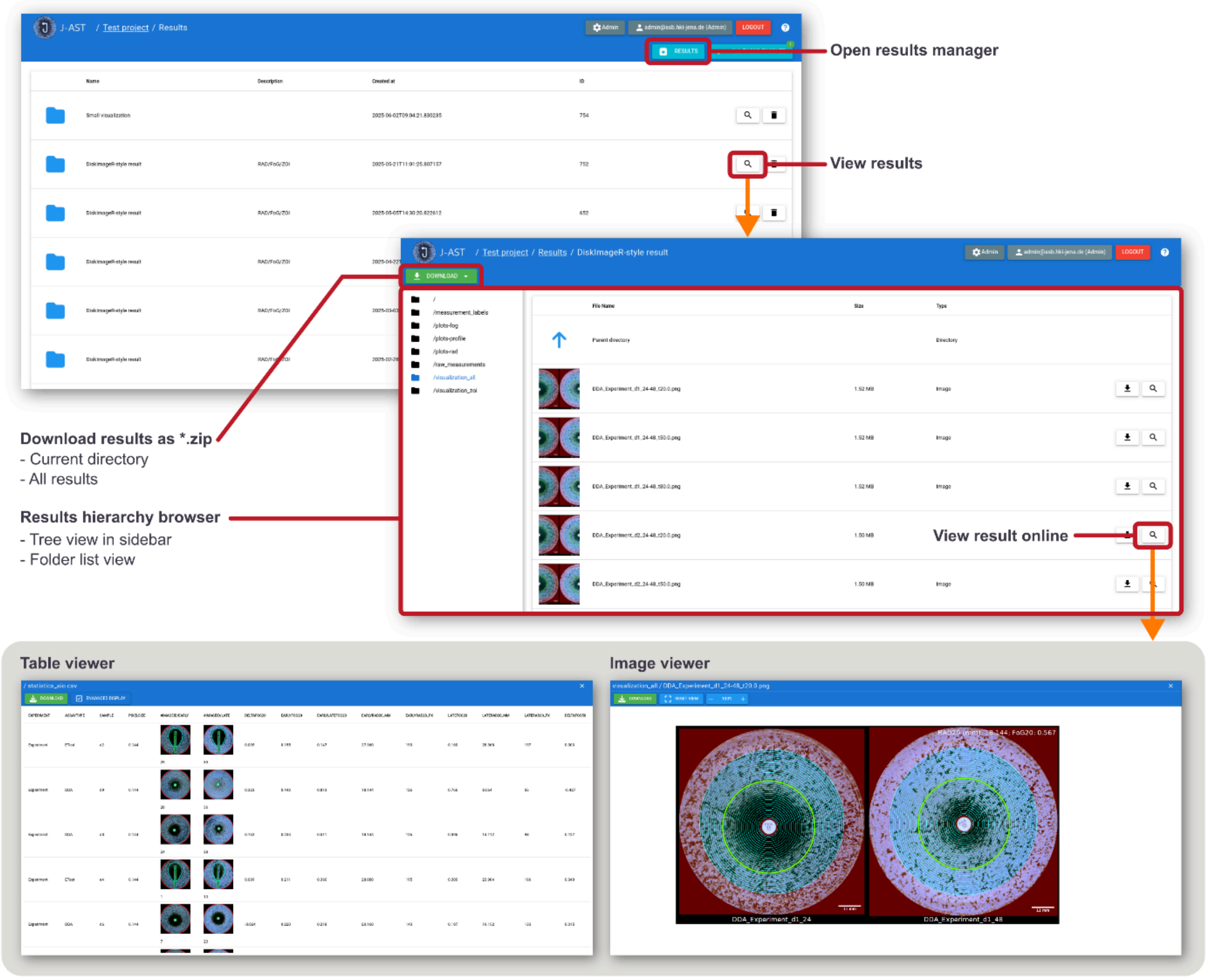

**Supplementary Figure 4:** Results browser. The list of results can be opened by clicking the “Results” button while a project is open. On selecting an item, the hierarchy of result files can be browsed directly within the browser. Image and CSV table files are directly supported by the J-AST viewer and can be reviewed without downloading. Results can be either downloaded individually or as a ZIP package.

### 2 Image analysis algorithms

#### 2.1 Plate detection

##### 2.1.1 Slow algorithm

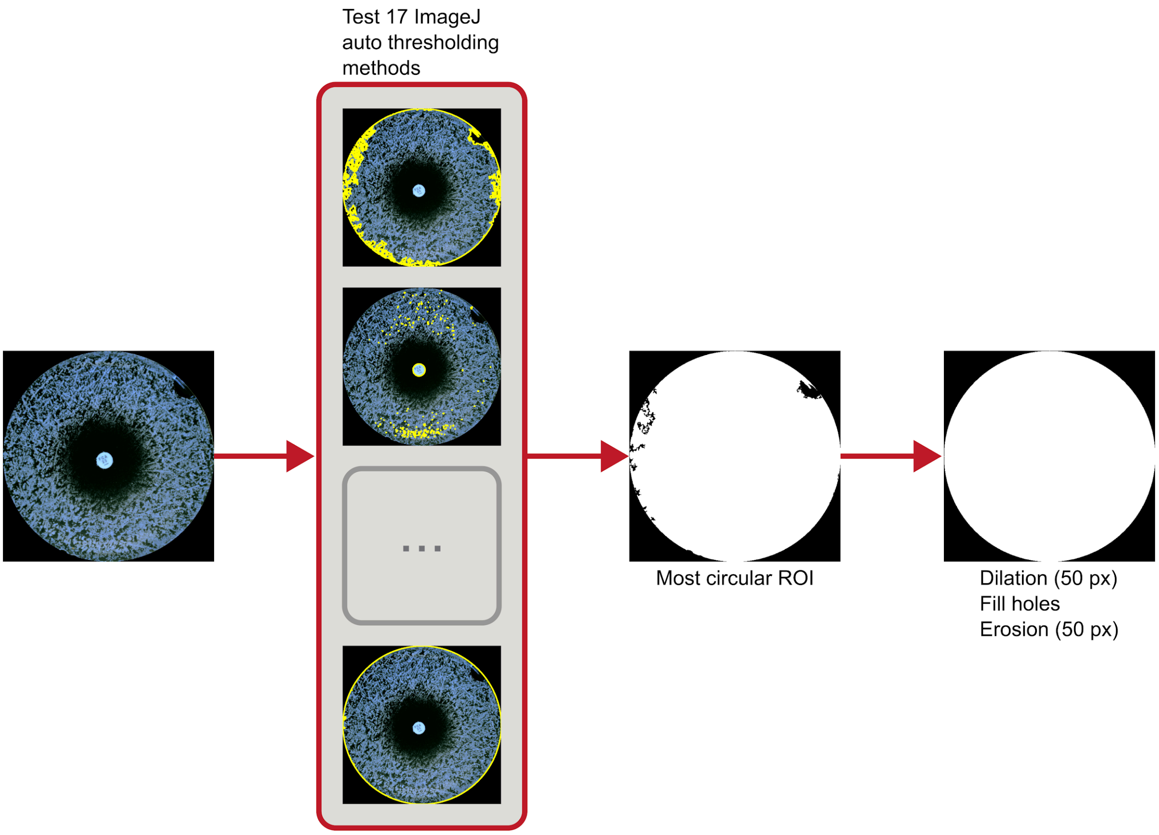

**Supplementary Figure 5:** Slow plate detection algorithm. All available ImageJ auto-thresholding methods are applied on the raw image. From the resulting masks, the most circular region is selected and postprocessed with morphological dilation, hole filling, and erosion.

The following steps are applied by the slow plate detection algorithm. See **Supplementary Figure 5** for an overview and **Supplementary Figure 6** for a screenshot of the pipeline.

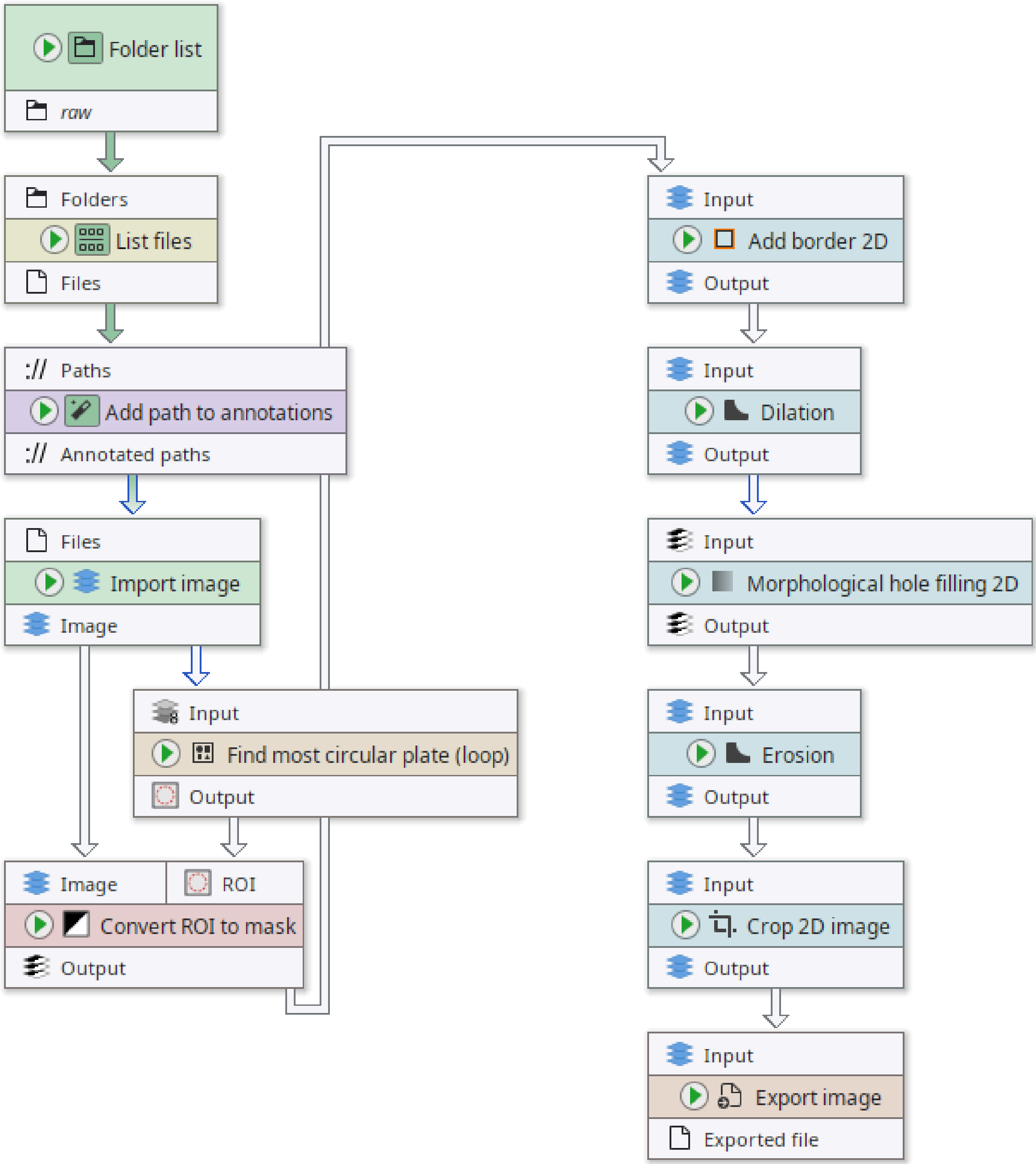

**Supplementary Figure 6:** Full pipeline of the slow plate detection algorithm.

##### *Node #1 "Folder list"*

This node obtains a reference to the folder containing the raw images.

- The parameter item #1 of "Folder paths" (folder-paths) is set to **"raw"**

##### *Node #2 "List files"*

This node lists the files in the input folder.

- Input "Folders" of node #2 receives data from output "Folder paths" of node #1

- The parameter "Keep file if ..." (filters) is set to **`STRING_MATCHES_GLOB(name, "*.png")`**

*Node #3 "Add path to annotations" of type "Annotate with file/directory name"*

This node annotates the image ID to each input file.

- Input "Paths" of node #3 receives data from output "Files" of node #2
- The parameter "Generated annotation" (generated-annotation) is set to **"#ImageId"**

*Node #4 "Import image"*

This node imports the input file(s) as ImageJ image(s).

- Input "Files" of node #4 receives data from output "Annotated paths" of node #3

*Node #5 "Find most circular plate (loop)" of type "Group"*

A group node that encapsulates the testing of multiple auto-thresholding methods to enhance the readability.

- Input "Input" of node #5 receives data from output "Image" of node #4
- This node contains a sub-graph. The "Group Input" and "Group Output" nodes contained in the graph are internally connected to the inputs and outputs of this node.

*Node #6 "Define multiple parameters"*

This node contains the auto-thresholding methods to be tested.

- The parameter "Parameters" (parameter-table) is set to contain the methods **Default, Huang, MaxEntropy, Mean, Minimum, Otsu, Percentile, Yen, Shanbhag, Intermodes, Triangle, MinError, Moments, IsoData, IJ_IsoData, RenyiEntropy,** and **Li**

*Node #7 "Group input"*

This node acts as input for the the group (Node #5).

*Node #8 "Annotate with image properties"*

This node adds the image width and height into the text annotations set.

- Input "Image" of node #8 receives data from output "Input" of node #7
- The parameter "Annotate with image height" (height-annotation) is set to **"imh"**
- The parameter "Annotate with image width" (width-annotation) is set to **"imw"**

*Node #9 "Auto threshold 2D"*

This node applies the auto-thresholding. It receives the parameters from Node #6, meaning that it will apply its workload per parameter set.

- Input "Input" of node #9 receives data from output "Annotated image" of node #8
- Input "{Parameters}" of node #9 receives data from output "Parameters" of node #6
- External parameters are enabled

*Node #10 "Find particles 2D"*

This node converts the auto-threshold outputs into ImageJ ROI lists.

- Input "Mask" of node #10 receives data from output "Output" of node #9

***Node #11 "Merge ROI lists"***

This node merges all generated ROI lists into a singular one.

- Input "Input" of node #11 receives data from output "ROI" of node #10

***Node #12 "Filter ROI by statistics"***

This node selects only the ROIs that cover at least 40% of the image area.

- Input "ROI" of node #12 receives data from output "Output" of node #11
- The parameter "Keep ROI if" (filter) is set to **`(Area / (NUM(imw) * NUM(imh))) >= GET_ITEM(custom,"min-coverage")`**
- The parameter "Measurements" (measurements) is set to only measure the area

***Node #13 "Annotate with ROI properties"***

This node adds the ROI count as a “Count” annotation of each ROI list.

- Input "Input" of node #13 receives data from output "Output" of node #12

***Node #14 "Split & filter by annotation"***

This node selects only the ROI lists that are not empty.

- Input "Input" of node #14 receives data from output "Output" of node #13
- The parameter "Output" in category "Filters" (target-slots/Output) is set to **`TO_NUMBER(Count) > 0`**

***Node #15 "Remove annotation"***

This node deletes the “Count” annotation, as it's not needed anymore.

- Input "Input" of node #15 receives data from output "Output" of node #14
- The parameter "Removed annotations" (annotation-type) is set to **"Count"**

***Node #16 "Sort and extract ROI by statistics"***

This node selects the ROI with the highest circularity.

- Input "ROI" of node #16 receives data from output "Output" of node #15
- The parameter "Selected indices" (selected-indices) is set to **0**
- The parameter item #1 of "Sort order" (sort-orders) is set to **("ShapeDescriptorCircularity", "Descending")**

***Node #17 "Group output"***

This node acts as an interface to the outside of the group.

- Input "Output" of node #17 receives data from output "Output" of node #16

***Node #18 "Convert ROI to mask" of type "Convert 2D ROI to mask"***

This node converts the ROI into a mask.

- Input "Image" of node #18 receives data from output "Image" of node #4
- Input "ROI" of node #18 receives data from output "Output" of node #5

##### *Node #19 "Add border 2D" of type "Add border 2D"*

This node adds a black border around the mask.

- Input "Input" of node #19 receives data from output "Output" of node #18
- The parameter "Margin right" (margin-right) is set to **250**
- The parameter "Margin left" (margin-left) is set to **250**
- The parameter "Margin top" (margin-top) is set to **250**
- The parameter "Margin bottom" (margin-bottom) is set to **250**

##### *Node #20 "Dilation" of type "Morphological operation 2D"*

This node applies a morphological dilation with radius 50 to smooth the plate.

- Input "Input" of node #20 receives data from output "Output" of node #19
- The parameter "Radius" (radius) is set to **50**

##### *Node #21 "Morphological hole filling 2D" of type "Morphological hole filling 2D"*

This node fills all inside holes by applying a flood-filling algorithm.

- Input "Input" of node #21 receives data from output "Output" of node #20

##### *Node #22 "Erosion" of type "Morphological operation 2D"*

This node erodes the plate by 50 pixels to retain the original area.

- Input "Input" of node #22 receives data from output "Output" of node #21
- The parameter "Radius" (radius) is set to **50**
- The parameter "Operation" (operation) is set to **"EROSION"**

##### *Node #23 "Crop 2D image"*

This node removes the border around the image.

- Input "Input" of node #23 receives data from output "Output" of node #22
- Left, top, right, and bottom set to 250 pixels, Center/Center anchor.

##### *Node #24 "Export image"*

This node exports the final mask into the expected directory.

- Input "Input" of node #24 receives data from output "Output" of node #23
- The parameter "File path" (file-path) is set to **`PATH_COMBINE(project_data_dir.tmp_dir, "plate", #ImageId)`**

#### 2.1.2 Fast algorithm

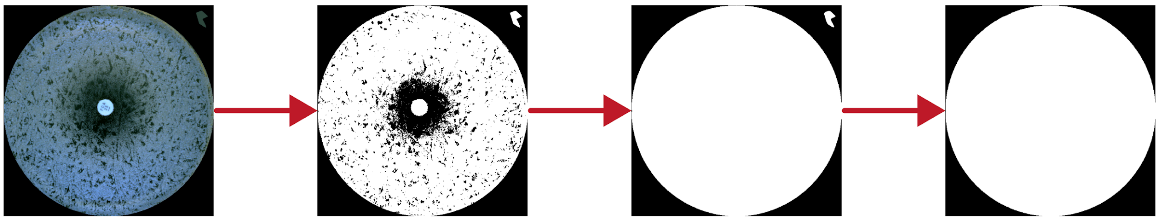

Otsu thresholding

Dilation (50 px)
Fill holes
Erosion (50 px)

Keep largest object
Fit circle

**Supplementary Figure 7:** Fast plate detection algorithm. The raw image is thresholded using Otsu's method. The resulting mask is postprocessed by applying a morphological dilation, hole filling, and erosion. Only the largest object is kept. An artifact was added to highlight the usage of the "Keep largest object" operation.

The following steps are applied by the fast plate detection algorithm. See **Supplementary Figure 7** for an overview and **Supplementary Figure 8** for a screenshot of the pipeline.

##### *Node #1 "Folder list"*

This node obtains a reference to the directory containing the raw inputs.

- The parameter item #1 of "Folder paths" (folder-paths) is set to **"raw"**

##### *Node #2 "List files"*

This node lists the files inside the input directory.

- Input "Folders" of node #2 receives data from output "Folder paths" of node #1
- The parameter "Keep file if ..." (filters) is set to **`STRING_MATCHES_GLOB(name, "*.png")`**

##### *Node #3 "Add path to annotations" of type "Annotate with file/directory name"*

This node attaches the image ID to each input file.

- Input "Paths" of node #3 receives data from output "Files" of node #2
- The parameter "Generated annotation" (generated-annotation) is set to **"#ImageId"**

##### *Node #4 "Import image"*

This node imports the incoming input files as ImageJ image.

- Input "Files" of node #4 receives data from output "Annotated paths" of node #3

##### *Node #5 "Auto threshold 2D"*

This node applies Otsu thresholding.

- Input "Input" of node #5 receives data from output "Image" of node #4
- The parameter "Method" (method) is set to **"Otsu"**

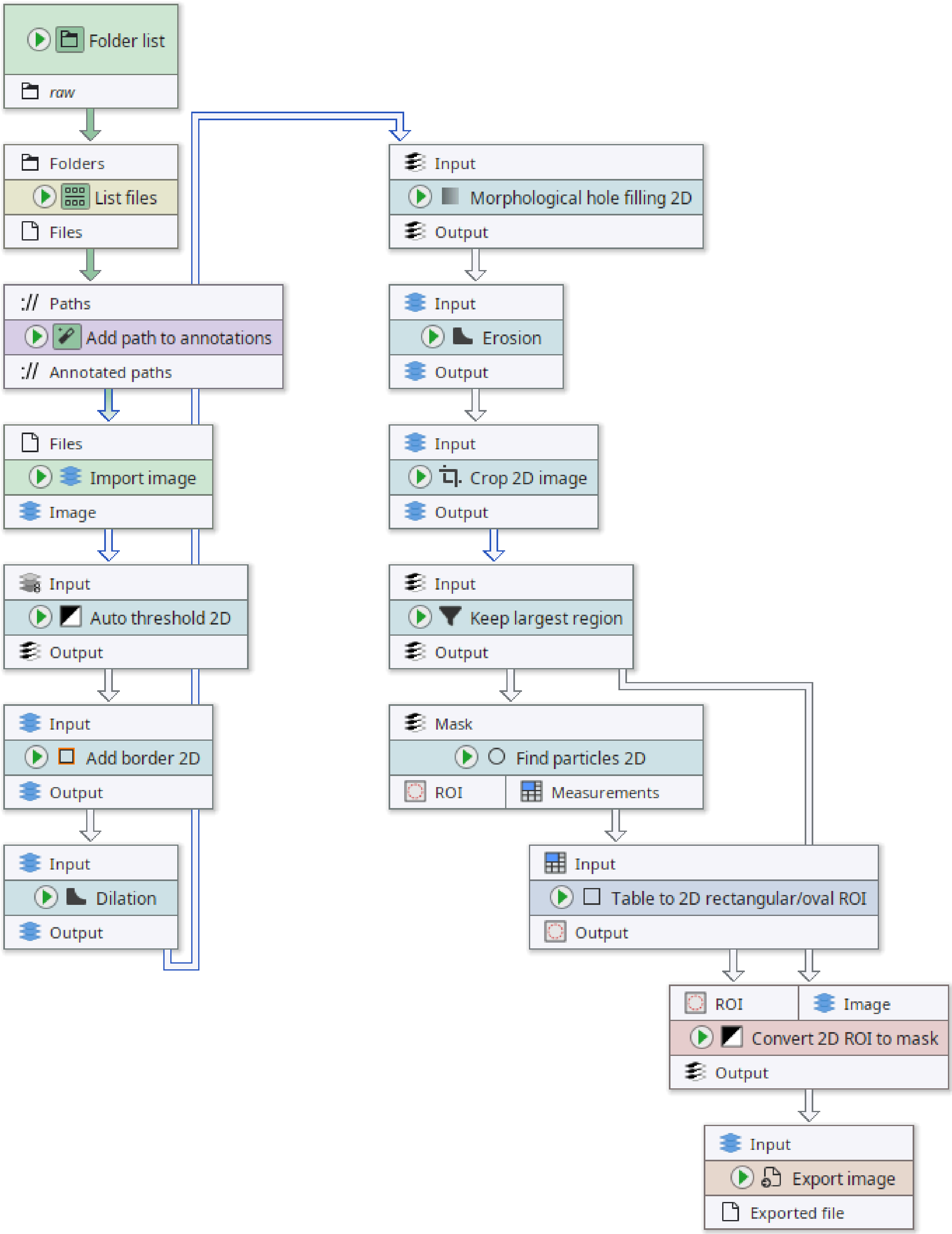

**Supplementary Figure 8:** Full pipeline of the fast plate detection algorithm.

##### *Node #6 "Add border 2D"*

This node adds a border around the image.

- Input "Input" of node #6 receives data from output "Output" of node #5

- The parameter "Margin right" (margin-right) is set to **250**
- The parameter "Margin left" (margin-left) is set to **250**
- The parameter "Margin top" (margin-top) is set to **250**
- The parameter "Margin bottom" (margin-bottom) is set to **250**

##### *Node #7 "Dilation" of type "Morphological operation 2D"*

This node applies morphological dilation with radius 50 px to smooth the mask.

- Input "Input" of node #7 receives data from output "Output" of node #6
- The parameter "Radius" (radius) is set to **50**

##### *Node #8 "Morphological hole filling 2D"*

This node fills holes inside the plate.

- Input "Input" of node #8 receives data from output "Output" of node #7

##### *Node #9 "Erosion" of type "Morphological operation 2D"*

This node restores the original plate area after the dilation.

- Input "Input" of node #9 receives data from output "Output" of node #8
- The parameter "Radius" (radius) is set to **50**
- The parameter "Operation" (operation) is set to **"EROSION"**

##### *Node #10 "Crop 2D image"*

This node removes the border that was added in an earlier step.

- Input "Input" of node #10 receives data from output "Output" of node #9
- Left, top, right, and bottom set to 250 pixels, Center/Center anchor.

##### *Node #11 "Keep largest region"*

This node removes all connected components except the largest one.

- Input "Input" of node #11 receives data from output "Output" of node #10

##### *Node #12 "Find particles 2D"*

This node applies the ImageJ particle finder, converting a mask into a set of ROIs.

- Input "Mask" of node #12 receives data from output "Output" of node #11

##### *Node #13 "Table to 2D rectangular/oval ROI"*

This node converts a table into a set of circular ROI by going through each row and extracting information from the column values.

- Input "Input" of node #13 receives data from output "Measurements" of node #12
- The parameter "Column 'X1'" (column-x1) is set to **("ExistingColumn", "BX")**
- The parameter "Column 'X2'" (column-x2) is set to **("Generate", 0)**
- The parameter "Created ROI type" (mode) is set to **"MinCircle"**
- The parameter "Column 'Y2'" (column-y2) is set to **("Generate", 0)**
- The parameter "Column 'Y1'" (column-y1) is set to **("ExistingColumn", "BY")**
- The parameter "ROI name" in category "ROI properties" (roi-properties/roi-name) is set to **"unnamed"**

##### *Node #14 "Convert 2D ROI to mask"*

This node converts a ROI list into a mask using an image as size reference.

- Input "ROI" of node #14 receives data from output "Output" of node #13
- Input "Image" of node #14 receives data from output "Output" of node #11

##### *Node #15 "Export image"*

This node exports the final mask as image file.

- Input "Input" of node #15 receives data from output "Output" of node #14
- The parameter "File path" (file-path) is set to **`PATH_COMBINE(project_data_dir.tmp_dir, "plate", #ImageId)`**

#### 2.2 Pixel size calibration

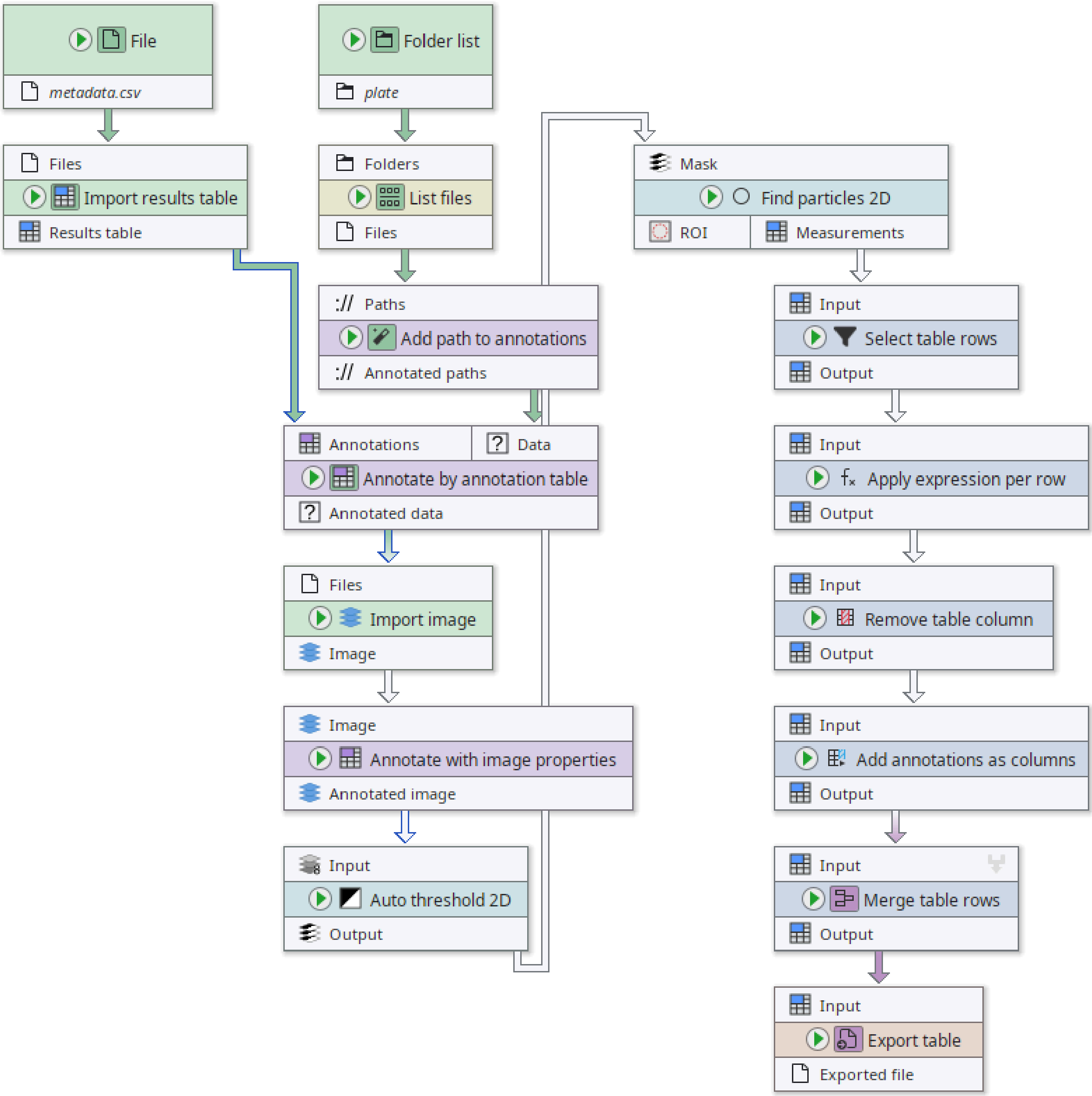

**Supplementary Figure 9:** Screenshot of the pipeline for pixel size calibration.

The following steps are applied by the algorithm that calculates the physical pixel size in millimeters from the plate annotation. A screenshot can be found in **Supplementary Figure 9**.

##### *Node #1 "File"*

This node references the metadata file that associates the image ID to other metadata fields.

- The parameter "File name" (file-name) is set to **"metadata.csv"**

##### *Node #2 "Folder list"*

This node references the directory of plate annotation masks.

- The parameter item #1 of "Folder paths" (folder-paths) is set to **"plate"**

***Node #3 "Import results table"***
This node imports the metadata table into a JIPipe data structure.

- Input "Files" of node #3 receives data from output "Filename" of node #1

***Node #4 "List files"***
This node lists all image files inside the directory that contains the plate annotation masks.

- Input "Folders" of node #4 receives data from output "Folder paths" of node #2
- The parameter "Keep file if ..." (filters) is set to **`STRING_MATCHES_GLOB(name, "*.png")`**

***Node #5 "Add path to annotations" of type "Annotate with file/directory name"***
This node annotates the image files with their unique ID.

- Input "Paths" of node #5 receives data from output "Files" of node #4
- The parameter "Generated annotation" (generated-annotation) is set to **"#ImageId"**

***Node #6 "Annotate by annotation table"***
This node associates metadata from the metadata table to the input images.

- Input "Annotations" of node #6 receives data from output "Results table" of node #3
- Input "Data" of node #6 receives data from output "Annotated paths" of node #5

***Node #7 "Import image"***
This node imports the plate masks.

- Input "Files" of node #7 receives data from output "Annotated data" of node #6

***Node #8 "Annotate with image properties"***
This node stores the image width as annotation.

- Input "Image" of node #8 receives data from output "Image" of node #7
- The parameter "Annotate with image width" (width-annotation) is set to **"image_width"**

***Node #9 "Auto threshold 2D"***
This node applies an auto-thresholding to ensure that the plate mask is a binary image.

- Input "Input" of node #9 receives data from output "Annotated image" of node #8

***Node #10 "Find particles 2D"***
This node converts the plate mask into a set of ImageJ ROIs and measures the bounding rectangle.

- Input "Mask" of node #10 receives data from output "Output" of node #9
- The parameter "Extracted measurements" (measurements) is set to **{"values":["BoundingRectangle"],"collapsed":false}**

***Node #11 "Select table rows"***

This node extracts only the first row from the bounding rectangle measurements.

- Input "Input" of node #11 receives data from output "Measurements" of node #10
- The parameter "Limit" (limit) is set to **0**

*Node #12 "Apply expression per row"*

This node calculates the pixel size based on the width of the plate ROI bounding box and the image width. It requires the plate diameter in millimeters (default: 90 mm).

- Input "Input" of node #12 receives data from output "Output" of node #11
- The parameter item #1 of "Generated values" (entries) is set to **column-name = `"PixelSize"`, value = `ROUNDD(custom.plate_diameter_mm / Width ,3)`**
- The parameter "Plate diameter (mm)" in category "Custom variables" (jipipe:algorithm:custom-expression-variables/plate_diameter_mm) is set to **90.0**

*Node #13 "Remove table column"*

This node removes all table columns except “PixelSize”.

- Input "Input" of node #13 receives data from output "Output" of node #12
- The parameter "Filters" (filters) is set to **`value != "PixelSize"`**

*Node #14 "Add annotations as columns"*

This node adds the annotations of the current table into the table as columns, with the exception of the “PixelSize” annotation. As Node #6 added all existing metadata as annotations, this effectively reconstructs the metadata table with an updated pixel size.

- Input "Input" of node #14 receives data from output "Output" of node #13
- The parameter "Annotation name filter" (annotation-name-filter) is set to **`value != "PixelSize"`**

*Node #15 "Merge table rows"*

This node creates a singular table for all input images.

- Input "Input" of node #15 receives data from output "Output" of node #14
- The parameter "Grouping method" in category "Merging iteration step generation" (jipipe:data-batch-generation/column-matching) is set to **"MergeAll"**

*Node #16 "Export table"*

This node exports the updated metadata table to be read by J-AST.

- Input "Input" of node #16 receives data from output "Output" of node #15
- The parameter "File path" (file-path) is set to **`PATH_COMBINE( project_data_dir.tmp_dir , "metadata_updated.csv")`**

#### 2.3 DDA disk detection

##### 2.3.1 Variant 1 (Slow)

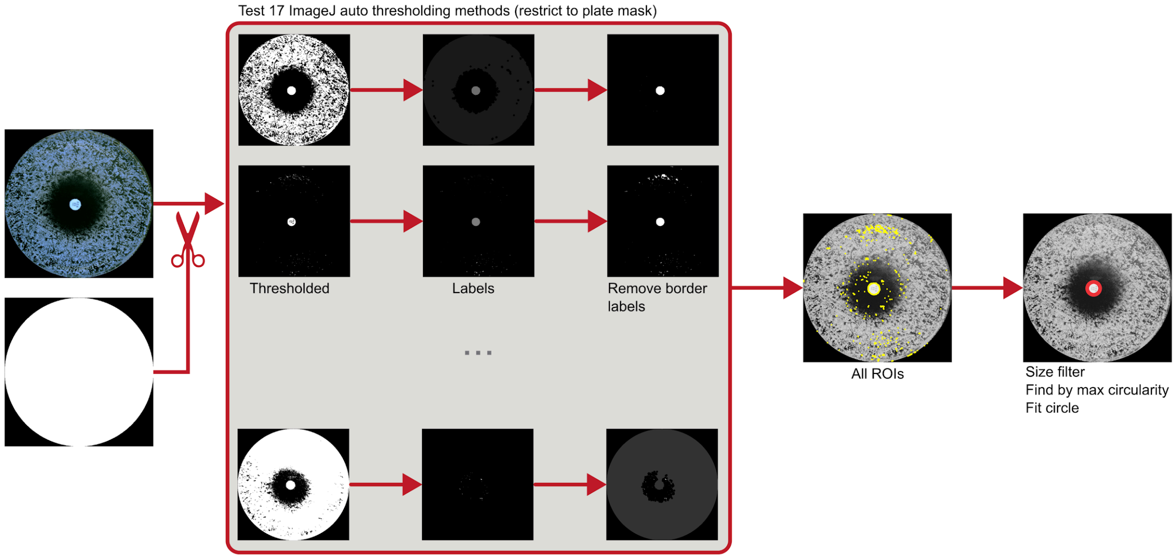

**Supplementary Figure 10:** First variant of the DDA disk detection; slow version. The regions outside the plate are removed. This is followed by a set of operations that are applied per ImageJ auto-thresholding method: the image is thresholded and objects that touch the image border are removed. The resulting ROI over all thresholding methods are collected and filtered by size. The object with the maximum circularity is selected and further processed using a circle fitting algorithm.

The following steps are applied by the first algorithm (slow version) that finds the disk region inside DDA. See **Supplementary Figure 10** for an overview and **Supplementary Figure 11** for a screenshot of the pipeline.

###### *Node #1 "Folder list"*

This node references the directory that contains the raw images.

- The parameter item #1 of "Folder paths" (folder-paths) is set to **"raw"**

###### *Node #2 "Folder list"*

This node references the directory that contains the plate annotations.

- The parameter item #1 of "Folder paths" (folder-paths) is set to **"plate"**

###### *Node #3 "File"*

This node references a CSV table file that associates image IDs with their metadata.

- The parameter "File name" (file-name) is set to **"metadata.csv"**

###### *Node #4 "List files"*

This node lists all PNG files in the directory that contains the raw images.

- Input "Folders" of node #4 receives data from output "Folder paths" of node #1

- The parameter "Keep file if ..." (filters) is set to **`STRING_MATCHES_GLOB(name, "*.png")`**

##### *Node #5 "List files"*

This node lists all PNG files in the directory that contains the plate masks.

- Input "Folders" of node #5 receives data from output "Folder paths" of node #2
- The parameter "Keep file if ..." (filters) is set to **`STRING_MATCHES_GLOB(name, "*.png")`**

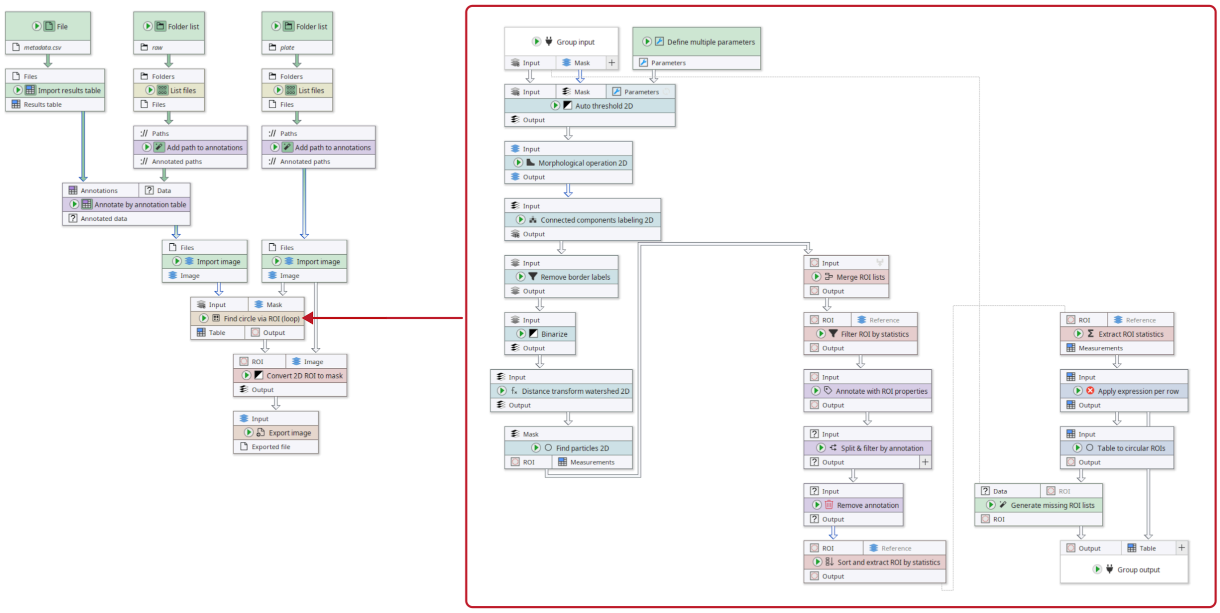

**Supplementary Figure 11:** Screenshot of the DDA disk detection pipeline (Variant 1, slow). The pipeline contains a group node that has an embedded pipeline (red box).

##### *Node #6 "Import results table"*

This node imports a CSV file into a table data structure.

- Input "Files" of node #6 receives data from output "Filename" of node #3

##### *Node #7 "Add path to annotations" of type "Annotate with file/directory name"*

This node attaches the image ID to the image file.

- Input "Paths" of node #7 receives data from output "Files" of node #4
- The parameter "Generated annotation" (generated-annotation) is set to **"#ImageId"**

##### *Node #8 "Add path to annotations" of type "Annotate with file/directory name"*

This node attaches the image ID to the image file.

- Input "Paths" of node #8 receives data from output "Files" of node #5
- The parameter "Generated annotation" (generated-annotation) is set to **"#ImageId"**

*Node #9 "Annotate by annotation table"*

This node associates metadata stored inside the input CSV table to the image file.

- Input "Annotations" of node #9 receives data from output "Results table" of node #6
- Input "Data" of node #9 receives data from output "Annotated paths" of node #7

*Node #10 "Import image"*

This node imports the image file as ImageJ image.

- Input "Files" of node #10 receives data from output "Annotated paths" of node #8

*Node #11 "Import image"*

This node imports the image file as ImageJ image.

- Input "Files" of node #11 receives data from output "Annotated data" of node #9

*Node #12 "Find circle via ROI (loop)" of type "Group"*

This node is a group that encapsulates the detection of the DDA disk.

- Input "Input" of node #12 receives data from output "Image" of node #11
- Input "Mask" of node #12 receives data from output "Image" of node #10
- This node contains a sub-graph. The "Group Input" and "Group Output" nodes contained in the graph are internally connected to the inputs and outputs of this node.

*Node #13 "Define multiple parameters"*

This node contains parameters that refer to all available ImageJ auto-thresholding methods.

- The parameter "Parameters" (parameter-table) is set to contain the methods **Default, Huang, MaxEntropy, Mean, Minimum, Otsu, Percentile, Yen, Shanbhag, Intermodes, Triangle, MinError, Moments, IsoData, IJ_IsoData, RenyiEntropy,** and **Li**

*Node #14 "Group input"*

This node acts as interface between the outside and inside of the group (node #12).

*Node #15 "Auto threshold 2D"*

This node applies auto-thresholding of the raw image, limited to the plate region. The node is configured to repeat its workload per parameter set. Those parameter sets are received from node #13.

- Input "Input" of node #15 receives data from output "Input" of node #14
- Input "Mask" of node #15 receives data from output "Mask" of node #14
- Input "{Parameters}" of node #15 receives data from output "Parameters" of node #13
- The parameter "Calculate threshold based on ..." (source-area) is set to **"InsideMask"**
- The parameter "Multiple parameters" in category "Multi-parameter settings" (jipipe:parameter-slot-algorithm/has-parameter-slot) is set to **true**
- The parameter "Attach parameter annotations" in category "Multi-parameter settings" (jipipe:parameter-slot-algorithm/attach-parameter-annotations) is set to **false**

*Node #16 "Morphological operation 2D"*

This node applies a morphological closing operation with radius 5 px.

- Input "Input" of node #16 receives data from output "Output" of node #15
- The parameter "Radius" (radius) is set to **5**
- The parameter "Operation" (operation) is set to **"CLOSING"**

***Node #17 "Connected components labeling 2D"***

This node converts the binary mask into a label image.

- Input "Input" of node #17 receives data from output "Output" of node #16

***Node #18 "Remove border labels"***

This node removes all labels that touch the image border.

- Input "Input" of node #18 receives data from output "Output" of node #17

***Node #19 "Binarize"***

This node converts the label image back into a mask.

- Input "Input" of node #19 receives data from output "Output" of node #18

***Node #20 "Distance transform watershed 2D"***

This node applies a distance transform watershed to ensure that there are no holes.

- Input "Input" of node #20 receives data from output "Output" of node #19

***Node #21 "Find particles 2D"***

This node applies the ImageJ particle finder for ROI detection.

- Input "Mask" of node #21 receives data from output "Output" of node #20

***Node #22 "Merge ROI lists"***

This node merges all generated per-threshold-algorithm ROI lists into a singular one.

- Input "Input" of node #22 receives data from output "ROI" of node #21

***Node #23 "Filter ROI by statistics" of type "Filter 2D ROI by statistics"***

This node applies a size and circularity filter on the ROIs detected by node #22.

- Input "ROI" of node #23 receives data from output "Output" of node #22
- The parameter "Keep ROI if" (filter) is set to **`SET_VARIABLE("cminf_px", custom.minFeret / NUM(PixelSize)); SET_VARIABLE("cmaxf_px", custom.maxFeret / NUM(PixelSize)); MinFeret >= cminf_px AND Feret >= cminf_px AND MinFeret <= cmaxf_px AND Feret <= cmaxf_px AND Circ. >= custom.minCirc`**
- The parameter "Measurements" (measurements) is set to **{"values":["FeretDiameter","ShapeDescriptors"],"collapsed":false}**
- The parameter "Minimum diameter (mm)" in category "Custom variables" (jipipe:algorithm:custom-expression-variables/minFeret) is set to **2.0**
- The parameter "Maximum diameter (mm)" in category "Custom variables" (jipipe:algorithm:custom-expression-variables/maxFeret) is set to **12.0**
- The parameter "Minimum circularity" in category "Custom variables" (jipipe:algorithm:custom-expression-variables/minCirc) is set to **0.5**

***Node #24 "Annotate with ROI properties"***

This node attaches the number ROIs to the currently processed ROI list.

- Input "Input" of node #24 receives data from output "Output" of node #23

***Node #25 "Split & filter by annotation"***

This node only selects ROI lists with at least one ROI.

- Input "Input" of node #25 receives data from output "Output" of node #24
- The parameter "Output" in category "Filters" (target-slots/Output) is set to **`TO_NUMBER(Count) > 0`**

***Node #26 "Remove annotation"***

This node removes the annotation added by node #24.

- Input "Input" of node #26 receives data from output "Output" of node #25
- The parameter "Removed annotations" (annotation-type) is set to **"Count"**

***Node #27 "Sort and extract ROI by statistics" of type "Sort and extract 2D ROI by statistics"***

This node selects the most circular ROI from each ROI list.

- Input "ROI" of node #27 receives data from output "Output" of node #26
- The parameter "Selected indices" (selected-indices) is set to **0**
- The parameter item #1 of "Sort order" (sort-orders) is set to **("ShapeDescriptorCircularity", "Descending")**

***Node #28 "Extract ROI statistics" of type "Extract 2D ROI statistics"***

This node measures the centroid and Feret diameter of the selected ROI.

- Input "ROI" of node #28 receives data from output "Output" of node #27
- The parameter "Extracted measurements" (measurements) is set to **{"values":["Centroid","FeretDiameter"],"collapsed":false}**

***Node #29 "Apply expression per row"***

This node calculates the radius of the DDA disk by dividing its Feret diameter by two.

- Input "Input" of node #29 receives data from output "Measurements" of node #28
- The parameter item #1 of "Expressions" (expression-list) is set to **(`Feret / 2`, `"Radius"`)**

***Node #30 "Table to circular ROIs" of type "Table to 2D circular ROI"***

This node creates a circular ROI from the table generated in node #29.

- Input "Input" of node #30 receives data from output "Output" of node #29
- The parameter "ROI name" in category "ROI properties" (roi-properties/roi-name) is set to **"unnamed"**

***Node #31 "Generate missing ROI lists" of type "Generate missing 2D ROI lists"***

This node creates empty ROI lists where data is missing.

- Input "Data" of node #31 receives data from output "Input" of node #14
- Input "ROI" of node #31 receives data from output "Output" of node #30

###### *Node #32 "Group output"*

This node interfaces the data from the group to the outer pipeline.

- Input "Output" of node #32 receives data from output "ROI" of node #31
- Input "Table" of node #32 receives data from output "Output" of node #29

###### *Node #33 "Convert 2D ROI to mask"*

This node converts ImageJ ROIs into a mask.

- Input "ROI" of node #33 receives data from output "Output" of node #12
- Input "Image" of node #33 receives data from output "Image" of node #10

###### *Node #34 "Export image"*

This node exports the final DDA disk annotation.

- Input "Input" of node #34 receives data from output "Output" of node #33
- The parameter "File path" (file-path) is set to **`PATH_COMBINE(project_data_dir.tmp_dir, "strip-disk", #ImageId)`**

##### 2.3.2 Variant 1 (Fast)

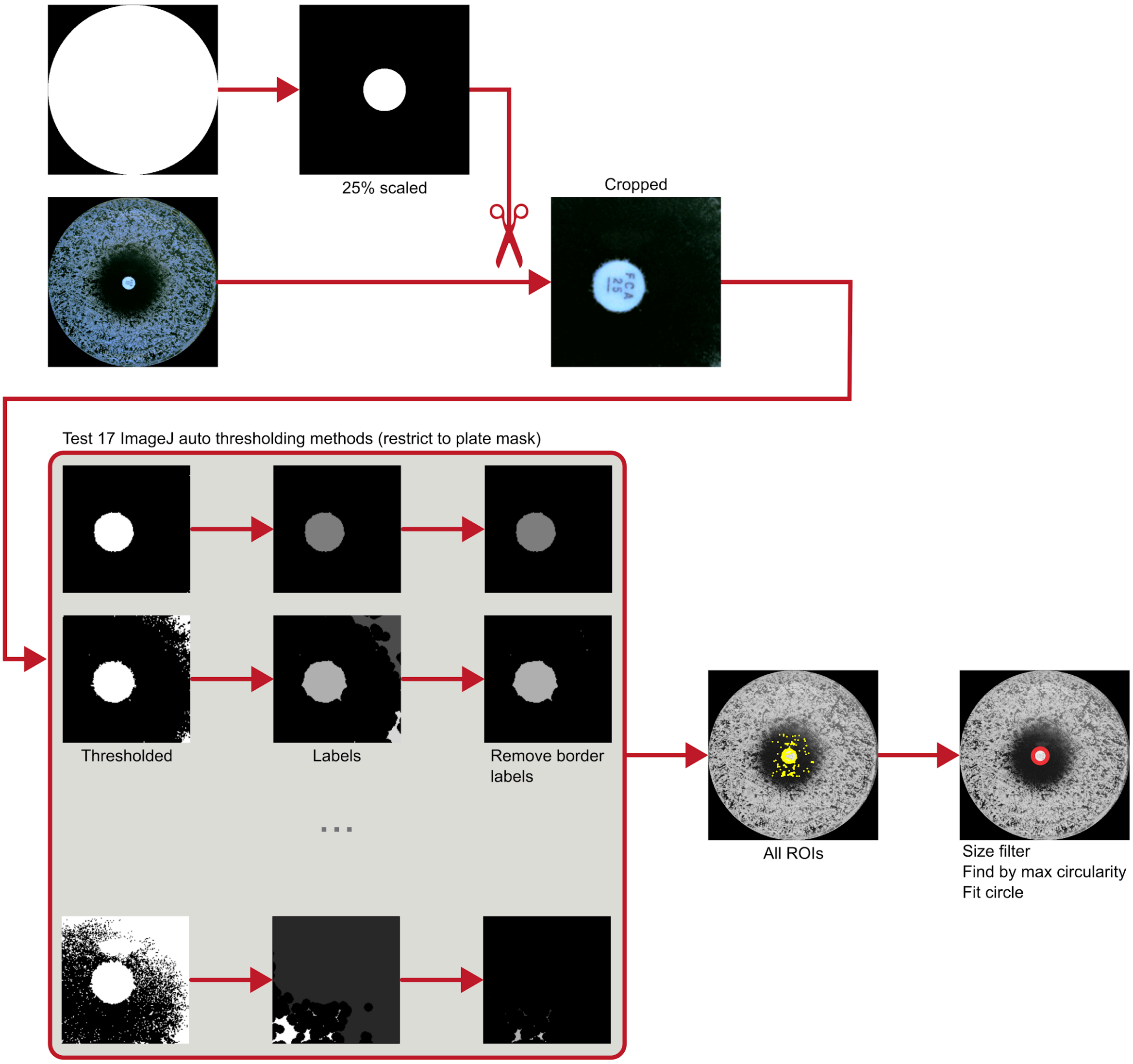

**Supplementary Figure 12:** First variant of the DDA disk detection; fast version. The plate mask is scaled to 25% of its size and used to extract only the central part of the image. This is followed by a set of operations that are applied per ImageJ auto-thresholding method: the image is thresholded and objects that touch the image border are removed. The resulting ROI over all thresholding methods are collected and filtered by size. The object with the maximum circularity is selected and further processed using a circle fitting algorithm.

The following steps are applied by the first algorithm (fast version) that finds the disk region inside DDA. See **Supplementary Figure 12** for an overview and **Supplementary Figure 13** for a screenshot of the pipeline.

###### *Node #1 "Folder list"*

This node references the directory that contains the raw images.

- The parameter item #1 of "Folder paths" (folder-paths) is set to **"raw"**

###### *Node #2 "Folder list"*

This node references the directory that contains the plate annotations.

- The parameter item #1 of "Folder paths" (folder-paths) is set to **"plate"**

##### *Node #3 "File"*

This node references a CSV file that associates metadata to each image ID.

- The parameter "File name" (file-name) is set to **"metadata.csv"**

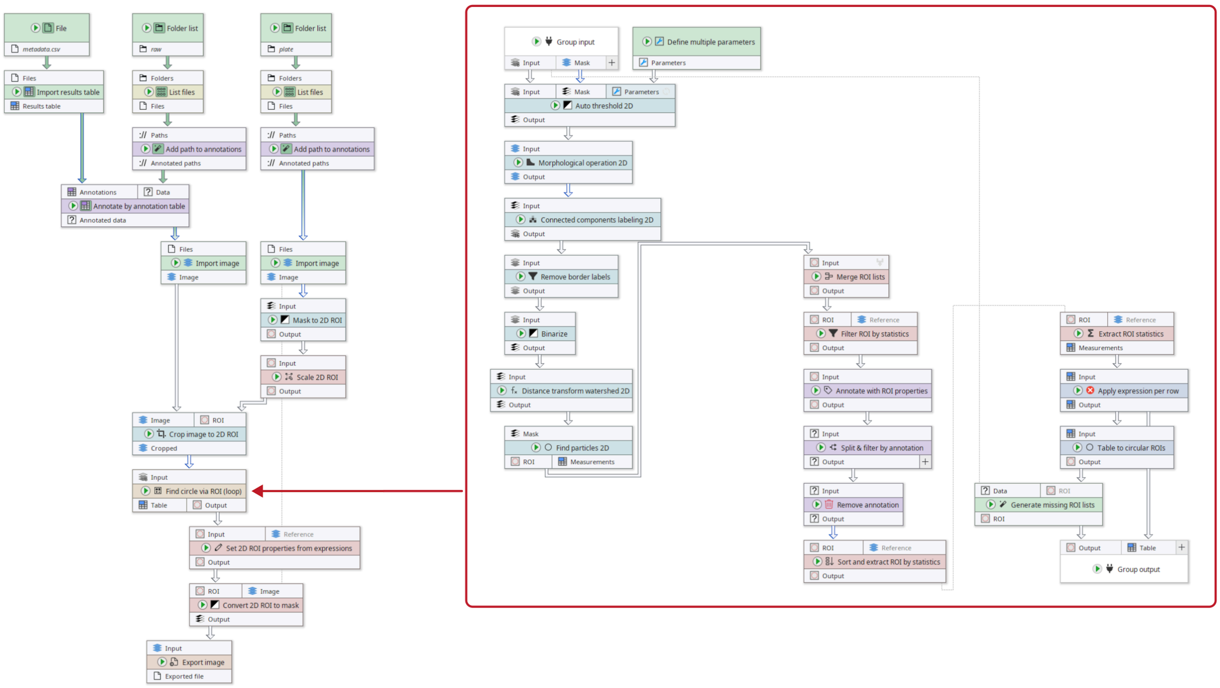

**Supplementary Figure 13:** Screenshot of the DDA disk detection pipeline (Variant 1, fast). The pipeline contains a group node that has an embedded pipeline (red box).

##### *Node #4 "List files"*

This node finds all PNG images in the input directory.

- Input "Folders" of node #4 receives data from output "Folder paths" of node #1
- The parameter "Keep file if ..." (filters) is set to **`STRING_MATCHES_GLOB(name, "*.png")`**

##### *Node #5 "List files"*

This node finds all PNG images in the input directory.

- Input "Folders" of node #5 receives data from output "Folder paths" of node #2
- The parameter "Keep file if ..." (filters) is set to **`STRING_MATCHES_GLOB(name, "*.png")`**

##### *Node #6 "Import results table"*

This node imports a CSV file into an ImageJ table.

- Input "Files" of node #6 receives data from output "Filename" of node #3

***Node #7 "Add path to annotations" of type "Annotate with file/directory name"***

This node attaches the image ID to the processed image file.

- Input "Paths" of node #7 receives data from output "Files" of node #4
- The parameter "Generated annotation" (generated-annotation) is set to **"#ImageId"**

***Node #8 "Add path to annotations" of type "Annotate with file/directory name"***

This node attaches the image ID to the processed image file.

- Input "Paths" of node #8 receives data from output "Files" of node #5
- The parameter "Generated annotation" (generated-annotation) is set to **"#ImageId"**

***Node #9 "Annotate by annotation table"***

This node attaches the image metadata from the metadata table to each image.

- Input "Annotations" of node #9 receives data from output "Results table" of node #6
- Input "Data" of node #9 receives data from output "Annotated paths" of node #7

***Node #10 "Import image"***

This node loads the image file using ImageJ.

- Input "Files" of node #10 receives data from output "Annotated paths" of node #8

***Node #11 "Import image"***

This node loads the image file using ImageJ.

- Input "Files" of node #11 receives data from output "Annotated data" of node #9

***Node #12 "Mask to 2D ROI"***

This node converts a mask image into an ImageJ ROI.

- Input "Input" of node #12 receives data from output "Image" of node #10

***Node #13 "Scale 2D ROI" of type "Scale 2D ROI (old)"***

This node scales the ROI down to 25% of its original size.

- Input "Input" of node #13 receives data from output "Output" of node #12
- The parameter "Scale X" (scale-x) is set to **0.25**
- The parameter "Scale Y" (scale-y) is set to **0.25**
- The parameter "Center scale" (center-scale) is set to **true**

***Node #14 "Crop image to 2D ROI"***

This node crops the raw image to the area defined by the scaled ROI (node #13).

- Input "Image" of node #14 receives data from output "Image" of node #11
- Input "ROI" of node #14 receives data from output "Output" of node #13
- The parameter "Crop channel plane" (crop-c) is set to **false**
- The parameter "Crop Z plane" (crop-z) is set to **false**
- The parameter "Annotate with Y location" (annotation-y) is set to **"crop.y"**
- The parameter "Annotate with X location" (annotation-x) is set to **"crop.x"**
- The parameter "Crop frame plane" (crop-t) is set to **false**

*Node #15 "Find circle via ROI (loop)" of type "Group"*

This node groups the disk detection into for organizational purposes.

- Input "Input" of node #15 receives data from output "Cropped" of node #14
- This node contains a sub-graph. The "Group Input" and "Group Output" nodes contained in the graph are internally connected to the inputs and outputs of this node.

*Node #16 "Define multiple parameters"*

This node produces parameter sets, one for each ImageJ auto-thresholding method.

- The parameter "Parameters" (parameter-table) is set to contain the methods **Default, Huang, MaxEntropy, Mean, Minimum, Otsu, Percentile, Yen, Shanbhag, Intermodes, Triangle, MinError, Moments, IsoData, IJ_IsoData, RenyiEntropy,** and **Li**

*Node #17 "Group input" of type "Group input"*

This node acts as an interface between the group (node #15) and its inside nodes.

*Node #18 "Auto threshold 2D"*

This node applies ImageJ auto-thresholding. As it receives multiple parameter sets from node #16, the workload is repeated for each ImageJ auto-thresholding method.

- Input "Input" of node #18 receives data from output "Input" of node #17
- Input "{Parameters}" of node #18 receives data from output "Parameters" of node #16
- The parameter "Multiple parameters" in category "Multi-parameter settings" (jipipe:parameter-slot-algorithm/has-parameter-slot) is set to **true**
- The parameter "Attach parameter annotations" in category "Multi-parameter settings" (jipipe:parameter-slot-algorithm/attach-parameter-annotations) is set to **false**

*Node #19 "Morphological operation 2D"*

This node applies a morphological closing operation with radius 5 px.

- Input "Input" of node #19 receives data from output "Output" of node #18
- The parameter "Radius" (radius) is set to **5**
- The parameter "Operation" (operation) is set to **"CLOSING"**

*Node #20 "Connected components labeling 2D"*

This node finds the connected mask components and produces a label image.

- Input "Input" of node #20 receives data from output "Output" of node #19

*Node #21 "Remove border labels"*

This node removes all labels that touch the image border.

- Input "Input" of node #21 receives data from output "Output" of node #20

*Node #22 "Binarize"*

This node converts the label image into a mask.

- Input "Input" of node #22 receives data from output "Output" of node #21

*Node #23 "Distance transform watershed 2D"*

This node applies distance transform watershed to remove holes.

- Input "Input" of node #23 receives data from output "Output" of node #22

*Node #24 "Find particles 2D"*

This node applies the ImageJ particle finder to convert masks into ROI lists.

- Input "Mask" of node #24 receives data from output "Output" of node #23

*Node #25 "Merge ROI lists" of type "Merge 2D ROI lists"*

This node merges the ROI lists for all thresholding methods into one list.

- Input "Input" of node #25 receives data from output "ROI" of node #24

*Node #26 "Filter ROI by statistics" of type "Filter 2D ROI by statistics"*

This node applies a size and circularity filter to the ROIs.

- Input "ROI" of node #26 receives data from output "Output" of node #25
- The parameter "Keep ROI if" (filter) is set to **`SET_VARIABLE("cminf_px", custom.minFeret / NUM(PixelSize)); SET_VARIABLE("cmaxf_px", custom.maxFeret / NUM(PixelSize)); MinFeret >= cminf_px AND Feret >= cminf_px AND MinFeret <= cmaxf_px AND Feret <= cmaxf_px AND Circ. >= custom.minCirc`**
- The parameter "Measurements" (measurements) is set to **{"values":["FeretDiameter","ShapeDescriptors"],"collapsed":false}**
- The parameter "Minimum diameter (mm)" in category "Custom variables" (jipipe:algorithm:custom-expression-variables/minFeret) is set to **2.0**
- The parameter "Maximum diameter (mm)" in category "Custom variables" (jipipe:algorithm:custom-expression-variables/maxFeret) is set to **12.0**
- The parameter "Minimum circularity" in category "Custom variables" (jipipe:algorithm:custom-expression-variables/minCirc) is set to **0.5**

*Node #27 "Annotate with ROI properties" of type "Annotate with 2D ROI properties"*

This node adds the number of ROIs as annotation to each ROI list.

- Input "Input" of node #27 receives data from output "Output" of node #26

*Node #28 "Split & filter by annotation"*

This node selects only the ROI lists that have at least one ROI.

- Input "Input" of node #28 receives data from output "Output" of node #27
- The parameter "Output" in category "Filters" (target-slots/Output) is set to **`TO_NUMBER(Count) > 0`**

*Node #29 "Remove annotation"*

This node removes the annotation generated by node #27.

- Input "Input" of node #29 receives data from output "Output" of node #28
- The parameter "Removed annotations" (annotation-type) is set to **"Count"**

##### *Node #30 "Sort and extract ROI by statistics" of type "Sort and extract 2D ROI by statistics"*

This node selects the ROI with the highest circularity.

- Input "ROI" of node #30 receives data from output "Output" of node #29
- The parameter "Selected indices" (selected-indices) is set to **0**
- The parameter item #1 of "Sort order" (sort-orders) is set to **("ShapeDescriptorCircularity", "Descending")**

##### *Node #31 "Extract ROI statistics" of type "Extract 2D ROI statistics"*

This node extracts the centroid and Feret diameter of the ROI.

- Input "ROI" of node #31 receives data from output "Output" of node #30
- The parameter "Extracted measurements" (measurements) is set to **{"values":["Centroid","FeretDiameter"],"collapsed":false}**

##### *Node #32 "Apply expression per row"*

This node calculates the DDA disk radius by dividing the Feret radius by two.

- Input "Input" of node #32 receives data from output "Measurements" of node #31
- The parameter item #1 of "Expressions" (expression-list) is set to **`(Feret / 2, "Radius")`**

##### *Node #33 "Table to circular ROIs" of type "Table to 2D circular ROI"*

This node creates a circular ROI based on the information stored inside the table.

- Input "Input" of node #33 receives data from output "Output" of node #32
- The parameter "ROI name" in category "ROI properties" (roi-properties/roi-name) is set to `"`**`unnamed`**`"`

##### *Node #34 "Generate missing ROI lists" of type "Generate missing 2D ROI lists"*

This node generates an empty ROI list if one is missing for an image ID.

- Input "Data" of node #34 receives data from output "Input" of node #17
- Input "ROI" of node #34 receives data from output "Output" of node #33

##### *Node #35 "Group output"*

This node acts as an interface that moves data from inside the group to the surrounding pipeline.

- Input "Output" of node #35 receives data from output "ROI" of node #34
- Input "Table" of node #35 receives data from output "Output" of node #32

##### *Node #36 "Set 2D ROI properties from expressions"*

This node reverses the cropping by node #14 by moving the ROIs to the correct position.

- Input "Input" of node #36 receives data from output "Output" of node #15
- The parameter "Location (X)" (position-x) is set to **`x + NUM(crop.x)`**
- The parameter "Location (Y)" (position-y) is set to **`y + NUM(crop.y)`**

##### *Node #37 "Convert 2D ROI to mask"*

This node converts the DDA disk ROI into a mask.

- Input "ROI" of node #37 receives data from output "Output" of node #36
- Input "Image" of node #37 receives data from output "Image" of node #10

###### *Node #38 "Export image"*

This node exports the mask into a PNG file.

- Input "Input" of node #38 receives data from output "Output" of node #37
- The parameter "File path" (file-path) is set to **`PATH_COMBINE(project_data_dir.tmp_dir, "strip-disk", #ImageId)`**

##### 2.3.3 Variant 2

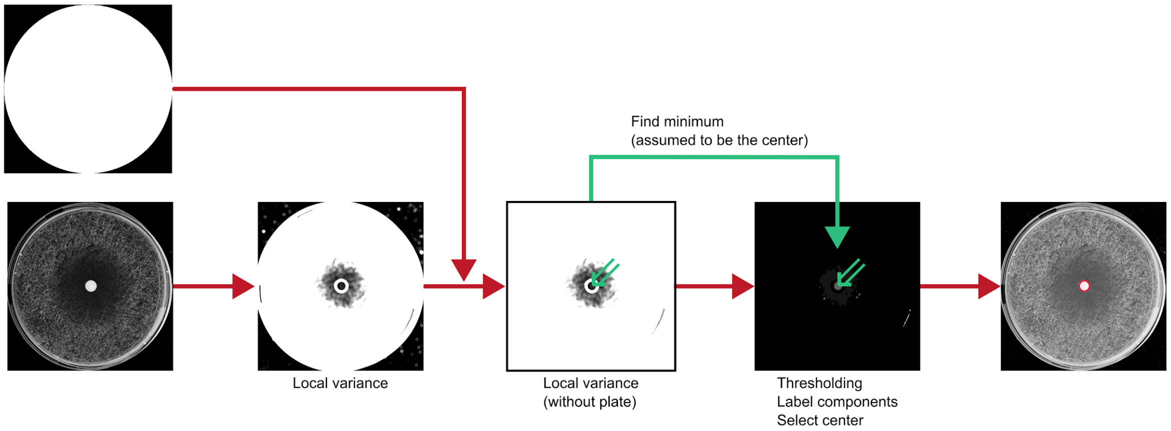

**Supplementary Figure 14:** Second variant of the DDA disk detection. The raw image is processed with a local variance filter. The center of the DDA is detected by finding the minimum inside the plate, which is then utilized to select the disk component from a mask created by an auto-thresholding algorithm.

The following steps are applied by the second algorithm that finds the disk region inside DDA. An overview can be found in **Supplementary Figure 14** and a screenshot is provided in **Supplementary Figure 15**.

###### *Node #1 "File"*

This node references a CSV file that associates the image ID to the image metadata.

- The parameter "File name" (file-name) is set to **"metadata.csv"**

###### *Node #2 "Folder list"*

This node references the directory that contains the raw images.

- The parameter item #1 of "Folder paths" (folder-paths) is set to **"raw"**

###### *Node #3 "Folder list"*

This node references the directory that contains the plate annotation masks.

- The parameter item #1 of "Folder paths" (folder-paths) is set to **"plate"**

##### *Node #4 "Import results table"*

This node imports the metadata CSV file as an ImageJ table.

- Input "Files" of node #4 receives data from output "Filename" of node #1

##### *Node #5 "List files"*

This node lists all PNG files in the directory containing the raw images.

- Input "Folders" of node #5 receives data from output "Folder paths" of node #2
- The parameter "Keep file if ..." (filters) is set to `STRING_MATCHES_GLOB(name, "*.png")`

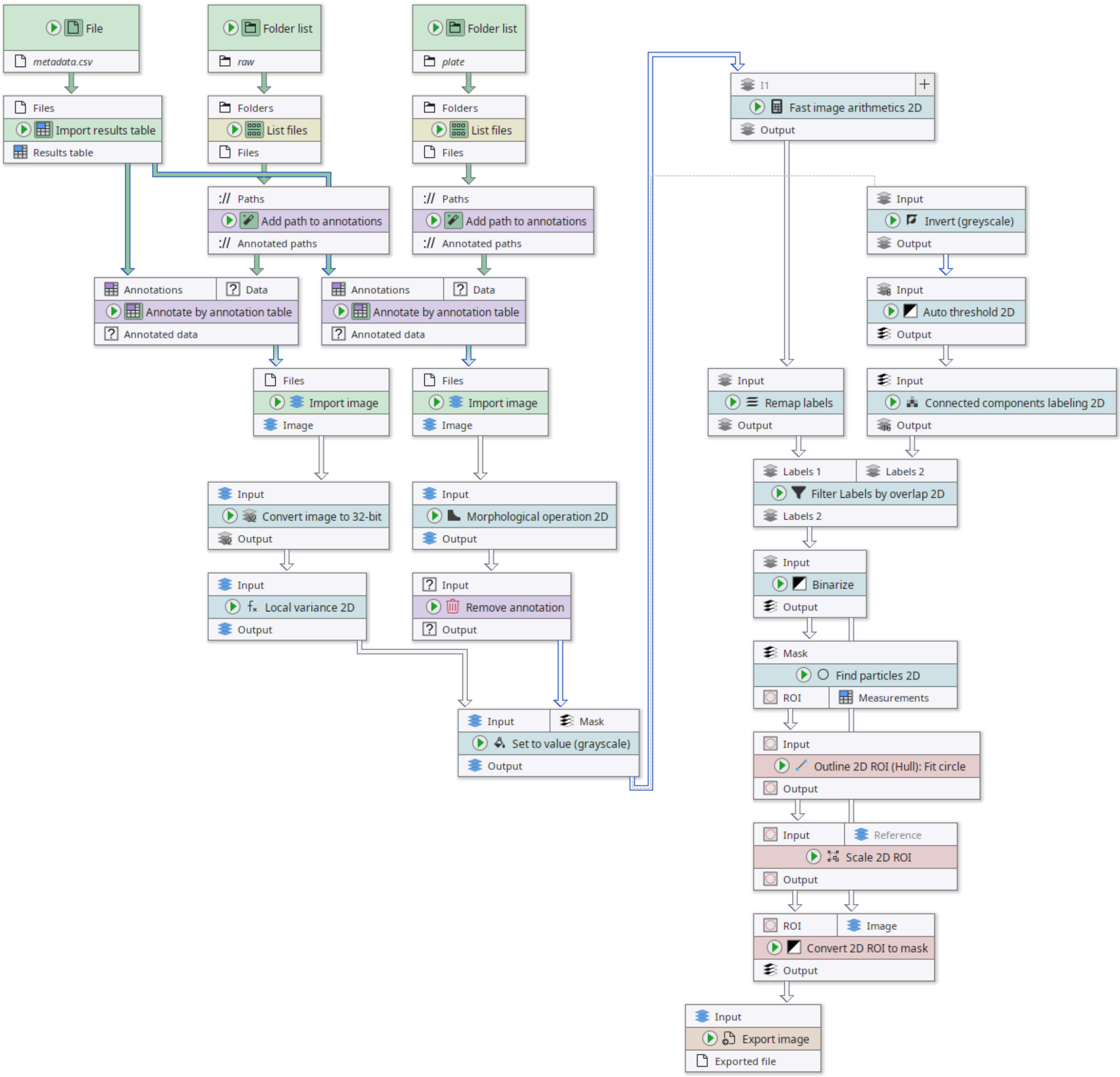

**Supplementary Figure 15:** Screenshot of the second variant of the DDA disk detection.

***Node #6 "List files"***
This node lists all PNG files in the directory that contains the plate masks.

- Input "Folders" of node #6 receives data from output "Folder paths" of node #3
- The parameter "Keep file if ..." (filters) is set to **`STRING_MATCHES_GLOB(name, "*.png")`**

***Node #7 "Add path to annotations" of type "Annotate with file/directory name"***
This node annotates an image file with its image ID.

- Input "Paths" of node #7 receives data from output "Files" of node #5
- The parameter "Generated annotation" (generated-annotation) is set to **"#ImageId"**

***Node #8 "Add path to annotations" of type "Annotate with file/directory name"***
This node annotates an image file with its image ID.

- Input "Paths" of node #8 receives data from output "Files" of node #6
- The parameter "Generated annotation" (generated-annotation) is set to **"#ImageId"**

***Node #9 "Annotate by annotation table"***
This node associates metadata from the metadata table to its respective image file.

- Input "Annotations" of node #9 receives data from output "Results table" of node #4
- Input "Data" of node #9 receives data from output "Annotated paths" of node #7

***Node #10 "Annotate by annotation table"***
This node associates metadata from the metadata table to its respective image file.

- Input "Annotations" of node #10 receives data from output "Results table" of node #4
- Input "Data" of node #10 receives data from output "Annotated paths" of node #8

***Node #11 "Import image"***
This node imports an image file into an ImageJ image.

- Input "Files" of node #11 receives data from output "Annotated data" of node #9

***Node #12 "Import image"***
This node imports an image file into an ImageJ image.

- Input "Files" of node #12 receives data from output "Annotated data" of node #10

***Node #13 "Convert image to 32-bit"***
This node ensures that the image is a 32-bit floating point image.

- Input "Input" of node #13 receives data from output "Image" of node #11

***Node #14 "Morphological operation 2D"***

This node applies a morphological erosion with radius $\frac{2}{PixelSize}$. The node is configured to add a safety border of 50 px around the image prior to executing the erosion. This border is removed afterwards.

- Input "Input" of node #14 receives data from output "Image" of node #12
- The parameter "Add border before applying operation" (add-border) is set to **true**
- The parameter "Radius" (radius) is set to **10**
- The parameter "Operation" (operation) is set to **"EROSION"**
- The parameter item #1 of "Overridden parameters" in category "Adaptive parameters" (jipipe:adaptive-parameters/overridden-parameters) is set to **`(2 / NUM(PixelSize), "radius")`**
- The parameter "Margin right" in category "Border settings" (border-parameters/margin-right) is set to **50**
- The parameter "Margin left" in category "Border settings" (border-parameters/margin-left) is set to **50**
- The parameter "Margin top" in category "Border settings" (border-parameters/margin-top) is set to **50**
- The parameter "Margin bottom" in category "Border settings" (border-parameters/margin-bottom) is set to **50**

###### *Node #15 "Local variance 2D"*

This node applies a local variance filter with a radius $\frac{ExpectedDiameter\ /\ 6}{PixelSize}$.

- Input "Input" of node #15 receives data from output "Output" of node #13
- The parameter "Radius" (radius) is set to **40.0**
- The parameter item #1 of "Overridden parameters" in category "Adaptive parameters" (jipipe:adaptive-parameters/overridden-parameters) is set to **`((_global.expectedDiskDiameter / 6) / NUM(PixelSize), "radius")`**

###### *Node #16 "Remove annotation"*

This node removes all annotations except the image ID.

- Input "Input" of node #16 receives data from output "Output" of node #14
- The parameter "Removed annotations" (annotation-type) is set to **`key != "#ImageId"`**

###### *Node #17 "Set to value (grayscale)"*

This node sets all values outside the plate to 255.

- Input "Input" of node #17 receives data from output "Output" of node #15
- Input "Mask" of node #17 receives data from output "Output" of node #16
- The parameter "Only apply to ..." (roi:target-area) is set to **"OutsideMask"**
- The parameter "Value" (value) is set to **255.0**

###### *Node #18 "Invert (greyscale)" of type "Invert pixel values"*

This node inverts the image.

- Input "Input" of node #18 receives data from output "Output" of node #17

###### *Node #19 "Fast image arithmetics 2D"*

This node sets all pixel values that are equal to the minimum value to 255. Other values are set to zero.

- Input "I1" of node #19 receives data from output "Output" of node #17
- The parameter "Expression" (expression) is set to **`I1 == SLICE_MIN(I1)`**
- The parameter "Bit depth" (bit-depth) is set to **"Grayscale8u"**

###### *Node #20 "Auto threshold 2D"*

This node applies the “Default” auto thresholding method from ImageJ.

- Input "Input" of node #20 receives data from output "Output" of node #18

###### *Node #21 "Remap labels"*

This node ensures that label values are consecutive and start with 1.

- Input "Input" of node #21 receives data from output "Output" of node #19

###### *Node #22 "Connected components labeling 2D"*

This node converts a mask into a label image.

- Input "Input" of node #22 receives data from output "Output" of node #20

###### *Node #23 "Filter Labels by overlap 2D"*

This node selects only the labels generated from the thresholding that overlap with the DDA center.

- Input "Labels 1" of node #23 receives data from output "Output" of node #21
- Input "Labels 2" of node #23 receives data from output "Output" of node #22
- The parameter "Overlap filter measurements" (overlap-filter-measurements) is set to **{"values":[],"collapsed":false}**
- The parameter "Enabled" in category "Labels 1 filter" (labels1/enabled) is set to **false**

###### *Node #24 "Binarize"*

This node converts the label image into a binary image.

- Input "Input" of node #24 receives data from output "Labels 2" of node #23

###### *Node #25 "Find particles 2D"*

This node converts a binary mask into ImageJ ROI lists.

- Input "Mask" of node #25 receives data from output "Output" of node #24

###### *Node #26 "Outline 2D ROI (Hull): Fit circle" of type "Outline 2D ROI (Hull)"*

This node applies a circle fitting to the incoming ROIs.

- Input "Input" of node #26 receives data from output "ROI" of node #25
- The parameter "Outline method" (outline) is set to **"FitCircle"**

###### *Node #27 "Scale 2D ROI" of type "Transform 2D ROI"*

This node scales the circular ROIs to fit the expected DDA size.

- Input "Input" of node #27 receives data from output "Output" of node #26

- The parameter "Measure in physical units" (measure-in-physical-units) is set to **false**
- The parameter "Scale X" (scale-x) is set to **`_global.expectedDiskDiameter / (NUM(PixelSize) * Feret)`**
- The parameter "Scale Y" (scale-y) is set to **`_global.expectedDiskDiameter / (NUM(PixelSize) * Feret)`**
- The parameter "Measurements" (measurements) is set to **{"values":["FeretDiameter","BoundingRectangle"],"collapsed":false}**

##### *Node #28 "Convert 2D ROI to mask"*

This node converts ROIs into a binary mask.

- Input "ROI" of node #28 receives data from output "Output" of node #27
- Input "Image" of node #28 receives data from output "Output" of node #24

##### *Node #29 "Export image"*

This node exports the generated mask.

- Input "Input" of node #29 receives data from output "Output" of node #28
- The parameter "File path" (file-path) is set to **`PATH_COMBINE(project_data_dir.tmp_dir, "strip-disk", #ImageId)`**

#### 2.3.4 Variant 3

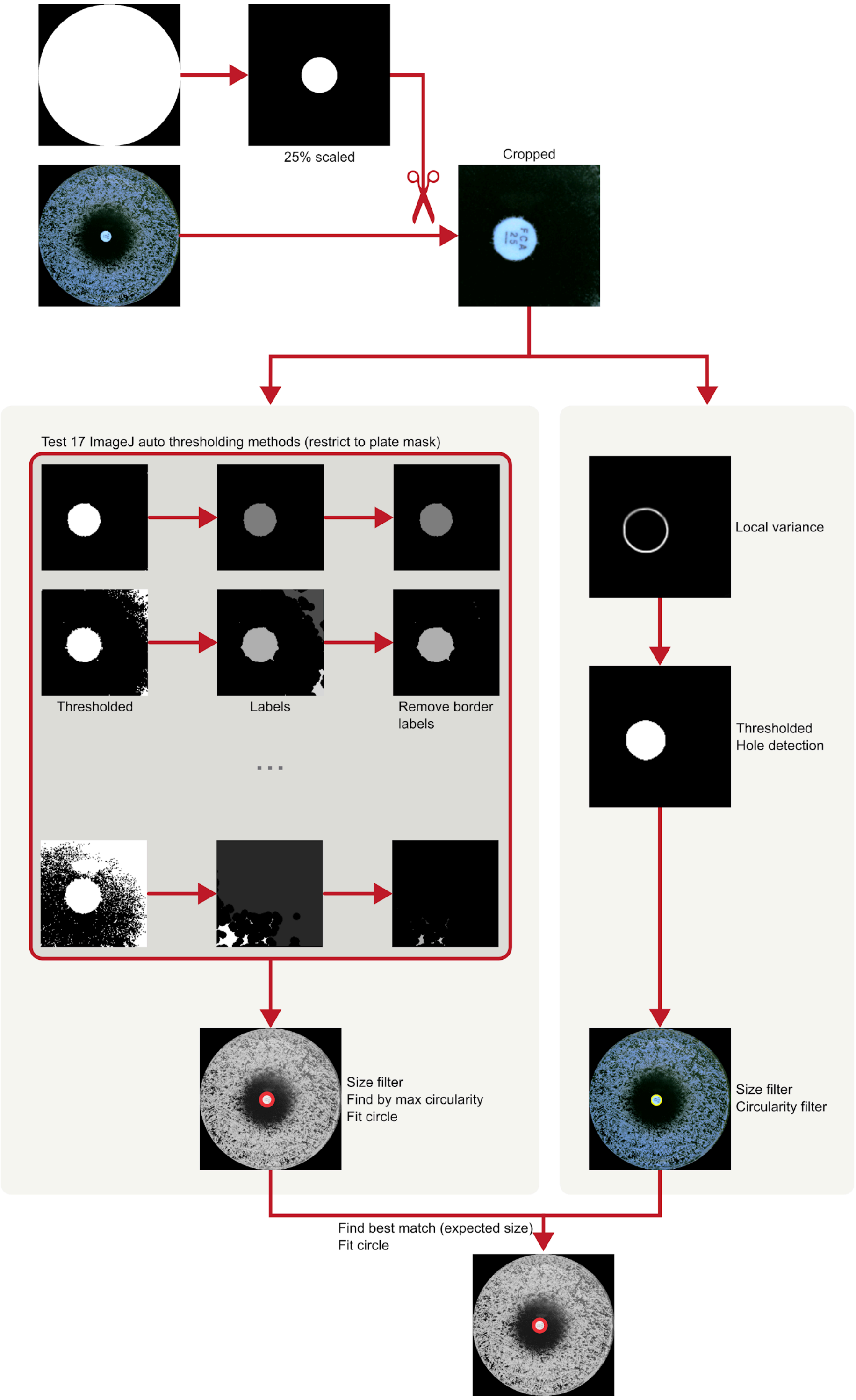

**Supplementary Figure 16:** Third variant of the DDA disk detection. The plate mask is scaled to 25% of its size and used to extract only the central part of the image. The algorithm is split into two branches (grey boxes). The first branch tests a set of operations per ImageJ auto-thresholding methods to generate candidates for DDA disks. The resulting objects are filtered and the most circular ROI is selected. In the second branch the local variance is used to highlight the edge of the disk. This is followed by an auto-thresholding and hole detection, as well as additional filtering. The results from both branches are compared and the object with the size closest to the expected value is selected.

The following steps are applied by the third algorithm that finds the disk region inside DDA. An overview can be found in **Supplementary Figure 16** and a screenshot is provided in **Supplementary Figure 17**.

##### *Node #1 "File"*

This node references the CSV table that associates the image ID to its metadata.

- The parameter "File name" (file-name) is set to **"metadata.csv"**

##### *Node #2 "Folder list"*

This node references the directory containing the raw images.

- The parameter item #1 of "Folder paths" (folder-paths) is set to **"raw"**

##### *Node #3 "Folder list"*

This node references the directory that contains the plate masks.

- The parameter item #1 of "Folder paths" (folder-paths) is set to **"plate"**

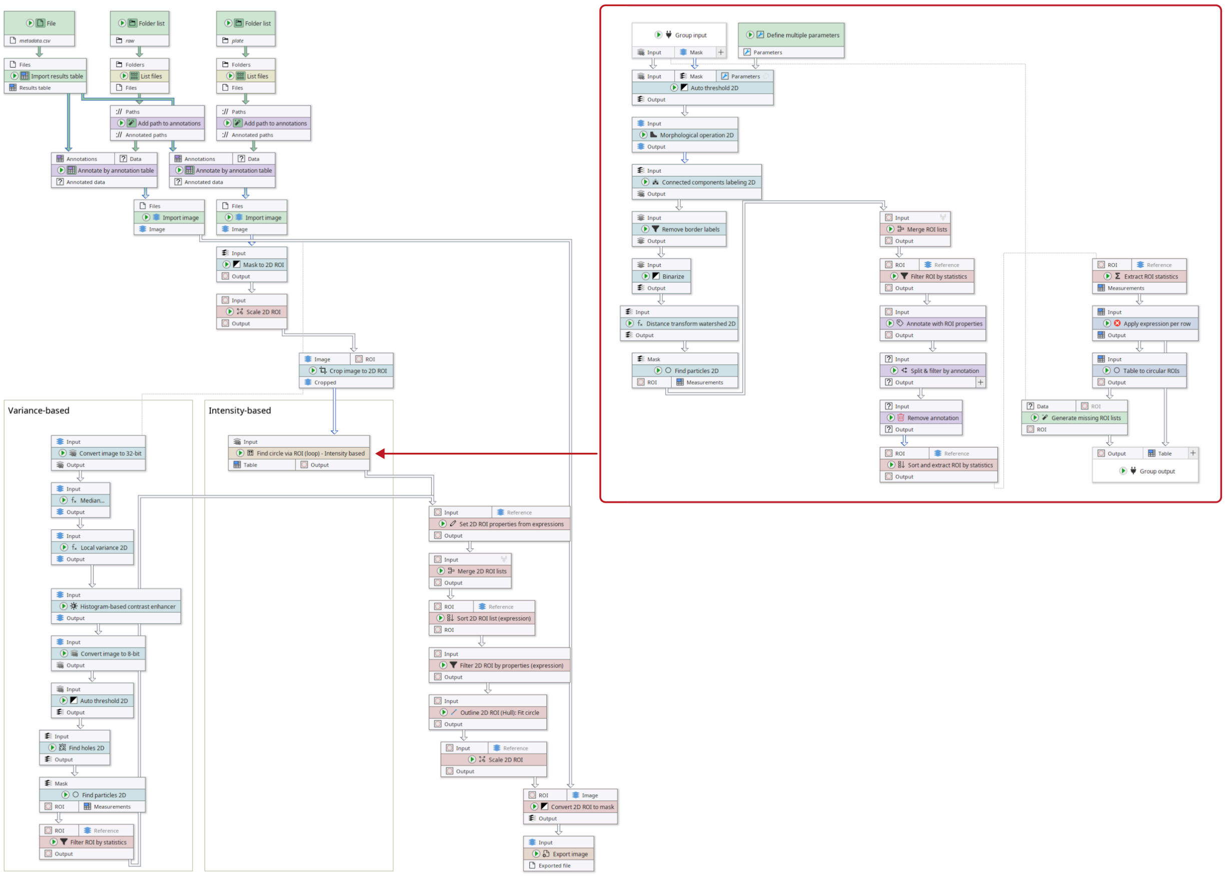

**Supplementary Figure 17:** Screenshot of the DDA disk detection pipeline (Variant 3). The pipeline contains a group node that has an embedded pipeline (red box).

*Node #4 "Import results table"*

This node imports the metadata CSV file as an ImageJ table.

- Input "Files" of node #4 receives data from output "Filename" of node #1

*Node #5 "List files"*

This node lists all PNG files in the input directory.

- Input "Folders" of node #5 receives data from output "Folder paths" of node #2
- The parameter "Keep file if ..." (filters) is set to **`STRING_MATCHES_GLOB(name, "*.png")`**

*Node #6 "List files"*

This node lists all PNG files in the input directory.

- Input "Folders" of node #6 receives data from output "Folder paths" of node #3
- The parameter "Keep file if ..." (filters) is set to **`STRING_MATCHES_GLOB(name, "*.png")`**

*Node #7 "Add path to annotations" of type "Annotate with file/directory name"*

This node attaches the image ID to the image file.

- Input "Paths" of node #7 receives data from output "Files" of node #5
- The parameter "Generated annotation" (generated-annotation) is set to **"#ImageId"**

*Node #8 "Add path to annotations" of type "Annotate with file/directory name"*

This node attaches the image ID to the image file.

- Input "Paths" of node #8 receives data from output "Files" of node #6
- The parameter "Generated annotation" (generated-annotation) is set to **"#ImageId"**

*Node #9 "Annotate by annotation table"*

This node attaches metadata from the metadata table to the image file.

- Input "Annotations" of node #9 receives data from output "Results table" of node #4
- Input "Data" of node #9 receives data from output "Annotated paths" of node #7

*Node #10 "Annotate by annotation table"*

This node attaches metadata from the metadata table to the image file.

- Input "Annotations" of node #10 receives data from output "Results table" of node #4
- Input "Data" of node #10 receives data from output "Annotated paths" of node #8

*Node #11 "Import image"*

This node imports an image file as ImageJ image.

- Input "Files" of node #11 receives data from output "Annotated data" of node #9

*Node #12 "Import image"*

This node imports an image file as ImageJ image.

- Input "Files" of node #12 receives data from output "Annotated data" of node #10

***Node #13 "Mask to 2D ROI"***
This node converts the plate mask annotation into an ImageJ ROI list.

- Input "Input" of node #13 receives data from output "Image" of node #12

***Node #14 "Scale 2D ROI" of type "Scale 2D ROI (old)"***
This node scales the plate ROI down to 25% of its size.

- Input "Input" of node #14 receives data from output "Output" of node #13
- The parameter "Scale X" (scale-x) is set to **0.25**
- The parameter "Scale Y" (scale-y) is set to **0.25**
- The parameter "Center scale" (center-scale) is set to **true**

***Node #15 "Crop image to 2D ROI"***
This node crops the raw image to the bounding box of the scaled plate ROI.

- Input "Image" of node #15 receives data from output "Image" of node #11
- Input "ROI" of node #15 receives data from output "Output" of node #14
- The parameter "Crop channel plane" (crop-c) is set to **false**
- The parameter "Crop Z plane" (crop-z) is set to **false**
- The parameter "Annotate with Y location" (annotation-y) is set to **"crop.y"**
- The parameter "Annotate with X location" (annotation-x) is set to **"crop.x"**
- The parameter "Crop frame plane" (crop-t) is set to **false**

***Node #16 "Find circle via ROI (loop) - Intensity based" of type "Group"***
This node is a group responsible for finding the DDA disk using intensity thresholding. The nodes are separated from the outside pipeline for organizational purposes.

- Input "Input" of node #16 receives data from output "Cropped" of node #15
- This node contains a sub-graph. The "Group Input" and "Group Output" nodes contained in the graph are internally connected to the inputs and outputs of this node.

***Node #17 "Define multiple parameters"***
This node defines parameter sets that set up the application of multiple ImageJ thresholding methods.

- The parameter "Parameters" (parameter-table) is set to contain the methods **Default, Huang, MaxEntropy, Mean, Minimum, Otsu, Percentile, Yen, Shanbhag, Intermodes, Triangle, MinError, Moments, IsoData, IJ_IsoData, RenyiEntropy,** and **Li**

***Node #18 "Group input"***
This node acts as interface from the outer pipeline to the group (node #16).

***Node #19 "Auto threshold 2D"***
This node applies ImageJ auto-thresholding. The node receives external parameters from node #17, thus repeating its workload for each parameter set.

- Input "Input" of node #19 receives data from output "Input" of node #18

- Input "{Parameters}" of node #19 receives data from output "Parameters" of node #17
- The parameter "Multiple parameters" in category "Multi-parameter settings" (jipipe:parameter-slot-algorithm/has-parameter-slot) is set to **true**
- The parameter "Attach parameter annotations" in category "Multi-parameter settings" (jipipe:parameter-slot-algorithm/attach-parameter-annotations) is set to **false**

###### *Node #20 "Morphological operation 2D"*

This node applies a morphological closing operation with a radius of 5 pixels.

- Input "Input" of node #20 receives data from output "Output" of node #19
- The parameter "Radius" (radius) is set to **5**
- The parameter "Operation" (operation) is set to **"CLOSING"**

###### *Node #21 "Connected components labeling 2D"*

This node converts a binary mask into a label image.

- Input "Input" of node #21 receives data from output "Output" of node #20

###### *Node #22 "Remove border labels"*

This node removes all labels that touch the image border.

- Input "Input" of node #22 receives data from output "Output" of node #21

###### *Node #23 "Binarize"*

This node converts the label image into a binary image.

- Input "Input" of node #23 receives data from output "Output" of node #22

###### *Node #24 "Distance transform watershed 2D"*

This node applies distance transform watershed to remove holes.

- Input "Input" of node #24 receives data from output "Output" of node #23

###### *Node #25 "Find particles 2D"*

This node uses the ImageJ particle finder to convert masks into ROI lists.

- Input "Mask" of node #25 receives data from output "Output" of node #24

###### *Node #26 "Merge ROI lists" of type "Merge 2D ROI lists"*

This node merges the per-thresholding-method ROI lists into a single list.

- Input "Input" of node #26 receives data from output "ROI" of node #25

###### *Node #27 "Filter ROI by statistics" of type "Filter 2D ROI by statistics"*

This node applies a size and circularity filter to the incoming ROI lists.

- Input "ROI" of node #27 receives data from output "Output" of node #26
- The parameter "Keep ROI if" (filter) is set to **`SET_VARIABLE("expectedPx", _global.expectedDiskDiameter / NUM(PixelSize)); SET_VARIABLE("expectedPxDiff", ABS((Feret - expectedPx) / expectedPx)); expectedPxDiff <=`**

`_global.expectedDiskDiameterMaxDiffPerc AND Circ. >= _global.minCirc`

- The parameter "Measurements" (measurements) is set to **{"values":["FeretDiameter","ShapeDescriptors"],"collapsed":false}**

##### *Node #28 "Annotate with ROI properties" of type "Annotate with 2D ROI properties"*

This node annotates each ROI list with the ROI count.

- Input "Input" of node #28 receives data from output "Output" of node #27

##### *Node #29 "Split & filter by annotation"*

This node filters out all empty ROI lists.

- Input "Input" of node #29 receives data from output "Output" of node #28
- The parameter "Output" in category "Filters" (target-slots/Output) is set to **TO_NUMBER(Count) > 0**

##### *Node #30 "Remove annotation"*

This node removes the annotation added by node #28.

- Input "Input" of node #30 receives data from output "Output" of node #29
- The parameter "Removed annotations" (annotation-type) is set to **"Count"**

##### *Node #31 "Sort and extract ROI by statistics" of type "Sort and extract 2D ROI by statistics"*

This node selects the ROI with the highest circularity.

- Input "ROI" of node #31 receives data from output "Output" of node #30
- The parameter "Selected indices" (selected-indices) is set to **0**
- The parameter item #1 of "Sort order" (sort-orders) is set to **("ShapeDescriptorCircularity", "Descending")**

##### *Node #32 "Extract ROI statistics" of type "Extract 2D ROI statistics"*

This node extracts the centroid and Feret diameter of the input ROI.

- Input "ROI" of node #32 receives data from output "Output" of node #31
- The parameter "Extracted measurements" (measurements) is set to **{"values":["Centroid","FeretDiameter"],"collapsed":false}**

##### *Node #33 "Apply expression per row"*

This node calculates the radius from the Feret diameter.

- Input "Input" of node #33 receives data from output "Measurements" of node #32
- The parameter item #1 of "Expressions" (expression-list) is set to `(Feret / 2, "Radius")`

##### *Node #34 "Table to circular ROIs" of type "Table to 2D circular ROI"*

This node draws a circular ROI based on the information contained within the input table.

- Input "Input" of node #34 receives data from output "Output" of node #33

- The parameter "ROI name" in category "ROI properties" (roi-properties/roi-name) is set to **"unnamed"**

***Node #35 "Generate missing ROI lists" of type "Generate missing 2D ROI lists"***

This node generates empty ROI lists if a ROI list is missing.

- Input "Data" of node #35 receives data from output "Input" of node #18
- Input "ROI" of node #35 receives data from output "Output" of node #34

***Node #36 "Group output"***

This node transfers the final ROI lists out of the group into the surrounding pipeline.

- Input "Output" of node #36 receives data from output "ROI" of node #35
- Input "Table" of node #36 receives data from output "Output" of node #33

***Node #37 "Convert image to 32-bit"***

This node ensures that the image is a 32-bit floating point greyscale image.

- Input "Input" of node #37 receives data from output "Cropped" of node #15

***Node #38 "Median..." of type "Median filter 2D"***

This node applies a median filter with radius 3 px.

- Input "Input" of node #38 receives data from output "Output" of node #37
- The parameter "Radius" (radius) is set to **3.0**

***Node #39 "Local variance 2D"***

This node applies a local variance filter with radius $\frac{ExpectedDiameter\ /\ 6}{PixelSize}$.

- Input "Input" of node #39 receives data from output "Output" of node #38
- The parameter "Radius" (radius) is set to **40.0**
- The parameter item #1 of "Overridden parameters" in category "Adaptive parameters" (jipipe:adaptive-parameters/overridden-parameters) is set to **`((_global.expectedDiskDiameter / 6) / NUM(PixelSize), "radius")`**

***Node #40 "Histogram-based contrast enhancer"***

This node enhances the image contrast.

- Input "Input" of node #40 receives data from output "Output" of node #39

***Node #41 "Convert image to 8-bit"***

This node ensures that the image is an 8-bit greyscale image.

- Input "Input" of node #41 receives data from output "Output" of node #40

***Node #42 "Auto threshold 2D"***

This node applies a “Default” ImageJ auto-thresholding.

- Input "Input" of node #42 receives data from output "Output" of node #41

***Node #43 "Find holes 2D"***

This node detects holes using a flood-filling approach.

- Input "Input" of node #43 receives data from output "Output" of node #42

***Node #44 "Find particles 2D"***

This node uses the ImageJ particle finder to convert the incoming mask into a ROI list.

- Input "Mask" of node #44 receives data from output "Output" of node #43

***Node #45 "Filter ROI by statistics" of type "Filter 2D ROI by statistics"***

This node applies a size and circularity filter to the incoming ROI.

- Input "ROI" of node #45 receives data from output "ROI" of node #44
- The parameter "Keep ROI if" (filter) is set to **`SET_VARIABLE("expectedPx", _global.expectedDiskDiameter / NUM(PixelSize)); SET_VARIABLE("expectedPxDiff", ABS((Feret - expectedPx) / expectedPx)); expectedPxDiff <= _global.expectedDiskDiameterMaxDiffPerc AND Circ. >= _global.minCirc`**
- The parameter "Measurements" (measurements) is set to **{"values":["FeretDiameter","ShapeDescriptors"],"collapsed":false}**

***Node #46 "Set 2D ROI properties from expressions"***

This node moves the ROI to the correct global position, which is needed due to the earlier cropping operation.

- Input "Input" of node #46 receives data from output "Output" of node #16
- Input "Input" of node #46 receives data from output "Output" of node #45
- The parameter "Location (X)" (position-x) is set to **`x + NUM(crop.x)`**
- The parameter "Location (Y)" (position-y) is set to **`y + NUM(crop.y)`**

***Node #47 "Merge 2D ROI lists"***

This node merges all DDA disk candidate ROIs into one list.

- Input "Input" of node #47 receives data from output "Output" of node #46

***Node #48 "Sort 2D ROI list (expression)"***

This node sorts the ROIs by the difference of their size to the expected DDA disk size.

- Input "ROI" of node #48 receives data from output "Output" of node #47
- The parameter "Expression" (expression) is set to **`SET_VARIABLE("expectedPx", _global.expectedDiskDiameter / NUM(PixelSize)); SET_VARIABLE("expectedPxDiff", ABS((Feret - expectedPx) / expectedPx)); expectedPxDiff`**

***Node #49 "Filter 2D ROI by properties (expression)"***

This node selects the first ROI, which will be the one with a diameter closest to the expected size.

- Input "Input" of node #49 receives data from output "ROI" of node #48
- The parameter "Keep ROI if" (filter) is set to **`index == 0`**

##### *Node #50 "Outline 2D ROI (Hull): Fit circle" of type "Outline 2D ROI (Hull)"*

This node applies an additional circle fitting.

- Input "Input" of node #50 receives data from output "Output" of node #49
- The parameter "Outline method" (outline) is set to **"FitCircle"**

##### *Node #51 "Scale 2D ROI" of type "Transform 2D ROI"*

This node ensures that the ROIs have the expected size.

- Input "Input" of node #51 receives data from output "Output" of node #50
- The parameter "Measure in physical units" (measure-in-physical-units) is set to **false**
- The parameter "Scale X" (scale-x) is set to **`_global.expectedDiskDiameter / (NUM(PixelSize) * Feret)`**
- The parameter "Scale Y" (scale-y) is set to **`_global.expectedDiskDiameter / (NUM(PixelSize) * Feret)`**
- The parameter "Measurements" (measurements) is set to **{"values":["FeretDiameter","BoundingRectangle"],"collapsed":false}**

##### *Node #52 "Convert 2D ROI to mask"*

This node converts the ImageJ ROIs into masks.

- Input "ROI" of node #52 receives data from output "Output" of node #51
- Input "Image" of node #52 receives data from output "Image" of node #11

##### *Node #53 "Export image"*

This node exports the masks into PNG files.

- Input "Input" of node #53 receives data from output "Output" of node #52
- The parameter "File path" (file-path) is set to **`PATH_COMBINE(project_data_dir.tmp_dir, "strip-disk", #ImageId)`**

#### 2.4 Etest strip detection

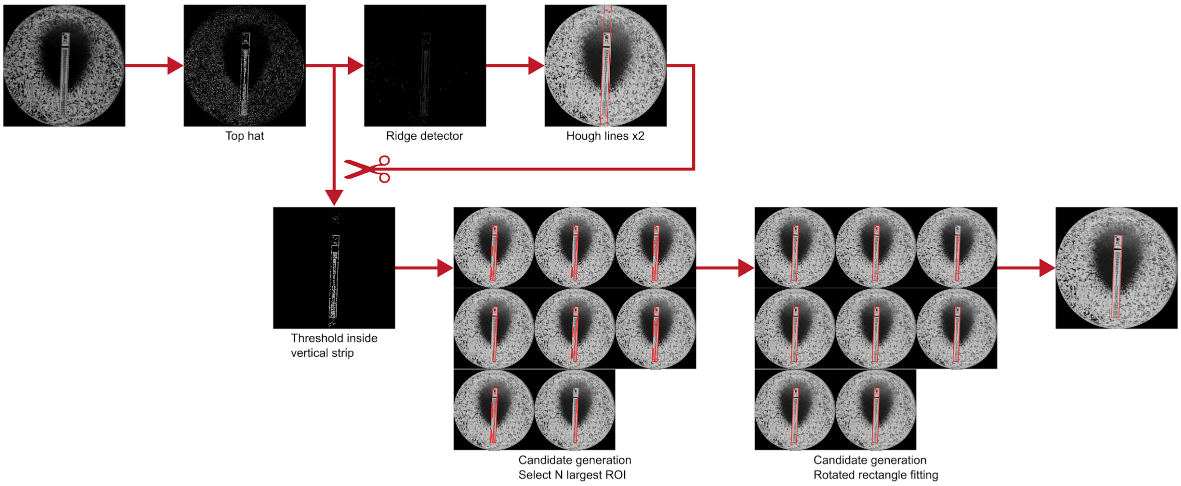

**Supplementary Figure 18:** Etest strip detection algorithm. Edges are first highlighted using a morphological top hat operation and detected using the ImageJ Ridge Detector. Two Hough line detection operations find the major sides of the strip that are used to apply an auto-thresholding on the top hat image exclusively to the detected area. Candidate strip ROI are generated by iteratively generating ROI that are composed of the N largest objects, followed by a rotated rectangle fitting. The candidate that matches the expected aspect ratio is selected.

The following steps are applied by the third algorithm that finds the strip region inside Etests. An overview can be found in **Supplementary Figure 18** and a screenshot is provided in **Supplementary Figure 19**.

*Node #1 "Folder list"*

This node references the directory that contains the raw images.

- The parameter item #1 of "Folder paths" (folder-paths) is set to **"raw"**

*Node #2 "Folder list"*

This node references the directory that contains the plate annotation masks.

- The parameter item #1 of "Folder paths" (folder-paths) is set to **"plate"**

*Node #3 "File"*

This node references a CSV file that associates the image ID to the image metadata.

- The parameter "File name" (file-name) is set to **"metadata.csv"**

*Node #4 "Define multiple parameters"*

This node creates the following parameter sets that will be used in another node:

| Selected indices |
|---|
| 0 |
| 0; 1 |
| 0; 1; 2 |
| 0; 1; 2; 3 |
| 0; 1; 2; 3; 4 |
| 0; 1; 2; 3; 4; 5 |
| 0; 1; 2; 3; 4; 5; 6 |
| 0; 1; 2; 3; 4; 5; 6; 7 |

*Node #5 "List files"*

This node lists all PNG files in the input directory.

- Input "Folders" of node #5 receives data from output "Folder paths" of node #1
- The parameter "Keep file if ..." (filters) is set to `STRING_MATCHES_GLOB(name, "*.png")`

***Node #6 "List files"***

This node lists all PNG files in the input directory.

- Input "Folders" of node #6 receives data from output "Folder paths" of node #2
- The parameter "Keep file if ..." (filters) is set to **`STRING_MATCHES_GLOB(name, "*.png")`**

***Node #7 "Import results table"***

This node imports the metadata CSV file as an ImageJ table.

- Input "Files" of node #7 receives data from output "Filename" of node #3

***Node #8 "Add path to annotations" of type "Annotate with file/directory name"***

This node attaches the image ID to the incoming file.

- Input "Paths" of node #8 receives data from output "Files" of node #5
- The parameter "Generated annotation" (generated-annotation) is set to **"#ImageId"**

***Node #9 "Add path to annotations" of type "Annotate with file/directory name"***

This node attaches the image ID to the incoming file.

- Input "Paths" of node #9 receives data from output "Files" of node #6
- The parameter "Generated annotation" (generated-annotation) is set to **"#ImageId"**

***Node #10 "Annotate by annotation table" of type "Annotate by annotation table"***

This node attaches metadata from the metadata table to the currently processed file.

- Input "Annotations" of node #10 receives data from output "Results table" of node #7
- Input "Data" of node #10 receives data from output "Annotated paths" of node #8

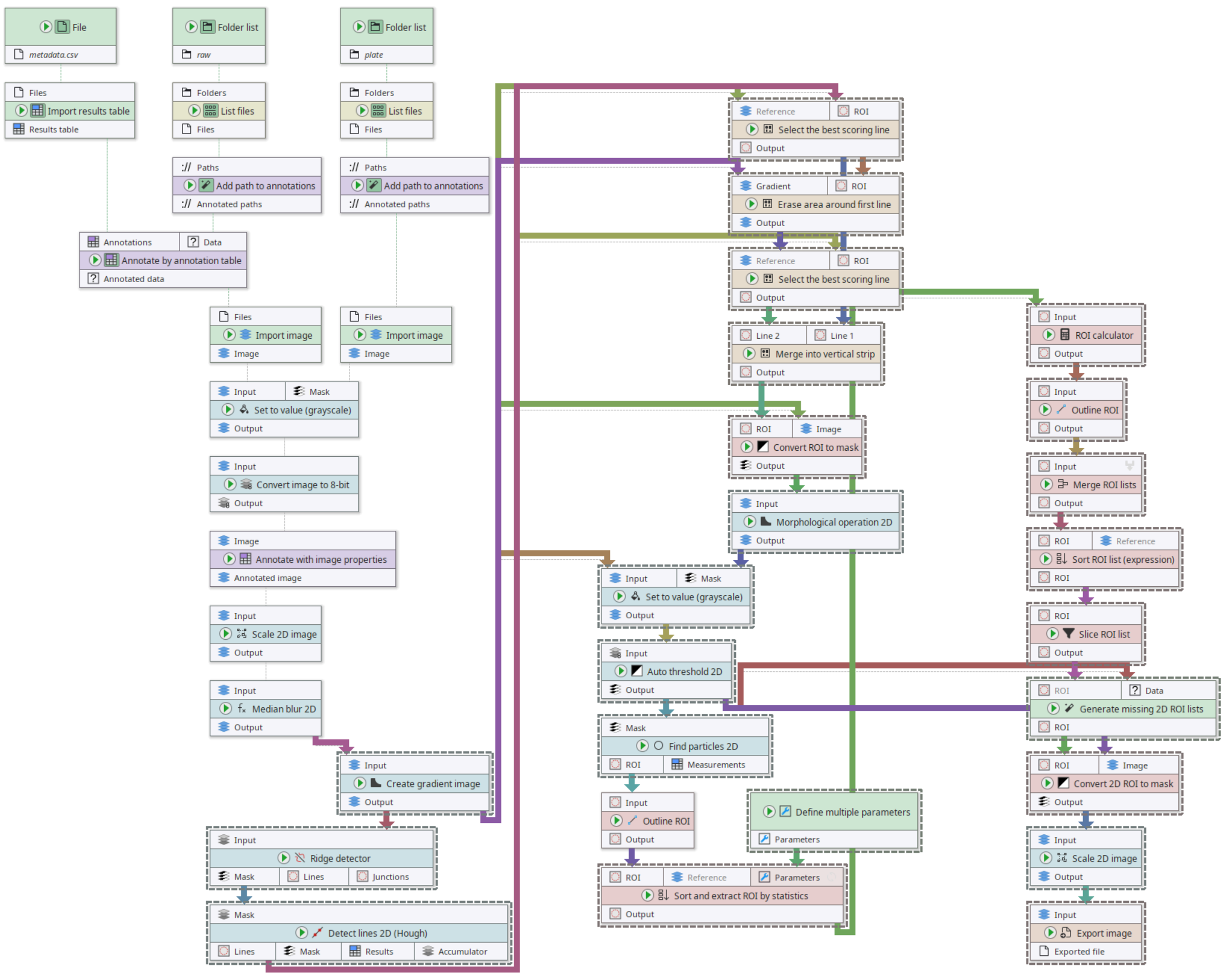

**Supplementary Figure 19:** Full Etest strip detection algorithm pipeline. We selected a few nodes to allow for a better visualization of the connections.

##### *Node #11 "Import image"*

This node imports an image file as ImageJ image.

- Input "Files" of node #11 receives data from output "Annotated paths" of node #9

##### *Node #12 "Import image"*

This node imports an image file as ImageJ image.

- Input "Files" of node #12 receives data from output "Annotated data" of node #10

##### *Node #13 "Set to value (grayscale)"*

This node sets the image areas outside the plate mask to zero.

- Input "Input" of node #13 receives data from output "Image" of node #12
- Input "Mask" of node #13 receives data from output "Image" of node #11
- The parameter "Only apply to ..." (roi:target-area) is set to **"OutsideMask"**

##### *Node #14 "Convert image to 8-bit"*

This node ensures that an image is an 8-bit greyscale image.

- Input "Input" of node #14 receives data from output "Output" of node #13

##### *Node #15 "Annotate with image properties"*

This node attaches the image width and height to the currently processed image in the form of an annotation.

- Input "Image" of node #15 receives data from output "Output" of node #14
- The parameter "Annotate with image height" (height-annotation) is set to **"original_img_h"**
- The parameter "Annotate with image width" (width-annotation) is set to **"original_img_w"**

##### *Node #16 "Scale 2D image"*

This node scales the incoming images to 2048x2048 pixels.

- Input "Input" of node #16 receives data from output "Annotated image" of node #15
- The parameter "Y axis" (y-axis) is set to **2048**
- The parameter "X axis" (x-axis) is set to **2048**
- The parameter "Interpolation" (interpolation-method) is set to **"None"**
- The parameter "Limit" in category "Input management" (jipipe:data-batch-generation/limit) is set to **[Disabled]**

##### *Node #17 "Median blur 2D" of type “Median filter 2D”*

This node applies a median filter with radius 1 px.

- Input "Input" of node #17 receives data from output "Output" of node #16

##### *Node #18 "Create gradient image" of type "Morphological operation 2D"*

This node applies a morphological bottom hat operation with a radius of 5 px.

- Input "Input" of node #18 receives data from output "Output" of node #17
- The parameter "Add border before applying operation" (add-border) is set to **true**
- The parameter "Radius" (radius) is set to **5**
- The parameter "Operation" (operation) is set to **"BOTTOMHAT"**

##### *Node #19 "Ridge detector"*

This node applies an ImageJ ridge detector.

- Input "Input" of node #19 receives data from output "Output" of node #18

##### *Node #20 "Detect lines 2D (Hough)" of type "Detect global lines 2D (Hough, legacy)"*

This node applies a Hough line detection algorithm.

- Input "Mask" of node #20 receives data from output "Mask" of node #19

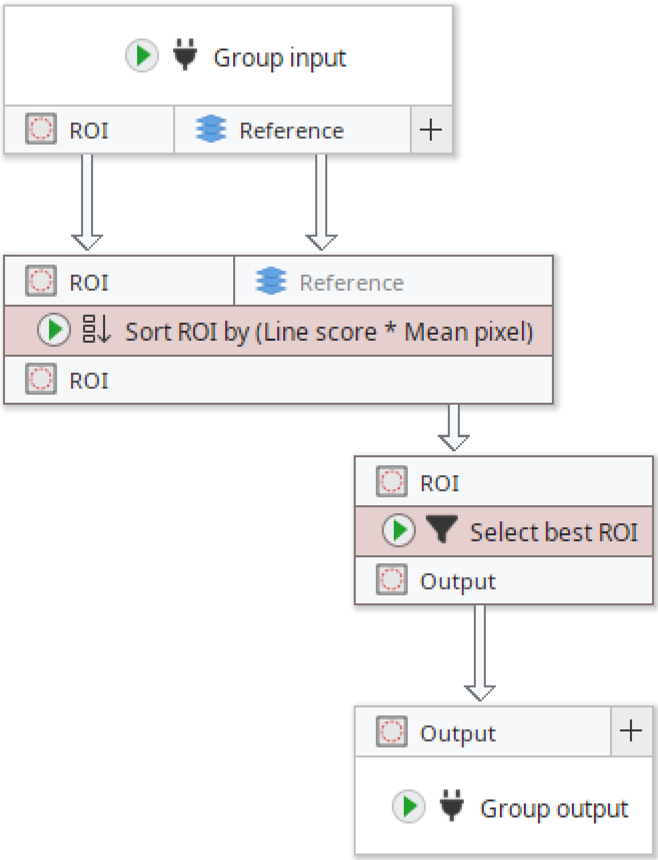

**Supplementary Figure 20:** Full Etest strip detection algorithm pipeline; group for selecting best-scoring line.

##### *Node #21 "Select the best scoring line" of type "Group"*

This is a group node that contains a sub-pipeline that finds the line with the highest score. See **Supplementary Figure 20** for a screenshot.

- Input "Reference" of node #21 receives data from output "Output" of node #18
- Input "ROI" of node #21 receives data from output "Lines" of node #20
- This node contains a sub-graph. The "Group Input" and "Group Output" nodes contained in the graph are internally connected to the inputs and outputs of this node.

##### *Node #22 "Group input"*

This node acts as an interface for the group node #21 where inputs are passed through.

##### *Node #23 "Sort ROI by (Line score * Mean pixel)" of type "Sort 2D ROI list (expression)"*

This node sorts the line ROI extracted by the Hough transform by the mean intensity on the gradient image.

- Input "ROI" of node #23 receives data from output "ROI" of node #22
- Input "Reference" of node #23 receives data from output "Reference" of node #22
- The parameter "Expression" (expression) is set to `TO_NUMBER(Name) * Mean`
- The parameter "Reverse sort order" (reverse-sort-order) is set to **true**

##### *Node #24 "Select best ROI" of type "Slice 2D ROI list"*

This node selects the first line ROI from the output of node #23, which yields the brightest line.

- Input "ROI" of node #24 receives data from output "ROI" of node #23
- The parameter "Selected indices" (selected-indices) is set to **0**

##### *Node #25 "Group output"*

This node acts as an interface to pass the detected line ROI to the outer pipeline.

- Input "Output" of node #25 receives data from output "Output" of node #24

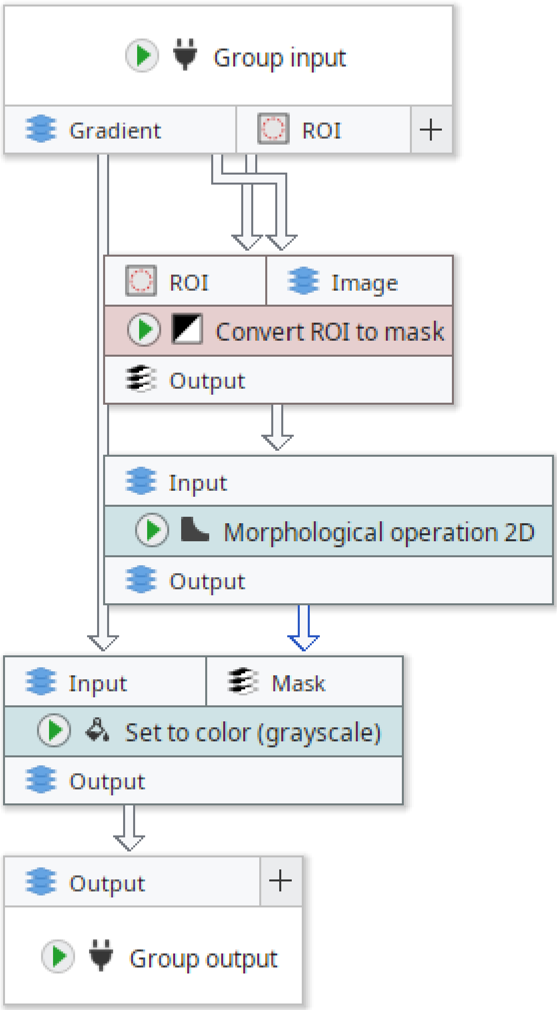

**Supplementary Figure 21:** Full Etest strip detection algorithm pipeline; group for erasing the area around a line.

##### *Node #26 "Erase area around first line" of type "Group"*

This node is a group that is responsible for erasing the line detected by node #21. See **Supplementary Figure 21** for a screenshot.

- Input "Gradient" of node #26 receives data from output "Output" of node #18
- Input "ROI" of node #26 receives data from output "Output" of node #21
- This node contains a sub-graph. The "Group Input" and "Group Output" nodes contained in the graph are internally connected to the inputs and outputs of this node.

***Node #27 "Group input"***

This node acts as interface for the inputs of node #26.

***Node #28 "Convert ROI to mask" of type "Convert 2D ROI to mask"***

This node converts the line ROI into a mask.

- Input "ROI" of node #28 receives data from output "ROI" of node #27
- Input "Image" of node #28 receives data from output "Gradient" of node #27

***Node #29 "Morphological operation 2D"***

This node dilates the mask generated by node #28 by a radius of 100.

- Input "Input" of node #29 receives data from output "Output" of node #28
- The parameter "Radius" (radius) is set to **100**

***Node #30 "Set to color (grayscale)" of type "Set to value (grayscale)"***

This node sets the value of the gradient image defined by the mask generated by node #29 to zero, thus erasing the first line.

- Input "Input" of node #30 receives data from output "Gradient" of node #27
- Input "Mask" of node #30 receives data from output "Output" of node #29
- The parameter "Only apply to ..." (roi:target-area) is set to **"InsideMask"**

***Node #31 "Group output"***

This node passes the modified gradient image back into the outer pipeline.

- Input "Output" of node #31 receives data from output "Output" of node #30

***Node #32 "Select the best scoring line" of type "Group"***

This is a group node that contains a sub-pipeline that finds the line with the highest score. See **Supplementary Figure 17** for a screenshot.

- Input "Reference" of node #32 receives data from output "Output" of node #26
- Input "ROI" of node #32 receives data from output "Lines" of node #20
- This node contains a sub-graph. The "Group Input" and "Group Output" nodes contained in the graph are internally connected to the inputs and outputs of this node.

***Node #22 "Group input"***

This node acts as an interface for the input data.

***Node #23 "Sort ROI by (Line score * Mean pixel)" of type "Sort 2D ROI list (expression)"***

This node sorts the line ROI extracted by the Hough transform by the mean intensity on the gradient image.

- Input "ROI" of node #23 receives data from output "ROI" of node #22
- Input "Reference" of node #23 receives data from output "Reference" of node #22
- The parameter "Expression" (expression) is set to **`TO_NUMBER(Name) * Mean`**
- The parameter "Reverse sort order" (reverse-sort-order) is set to **true**

***Node #24 "Select best ROI" of type "Slice 2D ROI list"***

This node selects the first line ROI from the output of node #23, which yields the brightest line.

- Input "ROI" of node #24 receives data from output "ROI" of node #23
- The parameter "Selected indices" (selected-indices) is set to **0**

***Node #25 "Group output"***

This node passes the ROI to the outer pipeline.

- Input "Output" of node #25 receives data from output "Output" of node #24

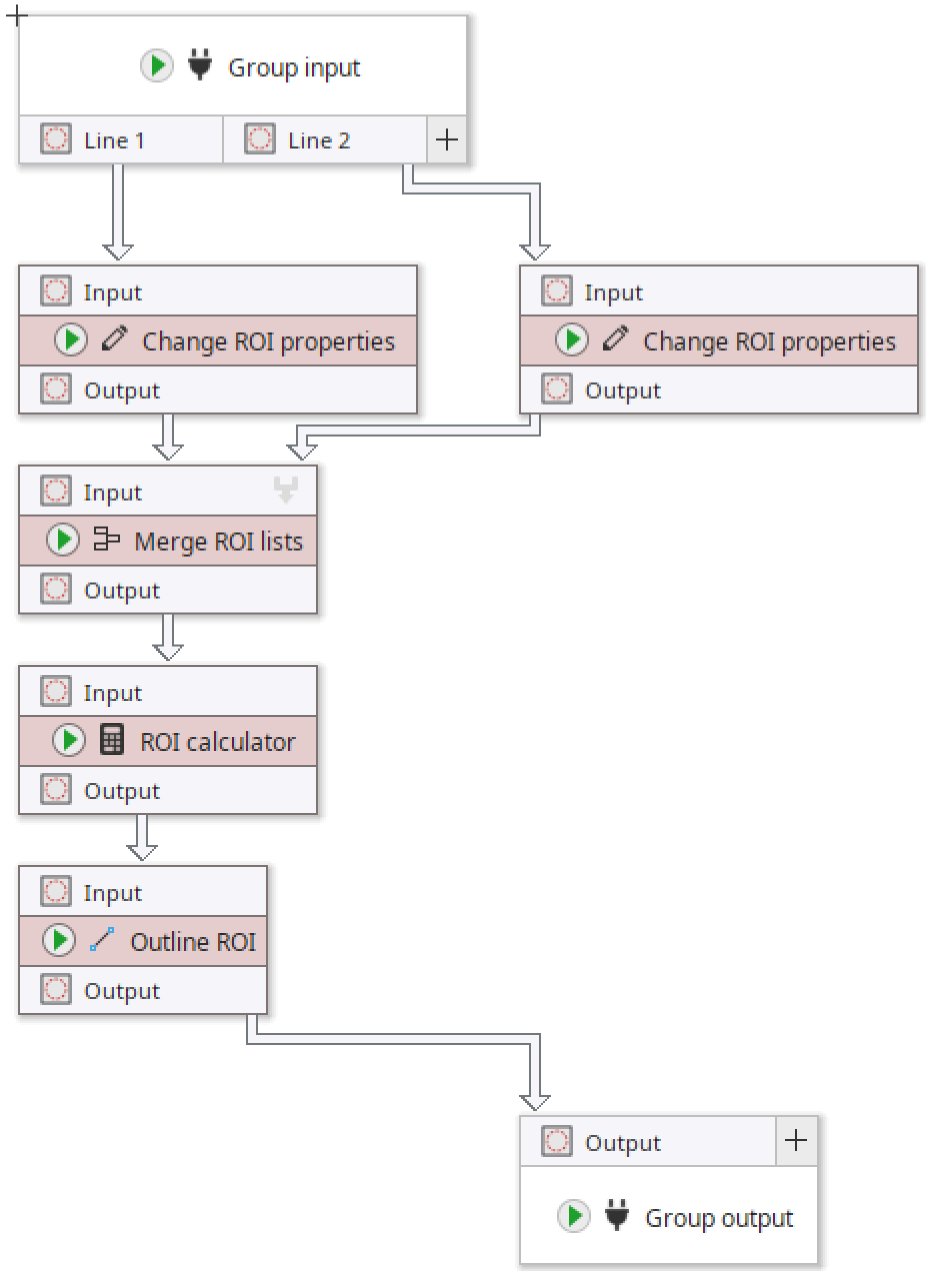

**Supplementary Figure 22:** Full Etest strip detection algorithm pipeline; group for creating vertical strip candidates.

***Node #33 "Merge into vertical strip" of type "Group"***

This node is a group that generates ROIs that will be later assembled into the final strip. A screenshot can be found in **Supplementary Figure 22**.

- Input "Line 2" of node #33 receives data from output "Output" of node #32
- Input "Line 1" of node #33 receives data from output "Output" of node #21
- This node contains a sub-graph. The "Group Input" and "Group Output" nodes contained in the graph are internally connected to the inputs and outputs of this node.

***Node #34 "Group input"***

This node acts as interface for the inputs of the group defined by node #33.

***Node #35 "Change ROI properties" of type "Set 2D ROI properties"***

This node sets the line color of the incoming ROIs.

- Input "Input" of node #35 receives data from output "Line 1" of node #34
- The parameter "Line color" (line-color) is set to **"#33FF00"**

***Node #36 "Change ROI properties" of type "Set 2D ROI properties"***

This node sets the line color of the incoming ROIs.

- Input "Input" of node #36 receives data from output "Line 2" of node #34
- The parameter "Line color" (line-color) is set to **"#FF0000"**

***Node #37 "Merge ROI lists" of type "Merge 2D ROI lists"***

This node merges multiple ROI lists into a single one.

- Input "Input" of node #37 receives data from output "Output" of node #35
- Input "Input" of node #37 receives data from output "Output" of node #36

***Node #38 "ROI calculator" of type "2D ROI calculator (AND/OR/XOR)"***

This node merges all ROIs in a ROI list into a single ROI.

- Input "Input" of node #38 receives data from output "Output" of node #37
- The parameter "Split after operation" (split-afterwards) is set to **false**
- The parameter "Operation" (operation) is set to **"LogicalOr"**

***Node #39 "Outline ROI" of type "Outline 2D ROI (Hull)"***

This node fits a convex hull around each ROI.

- Input "Input" of node #39 receives data from output "Output" of node #38
- The parameter "Outline method" (outline) is set to **"ConvexHull"**

***Node #40 "Group output"***

This node passes the generated ROIs to the outer pipeline.

- Input "Output" of node #40 receives data from output "Output" of node #39

*Node #41 "Convert ROI to mask" of type "Convert 2D ROI to mask"*

This node converts ROIs into masks.

- Input "ROI" of node #41 receives data from output "Output" of node #33
- Input "Image" of node #41 receives data from output "Output" of node #18

*Node #42 "Morphological operation 2D"*

This node applies a morphological dilation with radius 5 px.

- Input "Input" of node #42 receives data from output "Output" of node #41
- The parameter "Add border before applying operation" (add-border) is set to **true**
- The parameter "Radius" (radius) is set to **5**

*Node #43 "Set to value (grayscale)"*

This node deletes from the gradient image regions outside the known strip region at this point.

- Input "Input" of node #43 receives data from output "Output" of node #18
- Input "Mask" of node #43 receives data from output "Output" of node #42
- The parameter "Only apply to ..." (roi:target-area) is set to **"OutsideMask"**

*Node #44 "Auto threshold 2D" of type "Auto threshold 2D"*

This node applies auto-thresholding to detect the parts of the strip.

- Input "Input" of node #44 receives data from output "Output" of node #43

*Node #45 "Find particles 2D" of type "Find particles 2D"*

This node converts a mask into an ImageJ ROI list.

- Input "Mask" of node #45 receives data from output "Output" of node #44

*Node #46 "Outline ROI" of type "Outline 2D ROI (Hull)"*

This node applies a convex hull outline operation for each ROI.

- Input "Input" of node #46 receives data from output "ROI" of node #45
- The parameter "Outline method" (outline) is set to **"ConvexHull"**

*Node #47 "Sort and extract ROI by statistics" of type "Sort and extract 2D ROI by statistics"*

This node sorts each ROI list by the ROI area in a descending order, and then proceeds to select only the top N ROIs from the list. This node receives parameters from node #4, meaning that its workload is repeated per parameter set.

- Input "ROI" of node #47 receives data from output "Output" of node #46
- Input "{Parameters}" of node #47 receives data from output "Parameters" of node #4
- The parameter "Map line color" (map-line-color) is set to **[Disabled]**
- The parameter "Selected indices" (selected-indices) is set to **[Disabled]**
- The parameter item #1 of "Sort order" (sort-orders) is set to **("Area", "Descending")**
- The parameter "Multiple parameters" in category "Multi-parameter settings" (jipipe:parameter-slot-algorithm/has-parameter-slot) is set to **true**
- The parameter "Attach parameter annotations" in category "Multi-parameter settings" (jipipe:parameter-slot-algorithm/attach-parameter-annotations) is set to **false**

***Node #48 "ROI calculator" of type "2D ROI calculator (AND/OR/XOR)"***

This node merges all ROIs in a ROI list into a single ROI.

- Input "Input" of node #48 receives data from output "Output" of node #47
- The parameter "Split after operation" (split-afterwards) is set to **false**
- The parameter "Operation" (operation) is set to **"LogicalOr"**

***Node #49 "Outline ROI" of type "Outline 2D ROI (Hull)"***

This node fits a rotated rectangle around each ROI. This generates the strip candidate.

- Input "Input" of node #49 receives data from output "Output" of node #48
- The parameter "Outline method" (outline) is set to **"MinimumBoundingRectangle"**

***Node #50 "Merge ROI lists" of type "Merge 2D ROI lists"***

This node merges all strip candidates into a single ROI list.

- Input "Input" of node #50 receives data from output "Output" of node #49

***Node #51 "Sort ROI list (expression)" of type "Sort 2D ROI list (expression)"***

This node sorts the strip candidates by the difference in their aspect ratio to the expected value of 11.

- Input "ROI" of node #51 receives data from output "Output" of node #50
- The parameter "Expression" (expression) is set to **`ABS(AR - custom.expectedAR)`**
- The parameter "Expected AR" in category "Custom variables" (jipipe:algorithm:custom-expression-variables/expectedAR) is set to **11.0**

***Node #52 "Slice ROI list" of type "Slice 2D ROI list"***

This node selects the strip candidate ROI with the lowest difference in aspect ratio difference to the expected value.

- Input "ROI" of node #52 receives data from output "ROI" of node #51
- The parameter "Selected indices" (selected-indices) is set to **0**

***Node #53 "Generate missing 2D ROI lists"***

This node generates empty ROI lists if one is missing.

- Input "ROI" of node #53 receives data from output "Output" of node #52
- Input "Data" of node #53 receives data from output "Output" of node #44

***Node #54 "Convert 2D ROI to mask"***

This node converts ROI lists into masks.

- Input "ROI" of node #54 receives data from output "ROI" of node #53
- Input "Image" of node #54 receives data from output "Output" of node #44

***Node #55 "Scale 2D image"***

This node restores the original scale of the resulting mask image.

- Input "Input" of node #55 receives data from output "Output" of node #54
- The parameter "Y axis" (y-axis) is set to **original_img_h**

- The parameter "X axis" (x-axis) is set to **original_img_w**
- The parameter "Interpolation" (interpolation-method) is set to **"None"**

***Node #56 "Export image"***

This node exports the final mask as PNG.

- Input "Input" of node #56 receives data from output "Output" of node #55
- The parameter "File path" (file-path) is set to **`PATH_COMBINE(project_data_dir.tmp_dir, "strip-disk", #ImageId)`**

#### 2.5 Etest ZOI shape detection

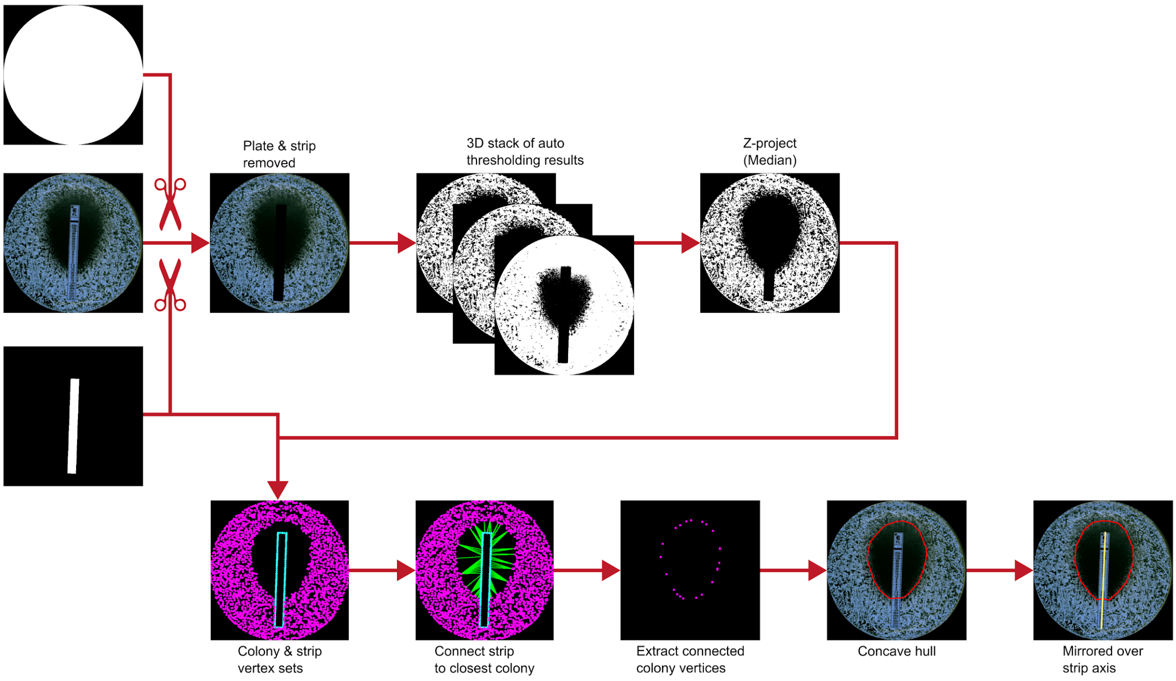

**Supplementary Figure 23:** Etest ZOI shape detection algorithm. Both the area outside the plate and the strip are erased from the image. This is followed by applying all available ImageJ auto-thresholding methods and creating a 3D stack out of the resulting masks. A median Z-projection generates a consensus thresholding. This colony mask and the strip annotation are used to generate colony and strip vertex sets using JIPipe’s filament processing library. Each strip vertex is connected to its closest colony vertex. Selecting only those connected colony vertices yields the key points for the shape that are converted into a symmetric concave ROI.

The following steps are applied by the third algorithm that finds the strip region inside Etests. An overview can be found in **Supplementary Figure 23** and a screenshot is provided in **Supplementary Figure 24**.

###### *Node #1 "Folder list"*

This node references the directory containing the raw images.

- The parameter item #1 of "Folder paths" (folder-paths) is set to **"raw"**

###### *Node #2 "Folder list"*

This node references the directory that contains the plate annotation masks.

- The parameter item #1 of "Folder paths" (folder-paths) is set to **"plate"**

###### *Node #3 "File"*

This node references a CSV file that associates the image ID to the image metadata.

- The parameter "File name" (file-name) is set to **"metadata.csv"**

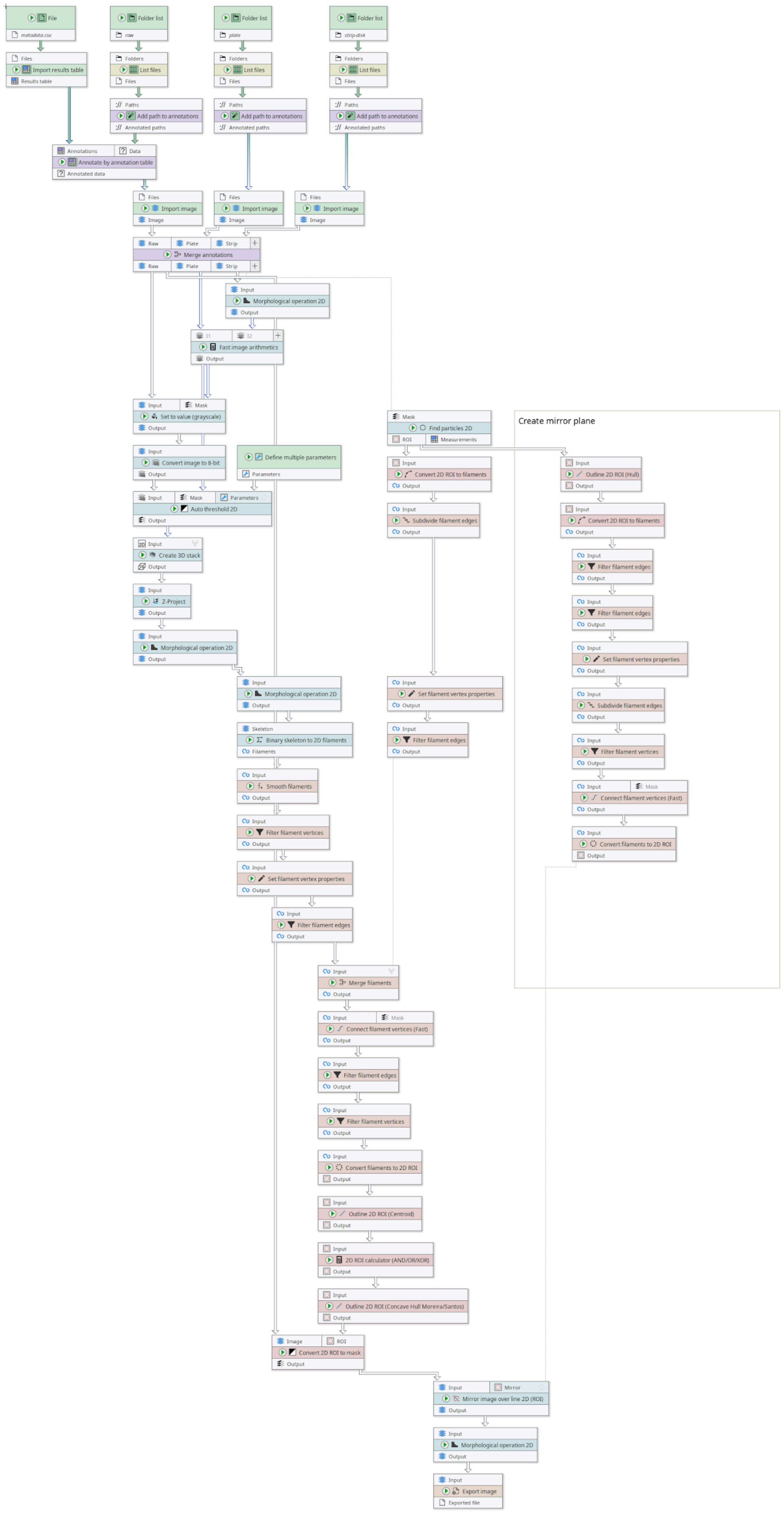

**Supplementary Figure 24:** Full pipeline of the Etest ZOI shape detection algorithm.

*Node #4 "Folder list"*
This node references a directory that contains the Etest strip annotation masks.

- The parameter item #1 of "Folder paths" (folder-paths) is set to **"strip-disk"**

*Node #5 "Define multiple parameters"*
This node contains parameter sets that allow to later apply multiple auto-thresholding methods.

- The parameter "Parameters" (parameter-table) is set to contain the methods **Default, Huang, MaxEntropy, Mean, Minimum, Otsu, Percentile, Yen, Shanbhag, Intermodes, Triangle, MinError, Moments, IsoData, IJ_IsoData, RenyiEntropy,** and **Li**

*Node #6 "List files"*
This node lists all PNG files in the input directory.

- Input "Folders" of node #6 receives data from output "Folder paths" of node #1
- The parameter "Keep file if ..." (filters) is set to **`STRING_MATCHES_GLOB(name, "*.png")`**

*Node #7 "List files"*
This node lists all PNG files in the input directory.

- Input "Folders" of node #7 receives data from output "Folder paths" of node #2
- The parameter "Keep file if ..." (filters) is set to **`STRING_MATCHES_GLOB(name, "*.png")`**

*Node #8 "Import results table"*
This node imports a CSV as ImageJ table.

- Input "Files" of node #8 receives data from output "Filename" of node #3

*Node #9 "List files"*
This node lists all PNG files in the input directory.

- Input "Folders" of node #9 receives data from output "Folder paths" of node #4
- The parameter "Keep file if ..." (filters) is set to **`STRING_MATCHES_GLOB(name, "*.png")`**

*Node #10 "Add path to annotations" of type "Annotate with file/directory name"*
This node attaches the image ID to the input file.

- Input "Paths" of node #10 receives data from output "Files" of node #6
- The parameter "Generated annotation" (generated-annotation) is set to **"#ImageId"**

*Node #11 "Add path to annotations" of type "Annotate with file/directory name"*
This node attaches the image ID to the input file.

- Input "Paths" of node #11 receives data from output "Files" of node #7

- The parameter "Generated annotation" (generated-annotation) is set to **"#ImageId"**

*Node #12 "Add path to annotations" of type "Annotate with file/directory name"*

This node attaches the image ID to the input file.

- Input "Paths" of node #12 receives data from output "Files" of node #9
- The parameter "Generated annotation" (generated-annotation) is set to **"#ImageId"**

*Node #13 "Annotate by annotation table"*

This node attaches the input files with metadata obtained from the table loaded by node #8.

- Input "Annotations" of node #13 receives data from output "Results table" of node #8
- Input "Data" of node #13 receives data from output "Annotated paths" of node #10

*Node #14 "Import image"*

This node loads the incoming PNG file as ImageJ image.

- Input "Files" of node #14 receives data from output "Annotated paths" of node #11

*Node #15 "Import image"*

This node loads the incoming PNG file as ImageJ image.

- Input "Files" of node #15 receives data from output "Annotated paths" of node #12

*Node #16 "Import image"*

This node loads the incoming PNG file as ImageJ image.

- Input "Files" of node #16 receives data from output "Annotated data" of node #13

*Node #17 "Merge annotations"*

This node copies annotations from the raw image to the plate and strip annotations.

- Input "Raw" of node #17 receives data from output "Image" of node #16
- Input "Plate" of node #17 receives data from output "Image" of node #14
- Input "Strip" of node #17 receives data from output "Image" of node #15

*Node #18 "Find particles 2D"*

This node utilizes the ImageJ particle finder to convert a mask into ImageJ ROI lists.

- Input "Mask" of node #18 receives data from output "Strip" of node #17

*Node #19 "Morphological operation 2D"*

This node applies a morphological dilation with a radius of $PixelSize \cdot 30$.

- Input "Input" of node #19 receives data from output "Strip" of node #17
- The parameter "Radius" (radius) is set to **5**
- The parameter "Attach parameter annotations" in category "Adaptive parameters" (jipipe:adaptive-parameters/attach-parameter-annotations) is set to **false**
- The parameter item #1 of "Overridden parameters" in category "Adaptive parameters" (jipipe:adaptive-parameters/overridden-parameters) is set to **`(NUM(PixelSize) * 30, "radius")`**

*Node #20 "Outline 2D ROI (Hull)"*

This node fits a minimum bounding rectangle around the input Etest strip ROIs.

- Input "Input" of node #20 receives data from output "ROI" of node #18
- The parameter "Outline method" (outline) is set to **"MinimumBoundingRectangle"**

*Node #21 "Convert 2D ROI to filaments"*

This node converts the ROI into a JIPipe filaments structure.

- Input "Input" of node #21 receives data from output "ROI" of node #18

*Node #22 "Fast image arithmetics" of type "Fast image arithmetics 2D"*

This node subtracts the I2 image from the I1 image.

- Input "I1" of node #22 receives data from output "Plate" of node #17
- Input "I2" of node #22 receives data from output "Output" of node #19
- The parameter "Expression" (expression) is set to **I1 - I2**

*Node #23 "Convert 2D ROI to filaments"*

This node converts the thresholded colonies into JIPipe filament structures.

- Input "Input" of node #23 receives data from output "Output" of node #20

*Node #24 "Subdivide filament edges"*

This node subdivides filament edges until the length is larger than 10 px.

- Input "Input" of node #24 receives data from output "Output" of node #21
- The parameter "Split if longer than ..." (maximum-length) is set to **10 px**
- The parameter "Maximum iterations" (max-num-iterations) is set to **[Disabled]**

*Node #25 "Set to value (grayscale)"*

This node erases the area outside the plate.

- Input "Input" of node #25 receives data from output "Raw" of node #17
- Input "Mask" of node #25 receives data from output "Output" of node #22
- The parameter "Only apply to ..." (roi:target-area) is set to **"OutsideMask"**

*Node #26 "Filter filament edges"*

This node removes all filament edges that are smaller than the maximum length over all edges.

- Input "Input" of node #26 receives data from output "Output" of node #23
- The parameter "Only keep edge if" (filter) is set to **`length < MAX(all.length)`**

*Node #27 "Set filament vertex properties"*

This node sets the color and value of the filament vertices.

- Input "Input" of node #27 receives data from output "Output" of node #24
- The parameter "Color" (color) is set to **"#33FFFF"**
- The parameter "Value" (value) is set to **0.0**

*Node #28 "Convert image to 8-bit"*

This node ensures that the incoming images are 8-bit greyscale.

- Input "Input" of node #28 receives data from output "Output" of node #25

##### *Node #29 "Filter filament edges"*

This node removes all filament edges that are smaller than the maximum length over all edges.

- Input "Input" of node #29 receives data from output "Output" of node #26
- The parameter "Only keep edge if" (filter) is set to **`length < MAX(all.length)`**

##### *Node #30 "Filter filament edges"*

This node removes all filament edges.

- Input "Input" of node #30 receives data from output "Output" of node #27
- The parameter "Only keep edge if" (filter) is set to **false**

##### *Node #31 "Auto threshold 2D"*

This node applies auto-thresholding. Statistics are only calculated from the pixel values inside the plate mask. This node applies all ImageJ auto-thresholding methods, as it receives parameter sets from a dedicated input.

- Input "Input" of node #31 receives data from output "Output" of node #28
- Input "Mask" of node #31 receives data from output "Output" of node #22
- Input "{Parameters}" of node #31 receives data from output "Parameters" of node #5
- The parameter "Calculate threshold based on ..." (source-area) is set to **"InsideMask"**
- The parameter "Multiple parameters" in category "Multi-parameter settings" (jipipe:parameter-slot-algorithm/has-parameter-slot) is set to **true**

##### *Node #32 "Set filament vertex properties"*

This node sets the color and the value of the filament vertices.

- Input "Input" of node #32 receives data from output "Output" of node #29
- The parameter "Color" (color) is set to **"#00FFFF"**
- The parameter "Value" (value) is set to **1.0**

##### *Node #33 "Create 3D stack"*

This node merges the different thresholding results received from node #31 into a single 3D stack.

- Input "Input" of node #33 receives data from output "Output" of node #31

##### *Node #34 "Subdivide filament edges"*

This node splits all filament edges once at the center.

- Input "Input" of node #34 receives data from output "Output" of node #32
- The parameter "Split if longer than ..." (maximum-length) is set to **[Disabled]**
- The parameter "Maximum iterations" (max-num-iterations) is set to **1**
- The parameter "Override value" (override-value) is set to **0.0**

##### *Node #35 "Z-Project"*

This node applies a Z-project using a median operation.

- Input "Input" of node #35 receives data from output "Output" of node #33
- The parameter "Method" (method) is set to **"Median"**

##### *Node #36 "Filter filament vertices"*

This node only keeps filament vertices with a value of zero, i.e. that were generated from the colonies.

- Input "Input" of node #36 receives data from output "Output" of node #34
- The parameter "Only keep vertex if" (filter) is set to **value == 0**

##### *Node #37 "Morphological operation 2D"*

This node applies a morphological opening operation with a radius `MAX(1, NUM(PixelSize) * 5)`.

- Input "Input" of node #37 receives data from output "Output" of node #35
- The parameter "Operation" (operation) is set to **"OPENING"**
- The parameter "Attach parameter annotations" in category "Adaptive parameters" (jipipe:adaptive-parameters/attach-parameter-annotations) is set to **false**
- The parameter item #1 of "Overridden parameters" in category "Adaptive parameters" (jipipe:adaptive-parameters/overridden-parameters) is set to **(`MAX(1, NUM(PixelSize) * 5),"radius"`)**

##### *Node #38 "Connect filament vertices (Fast)"*

This node connects each strip vertex with the closest colony vertex.

- Input "Input" of node #38 receives data from output "Output" of node #36
- The parameter "Edge length" (edge-length) is set to **x=0 y=Infinity**
- The parameter "Candidate vertex degree limit" (candidate-vertex-degree-limit) is set to **x=0 y=1**
- The parameter "Candidate vertex value limit" (candidate-vertex-value-limit) is set to **x=-Infinity y=Infinity**

##### *Node #39 "Morphological operation 2D"*

This node applies a morphological external gradient with radius 1.

- Input "Input" of node #39 receives data from output "Output" of node #37
- The parameter "Operation" (operation) is set to **"EXTERNAL_GRADIENT"**

##### *Node #40 "Convert filaments to 2D ROI"*

This node converts filaments into ImageJ ROI lists.

- Input "Input" of node #40 receives data from output "Output" of node #38
- The parameter "With vertices" (with-vertices) is set to **false**

##### *Node #41 "Binary skeleton to 2D filaments"*

This node converts a binary skeleton into JIPipe filaments.

- Input "Skeleton" of node #41 receives data from output "Output" of node #39

##### *Node #42 "Smooth filaments"*

This node smoothes the filaments, ensuring that vertices have an average distance of approximately 10 px.

- Input "Input" of node #42 receives data from output "Filaments" of node #41
- The parameter "Factor (X)" (factor-x) is set to **10.0**
- The parameter "Factor (Y)" (factor-y) is set to **10.0**

###### *Node #43 "Filter filament vertices"*

This node only selects vertices that have at least one connection.

- Input "Input" of node #43 receives data from output "Output" of node #42
- The parameter "Only keep vertex if" (filter) is set to **degree > 0**

###### *Node #44 "Set filament vertex properties"*

This node colors the vertices and sets the value to 1.

- Input "Input" of node #44 receives data from output "Output" of node #43
- The parameter "Color" (color) is set to **"#FF00FF"**
- The parameter "Value" (value) is set to **1.0**

###### *Node #45 "Filter filament edges"*

This node removes all filament edges.

- Input "Input" of node #45 receives data from output "Output" of node #44
- The parameter "Only keep edge if" (filter) is set to **false**

###### *Node #46 "Merge filaments"*

This node merges multiple filament sets.

- Input "Input" of node #46 receives data from output "Output" of node #45
- Input "Input" of node #46 receives data from output "Output" of node #30

###### *Node #47 "Connect filament vertices (Fast)"*

This node connects filament vertices with constraints.

- Input "Input" of node #47 receives data from output "Output" of node #46
- The parameter "Connect in 3D" (enable-3d) is set to **false**
- The parameter "Ignore if path exists" (ignore-if-has-path) is set to **false**
- The parameter "Limit created edges (source/target)" (limit-connections) is set to **1**
- The parameter "Source vertex filter" (source-vertex-filter) is set to **value == 0**
- The parameter "Target vertex filter" (target-vertex-filter) is set to **value == 1**
- The parameter "Bypass connection limit for targets" (ignore-limit-connections-for-target) is set to **true**
- The parameter "Edge length" (edge-length) is set to **x=0 y=Infinity**
- The parameter "Candidate vertex degree limit" (candidate-vertex-degree-limit) is set to **x=0 y=2147483647**
- The parameter "Candidate vertex value limit" (candidate-vertex-value-limit) is set to **x=-Infinity y=Infinity**

###### *Node #48 "Filter filament edges"*

This node only keeps edges with a minimum length of $PixelSize \cdot 100$.

- Input "Input" of node #48 receives data from output "Output" of node #47
- The parameter "Only keep edge if" (filter) is set to **`length > (NUM(PixelSize) * 100)`**

##### *Node #49 "Filter filament vertices"*

This node only keeps filament vertices with at least one connection and in the colony set.

- Input "Input" of node #49 receives data from output "Output" of node #48
- The parameter "Only keep vertex if" (filter) is set to **`degree > 0 AND value == 1`**

##### *Node #50 "Convert filaments to 2D ROI"*

This node converts filaments to ImageJ ROIs.

- Input "Input" of node #50 receives data from output "Output" of node #49
- The parameter "With edges" (with-edges) is set to **false**

##### *Node #51 "Outline 2D ROI (Centroid)"*

This node finds the centroid of each ROI.

- Input "Input" of node #51 receives data from output "Output" of node #50

##### *Node #52 "2D ROI calculator (AND/OR/XOR)"*

This node merges all ROIs in a list into a single one.

- Input "Input" of node #52 receives data from output "Output" of node #51
- The parameter "Operation" (operation) is set to **"LogicalOr"**

##### *Node #53 "Outline 2D ROI (Concave Hull Moreira/Santos)"*

This node finds the concave hull of the ROI.

- Input "Input" of node #53 receives data from output "Output" of node #52
- The parameter "Delete invalid ROIs" (skip-invalid-inputs) is set to **true**

##### *Node #54 "Convert 2D ROI to mask"*

This node converts ROI lists into a mask.

- Input "Image" of node #54 receives data from output "Raw" of node #17
- Input "ROI" of node #54 receives data from output "Output" of node #53

##### *Node #55 "Mirror image over line 2D (ROI)"*

This node ensures that the ZOI shape is symmetric.

- Input "Input" of node #55 receives data from output "Output" of node #54
- Input "Mirror" of node #55 receives data from output "Output" of node #40

##### *Node #56 "Morphological operation 2D"*

This node applies a morphological closing with a radius $PixelSize \cdot 50$.

- Input "Input" of node #56 receives data from output "Output" of node #55
- The parameter "Radius" (radius) is set to **5**
- The parameter "Operation" (operation) is set to **"CLOSING"**

- The parameter "Attach parameter annotations" in category "Adaptive parameters" (jipipe:adaptive-parameters/attach-parameter-annotations) is set to **false**
- The parameter item #1 of "Overridden parameters" in category "Adaptive parameters" (jipipe:adaptive-parameters/overridden-parameters) is set to **`(NUM(PixelSize) * 50, "radius")`**

***Node #57 "Export image"***

This node exports the ZOI shape mask as a PNG image.

- Input "Input" of node #57 receives data from output "Output" of node #56
- The parameter "File path" (file-path) is set to **`PATH_COMBINE(project_data_dir.tmp_dir, "zoi-shape", #ImageId)`**

#### 2.6 DiskImageR re-implementation

Rather than the *diskImageR* implementation, which generates multiple radial line profile measurements from the center of the DDA disk to the plate border, the *J-AST* algorithm (**Fig. 4a**) generates ring-shaped measurement regions with a configurable width of 1 mm around the DDA disk. To eliminate unwanted noise, all labels within a configurable distance of 10 mm from the plate border or outside the plate are removed. The average colony intensities $mean_{\tau, r}$ for each radius r and respective time point τ are extracted with a JIPipe operation, and converted into corresponding normalized values $relmean_{\tau, r}$ that are weighted relative to the minimum pixel intensity within the plate $bgr_{\tau}$, and the maximum over all $mean_{\tau, r}$: $relmean_{\tau, r} = \left| \frac{mean_{\tau,r} - bgr_{\tau}}{MAX_i(mean_{\tau,r}) - bgr_{\tau}} \right|$.
The resulting values from the early time point are used to find the $RAD_n$ as the first r where $(1 - relmean_{early, r})$ is larger or equal to the threshold n. The values are then used to determine $FoG_n = \left( \int_0^{RAD_n} relmean_{late,r}\, dr \right) / RAD_n$ by calculating the area under the $relmean_{late, r}$ curve on the late image and dividing it by the $RAD_n$.

The *J-AST* algorithm provides additional measurements, including a radius of inhibition $RAD_{late, n}$ that is calculated on the late time point image instead of the early time point image, $FoG_{early, n}$ and $FoG_{late, n}$, which are fractions of growth that are calculated only on the early or late time point images, respectively, as well as their difference $\Delta FoG_n = FoG_{late, n} - FoG_{early, n}$. *J-AST* additionally provides an analysis algorithm suitable for single images that again use the RAD and FoG values from the same time point (**Supplementary section 2.7**).

The following steps are applied by our re-implementation of the DiskImageR algorithm in JIPipe. An overview can be found in **Fig. 3**. The structure of the pipeline is shown in **Supplementary Figure 25**.

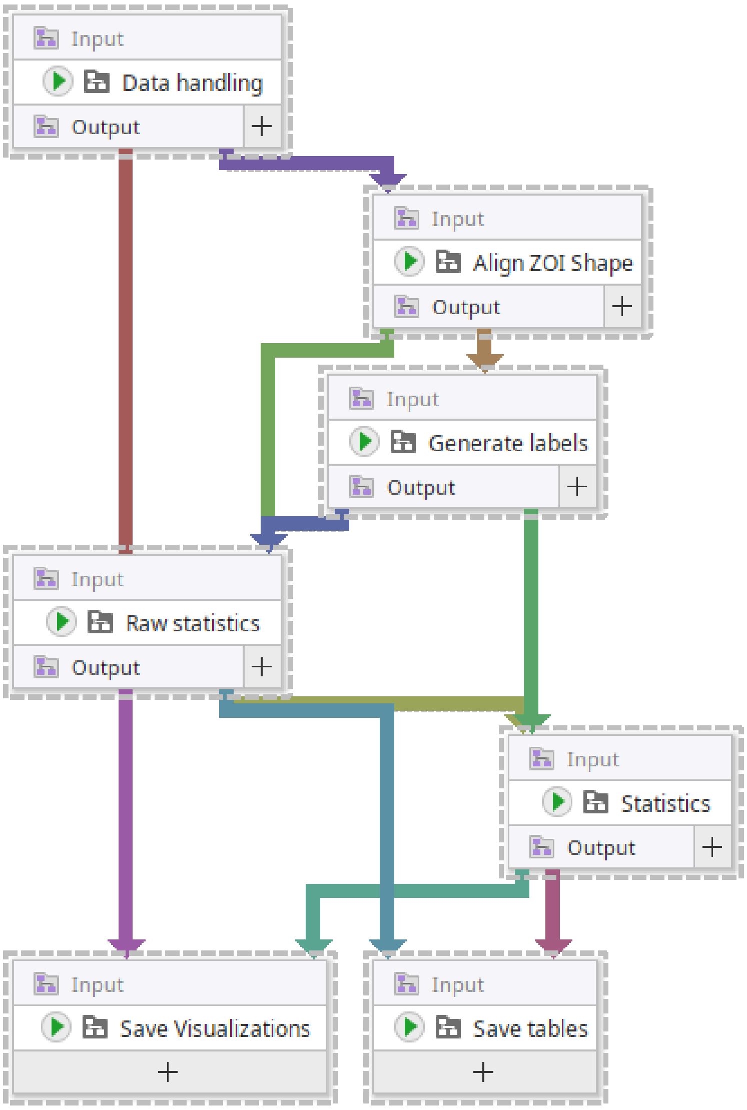

**Supplementary Figure 25:** Compartment structure of our DiskImageR re-implementation in JIPipe. We selected the nodes to better highlight the connections. This diagram also applies to the single-image variant.

##### 2.6.1.1 Compartment C1 "Data handling"

This compartment is responsible for finding, organizing, and importing the input images and associated metadata. A screenshot of the pipeline can be found in **Supplementary Figure 26**.

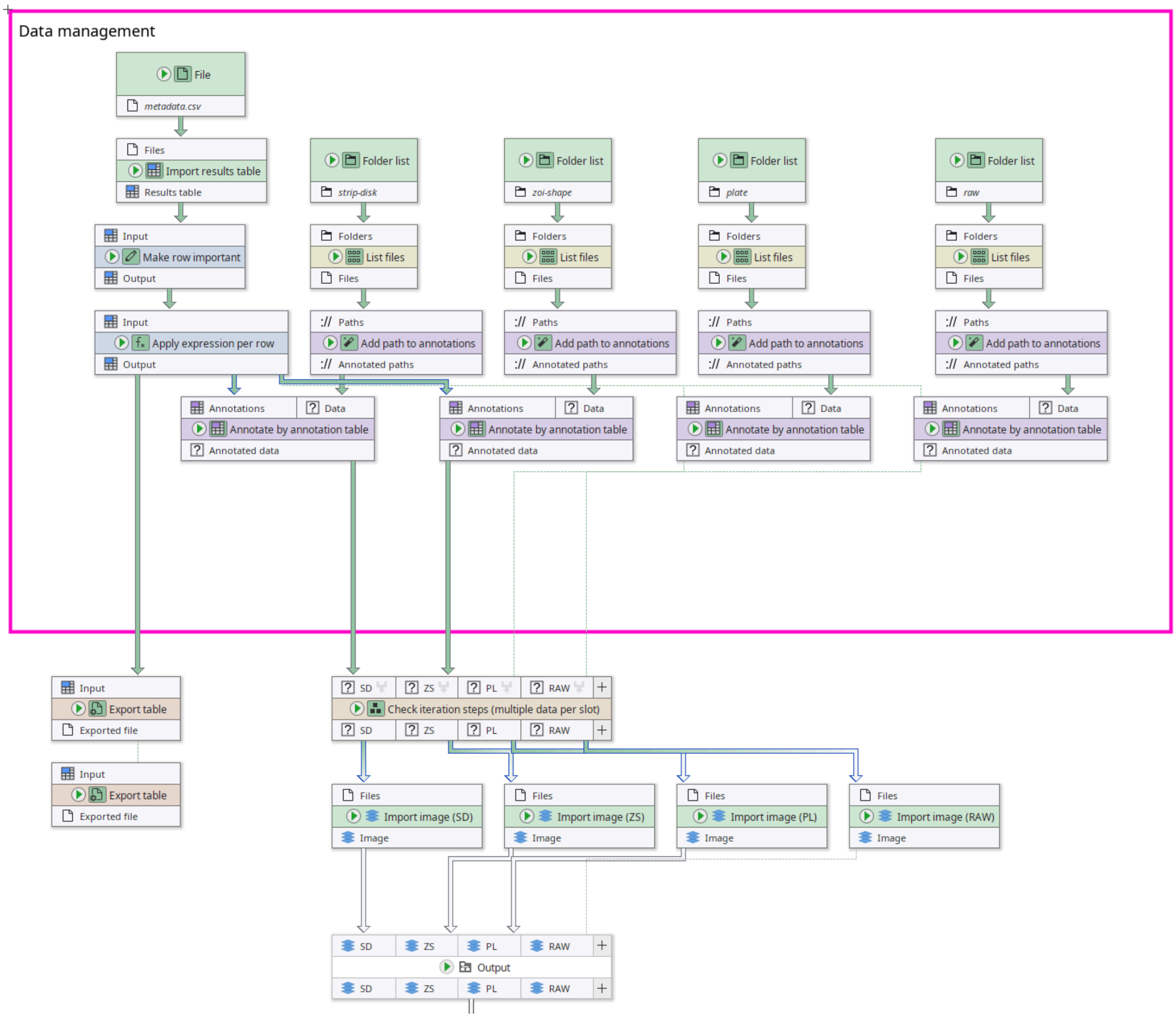

**Supplementary Figure 26:** Contents of the "Data Handling" compartment of our DiskImageR re-implementation in JIPipe.

##### *Node #1 "Folder list"*

This node references the directory that contains the DDA disk and Etest strip annotation masks.

- The parameter item #1 of "Folder paths" (folder-paths) is set to **"strip-disk"**

##### *Node #2 "Folder list"*

This node references the directory that contains the ZOI shape annotation masks.

- The parameter item #1 of "Folder paths" (folder-paths) is set to **"zoi-shape"**

##### *Node #3 "File"*

This node references a CSV file that associates the image ID to its corresponding image metadata.

- The parameter "File name" (file-name) is set to **"metadata.csv"**

##### *Node #4 "Folder list"*

This node references a directory that contains the plate annotation masks.

- The parameter item #1 of "Folder paths" (folder-paths) is set to **"plate"**

*Node #5 "Folder list"*
This node references a directory that contains the input images.

- The parameter item #1 of "Folder paths" (folder-paths) is set to **"raw"**

*Node #6 "List files"*
This node lists all PNG files in the input directory.

- Input "Folders" of node #6 receives data from output "Folder paths" of node #1
- The parameter "Keep file if ..." (filters) is set to **`STRING_MATCHES_GLOB(name, "*.png")`**

*Node #7 "List files"*
This node lists all PNG files in the input directory.

- Input "Folders" of node #7 receives data from output "Folder paths" of node #2
- The parameter "Keep file if ..." (filters) is set to **`STRING_MATCHES_GLOB(name, "*.png")`**

*Node #8 "Import results table"*
This node imports a CSV file as an ImageJ table.

- Input "Files" of node #8 receives data from output "Filename" of node #3

*Node #9 "List files"*
This node lists all PNG files in the input directory.

- Input "Folders" of node #9 receives data from output "Folder paths" of node #4
- The parameter "Keep file if ..." (filters) is set to **`STRING_MATCHES_GLOB(name, "*.png")`**

*Node #10 "List files"*
This node lists all PNG files in the input directory.

- Input "Folders" of node #10 receives data from output "Folder paths" of node #5
- The parameter "Keep file if ..." (filters) is set to **`STRING_MATCHES_GLOB(name, "*.png")`**

*Node #11 "Add path to annotations" of type "Annotate with file/directory name"*
This node attaches the image ID to the incoming file.

- Input "Paths" of node #11 receives data from output "Files" of node #6
- The parameter "Generated annotation" (generated-annotation) is set to **"#ImageId"**

*Node #12 "Add path to annotations" of type "Annotate with file/directory name"*
This node attaches the image ID to the incoming file.

- Input "Paths" of node #12 receives data from output "Files" of node #7
- The parameter "Generated annotation" (generated-annotation) is set to **"#ImageId"**

***Node #13 "Make row important" of type "Rename table column"***

This node renames the annotation “GroupRow” into “#GroupRow”, meaning that JIPipe uses that information to distinguish between data.

- Input "Input" of node #13 receives data from output "Results table" of node #8
- The parameter item #1 of "Renaming entries" (renaming-entries) is set to **source-column = "GroupRow", new-name = "#GroupRow"**

***Node #14 "Add path to annotations" of type "Annotate with file/directory name"***

This node attaches the image ID to the incoming file.

- Input "Paths" of node #14 receives data from output "Files" of node #9
- The parameter "Generated annotation" (generated-annotation) is set to **"#ImageId"**

***Node #15 "Add path to annotations" of type "Annotate with file/directory name"***

This node attaches the image ID to the incoming file.

- Input "Paths" of node #15 receives data from output "Files" of node #10
- The parameter "Generated annotation" (generated-annotation) is set to **"#ImageId"**

***Node #16 "Apply expression per row"***

This node finds the minimum value of the “GroupColumn” column in the input table and writes the result in a dedicated column “MinGroupColumn”.

- Input "Input" of node #16 receives data from output "Output" of node #13
- The parameter item #1 of "Generated values" (entries) is set to **column-name = `"MinGroupColumn"`, value = `MIN(all.GroupColumn)`**

***Node #17 "Annotate by annotation table"***

This node attaches metadata contained in a table to the incoming data.

- Input "Annotations" of node #17 receives data from output "Output" of node #16
- Input "Data" of node #17 receives data from output "Annotated paths" of node #12

***Node #18 "Annotate by annotation table"***

This node attaches metadata contained in a table to the incoming data.

- Input "Annotations" of node #18 receives data from output "Output" of node #16
- Input "Data" of node #18 receives data from output "Annotated paths" of node #15

***Node #19 "Export table"***

This node exports the incoming table as an Excel file.

- Input "Input" of node #19 receives data from output "Output" of node #16
- The parameter "File format" (file-format) is set to **"XLSX"**
- The parameter "File path" (file-path) is set to **`PATH_COMBINE(project_data_dir.tmp_dir, "results", "metadata")`**

***Node #20 "Annotate by annotation table"***

This node attaches metadata contained in a table to the incoming data.

- Input "Annotations" of node #20 receives data from output "Output" of node #16
- Input "Data" of node #20 receives data from output "Annotated paths" of node #11

##### *Node #21 "Export table"*

This node exports the incoming table as an Excel file.

- Input "Input" of node #21 receives data from output "Output" of node #16
- The parameter "File path" (file-path) is set to **`PATH_COMBINE(project_data_dir.tmp_dir, "results", "metadata")`**

##### *Node #22 "Annotate by annotation table"*

This node attaches metadata contained in a table to the incoming data.

- Input "Annotations" of node #22 receives data from output "Output" of node #16
- Input "Data" of node #22 receives data from output "Annotated paths" of node #14

##### *Node #23 "Check iteration steps (multiple data per slot)"*

This node checks if all necessary data is available and throws an error if this is not the case.

- Input "SD" of node #23 receives data from output "Annotated data" of node #20
- Input "ZS" of node #23 receives data from output "Annotated data" of node #17
- Input "PL" of node #23 receives data from output "Annotated data" of node #22
- Input "RAW" of node #23 receives data from output "Annotated data" of node #18
- The parameter "Grouping method" in category "Merging iteration step generation" (jipipe:data-batch-generation/column-matching) is set to **"Custom"**
- The parameter "Custom grouping columns" in category "Merging iteration step generation" (jipipe:data-batch-generation/custom-matched-columns-expression) is set to **`value == "#GroupRow"`**

##### *Node #24 "Import image (SD)" of type "Import image"*

This node imports a PNG file as an ImageJ image.

- Input "Files" of node #24 receives data from output "SD" of node #23

##### *Node #25 "Import image (ZS)" of type "Import image"*

This node imports a PNG file as an ImageJ image.

- Input "Files" of node #25 receives data from output "ZS" of node #23

##### *Node #26 "Import image (PL)" of type "Import image"*

This node imports a PNG file as an ImageJ image.

- Input "Files" of node #26 receives data from output "PL" of node #23

##### *Node #27 "Import image (RAW)" of type "Import image"*

This node imports a PNG file as an ImageJ image.

- Input "Files" of node #27 receives data from output "RAW" of node #23

##### *Node #28 "Output" of type "Compartment output"*

This node passes the data to other compartments.

- Input "SD" of node #28 receives data from output "Image" of node #24
- Input "ZS" of node #28 receives data from output "Image" of node #25
- Input "PL" of node #28 receives data from output "Image" of node #26
- Input "RAW" of node #28 receives data from output "Image" of node #27
- The output slot name is set to **"Output"**

###### 2.6.1.2 Compartment C2 "Align ZOI Shape"

This compartment is responsible for copying the ZOI shape from the early time point to the late time point using a registration function. A screenshot can be found in **Supplementary Figure 26**.

- The "Align ZOI Shape" compartment (C2) receives data from the "Data handling" compartment (C1)

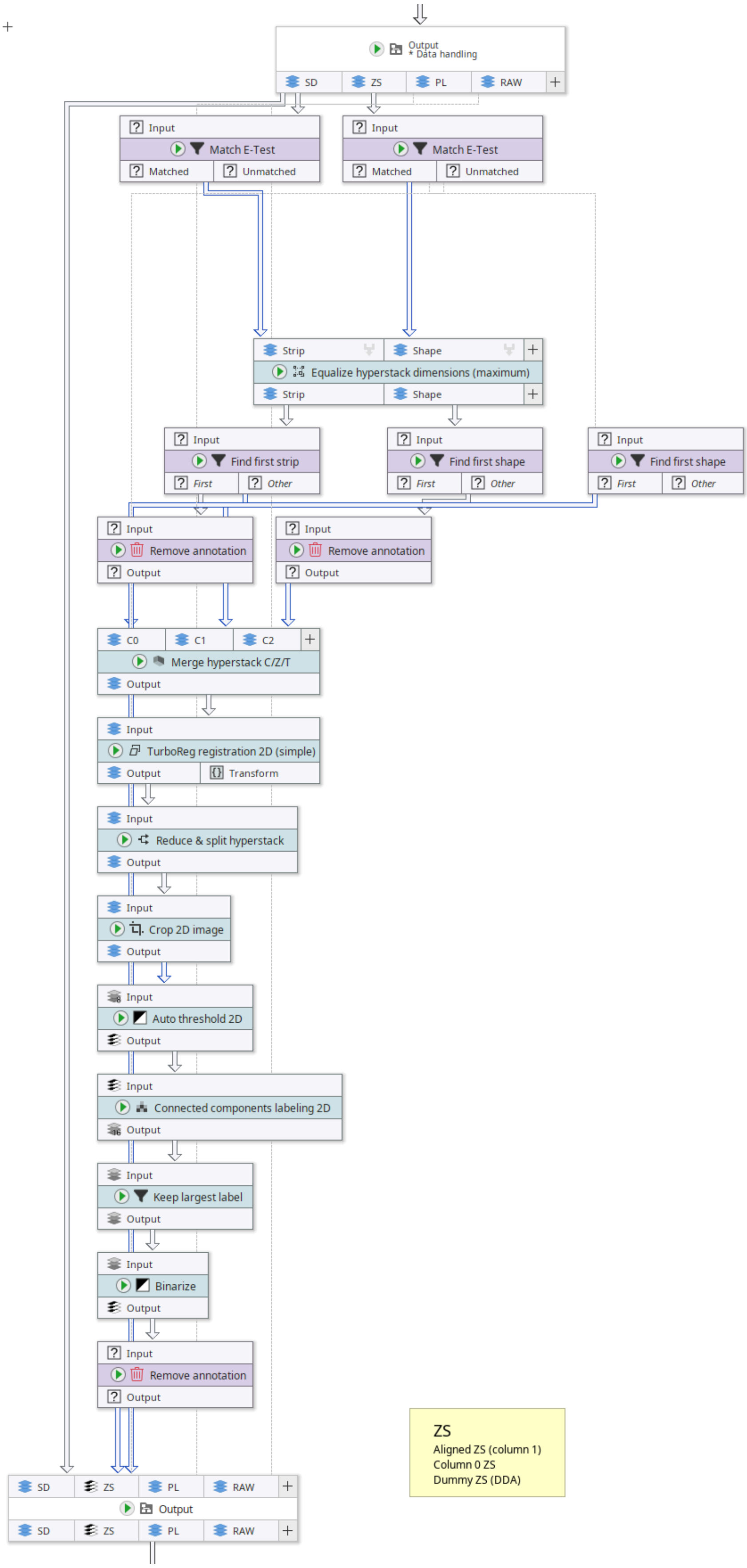

**Supplementary Figure 27:** Contents of the "Align ZOI Shape" compartment of our DiskImageR re-implementation in JIPipe.

*Node #28 "Output" of type "Compartment output"*

This node receives data from the “Data Handling” compartment.

- Input "SD" of node #28 receives data from output "Image" of node #24
- Input "ZS" of node #28 receives data from output "Image" of node #25
- Input "PL" of node #28 receives data from output "Image" of node #26
- Input "RAW" of node #28 receives data from output "Image" of node #27
- The output slot name is set to **"Output"**

*Node #29 "Match E-Test" of type "Filter by annotation (If else)"*

This node splits data flows by their assay type.

- Input "Input" of node #29 receives data from output "SD" of node #28
- The parameter "Only match data if" (filter) is set to **`#AssayType == "ETest"`**

*Node #30 "Match E-Test" of type "Filter by annotation (If else)"*

This node splits data flows by their assay type.

- Input "Input" of node #30 receives data from output "ZS" of node #28
- The parameter "Only match data if" (filter) is set to **`#AssayType == "ETest"`**

*Node #31 "Equalize hyperstack dimensions (maximum)"*

This node ensures that the width and the height inside the incoming image data sets are equal. This is required, as the registration algorithm only works on images with equal width and height.

- Input "Strip" of node #31 receives data from output "Matched" of node #29
- Input "Shape" of node #31 receives data from output "Matched" of node #30
- The parameter "Annotate with original width" (original-width-annotation) is set to **"oW"**
- The parameter "Equalize width and height" (equal-width-and-height) is set to **true**
- The parameter "Annotate with original height" (original-height-annotation) is set to **"oH"**
- The parameter "Grouping method" in category "Merging iteration step generation" (jipipe:data-batch-generation/column-matching) is set to **"MergeAll"**

*Node #32 "Find first shape" of type "Filter by annotation (If else)"*

This node selects the ZOI shapes from the early time point.

- Input "Input" of node #32 receives data from output "Matched" of node #30
- The parameter "Only match data if" (filter) is set to **`GroupColumn == MinGroupColumn`**

*Node #33 "Find first strip" of type "Filter by annotation (If else)"*

This node selects the E-Test strips from the early time point.

- Input "Input" of node #33 receives data from output "Strip" of node #31
- The parameter "Only match data if" (filter) is set to **`GroupColumn == MinGroupColumn`**

*Node #34 "Find first shape" of type "Filter by annotation (If else)"*

This node selects the ZOI shapes from the early time point.

- Input "Input" of node #34 receives data from output "Shape" of node #31

- The parameter "Only match data if" (filter) is set to `GroupColumn == MinGroupColumn`

*Node #35 "Remove annotation"*

This node removes all annotation except “#GroupRow”.

- Input "Input" of node #35 receives data from output "Matched" of node #33
- The parameter "Removed annotations" (annotation-type) is set to `key != "#GroupRow"`

*Node #36 "Remove annotation"*

This node removes the original image width and height annotations.

- Input "Input" of node #36 receives data from output "Unmatched" of node #34
- The parameter "Removed annotations" (annotation-type) is set to `key IN ARRAY("oW", "oH")`

*Node #37 "Merge hyperstack C/Z/T"*

This node creates a hyperstack out of the input image slices.

- Input "C0" of node #37 receives data from output "Output" of node #35
- Input "C1" of node #37 receives data from output "Unmatched" of node #33
- Input "C2" of node #37 receives data from output "Output" of node #36
- The parameter "C0" in category "Input order" (order/C0) is set to **0**
- The parameter "C1" in category "Input order" (order/C1) is set to **0**
- The parameter "C2" in category "Input order" (order/C2) is set to **0**

*Node #38 "TurboReg registration 2D (simple)"*

This node applies an image registration that yields and applies a transformation from the early time point strip to the late time point strip.

- Input "Input" of node #38 receives data from output "Output" of node #37
- The parameter item #1 of "Rules" (rules) is set to **condition = c == 0, reference-t-index = 0, rule-type = "CalculateTransformation", reference-z-index = 0, reference-c-index = 1**
- The parameter item #2 of "Rules" (rules) is set to **condition = c == 1, reference-t-index = t, rule-type = "Ignore", reference-z-index = z, reference-c-index = c**
- The parameter item #3 of "Rules" (rules) is set to **condition = c == 2, reference-t-index = t, rule-type = "UseTransformation", reference-z-index = z, reference-c-index = 0**

*Node #39 "Reduce & split hyperstack"*

This node selects the transformed ZOI shape from the registration output.

- Input "Input" of node #39 receives data from output "Output" of node #38
- The parameter "Indices (Channel)" (indices-c) is set to **2**
- The parameter "Annotate with C indices" (annotate-c) is set to **[Disabled]**
- The parameter "Annotate with Z indices" (annotate-z) is set to **[Disabled]**
- The parameter "Annotate with T indices" (annotate-t) is set to **[Disabled]**

*Node #40 "Crop 2D image"*

This node crops the registered ZOI shape mask back into the original size.

- Input "Input" of node #40 receives data from output "Output" of node #39
- The parameter "ROI" (roi) is set to **{"left":{"expression":"`0`"},"top":{"expression":"`0`"},"right":{"expression":"`0`"},"bottom":{"expression":"`0`"},"width":{"expression":"`NUM(oW)`"},"height":{"expression":"`NUM(oH)`"},"anchor":"TopLeft"}**

*Node #41 "Auto threshold 2D"*

This node applies a “Default” ImageJ auto-thresholding of the registered late ZOI shape.

- Input "Input" of node #41 receives data from output "Output" of node #40

*Node #42 "Connected components labeling 2D"*

This node finds the connected components and creates a label image.

- Input "Input" of node #42 receives data from output "Output" of node #41

*Node #43 "Keep largest label"*

This node deletes all labels except the largest one, which is assumed to be the ZOI shape.

- Input "Input" of node #43 receives data from output "Output" of node #42

*Node #44 "Binarize"*

This node turns the label image back into a mask.

- Input "Input" of node #44 receives data from output "Output" of node #43

*Node #45 "Remove annotation"*

This node removes the original width and height annotations.

- Input "Input" of node #45 receives data from output "Output" of node #44
- The parameter "Removed annotations" (annotation-type) is set to **`key IN ARRAY("oW", "oH")`**

*Node #46 "Output" of type "Compartment output"*

This node passes the generated data to other compartments.

- Input "SD" of node #46 receives data from output "SD" of node #28
- Input "ZS" of node #46 receives data from output "Unmatched" of node #30
- Input "ZS" of node #46 receives data from output "Matched" of node #32
- Input "ZS" of node #46 receives data from output "Output" of node #45
- Input "PL" of node #46 receives data from output "PL" of node #28
- Input "RAW" of node #46 receives data from output "RAW" of node #28
- The output slot name is set to **"Output"**

###### 2.6.1.3 Compartment C3 "Generate labels"

This compartment is responsible for generating the measurement labels that are used to extract the colony intensities over a radius axis. A Screenshot can be found in **Supplementary Figure 27**.

- The "Generate labels" compartment (C3) receives data from the "Align ZOI Shape" compartment (C2)

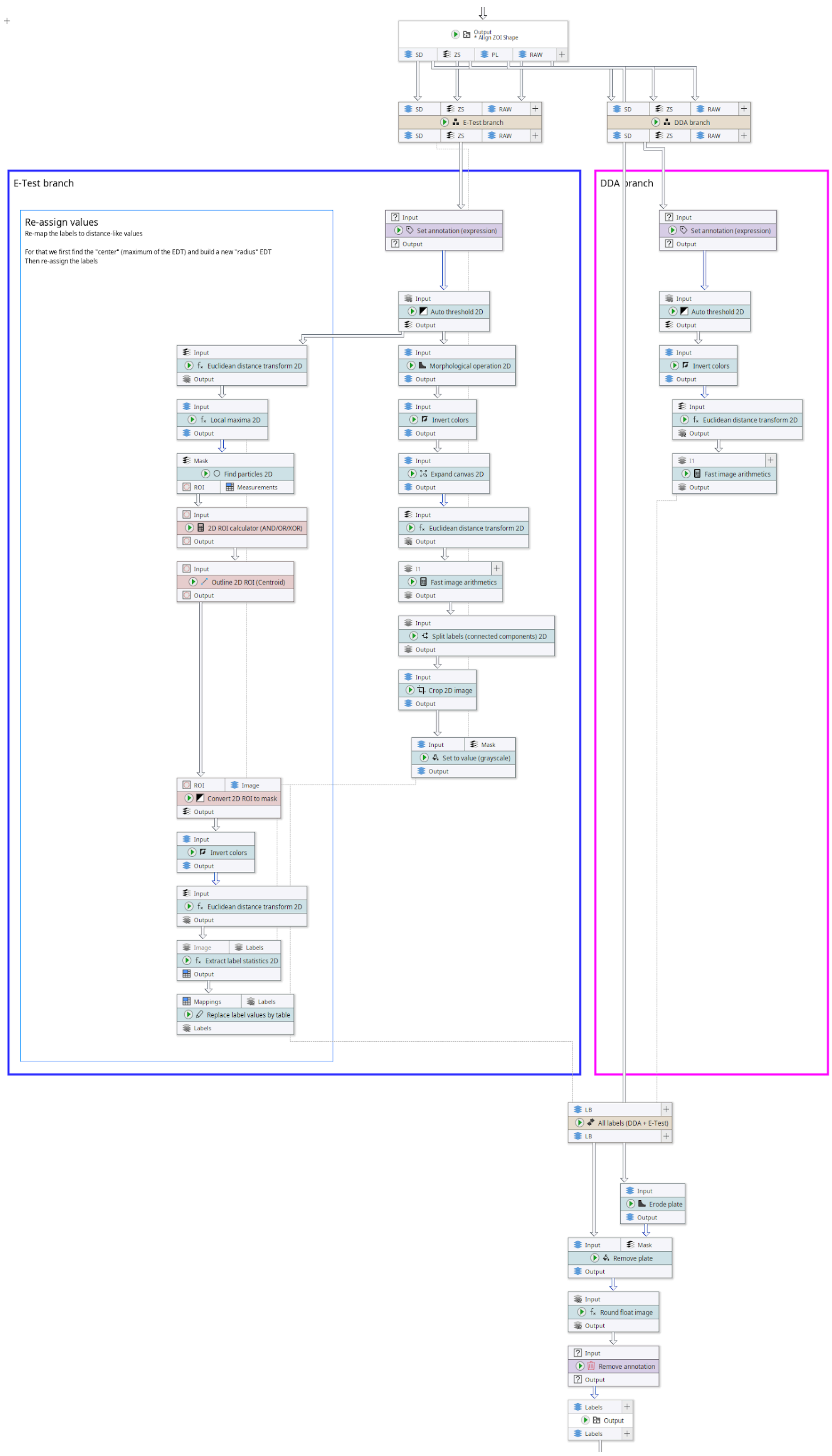

**Supplementary Figure 28:** Contents of the "Generate Labels" compartment of our DiskImageR re-implementation in JIPipe. This diagram also applies to the single-image variant.

***Node #46 "Output" of type "Compartment output"***

This node passes outputs from another compartment into the current one.

- Input "SD" of node #46 receives data from output "SD" of node #28
- Input "ZS" of node #46 receives data from output "Unmatched" of node #30
- Input "ZS" of node #46 receives data from output "Matched" of node #32
- Input "ZS" of node #46 receives data from output "Output" of node #45
- Input "PL" of node #46 receives data from output "PL" of node #28
- Input "RAW" of node #46 receives data from output "RAW" of node #28
- The output slot name is set to **"Output"**

***Node #47 "Erode plate" of type "Morphological operation 2D"***

This node erodes the plate by 10 mm to avoid noise introduced by the plate borders.

- Input "Input" of node #47 receives data from output "PL" of node #46
- The parameter "Add border before applying operation" (add-border) is set to **true**
- The parameter "Operation" (operation) is set to **"EROSION"**
- The parameter "Attach parameter annotations" in category "Adaptive parameters" (jipipe:adaptive-parameters/attach-parameter-annotations) is set to **false**
- The parameter item #1 of "Overridden parameters" in category "Adaptive parameters" (jipipe:adaptive-parameters/overridden-parameters) is set to **(`_global.plateShaveOffMillimeters / NUM(PixelSize),"radius"`)**
- The parameter "Margin right" in category "Border settings" (border-parameters/margin-right) is set to **100**
- The parameter "Margin left" in category "Border settings" (border-parameters/margin-left) is set to **100**
- The parameter "Margin top" in category "Border settings" (border-parameters/margin-top) is set to **100**
- The parameter "Margin bottom" in category "Border settings" (border-parameters/margin-bottom) is set to **100**

***Node #48 "DDA branch" of type "Check iteration steps (one data per slot)"***

This node ensures that all required data for the following steps are present.

- Input "SD" of node #48 receives data from output "SD" of node #46
- Input "ZS" of node #48 receives data from output "ZS" of node #46
- Input "RAW" of node #48 receives data from output "RAW" of node #46
- The parameter "Keep iteration step if" (filter) is set to **#AssayType != "ETest"**

***Node #49 "E-Test branch" of type "Check iteration steps (one data per slot)"***

This node ensures that all required data for the following steps are present.

- Input "SD" of node #49 receives data from output "SD" of node #46
- Input "ZS" of node #49 receives data from output "ZS" of node #46
- Input "RAW" of node #49 receives data from output "RAW" of node #46
- The parameter "Keep iteration step if" (filter) is set to **#AssayType == "ETest"**

***Node #50 "Set annotation (expression)"***

This node calculates the expected label thickness in pixels given the physical size of a pixel and the global parameter (1 mm).

- Input "Input" of node #50 receives data from output "SD" of node #48
- The parameter "Annotation name" (annotation-name) is set to **"LabelThickness"**
- The parameter "Annotation value" (annotation-value) is set to **`MAX(3, ROUND(_global.labelWidthMillimeters * (1 / NUM(PixelSize)))`**

*Node #51 "Set annotation (expression)"*

This node calculates the expected label thickness in pixels given the physical size of a pixel and the global parameter (1 mm).

- Input "Input" of node #51 receives data from output "ZS" of node #49
- The parameter "Annotation name" (annotation-name) is set to **"LabelThickness"**
- The parameter "Annotation value" (annotation-value) is set to **`MAX(3, ROUND(_global.labelWidthMillimeters * (1 / NUM(PixelSize)))`**

*Node #52 "Auto threshold 2D"*

This node applies an auto-thresholding algorithm.

- Input "Input" of node #52 receives data from output "Output" of node #50

*Node #53 "Auto threshold 2D"*

This node applies an auto-thresholding algorithm.

- Input "Input" of node #53 receives data from output "Output" of node #51

*Node #54 "Invert colors" of type "Invert pixel colors/values"*

This node inverts the greyscale color values (255 - x).

- Input "Input" of node #54 receives data from output "Output" of node #52

*Node #55 "Morphological operation 2D"*

This node applies a morphological internal gradient operation with radius 1 px, converting the ZOI shape area into an outline.

- Input "Input" of node #55 receives data from output "Output" of node #53
- The parameter "Operation" (operation) is set to **"INTERNAL_GRADIENT"**

*Node #56 "Euclidean distance transform 2D"*

This node applies an Euclidean distance transform.

- Input "Input" of node #56 receives data from output "Output" of node #53

*Node #57 "Euclidean distance transform 2D"*

This node applies an Euclidean distance transform.

- Input "Input" of node #57 receives data from output "Output" of node #54

*Node #58 "Invert colors" of type "Invert pixel colors/values"*

This node inverts the greyscale color values (255 - x).

- Input "Input" of node #58 receives data from output "Output" of node #55

***Node #59 "Local maxima 2D"***

This node finds all local maxima with a height tolerance of 10.

- Input "Input" of node #59 receives data from output "Output" of node #56

***Node #60 "Fast image arithmetics" of type "Fast image arithmetics 2D"***

This node rounds the distance transform output so that the thickness of the created bins is approximately 1 mm.

- Input "I1" of node #60 receives data from output "Output" of node #57
- The parameter "Expression" (expression) is set to **`ROUND(I1 / LabelThickness) * LabelThickness`**

***Node #61 "Expand canvas 2D"***

This node expands the image canvas size and fills created regions with white pixels. This ensures that no disconnected labels are generated.

- Input "Input" of node #61 receives data from output "Output" of node #58
- The parameter "Background color" (background-color) is set to **"#FFFFFF"**
- The parameter "Y axis" (y-axis) is set to **`default * 2`**
- The parameter "X axis" (x-axis) is set to **`default * 2`**

***Node #62 "Find particles 2D"***

This node uses the ImageJ particle finder to generate ROI lists.

- Input "Mask" of node #62 receives data from output "Output" of node #59

***Node #63 "Euclidean distance transform 2D"***

This node applies an Euclidean distance transform.

- Input "Input" of node #63 receives data from output "Output" of node #61

***Node #64 "2D ROI calculator (AND/OR/XOR)"***

This node merges all ROIs within a ROI list into a single composite ROI.

- Input "Input" of node #64 receives data from output "ROI" of node #62
- The parameter "Split after operation" (split-afterwards) is set to **false**
- The parameter "Operation" (operation) is set to **"LogicalOr"**

***Node #65 "Fast image arithmetics" of type "Fast image arithmetics 2D"***

This node rounds the distance transform output so that the thickness of the created bins is approximately 1 mm.

- Input "I1" of node #65 receives data from output "Output" of node #63
- The parameter "Expression" (expression) is set to **`(ROUND(I1 / LabelThickness) * LabelThickness) + 1`**

***Node #66 "Outline 2D ROI (Centroid)"***

This node finds the centroid of the all-in-one ROI generated by node #64.

- Input "Input" of node #66 receives data from output "Output" of node #64

***Node #67 "Split labels (connected components) 2D"***

This node ensures that labels that have the same value, but are not in the same connected component have a different value.

- Input "Input" of node #67 receives data from output "Output" of node #65

***Node #68 "Convert 2D ROI to mask"***

This node converts the incoming ROI lists into masks.

- Input "ROI" of node #68 receives data from output "Output" of node #66
- Input "Image" of node #68 receives data from output "Output" of node #59

***Node #69 "Crop 2D image"***

This node reverts the operation applied by node #61.

- Input "Input" of node #69 receives data from output "Output" of node #67
- The parameter "ROI" (roi) is set to **{"left":{"expression":"`width / 4`"},"top":{"expression":"`height / 4`"},"right":{"expression":"`0`"},"bottom":{"expression":"`0`"},"width":{"expression":"`width / 2`"},"height":{"expression":"`height / 2`"},"anchor":"TopLeft"}**

***Node #70 "Invert colors" of type "Invert pixel colors/values"***

This node inverts the greyscale image value.

- Input "Input" of node #70 receives data from output "Output" of node #68

***Node #71 "Set to value (grayscale)"***

This node erases the strip/disk from the input image.

- Input "Input" of node #71 receives data from output "Output" of node #69
- Input "Mask" of node #71 receives data from output "SD" of node #49
- The parameter "Only apply to ..." (roi:target-area) is set to **"InsideMask"**

***Node #72 "Euclidean distance transform 2D"***

This node applies an Euclidean distance transform.

- Input "Input" of node #72 receives data from output "Output" of node #70

***Node #73 "Extract label statistics 2D"***

This node measures the mean colony intensity for each measurement label.

- Input "Image" of node #73 receives data from output "Output" of node #72
- Input "Labels" of node #73 receives data from output "Output" of node #71
- The parameter "Measure in physical units" (measure-in-physical-units) is set to **false**
- The parameter "Measurements" (measurements) is set to **{"values":["PixelValueMean"],"collapsed":false}**

***Node #74 "Replace label values by table"***

This node re-maps label values from a table, making the labels correspond to the radius value.

- Input "Mappings" of node #74 receives data from output "Output" of node #73

- Input "Labels" of node #74 receives data from output "Output" of node #71
- The parameter "Old label column" (old-label-column) is set to **("ExistingColumn", "label_id")**
- The parameter "New label column" (new-label-column) is set to **("ExistingColumn", "Mean")**

***Node #75 "All labels (DDA + E-Test)" of type "IO Interface"***

This node merges the branches for DDA and Etest.

- Input "LB" of node #75 receives data from output "Output" of node #60
- Input "LB" of node #75 receives data from output "Labels" of node #74

***Node #76 "Remove plate" of type "Set to value (grayscale)"***

This node removes the plate from the measurement labels.

- Input "Input" of node #76 receives data from output "LB" of node #75
- Input "Mask" of node #76 receives data from output "Output" of node #47
- The parameter "Only apply to ..." (roi:target-area) is set to **"OutsideMask"**

***Node #77 "Round float image"***

This node ensures that all label values are integral.

- Input "Input" of node #77 receives data from output "Output" of node #76
- The parameter "Number of decimals" (decimals) is set to **0**

***Node #78 "Remove annotation"***

This node removes the pre-calculated label thickness annotation that was added in an earlier step.

- Input "Input" of node #78 receives data from output "Output" of node #77
- The parameter "Removed annotations" (annotation-type) is set to **key == "LabelThickness"**

***Node #79 "Output" of type "Compartment output"***

This node passes the measurement labels into other compartments.

- Input "Labels" of node #79 receives data from output "Output" of node #78
- The output slot name is set to **"Output"**

###### 2.6.1.4 Compartment C4 "Raw statistics"

This compartment is responsible for measuring the colony intensities using the measurement labels. A screenshot can be found in **Supplementary Figure 28**.

- The "Raw statistics" compartment (C4) receives data from the "Align ZOI Shape" compartment (C2)
- The "Raw statistics" compartment (C4) receives data from the "Generate labels" compartment (C3)

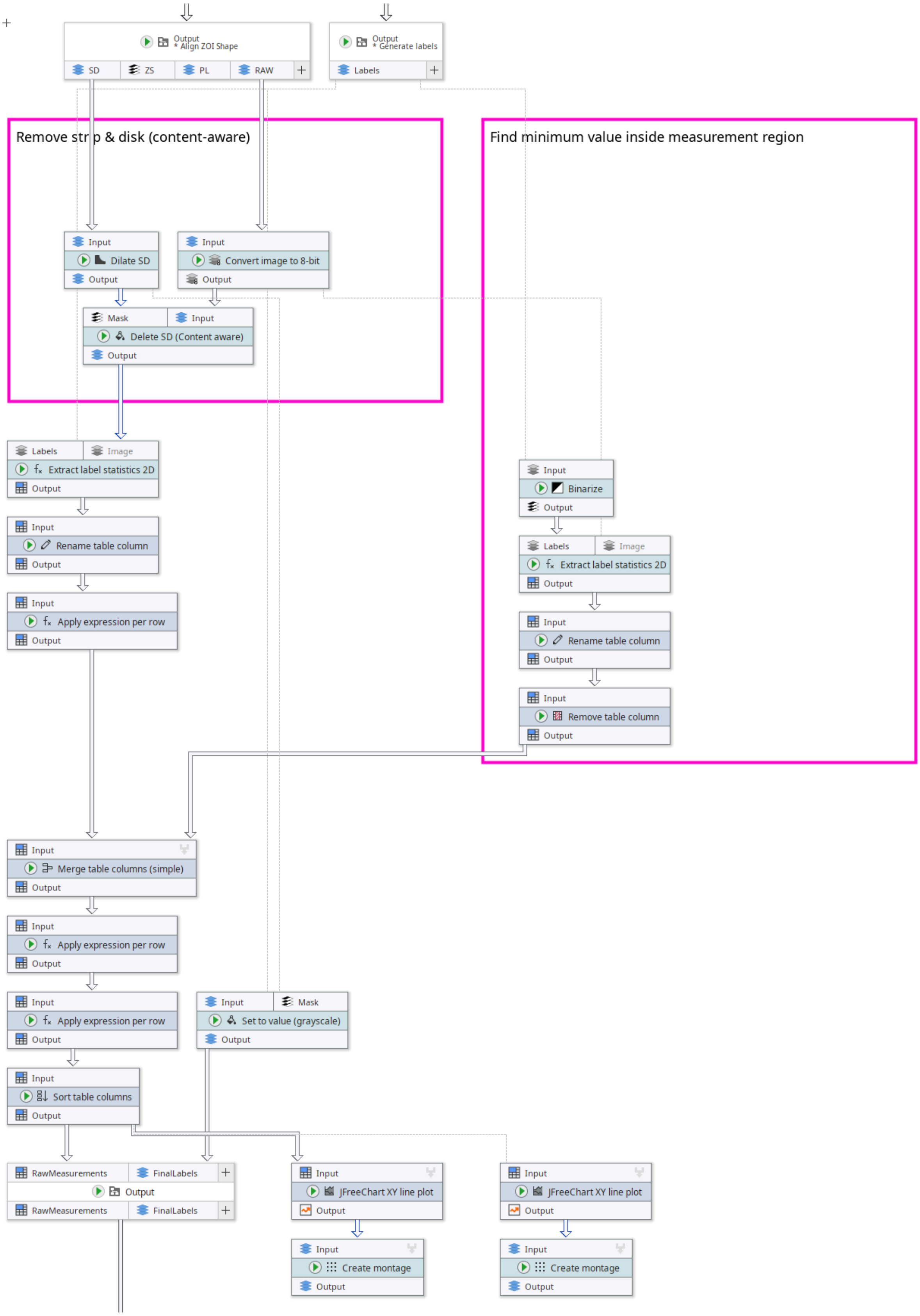

**Supplementary Figure 29:** Contents of the "Raw statistics" compartment of our DiskImageR re-implementation in JIPipe. This diagram also applies to the single-image variant

##### *Node #46 "Output" of type "Compartment output"*

This node receives results from another compartment.

- Input "SD" of node #46 receives data from output "SD" of node #28
- Input "ZS" of node #46 receives data from output "Unmatched" of node #30
- Input "ZS" of node #46 receives data from output "Matched" of node #32
- Input "ZS" of node #46 receives data from output "Output" of node #45
- Input "PL" of node #46 receives data from output "PL" of node #28
- Input "RAW" of node #46 receives data from output "RAW" of node #28
- The output slot name is set to **"Output"**

##### *Node #80 "Dilate SD" of type "Morphological operation 2D"*

This node dilates the DDA disk/Etest strip annotations by 1 mm.

- Input "Input" of node #80 receives data from output "SD" of node #46
- The parameter "Attach parameter annotations" in category "Adaptive parameters" (jipipe:adaptive-parameters/attach-parameter-annotations) is set to **false**
- The parameter item #1 of "Overridden parameters" in category "Adaptive parameters" (jipipe:adaptive-parameters/overridden-parameters) is set to **`(1 / NUM(PixelSize), "radius")`**
- The parameter "Margin right" in category "Border settings" (border-parameters/margin-right) is set to **100**
- The parameter "Margin left" in category "Border settings" (border-parameters/margin-left) is set to **100**
- The parameter "Margin top" in category "Border settings" (border-parameters/margin-top) is set to **100**
- The parameter "Margin bottom" in category "Border settings" (border-parameters/margin-bottom) is set to **100**

##### *Node #81 "Convert image to 8-bit"*

This node ensures that the input image is 8-bit greyscale.

- Input "Input" of node #81 receives data from output "RAW" of node #46

##### *Node #82 "Delete SD (Content aware)" of type "Set to content aware 2D"*

This node erases the DDA disk/Etest strip using a content-aware fill. Compared to erasing by setting the pixels to a specific color, this minimizes the impact on the measurements.

- Input "Mask" of node #82 receives data from output "Output" of node #80
- Input "Input" of node #82 receives data from output "Output" of node #81

##### *Node #79 "Output" of type "Compartment output"*

This node receives results from another compartment.

- Input "Labels" of node #79 receives data from output "Output" of node #78
- The output slot name is set to **"Output"**

##### *Node #83 "Set to value (grayscale)"*

This node erases the DDA disk/Etest strip from the input image.

- Input "Input" of node #83 receives data from output "Labels" of node #79
- Input "Mask" of node #83 receives data from output "Output" of node #80

- The parameter "Only apply to ..." (roi:target-area) is set to **"InsideMask"**

***Node #84 "Binarize"***
This node converts the labels into a binary mask.

- Input "Input" of node #84 receives data from output "Labels" of node #79

***Node #85 "Extract label statistics 2D"***
This node measures for each label the mean colony intensity.

- Input "Labels" of node #85 receives data from output "Labels" of node #79
- Input "Image" of node #85 receives data from output "Output" of node #82
- The parameter "Measurements" (measurements) is set to **{"values":["PixelValueMean"],"collapsed":false}**

***Node #86 "Extract label statistics 2D"***
This node measures for each label the minimum and maximum colony intensity.

- Input "Labels" of node #86 receives data from output "Output" of node #84
- Input "Image" of node #86 receives data from output "Output" of node #81
- The parameter "Measurements" (measurements) is set to **{"values":["PixelValueMinMax"],"collapsed":false}**

***Node #87 "Rename table column"***
This node renames the label ID column into “r_px”. This works because the label IDs already correspond to the radius.

- Input "Input" of node #87 receives data from output "Output" of node #85
- The parameter item #1 of "Renaming entries" (renaming-entries) is set to **source-column = "label_id", new-name = "r_px"**
- The parameter item #2 of "Renaming entries" (renaming-entries) is set to **source-column = "Mean", new-name = "mean"**

***Node #88 "Rename table column"***
This node renames the “Min” column into “background_intensity”.

- Input "Input" of node #88 receives data from output "Output" of node #86
- The parameter item #1 of "Renaming entries" (renaming-entries) is set to **source-column = "Min", new-name = "background_intensity"**

***Node #89 "Apply expression per row"***
This node converts the radius (pixels) into radius (mm).

- Input "Input" of node #89 receives data from output "Output" of node #87
- The parameter item #1 of "Generated values" (entries) is set to **column-name = "r_mm", value = `r_px * NUM(annotations @ "PixelSize")`**

***Node #90 "Remove table column"***
This node removes all values that are not the pre-determined background intensity.

- Input "Input" of node #90 receives data from output "Output" of node #88

- The parameter "Filters" (filters) is set to **`value != "background_intensity"`**

*Node #91 "Merge table columns (simple)"*

This node merges multiple tables column-wise.

- Input "Input" of node #91 receives data from output "Output" of node #89
- Input "Input" of node #91 receives data from output "Output" of node #90
- The parameter "Row normalization" (row-normalization) is set to **"LastValue"**

*Node #92 "Apply expression per row"*

This node finds the minimum and maximum colony intensities.

- Input "Input" of node #92 receives data from output "Output" of node #91
- The parameter item #1 of "Generated values" (entries) is set to **column-name = "min_intensity", value = `MIN(all.mean)`**
- The parameter item #2 of "Generated values" (entries) is set to **column-name = "max_intensity", value = `MAX(all.mean)`**

*Node #93 "Apply expression per row"*

This node calculates the relative colony intensity (rmean) and its related intensity reduction (reduction).

- Input "Input" of node #93 receives data from output "Output" of node #92
- The parameter item #1 of "Generated values" (entries) is set to **column-name = "rmean", value = `ABS(mean-background_intensity)/(max_intensity-background_intensity)`**
- The parameter item #2 of "Generated values" (entries) is set to **column-name = "reduction", value = `1 - rmean`**

*Node #94 "Sort table columns"*

This node sorts the table columns, so that the r_px, r_mm, and mean columns come first.

- Input "Input" of node #94 receives data from output "Output" of node #93
- The parameter item #1 of "Manual order" (manual-order) is set to **"r_px"**
- The parameter item #2 of "Manual order" (manual-order) is set to **"r_mm"**
- The parameter item #3 of "Manual order" (manual-order) is set to **"mean"**

*Node #95 "JFreeChart XY line plot"*

This node creates a plot of rmean over r_mm.

- Input "Input" of node #95 receives data from output "Output" of node #94
- The parameter "X" in category "Input columns" (input-columns/X) is set to **("ExistingColumn", "r_mm")**
- The parameter "Y" in category "Input columns" (input-columns/Y) is set to **("ExistingColumn", "rmean")**

*Node #96 "Output" of type "Compartment output"*

This node passes the raw measurements and labels to other compartments.

- Input "RawMeasurements" of node #96 receives data from output "Output" of node #94

- Input "FinalLabels" of node #96 receives data from output "Output" of node #83
- The output slot name is set to **"Output"**

###### *Node #97 "JFreeChart XY line plot"*

This node creates a plot of the reduction over r_mm.

- Input "Input" of node #97 receives data from output "Output" of node #94
- The parameter "X" in category "Input columns" (input-columns/X) is set to **("ExistingColumn", "r_mm")**
- The parameter "Y" in category "Input columns" (input-columns/Y) is set to **("ExistingColumn", "reduction")**

###### *Node #98 "Create montage"*

This plot merges multiple plots into a montage.

- Input "Input" of node #98 receives data from output "Output" of node #95
- The parameter "Force number of rows" in category "Montage" (montage-parameters/force-rows) is set to **1**
- The parameter "Grouping method" in category "Merging iteration step generation" (jipipe:data-batch-generation/column-matching) is set to **"Custom"**
- The parameter "Custom grouping columns" in category "Merging iteration step generation" (jipipe:data-batch-generation/custom-matched-columns-expression) is set to **"#GroupRow"**

###### *Node #99 "Create montage"*

This plot merges multiple plots into a montage.

- Input "Input" of node #99 receives data from output "Output" of node #97
- The parameter "Force number of rows" in category "Montage" (montage-parameters/force-rows) is set to **1**
- The parameter "Grouping method" in category "Merging iteration step generation" (jipipe:data-batch-generation/column-matching) is set to **"Custom"**
- The parameter "Custom grouping columns" in category "Merging iteration step generation" (jipipe:data-batch-generation/custom-matched-columns-expression) is set to **"#GroupRow"**

##### 2.6.1.5 Compartment C5 "Statistics"

This compartment is responsible for generating the output statistics, i.e. the RAD, FoG, and others. A screenshot of the pipeline can be found in **Supplementary Figure 30**.

- The "Statistics" compartment (C5) receives data from the "Raw statistics" compartment (C4)
- The "Statistics" compartment (C5) receives data from the "Generate labels" compartment (C3)

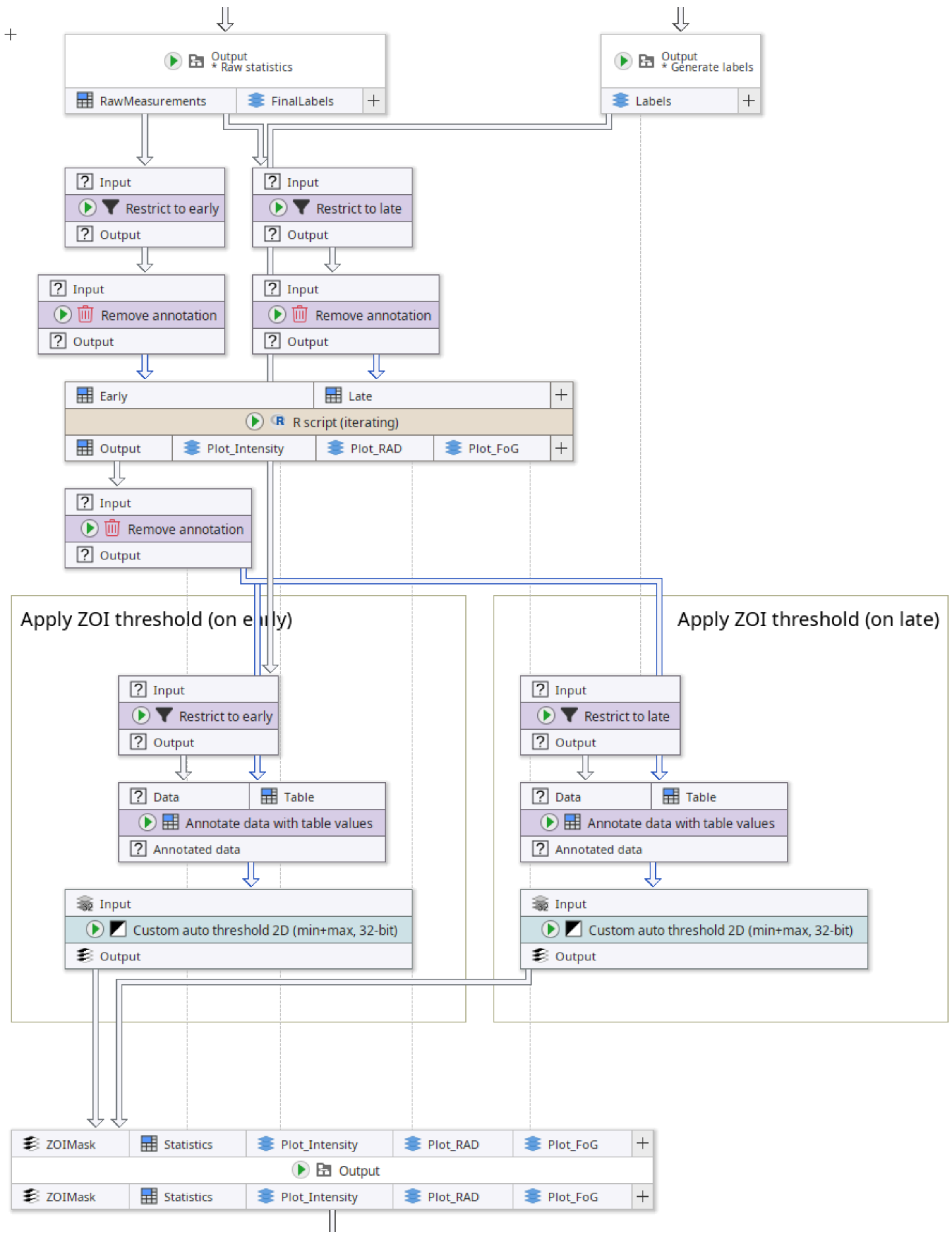

**Supplementary Figure 30:** Contents of the “Statistics” compartment of our DiskImageR re-implementation in JIPipe.

##### *Node #79 "Output" of type "Compartment output"*

This node passes the labels from the “Generate labels” compartment into the current compartment.

- Input "Labels" of node #79 receives data from output "Output" of node #78
- The output slot name is set to **"Output"**

***Node #100 "Restrict to early" of type "Filter by annotation"***

This node filters only data from the early time point.

- Input "Input" of node #100 receives data from output "Labels" of node #79
- The parameter "Only keep data if" (filter) is set to **`NUM(GroupColumn) == 0`**

***Node #101 "Restrict to late" of type "Filter by annotation"***

This node filters only data from the late time point.

- Input "Input" of node #101 receives data from output "Labels" of node #79
- The parameter "Only keep data if" (filter) is set to **`NUM(GroupColumn) == 1`**

***Node #96 "Output" of type "Compartment output"***

This node passes the raw measurements and final labels from the “Raw statistics” compartment into the current one.

- Input "RawMeasurements" of node #96 receives data from output "Output" of node #94
- Input "FinalLabels" of node #96 receives data from output "Output" of node #83
- The output slot name is set to **"Output"**

***Node #102 "Restrict to early" of type "Filter by annotation"***

This node filters only data from the early time point.

- Input "Input" of node #102 receives data from output "RawMeasurements" of node #96
- The parameter "Only keep data if" (filter) is set to **`NUM(GroupColumn) == 0`**

***Node #103 "Restrict to late" of type "Filter by annotation"***

This node filters only data from the late time point.

- Input "Input" of node #103 receives data from output "RawMeasurements" of node #96
- The parameter "Only keep data if" (filter) is set to **`NUM(GroupColumn) == 1`**

***Node #104 "Remove annotation"***

This node removes annotations that reference the time point, image id, and pixel size.

- Input "Input" of node #104 receives data from output "Output" of node #102
- The parameter "Removed annotations" (annotation-type) is set to **`key IN ARRAY("#TimePoint", "#ImageId", "PixelSize")`**

***Node #105 "Remove annotation"***

This node removes annotations that reference the time point, image id, and pixel size.

- Input "Input" of node #105 receives data from output "Output" of node #103
- The parameter "Removed annotations" (annotation-type) is set to **`key IN ARRAY("#TimePoint", "#ImageId", "PixelSize")`**

***Node #106 "R script (iterating)"***

This node runs an R script that handles the calculation of the RAD and FoG values.

- Input "Early" of node #106 receives data from output "Output" of node #104
- Input "Late" of node #106 receives data from output "Output" of node #105

- The parameter "Override R environment" (override-environment) is set to **[Disabled]**
- The parameter item #1 of "Thresholds" in category "Script variables" (variables/thresholds) is set to **0.2**
- The parameter item #2 of "Thresholds" in category "Script variables" (variables/thresholds) is set to **0.5**
- The parameter item #3 of "Thresholds" in category "Script variables" (variables/thresholds) is set to **0.8**

The pipeline has two variants, one using numeric integration (default), and another using R's function approximation. The following code is used by the variant using numeric integration:

```
library(ggplot2)
library(gridExtra)
library(tidyr)
library(reshape2)  # for melting data frames
theme_set(theme_minimal())
# Build dataset name from JIPipe annotations
dataset.name <- paste0(JIPipe.TextAnnotations[["#AssayType"]],
                       "_", JIPipe.TextAnnotations[["#Experiment"]],
                       "_", JIPipe.TextAnnotations[["#Sample"]])
# Get the input data frames
df.early <- JIPipe.GetInputAsDataFrame("Early")
df.late <- JIPipe.GetInputAsDataFrame("Late")
# Remove early frame rows with zero mean
df.early <- df.early[df.early$mean > 0,]
# Initialize intensity plots
plot.early <- ggplot(df.early, aes(x = r_mm, y = rmean)) +
    geom_line() +
    labs(title = paste0(dataset.name, " - Early Frame Intensity Profile
(earlyRAD)"),
         x = "Distance (mm)", y = "Relative mean intensity") +
    scale_x_continuous(limits = range(c(0, max(max(df.late$r_mm),
max(df.early$r_mm)))))
plot.late <- ggplot(df.late, aes(x = r_mm, y = rmean)) +
    geom_line() +
    labs(title = paste0(dataset.name, " - Late Frame Intensity Profile
(earlyRAD & lateRAD)"),
         x = "Distance (mm)", y = "Relative mean intensity") +
    scale_x_continuous(limits = range(c(0, max(max(df.late$r_mm),
max(df.early$r_mm)))))
# Define thresholds to use
# thresholds <- c(0.2, 0.5, 0.8)
# Robust AUC function using the trapezoidal rule on discrete data
calculate_auc <- function(x, y) {
  if(length(x) < 2 || length(y) < 2) {
    return(0) # Not enough points for integration
  }
  dx <- diff(x)
  y0 <- head(y, -1)
```

```
  y1 <- tail(y, -1)
  auc <- sum(dx * (y0 + y1) / 2)
  return(auc)
}
# Initialize a results data frame for all thresholds
results_df <- data.frame()
for(threshold in thresholds) {
    print(paste("Growth reduction", threshold))
    # Compute earlyRAD from the early table
    earlyRAD_mm <- df.early[df.early$rmean >= (1 - threshold),]$r_mm[1]
    earlyRAD_px <- df.early[df.early$rmean >= (1 - threshold),]$r_px[1]
    # Compute lateRAD from the late table
    lateRAD_mm <- df.late[df.late$rmean >= (1 - threshold),]$r_mm[1]
    lateRAD_px <- df.late[df.late$rmean >= (1 - threshold),]$r_px[1]

    # --- Calculate earlyLateFoG ---
    # Use earlyRAD applied to the late frame:
    df.late.filtered <- df.late[df.late$r_px <= earlyRAD_px,]
    mri_late_for_early <- max(df.late.filtered$rmean)
    auc_earlyLate <- calculate_auc(df.late.filtered$r_px,
df.late.filtered$rmean)
    amax_late_for_early <- earlyRAD_px * mri_late_for_early
    earlyLateFoG <- auc_earlyLate / amax_late_for_early

    # --- Calculate earlyFoG ---
    # Integration on the early frame up to earlyRAD:
    df.early.filtered <- df.early[df.early$r_px <= earlyRAD_px,]
    mri_early <- max(df.early.filtered$rmean)
    auc_earlyFoG <- calculate_auc(df.early.filtered$r_px,
df.early.filtered$rmean)
    amax_early <- earlyRAD_px * mri_early
    earlyFoG <- auc_earlyFoG / amax_early
    # --- Calculate lateFoG ---
    # Integration on the late frame using lateRAD:
    df.late.filtered2 <- df.late[df.late$r_px <= lateRAD_px,]
    mri_late <- max(df.late.filtered2$rmean)
    auc_lateFoG <- calculate_auc(df.late.filtered2$r_px,
df.late.filtered2$rmean)
    amax_late <- lateRAD_px * mri_late
    lateFoG <- auc_lateFoG / amax_late
    # --- Calculate deltaFoG ---
    deltaFoG <- lateFoG - earlyFoG
    # Compile current threshold results into a data frame row
    df.out <- data.frame(
        threshold      = threshold,
        earlyRAD_mm    = earlyRAD_mm,
        earlyRAD_px    = earlyRAD_px,
        lateRAD_mm     = lateRAD_mm,
```

```
        lateRAD_px      = lateRAD_px,
        earlyLateFoG    = earlyLateFoG,
        earlyFoG        = earlyFoG,
        lateFoG         = lateFoG,
        deltaFoG        = deltaFoG
    )
    # Accumulate the results
    results_df <- rbind(results_df, df.out)
    # Also output the data frame for this threshold using JIPipe
    JIPipe.AddOutputDataFrame(slot = "Output", data = df.out, annotations =
list("#Threshold" = as.character(threshold)))
    # Update the intensity plots:
    # For the early frame, add the earlyRAD vertical line (red dashed)
    plot.early <- plot.early + geom_vline(xintercept = earlyRAD_mm, color =
"red", linetype = "dashed")
    # For the late frame, add both the earlyRAD (red dashed) and lateRAD
(blue dotted) lines
    plot.late <- plot.late +
                 geom_vline(xintercept = earlyRAD_mm, color = "red",
linetype = "dashed") +
                 geom_vline(xintercept = lateRAD_mm, color = "blue",
linetype = "dotted")
}
### Save the combined intensity plots
combined_intensity_plot <- grid.arrange(plot.early, plot.late, ncol = 1)
plot_folder_intensity <- JIPipe.AddOutputFolder("Plot_Intensity")
ggsave(filename = paste0(plot_folder_intensity, "/data.png"),
       width = 10, height = 12, dpi = 120, plot = combined_intensity_plot)
### Create and save the RAD plot (earlyRAD vs lateRAD in mm vs threshold)
rad_df <- results_df[, c("threshold", "earlyRAD_mm", "lateRAD_mm")]
rad_long <- melt(rad_df, id.vars = "threshold", variable.name =
"Measurement", value.name = "RAD_mm")
rad_plot <- ggplot(rad_long, aes(x = threshold, y = RAD_mm, color =
Measurement)) +
            geom_line() + geom_point() +
            labs(title = paste0(dataset.name, " - RAD (mm) vs Threshold"),
x = "Threshold", y = "RAD (mm)") +
            theme_minimal()
plot_folder_rad <- JIPipe.AddOutputFolder("Plot_RAD")
ggsave(filename = paste0(plot_folder_rad, "/data.png"),
       width = 6, height = 4, dpi = 120, plot = rad_plot)
### Create and save the FoG plot (all FoG measurements vs threshold)
fog_df <- results_df[, c("threshold", "earlyLateFoG", "earlyFoG",
"lateFoG", "deltaFoG")]
fog_long <- melt(fog_df, id.vars = "threshold", variable.name =
"Measurement", value.name = "FoG")
fog_plot <- ggplot(fog_long, aes(x = threshold, y = FoG, color =
Measurement)) +
```

```
            geom_line() + geom_point() +
            labs(title =  paste0(dataset.name, "- FoG Measurements vs
Threshold"), x = "Threshold", y = "FoG") +
            theme_minimal()
plot_folder_fog <- JIPipe.AddOutputFolder("Plot_FoG")
ggsave(filename = paste0(plot_folder_fog, "/data.png"),
       width = 6, height = 4, dpi = 120, plot = fog_plot)
```

The following code is used by the pipeline variant that utilizes R's function approximation algorithm for the integration step:

```
library(ggplot2)
library(gridExtra)
library(tidyr)
library(reshape2)  # for melting data frames

theme_set(theme_minimal())

# Build dataset name from JIPipe annotations
dataset.name <- paste0(JIPipe.TextAnnotations[["#AssayType"]],
                       "_", JIPipe.TextAnnotations[["#Experiment"]],
                       "_", JIPipe.TextAnnotations[["#Sample"]])

# Get the input data frames
df.early <- JIPipe.GetInputAsDataFrame("Early")
df.late <- JIPipe.GetInputAsDataFrame("Late")

# Remove early frame rows with zero mean
df.early <- df.early[df.early$mean > 0,]

# Initialize intensity plots
plot.early <- ggplot(df.early, aes(x = r_mm, y = rmean)) +
    geom_line() +
    labs(title = paste0(dataset.name, " - Early Frame Intensity Profile
(earlyRAD)"),
         x = "Distance (mm)", y = "Relative mean intensity") +
    scale_x_continuous(limits = range(c(0, max(max(df.late$r_mm),
max(df.early$r_mm)))))

plot.late <- ggplot(df.late, aes(x = r_mm, y = rmean)) +
    geom_line() +
    labs(title = paste0(dataset.name, " - Late Frame Intensity Profile
(earlyRAD & lateRAD)"),
         x = "Distance (mm)", y = "Relative mean intensity") +
    scale_x_continuous(limits = range(c(0, max(max(df.late$r_mm),
max(df.early$r_mm)))))

# Define thresholds to use
```

```
# thresholds <- c(0.2, 0.5, 0.8)

# Initialize a results data frame for all thresholds
results_df <- data.frame()

for(threshold in thresholds) {
    print(paste("Growth reduction", threshold))

    # Compute earlyRAD from the early table
    earlyRAD_mm <- df.early[df.early$rmean >= (1 - threshold),]$r_mm[1]
    earlyRAD_px <- df.early[df.early$rmean >= (1 - threshold),]$r_px[1]

    # Compute lateRAD from the late table
    lateRAD_mm <- df.late[df.late$rmean >= (1 - threshold),]$r_mm[1]
    lateRAD_px <- df.late[df.late$rmean >= (1 - threshold),]$r_px[1]

    # Prepare interpolation functions for integration
    f.late  <- approxfun(x = df.late$r_px,  y = df.late$rmean,  method =
"linear", rule = 2, yright = 0)
    f.early <- approxfun(x = df.early$r_px, y = df.early$rmean, method =
"linear", rule = 2, yright = 0)

    # --- Calculate earlyLateFoG ---
    # Use earlyRAD applied to the late frame:
    df.late.filtered <- df.late[df.late$r_px <= earlyRAD_px,]
    mri_late_for_early <- max(df.late.filtered$rmean)
    auc_earlyLate <- integrate(f.late, lower = 0, upper = earlyRAD_px,
subdivisions = 2000)$value
    amax_late_for_early <- earlyRAD_px * mri_late_for_early
    earlyLateFoG <- auc_earlyLate / amax_late_for_early

    # --- Calculate earlyFoG ---
    # Integration on the early frame up to earlyRAD:
    df.early.filtered <- df.early[df.early$r_px <= earlyRAD_px,]
    mri_early <- max(df.early.filtered$rmean)
    auc_earlyFoG <- integrate(f.early, lower = 0, upper = earlyRAD_px,
subdivisions = 2000)$value
    amax_early <- earlyRAD_px * mri_early
    earlyFoG <- auc_earlyFoG / amax_early

    # --- Calculate lateFoG ---
    # Integration on the late frame using lateRAD:
    df.late.filtered2 <- df.late[df.late$r_px <= lateRAD_px,]
    mri_late <- max(df.late.filtered2$rmean)
    auc_lateFoG <- integrate(f.late, lower = 0, upper = lateRAD_px,
subdivisions = 2000)$value
    amax_late <- lateRAD_px * mri_late
    lateFoG <- auc_lateFoG / amax_late
```

```
    # --- Calculate deltaFoG ---
    deltaFoG <- lateFoG - earlyFoG

    # Compile current threshold results into a data frame row
    df.out <- data.frame(
        threshold      = threshold,
        earlyRAD_mm    = earlyRAD_mm,
        earlyRAD_px    = earlyRAD_px,
        lateRAD_mm     = lateRAD_mm,
        lateRAD_px     = lateRAD_px,
        earlyLateFoG   = earlyLateFoG,
        earlyFoG       = earlyFoG,
        lateFoG        = lateFoG,
        deltaFoG       = deltaFoG
    )

    # Accumulate the results
    results_df <- rbind(results_df, df.out)

    # Also output the data frame for this threshold using JIPipe
    JIPipe.AddOutputDataFrame(slot = "Output", data = df.out, annotations =
list("#Threshold" = as.character(threshold)))

    # Update the intensity plots:
    # For the early frame, add the earlyRAD vertical line (red dashed)
    plot.early <- plot.early + geom_vline(xintercept = earlyRAD_mm, color =
"red", linetype = "dashed")
    # For the late frame, add both the earlyRAD (red dashed) and lateRAD
(blue dotted) lines
    plot.late <- plot.late +
                 geom_vline(xintercept = earlyRAD_mm, color = "red",
linetype = "dashed") +
                 geom_vline(xintercept = lateRAD_mm, color = "blue",
linetype = "dotted")
}

### Save the combined intensity plots
combined_intensity_plot <- grid.arrange(plot.early, plot.late, ncol = 1)
plot_folder_intensity <- JIPipe.AddOutputFolder("Plot_Intensity")
ggsave(filename = paste0(plot_folder_intensity, "/data.png"),
       width = 10, height = 12, dpi = 120, plot = combined_intensity_plot)

### Create and save the RAD plot (earlyRAD vs lateRAD in mm vs threshold)
rad_df <- results_df[, c("threshold", "earlyRAD_mm", "lateRAD_mm")]
rad_long <- melt(rad_df, id.vars = "threshold", variable.name =
"Measurement", value.name = "RAD_mm")
```

```
rad_plot <- ggplot(rad_long, aes(x = threshold, y = RAD_mm, color =
Measurement)) +
            geom_line() + geom_point() +
            labs(title = paste0(dataset.name, " - RAD (mm) vs Threshold"),
x = "Threshold", y = "RAD (mm)") +
            theme_minimal()
plot_folder_rad <- JIPipe.AddOutputFolder("Plot_RAD")
ggsave(filename = paste0(plot_folder_rad, "/data.png"),
       width = 6, height = 4, dpi = 120, plot = rad_plot)

### Create and save the FoG plot (all FoG measurements vs threshold)
fog_df <- results_df[, c("threshold", "earlyLateFoG", "earlyFoG",
"lateFoG", "deltaFoG")]
fog_long <- melt(fog_df, id.vars = "threshold", variable.name =
"Measurement", value.name = "FoG")
fog_plot <- ggplot(fog_long, aes(x = threshold, y = FoG, color =
Measurement)) +
            geom_line() + geom_point() +
            labs(title =  paste0(dataset.name, "- FoG Measurements vs
Threshold"), x = "Threshold", y = "FoG") +
            theme_minimal()
plot_folder_fog <- JIPipe.AddOutputFolder("Plot_FoG")
ggsave(filename = paste0(plot_folder_fog, "/data.png"),
       width = 6, height = 4, dpi = 120, plot = fog_plot)
```

###### *Node #107 "Remove annotation"*

This node removes the group column annotation (indicator of time point).

- Input "Input" of node #107 receives data from output "Output" of node #106
- The parameter "Removed annotations" (annotation-type) is set to **key == "GroupColumn"**

###### *Node #108 "Annotate data with table values"*

This note attaches the RAD and FoG values obtained from the R script to the target data.

- Input "Data" of node #108 receives data from output "Output" of node #101
- Input "Table" of node #108 receives data from output "Output" of node #107
- The parameter item #1 of "Generated annotations" (generated-annotations) is set to **name = "rad_px", value = `LAST_OF(earlyRAD_px)`**
- The parameter item #2 of "Generated annotations" (generated-annotations) is set to **name = "rad_mm", value = `LAST_OF(earlyRAD_mm)`**
- The parameter item #3 of "Generated annotations" (generated-annotations) is set to **name = "FoG", value = `LAST_OF(earlyLateFoG)`**

###### *Node #109 "Annotate data with table values"*

This note attaches the RAD and FoG values obtained from the R script to the target data.

- Input "Data" of node #109 receives data from output "Output" of node #100

- Input "Table" of node #109 receives data from output "Output" of node #107
- The parameter item #1 of "Generated annotations" (generated-annotations) is set to **name = "rad_px", value = `LAST_OF(earlyRAD_px)`**
- The parameter item #2 of "Generated annotations" (generated-annotations) is set to **name = "rad_mm", value = `LAST_OF(earlyRAD_mm)`**
- The parameter item #3 of "Generated annotations" (generated-annotations) is set to **name = "FoG", value = `LAST_OF(earlyLateFoG)`**
- The parameter "Custom grouping columns" in category "Input management" (jipipe:data-batch-generation/custom-matched-columns-expression) is set to **`value == "#GroupRow"`**
- The parameter "Skip incomplete data sets" in category "Input management" (jipipe:data-batch-generation/skip-incomplete) is set to **true**

***Node #110 "Custom auto threshold 2D (min+max, 32-bit)"***

This node applies a RAD threshold to find the ZOI mask.

- Input "Input" of node #110 receives data from output "Annotated data" of node #108
- The parameter "Thresholding function" in category "Minimum threshold" (min-threshold/thresholding-function) is set to **1**
- The parameter "Thresholding function" in category "Maximum threshold" (max-threshold/thresholding-function) is set to **`NUM(rad_px)`**

***Node #111 "Custom auto threshold 2D (min+max, 32-bit)"***

This node applies a RAD threshold to find the ZOI mask.

- Input "Input" of node #111 receives data from output "Annotated data" of node #109
- The parameter "Thresholding function" in category "Minimum threshold" (min-threshold/thresholding-function) is set to **1**
- The parameter "Thresholding function" in category "Maximum threshold" (max-threshold/thresholding-function) is set to **`NUM(rad_px)`**

***Node #112 "Output" of type "Compartment output"***

This node passes the results to other compartments.

- Input "ZOIMask" of node #112 receives data from output "Output" of node #110
- Input "ZOIMask" of node #112 receives data from output "Output" of node #111
- Input "Statistics" of node #112 receives data from output "Output" of node #107
- Input "Plot_Intensity" of node #112 receives data from output "Plot_Intensity" of node #106
- Input "Plot_RAD" of node #112 receives data from output "Plot_RAD" of node #106
- Input "Plot_FoG" of node #112 receives data from output "Plot_FoG" of node #106
- The output slot name is set to **"Output"**

###### 2.6.1.6 Compartment C6 "Save Visualizations"

This compartment generates, collects, and exports visualizations and plots. A screenshot can be found in **Supplementary Figure 31**.

- The "Save Visualizations" compartment (C6) receives data from the "Raw statistics" compartment (C4)

- The "Save Visualizations" compartment (C6) receives data from the "Statistics" compartment (C5)
- The "Save Visualizations" compartment (C6) receives data from the "Data handling" compartment (C1)

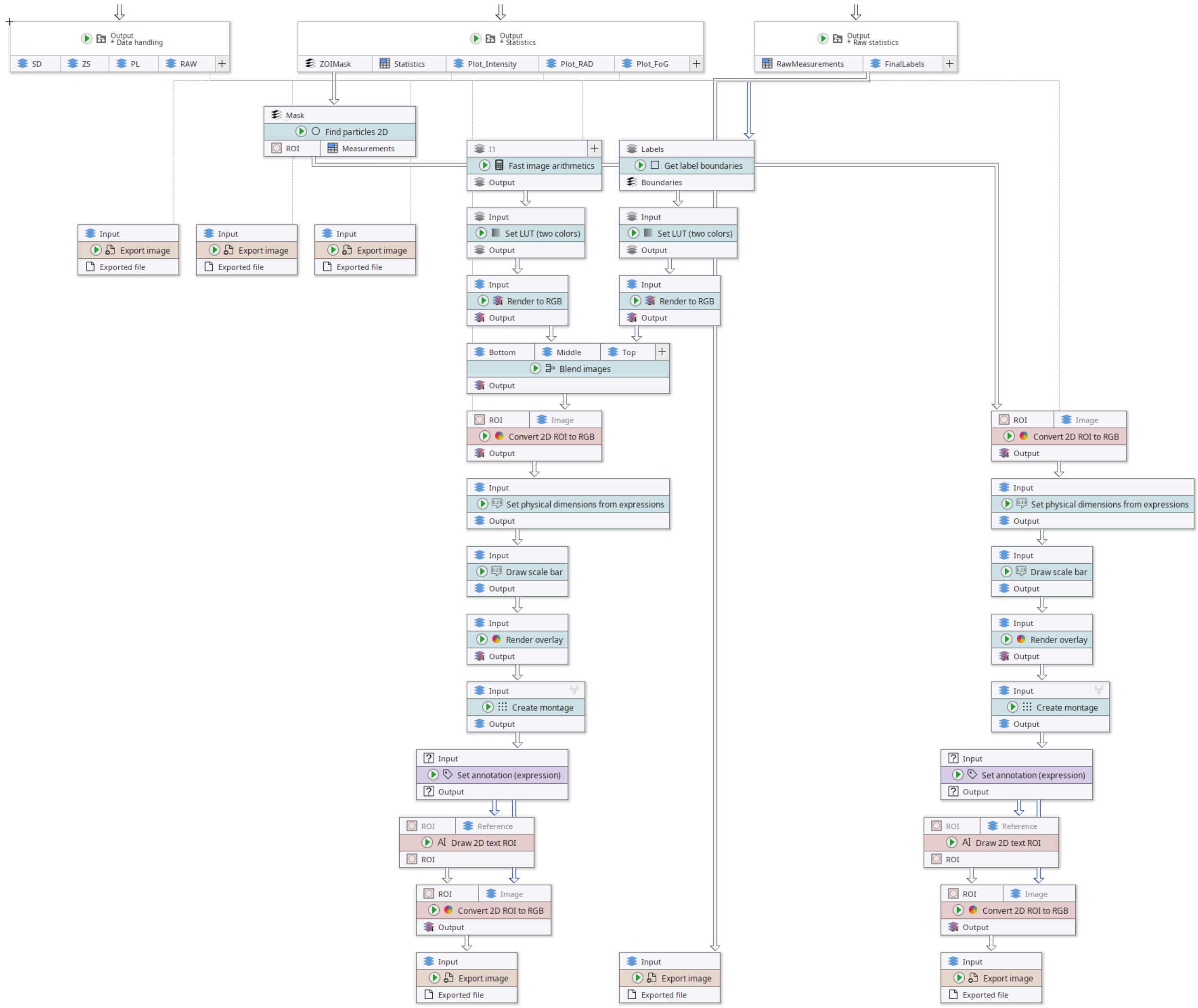

**Supplementary Figure 31:** Contents of the “Save Visualizations” compartment of our DiskImageR re-implementation in JIPipe. This diagram also applies to the single-image variant.

##### *Node #28 "Output" of type "Compartment output"*

This node passes results from another compartment into the current one.

- Input "SD" of node #28 receives data from output "Image" of node #24
- Input "ZS" of node #28 receives data from output "Image" of node #25
- Input "PL" of node #28 receives data from output "Image" of node #26
- Input "RAW" of node #28 receives data from output "Image" of node #27
- The output slot name is set to **"Output"**

##### *Node #96 "Output" of type "Compartment output"*

This node passes results from another compartment into the current one.

- Input "RawMeasurements" of node #96 receives data from output "Output" of node #94

- Input "FinalLabels" of node #96 receives data from output "Output" of node #83
- The output slot name is set to **"Output"**

##### *Node #113 "Get label boundaries"*

This node finds the measurement label boundaries for visualization purposes.

- Input "Labels" of node #113 receives data from output "FinalLabels" of node #96

##### *Node #114 "Fast image arithmetics"*

This node finds the zero-radius label.

- Input "I1" of node #114 receives data from output "FinalLabels" of node #96
- The parameter "Expression" (expression) is set to **I1 == 0**

##### *Node #115 "Export image"*

This node exports an image as a PNG file.

- Input "Input" of node #115 receives data from output "FinalLabels" of node #96
- The parameter "File format" (file-format) is set to **"TIFF"**
- The parameter "File path" (file-path) is set to **`PATH_COMBINE(project_data_dir.tmp_dir, "results", "measurement_labels", #AssayType + "_" + #Experiment + "_" + #Sample + "_" + #TimePoint)`**

##### *Node #116 "Set LUT (two colors)"*

This node sets the LUT of the incoming label boundaries to black-to-cyan.

- Input "Input" of node #116 receives data from output "Boundaries" of node #113
- The parameter "Second color" (second-color) is set to **"#00FFFF"**

##### *Node #117 "Set LUT (two colors)"*

This node sets the LUT of the incoming greyscale image to black-to-red.

- Input "Input" of node #117 receives data from output "Output" of node #114

##### *Node #118 "Render to RGB"*

This node renders the greyscale image with LUT into an RGB image.

- Input "Input" of node #118 receives data from output "Output" of node #116

##### *Node #119 "Render to RGB"*

This node renders the greyscale image with LUT into an RGB image.

- Input "Input" of node #119 receives data from output "Output" of node #117

##### *Node #120 "Blend images"*

This node creates the final visualization by overlaying multiple images.

- Input "Bottom" of node #120 receives data from output "RAW" of node #28
- Input "Middle" of node #120 receives data from output "Output" of node #119
- Input "Top" of node #120 receives data from output "Output" of node #118

- The parameter "Bottom" in category "Layers" (layers/Bottom) is set to **{"opacity":1.0}**
- The parameter "Middle" in category "Layers" (layers/Middle) is set to **{"opacity":0.4}**
- The parameter "Top" in category "Layers" (layers/Top) is set to **{"opacity":1.0}**

###### *Node #112 "Output" of type "Compartment output"*

This node passes results from another compartment into the current one.

- Input "ZOIMask" of node #112 receives data from output "Output" of node #110
- Input "ZOIMask" of node #112 receives data from output "Output" of node #111
- Input "Statistics" of node #112 receives data from output "Output" of node #107
- Input "Plot_Intensity" of node #112 receives data from output "Plot_Intensity" of node #106
- Input "Plot_RAD" of node #112 receives data from output "Plot_RAD" of node #106
- Input "Plot_FoG" of node #112 receives data from output "Plot_FoG" of node #106
- The output slot name is set to **"Output"**

###### *Node #121 "Find particles 2D"*

This node gets the ZOI mask as ImageJ ROI.

- Input "Mask" of node #121 receives data from output "ZOIMask" of node #112

###### *Node #122 "Export image"*

This node exports the incoming image as PNG.

- Input "Input" of node #122 receives data from output "Plot_Intensity" of node #112
- The parameter "File path" (file-path) is set to **`PATH_COMBINE(project_data_dir.tmp_dir, "results", "plots-profile", #AssayType + "_" + #Experiment + "_" + #Sample)`**

###### *Node #123 "Export image"*

This node exports the incoming image as PNG.

- Input "Input" of node #123 receives data from output "Plot_RAD" of node #112
- The parameter "File path" (file-path) is set to **`PATH_COMBINE(project_data_dir.tmp_dir, "results", "plots-rad", #AssayType + "_" + #Experiment + "_" + #Sample)`**

###### *Node #124 "Export image"*

This node exports the incoming image as PNG.

- Input "Input" of node #124 receives data from output "Plot_FoG" of node #112
- The parameter "File path" (file-path) is set to **`PATH_COMBINE(project_data_dir.tmp_dir, "results", "plots-fog", #AssayType + "_" + #Experiment + "_" + #Sample)`**

###### *Node #125 "Convert 2D ROI to RGB"*

This node draws an ImageJ ROI over the reference image.

- Input "ROI" of node #125 receives data from output "ROI" of node #121
- Input "Image" of node #125 receives data from output "Output" of node #120
- The parameter "Override line width" (override-line-width) is set to **4.0**

- The parameter "Override line color" (override-line-color) is set to **"#66FF00"**

##### *Node #126 "Convert 2D ROI to RGB"*
This node draws an ImageJ ROI over the reference image.

- Input "ROI" of node #126 receives data from output "ROI" of node #121
- Input "Image" of node #126 receives data from output "RAW" of node #28
- The parameter "Override line width" (override-line-width) is set to **4.0**
- The parameter "Override line color" (override-line-color) is set to **"#66FF00"**

##### *Node #127 "Set physical dimensions from expressions"*
This node sets the physical pixel dimension metadata of the incoming image.

- Input "Input" of node #127 receives data from output "Output" of node #125
- The parameter "Physical dimension (Y)" (physical-dimension-y) is set to **`PixelSize + " mm"`**
- The parameter "Physical dimension (X)" (physical-dimension-x) is set to **`PixelSize + " mm"`**

##### *Node #128 "Set physical dimensions from expressions"*
This node sets the physical pixel dimension metadata of the incoming image.

- Input "Input" of node #128 receives data from output "Output" of node #126
- The parameter "Physical dimension (Y)" (physical-dimension-y) is set to **`PixelSize + " mm"`**
- The parameter "Physical dimension (X)" (physical-dimension-x) is set to **`PixelSize + " mm"`**

##### *Node #129 "Draw scale bar"*
This node draws a scale bar.

- Input "Input" of node #129 receives data from output "Output" of node #127

##### *Node #130 "Draw scale bar"*
This node draws a scale bar.

- Input "Input" of node #130 receives data from output "Output" of node #128

##### *Node #131 "Render overlay"*
This node draws its associated ROI over the incoming image.

- Input "Input" of node #131 receives data from output "Output" of node #129
- The parameter "Draw outline" (draw-outline-mode) is set to **"IfAvailable"**

##### *Node #132 "Render overlay"*
This node draws its associated ROI over the incoming image.

- Input "Input" of node #132 receives data from output "Output" of node #130
- The parameter "Draw outline" (draw-outline-mode) is set to **"IfAvailable"**

##### *Node #133 "Create montage"*

This node creates an image montage.

- Input "Input" of node #133 receives data from output "Output" of node #131
- The parameter "Custom sorting label" in category "Montage" (montage-parameters/sorting-label-expression) is set to **`NUM(GroupColumn)`**
- The parameter "Label" in category "Montage" (montage-parameters/label-expression) is set to **`STRING_FORMAT("%s_%s_%s_%s", #AssayType, #Experiment, #Sample, #TimePoint)`**
- The parameter "Force number of rows" in category "Montage" (montage-parameters/force-rows) is set to **1**
- The parameter "Label font size" in category "Labels" (montage-parameters/label-parameters/label-font-size) is set to **22**
- The parameter "Border size" in category "Canvas" (montage-parameters/canvas-parameters/border-size) is set to **5**
- The parameter "Border color" in category "Canvas" (montage-parameters/canvas-parameters/border-color) is set to **"#000000"**
- The parameter "Grouping method" in category "Merging iteration step generation" (jipipe:data-batch-generation/column-matching) is set to **"Custom"**
- The parameter "Custom grouping columns" in category "Merging iteration step generation" (jipipe:data-batch-generation/custom-matched-columns-expression) is set to **`value IN ARRAY("#GroupRow", "#Threshold")`**

###### *Node #134 "Create montage"*

This node creates an image montage.

- Input "Input" of node #134 receives data from output "Output" of node #132
- The parameter "Custom sorting label" in category "Montage" (montage-parameters/sorting-label-expression) is set to **`NUM(GroupColumn)`**
- The parameter "Label" in category "Montage" (montage-parameters/label-expression) is set to **`STRING_FORMAT("%s_%s_%s_%s", #AssayType, #Experiment, #Sample, #TimePoint)`**
- The parameter "Force number of rows" in category "Montage" (montage-parameters/force-rows) is set to **1**
- The parameter "Label font size" in category "Labels" (montage-parameters/label-parameters/label-font-size) is set to **22**
- The parameter "Border size" in category "Canvas" (montage-parameters/canvas-parameters/border-size) is set to **5**
- The parameter "Border color" in category "Canvas" (montage-parameters/canvas-parameters/border-color) is set to **"#000000"**
- The parameter "Grouping method" in category "Merging iteration step generation" (jipipe:data-batch-generation/column-matching) is set to **"Custom"**
- The parameter "Custom grouping columns" in category "Merging iteration step generation" (jipipe:data-batch-generation/custom-matched-columns-expression) is set to **`value IN ARRAY("#GroupRow", "#Threshold")`**

###### *Node #135 "Set annotation (expression)"*

This node sets the correct time point annotation.

- Input "Input" of node #135 receives data from output "Output" of node #133
- The parameter "Annotation name" (annotation-name) is set to **`"#TimePoint"`**

- The parameter "Annotation value" (annotation-value) is set to **`JOIN_STRING(SORT_ASCENDING(PARSE_JSON(#TimePoint)) , "-")`**

##### *Node #136 "Set annotation (expression)"*

This node sets the correct time point annotation.

- Input "Input" of node #136 receives data from output "Output" of node #134
- The parameter "Annotation name" (annotation-name) is set to **"`#TimePoint`"**
- The parameter "Annotation value" (annotation-value) is set to **`JOIN_STRING(SORT_ASCENDING(PARSE_JSON(#TimePoint)) , "-")`**

##### *Node #137 "Draw 2D text ROI"*

This node draws the RAD and FoG over the image.

- Input "Reference" of node #137 receives data from output "Output" of node #135
- The parameter "Background margin" (background-margin) is set to **{"left":{"expression":"`2.0`"},"top":{"expression":"`2.0`"},"right":{"expression":"`2.0`"},"bottom":{"expression":"`2.0`"}}**
- The parameter "Font size" (font-size) is set to **22**
- The parameter "Location" (location) is set to **{"left":{"expression":"`0`"},"top":{"expression":"`10`"},"right":{"expression":"`10`"},"bottom":{"expression":"`0`"},"anchor":"TopRight"}**
- The parameter "Text" (text) is set to **`STRING_FORMAT("RAD%d (mm): %s; FoG%d: %s", TO_INTEGER(NUM(#Threshold) * 100), rad_mm, TO_INTEGER(NUM(#Threshold) * 100), D2S(NUM(FoG), 3))`**

##### *Node #138 "Draw 2D text ROI"*

This node draws the RAD and FoG over the image.

- Input "Reference" of node #138 receives data from output "Output" of node #136
- The parameter "Background margin" (background-margin) is set to **{"left":{"expression":"`2.0`"},"top":{"expression":"`2.0`"},"right":{"expression":"`2.0`"},"bottom":{"expression":"`2.0`"}}**
- The parameter "Font size" (font-size) is set to **22**
- The parameter "Location" (location) is set to **{"left":{"expression":"`0`"},"top":{"expression":"`10`"},"right":{"expression":"`10`"},"bottom":{"expression":"`0`"},"anchor":"TopRight"}**
- The parameter "Text" (text) is set to **`STRING_FORMAT("RAD%d (mm): %s; FoG%d: %s", TO_INTEGER(NUM(#Threshold) * 100), rad_mm, TO_INTEGER(NUM(#Threshold) * 100), D2S(NUM(FoG), 3))`**

##### *Node #139 "Convert 2D ROI to RGB"*

This node draws a ROI over the reference image.

- Input "ROI" of node #139 receives data from output "ROI" of node #137
- Input "Image" of node #139 receives data from output "Output" of node #135
- The parameter "Merge same annotation values" in category "Input management" (jipipe:data-batch-generation/annotation-merge-strategy) is set to **"OverwriteExisting"**

##### *Node #140 "Convert 2D ROI to RGB"*

- Input "ROI" of node #140 receives data from output "ROI" of node #138
- Input "Image" of node #140 receives data from output "Output" of node #136
- The parameter "Merge same annotation values" in category "Input management" (jipipe:data-batch-generation/annotation-merge-strategy) is set to **"OverwriteExisting"**

###### *Node #141 "Export image"*

This node exports the incoming image as PNG.

- Input "Input" of node #141 receives data from output "Output" of node #139
- The parameter "File path" (file-path) is set to **`PATH_COMBINE(project_data_dir.tmp_dir, "results", "visualization_all", #AssayType + "_" + #Experiment + "_" + #Sample + "_" + #TimePoint + "_t" + (NUM(#Threshold) * 100))`**

###### *Node #142 "Export image"*

This node exports the incoming image as PNG.

- Input "Input" of node #142 receives data from output "Output" of node #140
- The parameter "File path" (file-path) is set to **`PATH_COMBINE(project_data_dir.tmp_dir, "results", "visualization_zoi", #AssayType + "_" + #Experiment + "_" + #Sample + "_" + #TimePoint + "_t" + (NUM(#Threshold) * 100))`**

##### 2.6.1.7 Compartment C7 "Save tables"

This compartment is responsible for collecting the statistics and exporting them as Excel and CSV tables. A screenshot can be found in **Supplementary Figure 32**.

- The "Save tables" compartment (C7) receives data from the "Raw statistics" compartment (C4)
- The "Save tables" compartment (C7) receives data from the "Statistics" compartment (C5)

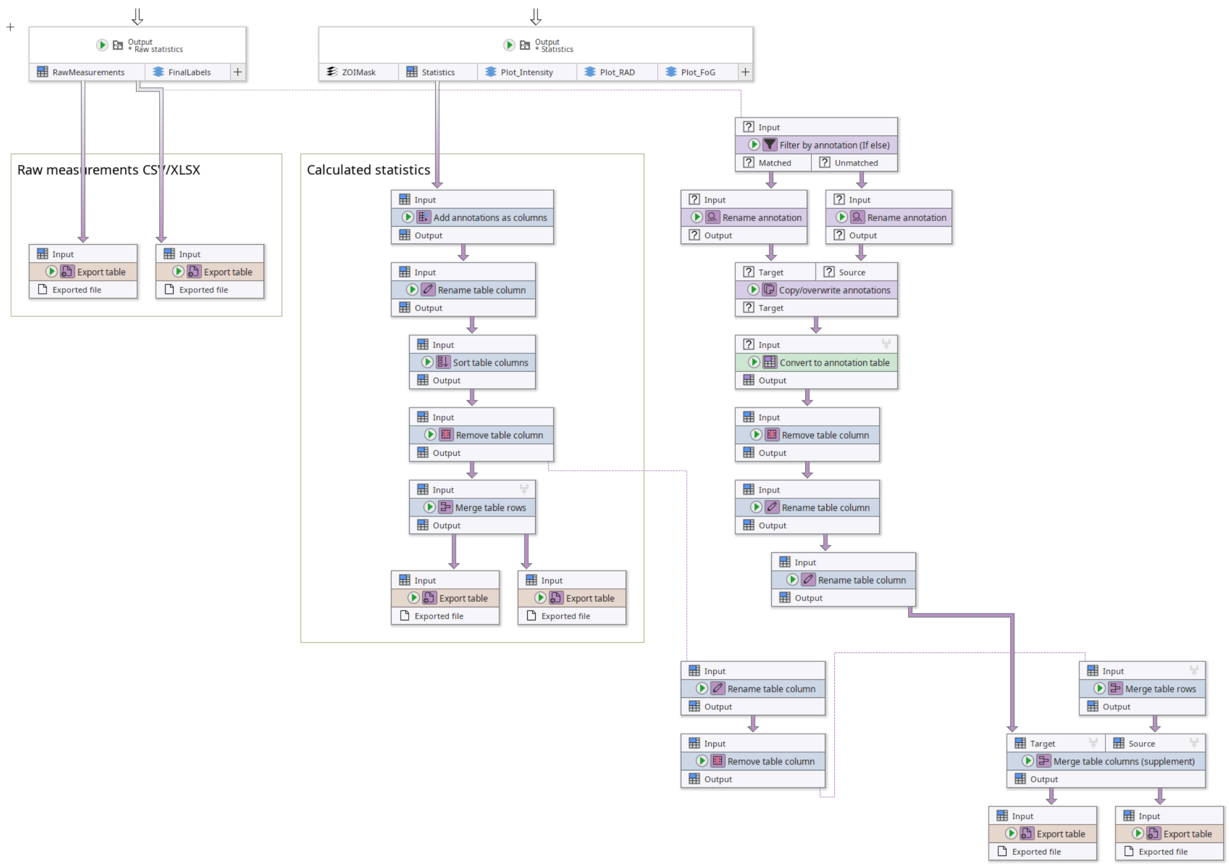

**Supplementary Figure 32:** Contents of the “Save tables” compartment of our DiskImageR re-implementation in JIPipe.

##### *Node #96 "Output" of type "Compartment output"*

This node passes results from another compartment into the current one.

- Input "RawMeasurements" of node #96 receives data from output "Output" of node #94
- Input "FinalLabels" of node #96 receives data from output "Output" of node #83
- The output slot name is set to **"Output"**

##### *Node #143 "Export table"*

This node exports a table into a CSV file.

- Input "Input" of node #143 receives data from output "RawMeasurements" of node #96
- The parameter "File path" (file-path) is set to **`PATH_COMBINE(project_data_dir.tmp_dir, "results", "raw_measurements", #AssayType + "_" + #Experiment + "_" + #Sample + "_" + #TimePoint)`**

##### *Node #144 "Export table"*

This node exports a table into an Excel file.

- Input "Input" of node #144 receives data from output "RawMeasurements" of node #96
- The parameter "File format" (file-format) is set to **"XLSX"**
- The parameter "File path" (file-path) is set to **`PATH_COMBINE(project_data_dir.tmp_dir, "results",`**

**"raw_measurements", #AssayType + "_" + #Experiment + "_" + #Sample + "_" + #TimePoint)**

##### *Node #145 "Filter by annotation (If else)"*

This node filters only the early time point from the raw measurements.

- Input "Input" of node #145 receives data from output "RawMeasurements" of node #96
- The parameter "Only match data if" (filter) is set to **`GroupColumn == "0"`**

##### *Node #146 "Rename annotation"*

This node renames the image ID and time point annotations.

- Input "Input" of node #146 receives data from output "Matched" of node #145
- The parameter item #1 of "Renaming items" (renaming-items) is set to **("#ImageId", "ImageId/Early")**
- The parameter item #2 of "Renaming items" (renaming-items) is set to **("#TimePoint", "TimePoint")**

##### *Node #147 "Rename annotation"*

This node renames the image ID and time point annotations.

- Input "Input" of node #147 receives data from output "Unmatched" of node #145
- The parameter item #1 of "Renaming items" (renaming-items) is set to **("#ImageId", "ImageId/Late")**
- The parameter item #2 of "Renaming items" (renaming-items) is set to **("#TimePoint", "TimePoint")**

##### *Node #148 "Copy/overwrite annotations"*

This node copies annotations between two data flows.

- Input "Target" of node #148 receives data from output "Output" of node #146
- Input "Source" of node #148 receives data from output "Output" of node #147
- The parameter "Remove existing annotations" (remove-existing-annotations-filter) is set to **false**
- The parameter "Selected source annotations" (source-annotation-filter) is set to **true**

##### *Node #149 "Convert to annotation table"*

This node converts the annotations of the incoming flow into a table.

- Input "Input" of node #149 receives data from output "Target" of node #148
- The parameter "Grouping method" in category "Merging iteration step generation" (jipipe:data-batch-generation/column-matching) is set to **"MergeAll"**

##### *Node #150 "Remove table column"*

This node removes unnecessary table columns.

- Input "Input" of node #150 receives data from output "Output" of node #149
- The parameter "Filters" (filters) is set to **`value IN ARRAY("GroupColumn", "#GroupRow", "MinGroupColumn", "TimePoint")`**

##### *Node #151 "Rename table column"*

This node removes the “#” in front of table column names where applicable.

- Input "Input" of node #151 receives data from output "Output" of node #150
- The parameter item #1 of "Renaming entries" (renaming-entries) is set to **source-column =** **`STRING_STARTS_WITH(value, "#")`**, **new-name =** **`SUBSTRING(current_name, 1)`**

*Node #152 "Rename table column"*

This node ensures that a column starting with “ImageId” starts with “#ImageId”. This is needed for enhanced J-AST visualization.

- Input "Input" of node #152 receives data from output "Output" of node #151
- The parameter item #1 of "Renaming entries" (renaming-entries) is set to **source-column =** **`STRING_STARTS_WITH(value, "ImageId")`**, **new-name =** **`"#" + current_name`**

*Node #112 "Output" of type "Compartment output"*

This node passes results from another compartment into the current one.

- Input "ZOIMask" of node #112 receives data from output "Output" of node #110
- Input "ZOIMask" of node #112 receives data from output "Output" of node #111
- Input "Statistics" of node #112 receives data from output "Output" of node #107
- Input "Plot_Intensity" of node #112 receives data from output "Plot_Intensity" of node #106
- Input "Plot_RAD" of node #112 receives data from output "Plot_RAD" of node #106
- Input "Plot_FoG" of node #112 receives data from output "Plot_FoG" of node #106
- The output slot name is set to **"Output"**

*Node #153 "Add annotations as columns"*

This node copies annotation into the currently processed table.

- Input "Input" of node #153 receives data from output "Statistics" of node #112
- The parameter "Annotation name filter" (annotation-name-filter) is set to **`value IN ARRAY("#Experiment", "#AssayType", "#Threshold", "#Sample", "PixelSize")`**

*Node #154 "Rename table column"*

This node removes the “#” in front of table column names where applicable.

- Input "Input" of node #154 receives data from output "Output" of node #153
- The parameter item #1 of "Renaming entries" (renaming-entries) is set to **source-column =** **`STRING_STARTS_WITH(value, "#")`**, **new-name =** **`SUBSTRING(current_name, 1)`**

*Node #155 "Sort table columns"*

This node ensures that the table columns follow a specific order.

- Input "Input" of node #155 receives data from output "Output" of node #154
- The parameter item #1 of "Manual order" (manual-order) is set to **"AssayType"**
- The parameter item #2 of "Manual order" (manual-order) is set to **"Experiment"**

- The parameter item #3 of "Manual order" (manual-order) is set to **"Sample"**
- The parameter item #4 of "Manual order" (manual-order) is set to **"PixelSize"**
- The parameter item #5 of "Manual order" (manual-order) is set to **"Threshold"**

*Node #156 "Remove table column"*

This node removes the “PixelSize” table column.

- Input "Input" of node #156 receives data from output "Output" of node #155
- The parameter "Filters" (filters) is set to **`value IN ARRAY("t", "PixelSize")`**

*Node #157 "Merge table rows"*

This node creates an all-in-one table.

- Input "Input" of node #157 receives data from output "Output" of node #156
- The parameter "Grouping method" in category "Merging iteration step generation" (jipipe:data-batch-generation/column-matching) is set to **"MergeAll"**

*Node #158 "Rename table column"*

This node renames table columns to the desired final name.

- Input "Input" of node #158 receives data from output "Output" of node #156
- The parameter item #1 of "Renaming entries" (renaming-entries) is set to **source-column = `value IN ARRAY("deltaFoG", "earlyFoG", "lateFoG", "earlyLateFoG")`, new-name = `current_name + TO_INTEGER(NUM(#Threshold) * 100)`**
- The parameter item #2 of "Renaming entries" (renaming-entries) is set to **source-column = `"earlyRAD_mm"`, new-name = `"earlyRAD" + TO_INTEGER(NUM(#Threshold) * 100) + "_mm"`**
- The parameter item #3 of "Renaming entries" (renaming-entries) is set to **source-column = `"lateRAD_mm"`, new-name = `"lateRAD" + TO_INTEGER(NUM(#Threshold) * 100) + "_mm"`**
- The parameter item #4 of "Renaming entries" (renaming-entries) is set to **source-column = `"earlyRAD_px"`, new-name = `"earlyRAD" + TO_INTEGER(NUM(#Threshold) * 100) + "_px"`**
- The parameter item #5 of "Renaming entries" (renaming-entries) is set to **source-column = `"lateRAD_px"`, new-name = `"lateRAD" + TO_INTEGER(NUM(#Threshold) * 100) + "_px"`**

*Node #159 "Export table"*

This node exports the incoming table as an Excel file.

- Input "Input" of node #159 receives data from output "Output" of node #157
- The parameter "File format" (file-format) is set to **"XLSX"**
- The parameter "File path" (file-path) is set to **`PATH_COMBINE(project_data_dir.tmp_dir, "results", "statistics")`**

*Node #160 "Export table"*

This node exports the incoming table as a CSV file.

- Input "Input" of node #160 receives data from output "Output" of node #157

- The parameter "File path" (file-path) is set to **`PATH_COMBINE(project_data_dir.tmp_dir, "results", "statistics")`**

##### *Node #161 "Remove table column"*

This node removes all remaining internal thresholding information from the results.

- Input "Input" of node #161 receives data from output "Output" of node #158
- The parameter "Filters" (filters) is set to **`value IN ARRAY("Threshold", "threshold")`**

##### *Node #162 "Merge table rows"*

This node creates a large all-in-one table per threshold.

- Input "Input" of node #162 receives data from output "Output" of node #161
- The parameter "Grouping method" in category "Merging iteration step generation" (jipipe:data-batch-generation/column-matching) is set to **"Custom"**
- The parameter "Custom grouping columns" in category "Merging iteration step generation" (jipipe:data-batch-generation/custom-matched-columns-expression) is set to **`"#Threshold"`**

##### *Node #163 "Merge table columns (supplement)"*

This node adds columns from one table into another. The algorithm uses specific reference columns to align the data.

- Input "Target" of node #163 receives data from output "Output" of node #152
- Input "Source" of node #163 receives data from output "Output" of node #162
- The parameter "Column filter" (column-filter) is set to **`value != "PixelSize"`**
- The parameter "Reference columns" (reference-columns) is set to **`value IN ARRAY("Experiment", "AssayType", "Sample")`**
- The parameter "Grouping method" in category "Merging iteration step generation" (jipipe:data-batch-generation/column-matching) is set to **"MergeAll"**

##### *Node #164 "Export table"*

This node exports the incoming table as a CSV file.

- Input "Input" of node #164 receives data from output "Output" of node #163
- The parameter "File path" (file-path) is set to **`PATH_COMBINE(project_data_dir.tmp_dir, "results", "statistics_aio")`**

##### *Node #165 "Export table"*

This node exports the incoming table as an Excel file.

- Input "Input" of node #165 receives data from output "Output" of node #163
- The parameter "File format" (file-format) is set to **"XLSX"**
- The parameter "File path" (file-path) is set to **`PATH_COMBINE(project_data_dir.tmp_dir, "results", "statistics_aio")`**

### 2.7 DiskImageR re-implementation (single image)

The following steps are applied by our single-image re-implementation of the DiskImageR algorithm in JIPipe. An overview can be found in **Fig. 3**. The structure of the pipeline is shown in **Supplementary Figure 25**.

###### 2.7.1.1 Compartment C1 "Data handling"

This compartment is responsible for organizing and importing the input images and associated metadata. A Screenshot can be found in **Supplementary Figure 26.**

***Node #1 "Folder list"***

This node references the directory that contains the DDA disk and Etest strip annotation masks.

- The parameter item #1 of "Folder paths" (folder-paths) is set to **"strip-disk"**

***Node #2 "Folder list"***

This node references the directory that contains the ZOI shape annotation masks.

- The parameter item #1 of "Folder paths" (folder-paths) is set to **"zoi-shape"**

***Node #3 "File"***

This node references a CSV file that associates the image ID to its corresponding image metadata.

- The parameter "File name" (file-name) is set to **"metadata.csv"**

***Node #4 "Folder list"***

This node references a directory that contains the plate annotation masks.

- The parameter item #1 of "Folder paths" (folder-paths) is set to **"plate"**

***Node #5 "Folder list"***

This node references a directory that contains the input images.

- The parameter item #1 of "Folder paths" (folder-paths) is set to **"raw"**

***Node #6 "List files"***

This node lists all PNG files in the input directory.

- Input "Folders" of node #6 receives data from output "Folder paths" of node #1
- The parameter "Keep file if ..." (filters) is set to **`STRING_MATCHES_GLOB(name, "*.png")`**

***Node #7 "List files"***

This node lists all PNG files in the input directory.

- Input "Folders" of node #7 receives data from output "Folder paths" of node #2

- The parameter "Keep file if ..." (filters) is set to **`STRING_MATCHES_GLOB(name, "*.png")`**

***Node #8 "Import results table"***

This node imports a CSV file as an ImageJ table.

- Input "Files" of node #8 receives data from output "Filename" of node #3

***Node #9 "List files"***

This node lists all PNG files in the input directory.

- Input "Folders" of node #9 receives data from output "Folder paths" of node #4
- The parameter "Keep file if ..." (filters) is set to **`STRING_MATCHES_GLOB(name, "*.png")`**

***Node #10 "List files"***

This node lists all PNG files in the input directory.

- Input "Folders" of node #10 receives data from output "Folder paths" of node #5
- The parameter "Keep file if ..." (filters) is set to **`STRING_MATCHES_GLOB(name, "*.png")`**

***Node #11 "Add path to annotations" of type "Annotate with file/directory name"***

This node attaches the image ID to the incoming file.

- Input "Paths" of node #11 receives data from output "Files" of node #6
- The parameter "Generated annotation" (generated-annotation) is set to **"#ImageId"**

***Node #12 "Add path to annotations" of type "Annotate with file/directory name"***

This node attaches the image ID to the incoming file.

- Input "Paths" of node #12 receives data from output "Files" of node #7
- The parameter "Generated annotation" (generated-annotation) is set to **"#ImageId"**

***Node #13 "Make row important" of type "Rename table column"***

This node renames the annotation “GroupRow” into “#GroupRow”, meaning that JIPipe uses that information to distinguish between data.

- Input "Input" of node #13 receives data from output "Results table" of node #8
- The parameter item #1 of "Renaming entries" (renaming-entries) is set to **source-column = "GroupRow", new-name = "#GroupRow"**

***Node #14 "Add path to annotations" of type "Annotate with file/directory name"***

This node attaches the image ID to the incoming file.

- Input "Paths" of node #14 receives data from output "Files" of node #9
- The parameter "Generated annotation" (generated-annotation) is set to **"#ImageId"**

***Node #15 "Add path to annotations" of type "Annotate with file/directory name"***

This node attaches the image ID to the incoming file.

- Input "Paths" of node #15 receives data from output "Files" of node #10
- The parameter "Generated annotation" (generated-annotation) is set to **"#ImageId"**

***Node #16 "Apply expression per row"***

This node finds the minimum value of the “GroupColumn” column in the input table and writes the result in a dedicated column “MinGroupColumn”.

- Input "Input" of node #16 receives data from output "Output" of node #13
- The parameter item #1 of "Generated values" (entries) is set to **column-name = "MinGroupColumn", value = `MIN(all.GroupColumn)`**

***Node #17 "Annotate by annotation table"***

This node attaches metadata contained in a table to the incoming data.

- Input "Annotations" of node #17 receives data from output "Output" of node #16
- Input "Data" of node #17 receives data from output "Annotated paths" of node #11

***Node #18 "Annotate by annotation table"***

This node attaches metadata contained in a table to the incoming data.

- Input "Annotations" of node #18 receives data from output "Output" of node #16
- Input "Data" of node #18 receives data from output "Annotated paths" of node #15

***Node #19 "Export table"***

This node exports the incoming table as a CSV file.

- Input "Input" of node #19 receives data from output "Output" of node #16
- The parameter "File path" (file-path) is set to **`PATH_COMBINE(project_data_dir.tmp_dir, "results", "metadata")`**

***Node #20 "Export table"***

This node exports the incoming table as an Excel file.

- Input "Input" of node #20 receives data from output "Output" of node #16
- The parameter "File format" (file-format) is set to **"XLSX"**
- The parameter "File path" (file-path) is set to **`PATH_COMBINE(project_data_dir.tmp_dir, "results", "metadata")`**

***Node #21 "Annotate by annotation table"***

This node attaches metadata contained in a table to the incoming data.

- Input "Annotations" of node #21 receives data from output "Output" of node #16
- Input "Data" of node #21 receives data from output "Annotated paths" of node #14

***Node #22 "Annotate by annotation table"***

This node attaches metadata contained in a table to the incoming data.

- Input "Annotations" of node #22 receives data from output "Output" of node #16
- Input "Data" of node #22 receives data from output "Annotated paths" of node #12

***Node #23 "Check iteration steps (multiple data per slot)"***

This node checks if all necessary data is available and throws an error if this is not the case.

- Input "SD" of node #23 receives data from output "Annotated data" of node #17
- Input "ZS" of node #23 receives data from output "Annotated data" of node #22
- Input "PL" of node #23 receives data from output "Annotated data" of node #21
- Input "RAW" of node #23 receives data from output "Annotated data" of node #18
- The parameter "Grouping method" in category "Merging iteration step generation" (jipipe:data-batch-generation/column-matching) is set to **"Custom"**
- The parameter "Custom grouping columns" in category "Merging iteration step generation" (jipipe:data-batch-generation/custom-matched-columns-expression) is set to **value == "#GroupRow"**

***Node #24 "Import image (SD)" of type "Import image"***
This node imports a PNG file as an ImageJ image.

- Input "Files" of node #24 receives data from output "SD" of node #23

***Node #25 "Import image (ZS)" of type "Import image"***
This node imports a PNG file as an ImageJ image.

- Input "Files" of node #25 receives data from output "ZS" of node #23

***Node #26 "Import image (PL)" of type "Import image"***
This node imports a PNG file as an ImageJ image.

- Input "Files" of node #26 receives data from output "PL" of node #23

***Node #27 "Import image (RAW)" of type "Import image"***
This node imports a PNG file as an ImageJ image.

- Input "Files" of node #27 receives data from output "RAW" of node #23

***Node #28 "Output" of type "Compartment output"***
This node passes the data to other compartments.

- Input "SD" of node #28 receives data from output "Image" of node #24
- Input "ZS" of node #28 receives data from output "Image" of node #25
- Input "PL" of node #28 receives data from output "Image" of node #26
- Input "RAW" of node #28 receives data from output "Image" of node #27
- The output slot name is set to **"Output"**

###### 2.7.1.2 Compartment C2 "Align ZOI Shape"

This compartment applies no workload, as it is a remnant of the conversion process from the timeline to the single-image analysis. A screenshot can be found in **Supplementary Figure 33**.

- The "Align ZOI Shape" compartment (C2) receives data from the "Data handling" compartment (C1)

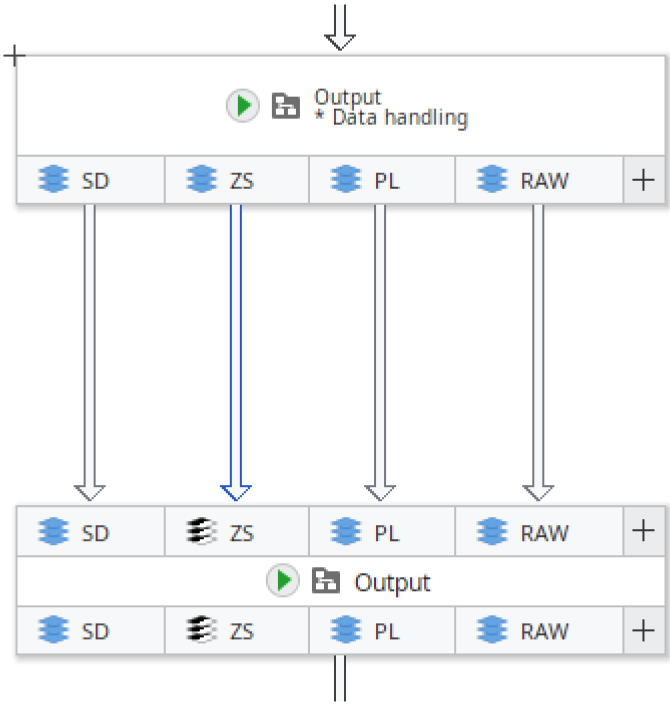

**Supplementary Figure 33:** Contents of the "Align ZOI Shape" compartment of our single-image DiskImageR implementation in JIPipe.

*Node #28 "Output" of type "Compartment output"*

- Input "SD" of node #28 receives data from output "Image" of node #24
- Input "ZS" of node #28 receives data from output "Image" of node #25
- Input "PL" of node #28 receives data from output "Image" of node #26
- Input "RAW" of node #28 receives data from output "Image" of node #27
- The output slot name is set to **"Output"**

*Node #29 "Output" of type "Compartment output"*

- Input "SD" of node #29 receives data from output "SD" of node #28
- Input "ZS" of node #29 receives data from output "ZS" of node #28
- Input "PL" of node #29 receives data from output "PL" of node #28
- Input "RAW" of node #29 receives data from output "RAW" of node #28
- The output slot name is set to **"Output"**

##### 2.7.1.3 Compartment C3 "Generate labels"

This compartment is responsible for generating the measurement labels that are used to extract the colony intensities over a radius axis. A Screenshot can be found in **Supplementary Figure 28**.

- The "Generate labels" compartment (C3) receives data from the "Align ZOI Shape" compartment (C2)

*Node #29 "Output" of type "Compartment output"*

This node passes outputs from another compartment into the current one.

- Input "SD" of node #29 receives data from output "SD" of node #28
- Input "ZS" of node #29 receives data from output "ZS" of node #28
- Input "PL" of node #29 receives data from output "PL" of node #28
- Input "RAW" of node #29 receives data from output "RAW" of node #28
- The output slot name is set to **"Output"**

##### *Node #30 "Erode plate" of type "Morphological operation 2D"*

This node erodes the plate by 10 mm to avoid noise introduced by the plate borders.

- Input "Input" of node #30 receives data from output "PL" of node #29
- The parameter "Add border before applying operation" (add-border) is set to **true**
- The parameter "Operation" (operation) is set to **"EROSION"**
- The parameter "Attach parameter annotations" in category "Adaptive parameters" (jipipe:adaptive-parameters/attach-parameter-annotations) is set to **false**
- The parameter item #1 of "Overridden parameters" in category "Adaptive parameters" (jipipe:adaptive-parameters/overridden-parameters) is set to **`(_global.plateShaveOffMillimeters / NUM(PixelSize), "radius")`**
- The parameter "Margin right" in category "Border settings" (border-parameters/margin-right) is set to **100**
- The parameter "Margin left" in category "Border settings" (border-parameters/margin-left) is set to **100**
- The parameter "Margin top" in category "Border settings" (border-parameters/margin-top) is set to **100**
- The parameter "Margin bottom" in category "Border settings" (border-parameters/margin-bottom) is set to **100**

##### *Node #31 "DDA branch" of type "Check iteration steps (one data per slot)"*

This node ensures that all required data for the following steps are present.

- Input "SD" of node #31 receives data from output "SD" of node #29
- Input "ZS" of node #31 receives data from output "ZS" of node #29
- Input "RAW" of node #31 receives data from output "RAW" of node #29
- The parameter "Keep iteration step if" (filter) is set to **`#AssayType != "ETest"`**

##### *Node #32 "E-Test branch" of type "Check iteration steps (one data per slot)"*

This node ensures that all required data for the following steps are present.

- Input "SD" of node #32 receives data from output "SD" of node #29
- Input "ZS" of node #32 receives data from output "ZS" of node #29
- Input "RAW" of node #32 receives data from output "RAW" of node #29
- The parameter "Keep iteration step if" (filter) is set to **`#AssayType == "ETest"`**

##### *Node #33 "Set annotation (expression)"*

This node calculates the expected label thickness in pixels given the physical size of a pixel and the global parameter (1 mm).

- Input "Input" of node #33 receives data from output "SD" of node #31
- The parameter "Annotation name" (annotation-name) is set to **"LabelThickness"**
- The parameter "Annotation value" (annotation-value) is set to **`MAX(3, ROUND(_global.labelWidthMillimeters * (1 / NUM(PixelSize))))`**

*Node #34 "Set annotation (expression)"*

This node calculates the expected label thickness in pixels given the physical size of a pixel and the global parameter (1 mm).

- Input "Input" of node #34 receives data from output "ZS" of node #32
- The parameter "Annotation name" (annotation-name) is set to **"LabelThickness"**
- The parameter "Annotation value" (annotation-value) is set to **`MAX(3, ROUND(_global.labelWidthMillimeters * (1 / NUM(PixelSize)))`**

*Node #35 "Auto threshold 2D"*

This node applies an auto-thresholding algorithm.

- Input "Input" of node #35 receives data from output "Output" of node #33

*Node #36 "Auto threshold 2D"*

This node applies an auto-thresholding algorithm.

- Input "Input" of node #36 receives data from output "Output" of node #34

*Node #37 "Invert colors" of type "Invert pixel colors/values"*

This node inverts the greyscale color values (255 - x).

- Input "Input" of node #37 receives data from output "Output" of node #35

*Node #38 "Euclidean distance transform 2D"*

This node applies an Euclidean distance transform.

- Input "Input" of node #38 receives data from output "Output" of node #36

*Node #39 "Morphological operation 2D"*

This node applies a morphological internal gradient operation with radius 1 px, converting the ZOI shape area into an outline.

- Input "Input" of node #39 receives data from output "Output" of node #36
- The parameter "Operation" (operation) is set to **"INTERNAL_GRADIENT"**

*Node #40 "Euclidean distance transform 2D"*

This node applies an Euclidean distance transform.

- Input "Input" of node #40 receives data from output "Output" of node #37

*Node #41 "Local maxima 2D"*

This node finds all local maxima with a height tolerance of 10.

- Input "Input" of node #41 receives data from output "Output" of node #38

*Node #42 "Invert colors" of type "Invert pixel colors/values"*

This node inverts the greyscale color values (255 - x).

- Input "Input" of node #42 receives data from output "Output" of node #39

*Node #43 "Fast image arithmetics"*

This node rounds the distance transform output so that the thickness of the created bins is approximately 1 mm.

- Input "I1" of node #43 receives data from output "Output" of node #40
- The parameter "Expression" (expression) is set to **`(ROUND(I1 / LabelThickness) * LabelThickness) + 1`**

*Node #44 "Find particles 2D"*

This node uses the ImageJ particle finder to generate ROI lists.

- Input "Mask" of node #44 receives data from output "Output" of node #41

*Node #45 "Expand canvas 2D"*

This node expands the image canvas size and fills created regions with white pixels. This ensures that no disconnected labels are generated.

- Input "Input" of node #45 receives data from output "Output" of node #42
- The parameter "Background color" (background-color) is set to **"#FFFFFF"**
- The parameter "Y axis" (y-axis) is set to **`default * 2`**
- The parameter "X axis" (x-axis) is set to **`default * 2`**

*Node #46 "2D ROI calculator (AND/OR/XOR)"*

This node merges all ROIs within a ROI list into a single composite ROI.

- Input "Input" of node #46 receives data from output "ROI" of node #44
- The parameter "Split after operation" (split-afterwards) is set to **false**
- The parameter "Operation" (operation) is set to **"LogicalOr"**

*Node #47 "Euclidean distance transform 2D"*

This node applies an Euclidean distance transform.

- Input "Input" of node #47 receives data from output "Output" of node #45

*Node #48 "Outline 2D ROI (Centroid)"*

This node finds the centroid of the all-in-one ROI generated by node #46.

- Input "Input" of node #48 receives data from output "Output" of node #46

*Node #49 "Fast image arithmetics"*

This node rounds the distance transform output so that the thickness of the created bins is approximately 1 mm.

- Input "I1" of node #49 receives data from output "Output" of node #47
- The parameter "Expression" (expression) is set to **`(ROUND(I1 / LabelThickness) * LabelThickness) + 1`**

*Node #50 "Convert 2D ROI to mask"*

This node converts the incoming ROI lists into masks.

- Input "ROI" of node #50 receives data from output "Output" of node #48
- Input "Image" of node #50 receives data from output "Output" of node #41

*Node #51 "Split labels (connected components) 2D"*

This node ensures that labels that have the same value, but are not in the same connected component have a different value.

- Input "Input" of node #51 receives data from output "Output" of node #49

*Node #52 "Invert colors" of type "Invert pixel colors/values"*

This node inverts the greyscale image value.

- Input "Input" of node #52 receives data from output "Output" of node #50

*Node #53 "Crop 2D image"*

This node reverts the operation applied by node #45.

- Input "Input" of node #53 receives data from output "Output" of node #51
- The parameter "ROI" (roi) is set to **{"left":{"expression":"`width / 4`"},"top":{"expression":"`height / 4`"},"right":{"expression":"`0`"},"bottom":{"expression":"`0`"},"width":{"expression":"`width / 2`"},"height":{"expression":"`height / 2`"},"anchor":"TopLeft"}**

*Node #54 "Euclidean distance transform 2D"*

This node applies an Euclidean distance transform.

- Input "Input" of node #54 receives data from output "Output" of node #52

*Node #55 "Set to value (grayscale)"*

This node erases the strip/disk from the input image.

- Input "Input" of node #55 receives data from output "Output" of node #53
- Input "Mask" of node #55 receives data from output "SD" of node #32
- The parameter "Only apply to ..." (roi:target-area) is set to **"InsideMask"**

*Node #56 "Extract label statistics 2D"*

This node measures the mean colony intensity for each measurement label.

- Input "Image" of node #56 receives data from output "Output" of node #54
- Input "Labels" of node #56 receives data from output "Output" of node #55
- The parameter "Measure in physical units" (measure-in-physical-units) is set to **false**
- The parameter "Measurements" (measurements) is set to **{"values":["PixelValueMean"],"collapsed":false}**

*Node #57 "Replace label values by table"*

This node re-maps label values from a table, making the labels correspond to the radius value.

- Input "Mappings" of node #57 receives data from output "Output" of node #56
- Input "Labels" of node #57 receives data from output "Output" of node #55
- The parameter "Old label column" (old-label-column) is set to **("ExistingColumn", "label_id")**
- The parameter "New label column" (new-label-column) is set to **("ExistingColumn", "Mean")**

***Node #58 "All labels (DDA + E-Test)" of type "IO Interface"***

This node merges the branches for DDA and Etest.

- Input "LB" of node #58 receives data from output "Output" of node #43
- Input "LB" of node #58 receives data from output "Labels" of node #57

***Node #59 "Remove plate" of type "Set to value (grayscale)"***

This node removes the plate from the measurement labels.

- Input "Input" of node #59 receives data from output "LB" of node #58
- Input "Mask" of node #59 receives data from output "Output" of node #30
- The parameter "Only apply to ..." (roi:target-area) is set to **"OutsideMask"**

***Node #60 "Round float image"***

This node ensures that all label values are integral.

- Input "Input" of node #60 receives data from output "Output" of node #59
- The parameter "Number of decimals" (decimals) is set to **0**

***Node #61 "Remove annotation"***

This node removes the pre-calculated label thickness annotation that was added in an earlier step.

- Input "Input" of node #61 receives data from output "Output" of node #60
- The parameter "Removed annotations" (annotation-type) is set to **key == "LabelThickness"**

***Node #62 "Output" of type "Compartment output"***

This node passes the measurement labels into other compartments.

- Input "Labels" of node #62 receives data from output "Output" of node #61
- The output slot name is set to **"Output"**

###### 2.7.1.4 Compartment C4 "Raw statistics"

This compartment is responsible for measuring the colony intensities using the measurement labels. A screenshot can be found in **Supplementary Figure 25**.

- The "Raw statistics" compartment (C4) receives data from the "Generate labels" compartment (C3)
- The "Raw statistics" compartment (C4) receives data from the "Align ZOI Shape" compartment (C2)

***Node #29 "Output" of type "Compartment output"***

This node receives results from another compartment.

- Input "SD" of node #29 receives data from output "SD" of node #28
- Input "ZS" of node #29 receives data from output "ZS" of node #28
- Input "PL" of node #29 receives data from output "PL" of node #28
- Input "RAW" of node #29 receives data from output "RAW" of node #28
- The output slot name is set to **"Output"**

*Node #63 "Dilate SD" of type "Morphological operation 2D"*

This node dilates the DDA disk/Etest strip annotations by 1 mm.

- Input "Input" of node #63 receives data from output "SD" of node #29
- The parameter "Attach parameter annotations" in category "Adaptive parameters" (jipipe:adaptive-parameters/attach-parameter-annotations) is set to **false**
- The parameter item #1 of "Overridden parameters" in category "Adaptive parameters" (jipipe:adaptive-parameters/overridden-parameters) is set to **(`1 / NUM(PixelSize)`, `"radius"`)**
- The parameter "Margin right" in category "Border settings" (border-parameters/margin-right) is set to **100**
- The parameter "Margin left" in category "Border settings" (border-parameters/margin-left) is set to **100**
- The parameter "Margin top" in category "Border settings" (border-parameters/margin-top) is set to **100**
- The parameter "Margin bottom" in category "Border settings" (border-parameters/margin-bottom) is set to **100**

*Node #64 "Convert image to 8-bit"*

This node ensures that the input image is 8-bit greyscale.

- Input "Input" of node #64 receives data from output "RAW" of node #29

*Node #65 "Delete SD (Content aware)" of type "Set to content aware 2D"*

This node erases the DDA disk/Etest strip using a content-aware fill. Compared to erasing by setting the pixels to a specific color, this minimizes the impact on the measurements.

- Input "Mask" of node #65 receives data from output "Output" of node #63
- Input "Input" of node #65 receives data from output "Output" of node #64

*Node #62 "Output" of type "Compartment output"*

This node receives results from another compartment.

- Input "Labels" of node #62 receives data from output "Output" of node #61
- The output slot name is set to **"Output"**

*Node #66 "Set to value (grayscale)"*

This node erases the DDA disk/Etest strip from the input image.

- Input "Input" of node #66 receives data from output "Labels" of node #62
- Input "Mask" of node #66 receives data from output "Output" of node #63
- The parameter "Only apply to ..." (roi:target-area) is set to **"InsideMask"**

*Node #67 "Extract label statistics 2D"*

This node measures for each label the mean colony intensity.

- Input "Labels" of node #67 receives data from output "Labels" of node #62
- Input "Image" of node #67 receives data from output "Output" of node #65
- The parameter "Measurements" (measurements) is set to **{"values":["PixelValueMean"],"collapsed":false}**

***Node #68 "Binarize"***

This node converts the labels into a binary mask.

- Input "Input" of node #68 receives data from output "Labels" of node #62

***Node #69 "Rename table column"***

This node renames the label ID column into “r_px”. This works because the label IDs already correspond to the radius.

- Input "Input" of node #69 receives data from output "Output" of node #67
- The parameter item #1 of "Renaming entries" (renaming-entries) is set to **source-column = "label_id", new-name = "r_px"**
- The parameter item #2 of "Renaming entries" (renaming-entries) is set to **source-column = "Mean", new-name = "mean"**

***Node #70 "Extract label statistics 2D"***

This node measures for each label the minimum and maximum colony intensity.

- Input "Labels" of node #70 receives data from output "Output" of node #68
- Input "Image" of node #70 receives data from output "Output" of node #64
- The parameter "Measurements" (measurements) is set to **{"values":["PixelValueMinMax"],"collapsed":false}**

***Node #71 "Apply expression per row"***

This node converts the radius (pixels) into radius (mm).

- Input "Input" of node #71 receives data from output "Output" of node #69
- The parameter item #1 of "Generated values" (entries) is set to **column-name = "r_mm", value = r_px * NUM(annotations @ "PixelSize")**

***Node #72 "Rename table column"***

This node renames the “Min” column into “background_intensity”.

- Input "Input" of node #72 receives data from output "Output" of node #70
- The parameter item #1 of "Renaming entries" (renaming-entries) is set to **source-column = "Min", new-name = "background_intensity"**

***Node #73 "Remove table column"***

This node renames the “Min” column into “background_intensity”.

- Input "Input" of node #73 receives data from output "Output" of node #72
- The parameter "Filters" (filters) is set to **value != "background_intensity"**

***Node #74 "Merge table columns (simple)”***

This node merges multiple tables column-wise.

- Input "Input" of node #74 receives data from output "Output" of node #71
- Input "Input" of node #74 receives data from output "Output" of node #73
- The parameter "Row normalization" (row-normalization) is set to **"LastValue"**

***Node #75 "Apply expression per row"***

This node finds the minimum and maximum colony intensities.

- Input "Input" of node #75 receives data from output "Output" of node #74
- The parameter item #1 of "Generated values" (entries) is set to **column-name = "min_intensity", value = `MIN(all.mean)`**
- The parameter item #2 of "Generated values" (entries) is set to **column-name = "max_intensity", value = `MAX(all.mean)`**

##### *Node #76 "Apply expression per row"*

This node calculates the relative colony intensity (rmean) and its related intensity reduction (reduction).

- Input "Input" of node #76 receives data from output "Output" of node #75
- The parameter item #1 of "Generated values" (entries) is set to **column-name = "rmean", value = `ABS(mean-background_intensity)/(max_intensity-background_intensity)`**
- The parameter item #2 of "Generated values" (entries) is set to **column-name = "reduction", value = `1 - rmean`**

##### *Node #77 "Sort table columns"*

This node sorts the table columns, so that the r_px, r_mm, and mean columns come first.

- Input "Input" of node #77 receives data from output "Output" of node #76
- The parameter item #1 of "Manual order" (manual-order) is set to **"r_px"**
- The parameter item #2 of "Manual order" (manual-order) is set to **"r_mm"**
- The parameter item #3 of "Manual order" (manual-order) is set to **"mean"**

##### *Node #78 "JFreeChart XY line plot"*

This node creates a plot of rmean over r_mm.

- Input "Input" of node #78 receives data from output "Output" of node #77
- The parameter "X" in category "Input columns" (input-columns/X) is set to **("ExistingColumn", "r_mm")**
- The parameter "Y" in category "Input columns" (input-columns/Y) is set to **("ExistingColumn", "rmean")**

##### *Node #79 "Output" of type "Compartment output"*

This node passes the raw measurements and labels to other compartments.

- Input "RawMeasurements" of node #79 receives data from output "Output" of node #77
- Input "FinalLabels" of node #79 receives data from output "Output" of node #66
- The output slot name is set to **"Output"**

##### *Node #80 "JFreeChart XY line plot"*

This node creates a plot of the reduction over r_mm.

- Input "Input" of node #80 receives data from output "Output" of node #77
- The parameter "X" in category "Input columns" (input-columns/X) is set to **("ExistingColumn", "r_mm")**

- The parameter "Y" in category "Input columns" (input-columns/Y) is set to **("ExistingColumn", "reduction")**

###### *Node #81 "Create montage"*

This plot merges multiple plots into a montage.

- Input "Input" of node #81 receives data from output "Output" of node #78
- The parameter "Force number of rows" in category "Montage" (montage-parameters/force-rows) is set to **1**
- The parameter "Grouping method" in category "Merging iteration step generation" (jipipe:data-batch-generation/column-matching) is set to **"Custom"**
- The parameter "Custom grouping columns" in category "Merging iteration step generation" (jipipe:data-batch-generation/custom-matched-columns-expression) is set to **"#GroupRow"**

###### *Node #82 "Create montage"*

This plot merges multiple plots into a montage.

- Input "Input" of node #82 receives data from output "Output" of node #80
- The parameter "Force number of rows" in category "Montage" (montage-parameters/force-rows) is set to **1**
- The parameter "Grouping method" in category "Merging iteration step generation" (jipipe:data-batch-generation/column-matching) is set to **"Custom"**
- The parameter "Custom grouping columns" in category "Merging iteration step generation" (jipipe:data-batch-generation/custom-matched-columns-expression) is set to **"#GroupRow"**

##### 2.7.1.5 Compartment C5 "Statistics"

This compartment is responsible for generating the output statistics, i.e. the RAD, FoG, and others. A screenshot of the pipeline can be found in **Supplementary Figure 34**.

- The "Statistics" compartment (C5) receives data from the "Generate labels" compartment (C3)
- The "Statistics" compartment (C5) receives data from the "Raw statistics" compartment (C4)

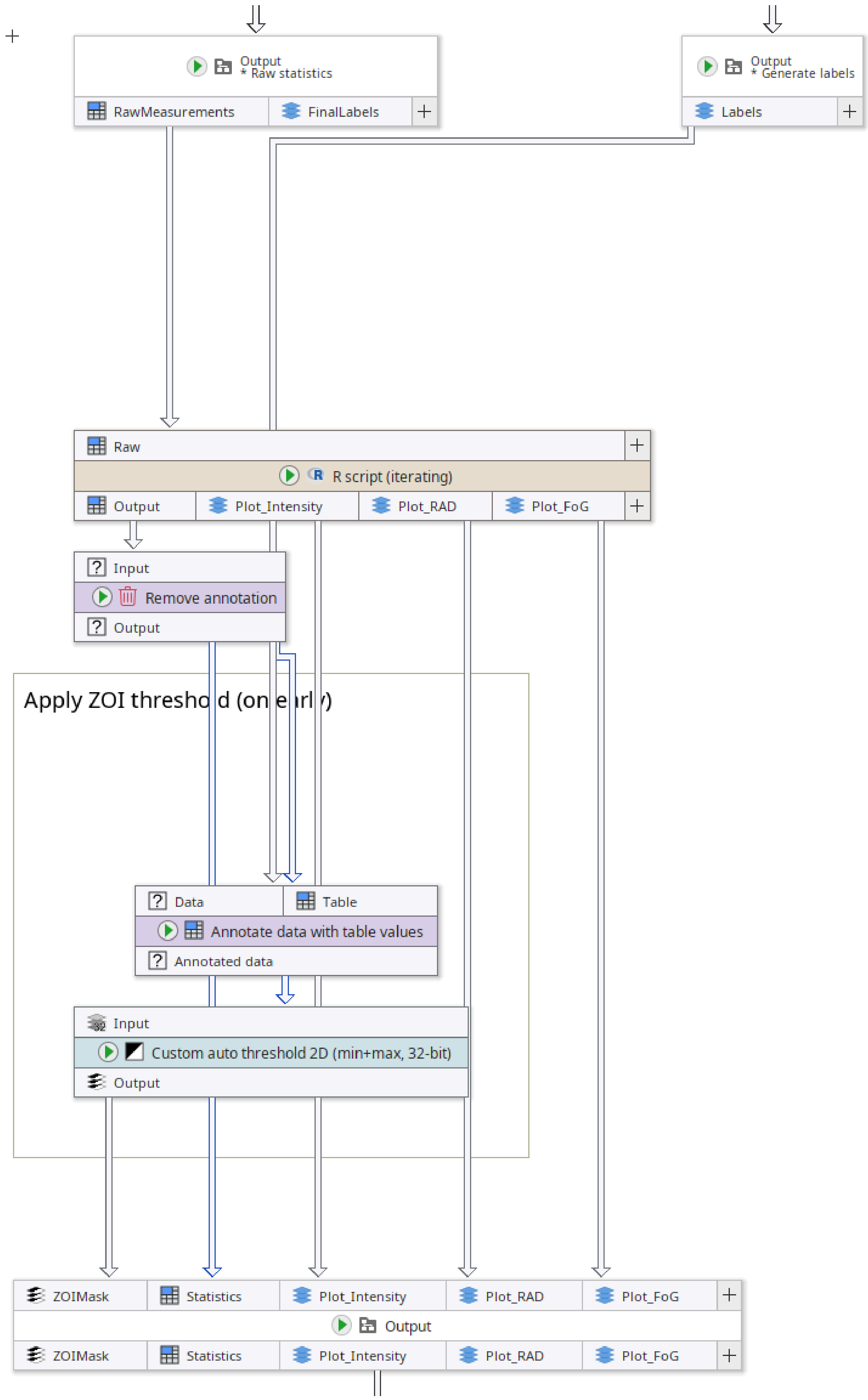

**Supplementary Figure 34:** Contents of the “Statistics” compartment of our single-image DiskImageR implementation in JIPipe.

*Node #62 "Output" of type "Compartment output"*

This node passes the labels from the “Generate labels” compartment into the current compartment.

- Input "Labels" of node #62 receives data from output "Output" of node #61
- The output slot name is set to **"Output"**

*Node #79 "Output" of type "Compartment output"*

This node passes the raw measurements and final labels from the “Raw statistics” compartment into the current one.

- Input "RawMeasurements" of node #79 receives data from output "Output" of node #77
- Input "FinalLabels" of node #79 receives data from output "Output" of node #66
- The output slot name is set to **"Output"**

*Node #83 "R script (iterating)"*

This node runs an R script that handles the calculation of the RAD and FoG values.

- Input "Raw" of node #83 receives data from output "RawMeasurements" of node #79
- The parameter "Override R environment" (override-environment) is set to **[Disabled]**
- The parameter item #1 of "Thresholds" in category "Script variables" (variables/thresholds) is set to **0.2**
- The parameter item #2 of "Thresholds" in category "Script variables" (variables/thresholds) is set to **0.5**
- The parameter item #3 of "Thresholds" in category "Script variables" (variables/thresholds) is set to **0.8**

The pipeline has two variants, one using numeric integration (default), and another using R’s function approximation. The following code is used by the variant using numeric integration:

```
library(ggplot2)
library(gridExtra)
library(reshape2)  # for melting data frames if needed

# Use a generic sans font to avoid font issues on Linux CLI
theme_set(theme_minimal(base_family = "sans"))

# Build dataset name from JIPipe annotations
dataset.name <- paste0(JIPipe.TextAnnotations[["#AssayType"]],
                       "_", JIPipe.TextAnnotations[["#Experiment"]],
                       "_", JIPipe.TextAnnotations[["#Sample"]])

# Retrieve the single data frame "Raw"
df <- JIPipe.GetInputAsDataFrame("Raw")

# Filter out rows with non-positive mean values
df <- df[df$mean > 0, ]

# Create an intensity profile of the raw frame
```

```r
plot.raw <- ggplot(df, aes(x = r_mm, y = rmean)) +
  geom_line() +
  labs(title = paste0(dataset.name, " - Raw Frame Intensity Profile"),
       x = "Distance (mm)", y = "Relative mean intensity") +
  scale_x_continuous(limits = c(0, max(df$r_mm)))

# Define thresholds to analyze
thresholds <- c(0.2, 0.5, 0.8)

# Colors for each threshold's vertical line (one per threshold)
line_colors <- c("red", "blue", "green")

# Robust AUC function using the trapezoidal rule on discrete data
calculate_auc <- function(x, y) {
  if(length(x) < 2 || length(y) < 2) {
    return(0) # Not enough points for integration
  }
  dx <- diff(x)
  y0 <- head(y, -1)
  y1 <- tail(y, -1)
  auc <- sum(dx * (y0 + y1) / 2)
  return(auc)
}

# Global results data frame for plotting
results_df <- data.frame()

# Loop over each threshold to compute measurements and output table per
threshold.
for(i in seq_along(thresholds)) {

  threshold <- thresholds[i]

  # Compute RAD as the first radius where rmean >= (1 - threshold)
  rad_mm <- df[df$rmean >= (1 - threshold), ]$r_mm[1]
  rad_px <- df[df$rmean >= (1 - threshold), ]$r_px[1]

  # Create an interpolation function based on r_px and rmean
  # f <- approxfun(x = df$r_px, y = df$rmean, method = "linear", rule = 2,
yright = 0)

  # Use only the portion of the data up to the calculated RAD (in pixels)
  df.filtered <- df[df$r_px <= rad_px, ]
  mri <- max(df.filtered$rmean)

  # Calculate the area under the curve (AUC) and the maximum possible area
(amax)
  # auc <- integrate(f, lower = 0, upper = rad_px, subdivisions =
```

```
2000)$value
  auc <- calculate_auc(df.filtered$r_px, df.filtered$rmean)
  amax <- rad_px * mri

  # FoG is the ratio of the AUC to the maximum area (amax)
  fog <- auc / amax

  # Create a results data frame for the current threshold
  df.out <- data.frame(threshold  = threshold,
                       RAD_mm     = rad_mm,
                       RAD_px     = rad_px,
                       FoG        = fog)

  # Append the current results to the global results (used for plotting
later)
  results_df <- rbind(results_df, df.out)

  # Output the data frame for this threshold using JIPipe with annotation
  JIPipe.AddOutputDataFrame(slot = "Output",
                            data = df.out,
                            annotations = list("#Threshold" =
as.character(threshold)))

  # Update the intensity plot: add a vertical dashed line marking the
computed RAD (in mm)
  plot.raw <- plot.raw +
    geom_vline(xintercept = rad_mm, color = line_colors[i], linetype =
"dashed", show.legend = FALSE)
}

### Save the combined intensity plot
plot_folder_intensity <- JIPipe.AddOutputFolder("Plot_Intensity")
ggsave(filename = paste0(plot_folder_intensity, "/data.png"),
       plot = plot.raw, width = 10, height = 6, dpi = 120)

### Create and save the RAD plot (RAD in mm vs. threshold)
rad_plot <- ggplot(results_df, aes(x = threshold, y = RAD_mm)) +
  geom_line() + geom_point() +
  labs(title = "RAD (mm) vs. Threshold", x = "Threshold", y = "RAD (mm)") +
  theme_minimal()
plot_folder_rad <- JIPipe.AddOutputFolder("Plot_RAD")
ggsave(filename = paste0(plot_folder_rad, "/data.png"),
       plot = rad_plot, width = 6, height = 4, dpi = 120)

### Create and save the FoG plot (FoG vs. threshold)
fog_plot <- ggplot(results_df, aes(x = threshold, y = FoG)) +
```

```
  geom_line() + geom_point() +
  labs(title = "FoG vs. Threshold", x = "Threshold", y = "FoG") +
  theme_minimal()
plot_folder_fog <- JIPipe.AddOutputFolder("Plot_FoG")
ggsave(filename = paste0(plot_folder_fog, "/data.png"),
       plot = fog_plot, width = 6, height = 4, dpi = 120)
```

The following code is used by the pipeline variant that utilizes R's function approximation algorithm for the integration step:

```
library(ggplot2)
library(gridExtra)
library(reshape2)  # for melting data frames if needed

# Use a generic sans font to avoid font issues on Linux CLI
theme_set(theme_minimal(base_family = "sans"))

# Build dataset name from JIPipe annotations
dataset.name <- paste0(JIPipe.TextAnnotations[["#AssayType"]],
                       "_", JIPipe.TextAnnotations[["#Experiment"]],
                       "_", JIPipe.TextAnnotations[["#Sample"]])

# Retrieve the single data frame "Raw"
df <- JIPipe.GetInputAsDataFrame("Raw")

# Filter out rows with non-positive mean values
df <- df[df$mean > 0, ]

# Create an intensity profile of the raw frame
plot.raw <- ggplot(df, aes(x = r_mm, y = rmean)) +
  geom_line() +
  labs(title = paste0(dataset.name, " - Raw Frame Intensity Profile"),
       x = "Distance (mm)", y = "Relative mean intensity") +
  scale_x_continuous(limits = c(0, max(df$r_mm)))

# Define thresholds to analyze
thresholds <- c(0.2, 0.5, 0.8)

# Colors for each threshold's vertical line (one per threshold)
line_colors <- c("red", "blue", "green")

# Global results data frame for plotting
results_df <- data.frame()

# Loop over each threshold to compute measurements and output table per
threshold.
```

```
for(i in seq_along(thresholds)) {

  threshold <- thresholds[i]

  # Compute RAD as the first radius where rmean >= (1 - threshold)
  rad_mm <- df[df$rmean >= (1 - threshold), ]$r_mm[1]
  rad_px <- df[df$rmean >= (1 - threshold), ]$r_px[1]

  # Create an interpolation function based on r_px and rmean
  f <- approxfun(x = df$r_px, y = df$rmean, method = "linear", rule = 2,
yright = 0)

  # Use only the portion of the data up to the calculated RAD (in pixels)
  df.filtered <- df[df$r_px <= rad_px, ]
  mri <- max(df.filtered$rmean)

  # Calculate the area under the curve (AUC) and the maximum possible area
(amax)
  auc <- integrate(f, lower = 0, upper = rad_px, subdivisions = 2000)$value
  amax <- rad_px * mri

  # FoG is the ratio of the AUC to the maximum area (amax)
  fog <- auc / amax

  # Create a results data frame for the current threshold
  df.out <- data.frame(threshold  = threshold,
                       RAD_mm     = rad_mm,
                       RAD_px     = rad_px,
                       FoG        = fog)

  # Append the current results to the global results (used for plotting
later)
  results_df <- rbind(results_df, df.out)

  # Output the data frame for this threshold using JIPipe with annotation
  JIPipe.AddOutputDataFrame(slot = "Output",
                            data = df.out,
                            annotations = list("#Threshold" =
as.character(threshold)))

  # Update the intensity plot: add a vertical dashed line marking the
computed RAD (in mm)
  plot.raw <- plot.raw +
    geom_vline(xintercept = rad_mm, color = line_colors[i], linetype =
"dashed", show.legend = FALSE)
}

### Save the combined intensity plot
```

```
plot_folder_intensity <- JIPipe.AddOutputFolder("Plot_Intensity")
ggsave(filename = paste0(plot_folder_intensity, "/data.png"),
       plot = plot.raw, width = 10, height = 6, dpi = 120)

### Create and save the RAD plot (RAD in mm vs. threshold)
rad_plot <- ggplot(results_df, aes(x = threshold, y = RAD_mm)) +
  geom_line() + geom_point() +
  labs(title = "RAD (mm) vs. Threshold", x = "Threshold", y = "RAD (mm)") +
  theme_minimal()
plot_folder_rad <- JIPipe.AddOutputFolder("Plot_RAD")
ggsave(filename = paste0(plot_folder_rad, "/data.png"),
       plot = rad_plot, width = 6, height = 4, dpi = 120)

### Create and save the FoG plot (FoG vs. threshold)
fog_plot <- ggplot(results_df, aes(x = threshold, y = FoG)) +
  geom_line() + geom_point() +
  labs(title = "FoG vs. Threshold", x = "Threshold", y = "FoG") +
  theme_minimal()
plot_folder_fog <- JIPipe.AddOutputFolder("Plot_FoG")
ggsave(filename = paste0(plot_folder_fog, "/data.png"),
       plot = fog_plot, width = 6, height = 4, dpi = 120)
```

*Node #84 "Remove annotation"*

This node removes the group column annotation (indicator of time point).

- Input "Input" of node #84 receives data from output "Output" of node #83
- The parameter "Removed annotations" (annotation-type) is set to **key == "GroupColumn"**

*Node #85 "Annotate data with table values”*

This note attaches the RAD and FoG values obtained from the R script to the target data.

- Input "Data" of node #85 receives data from output "Labels" of node #62
- Input "Table" of node #85 receives data from output "Output" of node #84
- The parameter item #1 of "Generated annotations" (generated-annotations) is set to **name = "rad_px", value = `LAST_OF(RAD_px)`**
- The parameter item #2 of "Generated annotations" (generated-annotations) is set to **name = "rad_mm", value = `LAST_OF(RAD_mm)`**
- The parameter item #3 of "Generated annotations" (generated-annotations) is set to **name = "FoG", value = `LAST_OF(FoG)`**
- The parameter "Custom grouping columns" in category "Input management" (jipipe:data-batch-generation/custom-matched-columns-expression) is set to `value == "#GroupRow"`

- The parameter "Skip incomplete data sets" in category "Input management" (jipipe:data-batch-generation/skip-incomplete) is set to **true**

***Node #86 "Custom auto threshold 2D (min+max, 32-bit)"***

This node applies a RAD threshold to find the ZOI mask.

- Input "Input" of node #86 receives data from output "Annotated data" of node #85
- The parameter "Thresholding function" in category "Minimum threshold" (min-threshold/thresholding-function) is set to **1**
- The parameter "Thresholding function" in category "Maximum threshold" (max-threshold/thresholding-function) is set to **`NUM(rad_px)`**

***Node #87 "Output" of type "Compartment output"***

This node passes the results to other compartments.

- Input "ZOIMask" of node #87 receives data from output "Output" of node #86
- Input "Statistics" of node #87 receives data from output "Output" of node #84
- Input "Plot_Intensity" of node #87 receives data from output "Plot_Intensity" of node #83
- Input "Plot_RAD" of node #87 receives data from output "Plot_RAD" of node #83
- Input "Plot_FoG" of node #87 receives data from output "Plot_FoG" of node #83
- The output slot name is set to **"Output"**

###### 2.7.1.6 Compartment C6 "Save Visualizations"

This compartment generates, collects, and exports visualizations and plots. A screenshot can be found in **Supplementary Figure 31**.

- The "Save Visualizations" compartment (C6) receives data from the "Raw statistics" compartment (C4)
- The "Save Visualizations" compartment (C6) receives data from the "Statistics" compartment (C5)
- The "Save Visualizations" compartment (C6) receives data from the "Data handling" compartment (C1)

***Node #28 "Output" of type "Compartment output"***

This node passes results from another compartment into the current one.

- Input "SD" of node #28 receives data from output "Image" of node #24
- Input "ZS" of node #28 receives data from output "Image" of node #25
- Input "PL" of node #28 receives data from output "Image" of node #26
- Input "RAW" of node #28 receives data from output "Image" of node #27
- The output slot name is set to **"Output"**

***Node #79 "Output" of type "Compartment output"***

This node passes results from another compartment into the current one.

- Input "RawMeasurements" of node #79 receives data from output "Output" of node #77
- Input "FinalLabels" of node #79 receives data from output "Output" of node #66
- The output slot name is set to **"Output"**

***Node #88 "Export image"***

This node exports an image as a PNG file.

- Input "Input" of node #88 receives data from output "FinalLabels" of node #79
- The parameter "File format" (file-format) is set to **"TIFF"**
- The parameter "File path" (file-path) is set to **`PATH_COMBINE(project_data_dir.tmp_dir, "results", "measurement_labels", #AssayType + "_" + #Experiment + "_" + #Sample + "_" + #TimePoint)`**

***Node #89 "Get label boundaries"***

This node finds the measurement label boundaries for visualization purposes.

- Input "Labels" of node #89 receives data from output "FinalLabels" of node #79

***Node #90 "Fast image arithmetics"***

This node finds the zero-radius label.

- Input "I1" of node #90 receives data from output "FinalLabels" of node #79
- The parameter "Expression" (expression) is set to **I1 == 0**

***Node #91 "Set LUT (two colors)"***

This node sets the LUT of the incoming label boundaries to black-to-cyan.

- Input "Input" of node #91 receives data from output "Boundaries" of node #89
- The parameter "Second color" (second-color) is set to **"#00FFFF"**

***Node #92 "Set LUT (two colors)"***

This node sets the LUT of the incoming greyscale image to black-to-red.

- Input "Input" of node #92 receives data from output "Output" of node #90

***Node #93 "Render to RGB"***

This node renders the greyscale image with LUT into an RGB image.

- Input "Input" of node #93 receives data from output "Output" of node #91

***Node #94 "Render to RGB"***

This node renders the greyscale image with LUT into an RGB image.

- Input "Input" of node #94 receives data from output "Output" of node #92

***Node #95 "Blend images"***

This node creates the final visualization by overlaying multiple images.

- Input "Bottom" of node #95 receives data from output "RAW" of node #28
- Input "Middle" of node #95 receives data from output "Output" of node #94
- Input "Top" of node #95 receives data from output "Output" of node #93
- The parameter "Bottom" in category "Layers" (layers/Bottom) is set to **{"opacity":1.0}**
- The parameter "Middle" in category "Layers" (layers/Middle) is set to **{"opacity":0.4}**
- The parameter "Top" in category "Layers" (layers/Top) is set to **{"opacity":1.0}**

***Node #87 "Output" of type "Compartment output"***

This node passes results from another compartment into the current one.

- Input "ZOIMask" of node #87 receives data from output "Output" of node #86
- Input "Statistics" of node #87 receives data from output "Output" of node #84
- Input "Plot_Intensity" of node #87 receives data from output "Plot_Intensity" of node #83
- Input "Plot_RAD" of node #87 receives data from output "Plot_RAD" of node #83
- Input "Plot_FoG" of node #87 receives data from output "Plot_FoG" of node #83
- The output slot name is set to **"Output"**

###### *Node #96 "Find particles 2D"*
This node gets the ZOI mask as ImageJ ROI.

- Input "Mask" of node #96 receives data from output "ZOIMask" of node #87

###### *Node #97 "Export image"*
This node exports the incoming image as PNG.

- Input "Input" of node #97 receives data from output "Plot_Intensity" of node #87
- The parameter "File path" (file-path) is set to **`PATH_COMBINE(project_data_dir.tmp_dir, "results", "plots-profile", #AssayType + "_" + #Experiment + "_" + #Sample)`**

###### *Node #98 "Export image"*
This node exports the incoming image as PNG.

- Input "Input" of node #98 receives data from output "Plot_RAD" of node #87
- The parameter "File path" (file-path) is set to **`PATH_COMBINE(project_data_dir.tmp_dir, "results", "plots-rad", #AssayType + "_" + #Experiment + "_" + #Sample)`**

###### *Node #99 "Export image"*
This node exports the incoming image as PNG.

- Input "Input" of node #99 receives data from output "Plot_FoG" of node #87
- The parameter "File path" (file-path) is set to **`PATH_COMBINE(project_data_dir.tmp_dir, "results", "plots-fog", #AssayType + "_" + #Experiment + "_" + #Sample)`**

###### *Node #100 "Convert 2D ROI to RGB"*
This node draws an ImageJ ROI over the reference image.

- Input "ROI" of node #100 receives data from output "ROI" of node #96
- Input "Image" of node #100 receives data from output "RAW" of node #28
- The parameter "Override line width" (override-line-width) is set to **4.0**
- The parameter "Override line color" (override-line-color) is set to **"#66FF00"**

###### *Node #101 "Convert 2D ROI to RGB"*
This node draws an ImageJ ROI over the reference image.

- Input "ROI" of node #101 receives data from output "ROI" of node #96
- Input "Image" of node #101 receives data from output "Output" of node #95

- The parameter "Override line width" (override-line-width) is set to **4.0**
- The parameter "Override line color" (override-line-color) is set to **"#66FF00"**

##### *Node #102 "Set physical dimensions from expressions"*

This node sets the physical pixel dimension metadata of the incoming image.

- Input "Input" of node #102 receives data from output "Output" of node #100
- The parameter "Physical dimension (Y)" (physical-dimension-y) is set to **`PixelSize + " mm"`**
- The parameter "Physical dimension (X)" (physical-dimension-x) is set to **`PixelSize + " mm"`**

##### *Node #103 "Set physical dimensions from expressions"*

This node sets the physical pixel dimension metadata of the incoming image.

- Input "Input" of node #103 receives data from output "Output" of node #101
- The parameter "Physical dimension (Y)" (physical-dimension-y) is set to **`PixelSize + " mm"`**
- The parameter "Physical dimension (X)" (physical-dimension-x) is set to **`PixelSize + " mm"`**

##### *Node #104 "Draw scale bar"*

This node draws a scale bar.

- Input "Input" of node #104 receives data from output "Output" of node #102

##### *Node #105 "Draw scale bar"*

This node draws a scale bar.

- Input "Input" of node #105 receives data from output "Output" of node #103

##### *Node #106 "Render overlay"*

This node draws its associated ROI over the incoming image.

- Input "Input" of node #106 receives data from output "Output" of node #104
- The parameter "Draw outline" (draw-outline-mode) is set to **"IfAvailable"**

##### *Node #107 "Render overlay"*

This node draws its associated ROI over the incoming image.

- Input "Input" of node #107 receives data from output "Output" of node #105
- The parameter "Draw outline" (draw-outline-mode) is set to **"IfAvailable"**

##### *Node #108 "Create montage"*

This node creates an image montage.

- Input "Input" of node #108 receives data from output "Output" of node #106
- The parameter "Custom sorting label" in category "Montage" (montage-parameters/sorting-label-expression) is set to **`NUM(GroupColumn)`**

- The parameter "Label" in category "Montage" (montage-parameters/label-expression) is set to **`STRING_FORMAT("%s_%s_%s_%s", #AssayType, #Experiment, #Sample, #TimePoint)`**
- The parameter "Force number of rows" in category "Montage" (montage-parameters/force-rows) is set to **1**
- The parameter "Label font size" in category "Labels" (montage-parameters/label-parameters/label-font-size) is set to **22**
- The parameter "Border size" in category "Canvas" (montage-parameters/canvas-parameters/border-size) is set to **5**
- The parameter "Border color" in category "Canvas" (montage-parameters/canvas-parameters/border-color) is set to **"#000000"**
- The parameter "Grouping method" in category "Merging iteration step generation" (jipipe:data-batch-generation/column-matching) is set to **"Custom"**
- The parameter "Custom grouping columns" in category "Merging iteration step generation" (jipipe:data-batch-generation/custom-matched-columns-expression) is set to **`value IN ARRAY("#GroupRow", "#Threshold")`**

***Node #109 "Create montage"***

This node creates an image montage.

- Input "Input" of node #109 receives data from output "Output" of node #107
- The parameter "Custom sorting label" in category "Montage" (montage-parameters/sorting-label-expression) is set to **`NUM(GroupColumn)`**
- The parameter "Label" in category "Montage" (montage-parameters/label-expression) is set to **`STRING_FORMAT("%s_%s_%s_%s", #AssayType, #Experiment, #Sample, #TimePoint)`**
- The parameter "Force number of rows" in category "Montage" (montage-parameters/force-rows) is set to **1**
- The parameter "Label font size" in category "Labels" (montage-parameters/label-parameters/label-font-size) is set to **22**
- The parameter "Border size" in category "Canvas" (montage-parameters/canvas-parameters/border-size) is set to **5**
- The parameter "Border color" in category "Canvas" (montage-parameters/canvas-parameters/border-color) is set to **"#000000"**
- The parameter "Grouping method" in category "Merging iteration step generation" (jipipe:data-batch-generation/column-matching) is set to **"Custom"**
- The parameter "Custom grouping columns" in category "Merging iteration step generation" (jipipe:data-batch-generation/custom-matched-columns-expression) is set to **`value IN ARRAY("#GroupRow", "#Threshold")`**

***Node #110 "Set annotation (expression)"***

This node sets the correct time point annotation.

- Input "Input" of node #110 receives data from output "Output" of node #108
- The parameter "Annotation name" (annotation-name) is set to **"#TimePoint"**
- The parameter "Annotation value" (annotation-value) is set to **`JOIN_STRING(SORT_ASCENDING(PARSE_JSON(#TimePoint)) , "-")`**

***Node #111 "Set annotation (expression)"***

This node sets the correct time point annotation.

- Input "Input" of node #111 receives data from output "Output" of node #109
- The parameter "Annotation name" (annotation-name) is set to **"#TimePoint"**
- The parameter "Annotation value" (annotation-value) is set to **`JOIN_STRING(SORT_ASCENDING(PARSE_JSON(#TimePoint)) , "-")`**

##### *Node #112 "Draw 2D text ROI"*

This node draws the RAD and FoG over the image.

- Input "Reference" of node #112 receives data from output "Output" of node #110
- The parameter "Background margin" (background-margin) is set to **{"left":{"expression":"2.0"},"top":{"expression":"2.0"},"right":{"expression":"2.0"},"bottom":{"expression":"2.0"}}**
- The parameter "Font size" (font-size) is set to **22**
- The parameter "Location" (location) is set to **{"left":{"expression":"0"},"top":{"expression":"10"},"right":{"expression":"10"},"bottom":{"expression":"0"},"anchor":"TopRight"}**
- The parameter "Text" (text) is set to **`STRING_FORMAT("RAD%d (mm): %s; FoG%d: %s", TO_INTEGER(NUM(#Threshold) * 100), rad_mm, TO_INTEGER(NUM(#Threshold) * 100), D2S(NUM(FoG), 3))`**

##### *Node #113 "Draw 2D text ROI"*

This node draws the RAD and FoG over the image.

- Input "Reference" of node #113 receives data from output "Output" of node #111
- The parameter "Background margin" (background-margin) is set to **{"left":{"expression":"2.0"},"top":{"expression":"2.0"},"right":{"expression":"2.0"},"bottom":{"expression":"2.0"}}**
- The parameter "Font size" (font-size) is set to **22**
- The parameter "Location" (location) is set to **{"left":{"expression":"0"},"top":{"expression":"10"},"right":{"expression":"10"},"bottom":{"expression":"0"},"anchor":"TopRight"}**
- The parameter "Text" (text) is set to **`STRING_FORMAT("RAD%d (mm): %s; FoG%d: %s", TO_INTEGER(NUM(#Threshold) * 100), rad_mm, TO_INTEGER(NUM(#Threshold) * 100), D2S(NUM(FoG), 3))`**

##### *Node #114 "Convert 2D ROI to RGB"*

- Input "ROI" of node #114 receives data from output "ROI" of node #112
- Input "Image" of node #114 receives data from output "Output" of node #110
- The parameter "Merge same annotation values" in category "Input management" (jipipe:data-batch-generation/annotation-merge-strategy) is set to **"OverwriteExisting"**

##### *Node #115 "Convert 2D ROI to RGB"*

This node draws a ROI over the reference image.

- Input "ROI" of node #115 receives data from output "ROI" of node #113
- Input "Image" of node #115 receives data from output "Output" of node #111
- The parameter "Merge same annotation values" in category "Input management" (jipipe:data-batch-generation/annotation-merge-strategy) is set to **"OverwriteExisting"**

***Node #116 "Export image"***

This node exports the incoming image as PNG.

- Input "Input" of node #116 receives data from output "Output" of node #114
- The parameter "File path" (file-path) is set to `PATH_COMBINE(project_data_dir.tmp_dir, "results", "visualization_zoi", #AssayType + "_" + #Experiment + "_" + #Sample + "_" + #TimePoint + "_t" + (NUM(#Threshold) * 100))`

***Node #117 "Export image"***

This node exports the incoming image as PNG.

- Input "Input" of node #117 receives data from output "Output" of node #115
- The parameter "File path" (file-path) is set to `PATH_COMBINE(project_data_dir.tmp_dir, "results", "visualization_all", #AssayType + "_" + #Experiment + "_" + #Sample + "_" + #TimePoint + "_t" + (NUM(#Threshold) * 100))`

###### 2.7.1.7 Compartment C7 "Save tables"

This compartment is responsible for collecting the statistics and exporting them as Excel and CSV tables. A screenshot can be found in **Supplementary Figure 35**.

- The "Save tables" compartment (C7) receives data from the "Raw statistics" compartment (C4)
- The "Save tables" compartment (C7) receives data from the "Statistics" compartment (C5)

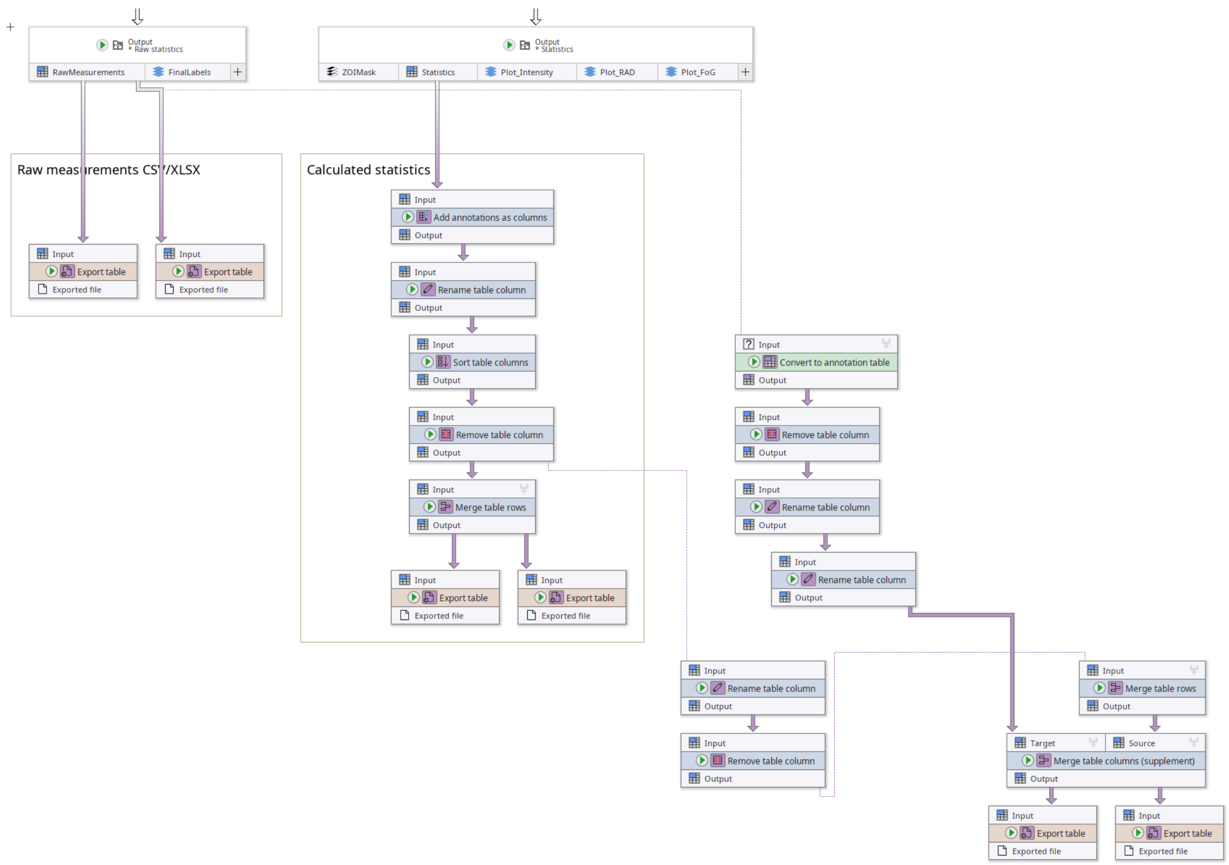

**Supplementary Figure 35:** Contents of the “Save tables” compartment of our single-image DiskImageR implementation in JIPipe.

##### *Node #79 "Output" of type "Compartment output"*

This node passes results from another compartment into the current one.

- Input "RawMeasurements" of node #79 receives data from output "Output" of node #77
- Input "FinalLabels" of node #79 receives data from output "Output" of node #66
- The output slot name is set to **"Output"**

##### *Node #118 "Export table"*

This node exports a table into an Excel file.

- Input "Input" of node #118 receives data from output "RawMeasurements" of node #79
- The parameter "File format" (file-format) is set to **"XLSX"**
- The parameter "File path" (file-path) is set to **`PATH_COMBINE(project_data_dir.tmp_dir, "results", "raw_measurements", #AssayType + "_" + #Experiment + "_" + #Sample + "_" + #TimePoint)`**

##### *Node #119 "Export table"*

This node exports a table into a CSV file.

- Input "Input" of node #119 receives data from output "RawMeasurements" of node #79
- The parameter "File path" (file-path) is set to **`PATH_COMBINE(project_data_dir.tmp_dir, "results",`**

**"raw_measurements", #AssayType + "_" + #Experiment + "_" + #Sample + "_" + #TimePoint)**

*Node #120 "Convert to annotation table"*

- Input "Input" of node #120 receives data from output "RawMeasurements" of node #79
- The parameter "Grouping method" in category "Merging iteration step generation" (jipipe:data-batch-generation/column-matching) is set to **"MergeAll"**

*Node #121 "Remove table column"*

- Input "Input" of node #121 receives data from output "Output" of node #120
- The parameter "Filters" (filters) is set to **`value IN ARRAY("GroupColumn", "#GroupRow", "MinGroupColumn")`**

*Node #122 "Rename table column"*

This node removes the "#" in front of table column names where applicable.

- Input "Input" of node #122 receives data from output "Output" of node #121
- The parameter item #1 of "Renaming entries" (renaming-entries) is set to **source-column = `STRING_STARTS_WITH(value, "#")`, new-name = `SUBSTRING(current_name, 1)`**

*Node #123 "Rename table column"*

This node ensures that a column starting with "ImageId" starts with "#ImageId". This is needed for enhanced J-AST visualization.

- Input "Input" of node #123 receives data from output "Output" of node #122
- The parameter item #1 of "Renaming entries" (renaming-entries) is set to **source-column = `STRING_STARTS_WITH(value, "ImageId")`, new-name = `"#" + current_name`**

*Node #87 "Output" of type "Compartment output"*

This node passes results from another compartment into the current one.

- Input "ZOIMask" of node #87 receives data from output "Output" of node #86
- Input "Statistics" of node #87 receives data from output "Output" of node #84
- Input "Plot_Intensity" of node #87 receives data from output "Plot_Intensity" of node #83
- Input "Plot_RAD" of node #87 receives data from output "Plot_RAD" of node #83
- Input "Plot_FoG" of node #87 receives data from output "Plot_FoG" of node #83
- The output slot name is set to **"Output"**

*Node #124 "Add annotations as columns"*

This node copies annotation into the currently processed table.

- Input "Input" of node #124 receives data from output "Statistics" of node #87
- The parameter "Annotation name filter" (annotation-name-filter) is set to **`value IN ARRAY("#Experiment", "#AssayType", "#Threshold", "#Sample", "PixelSize")`**

*Node #125 "Rename table column"*

This node removes the “#” in front of table column names where applicable.

- Input "Input" of node #125 receives data from output "Output" of node #124
- The parameter item #1 of "Renaming entries" (renaming-entries) is set to **source-column = `STRING_STARTS_WITH(value, "#")`, new-name = `SUBSTRING(current_name, 1)`**

###### *Node #126 "Sort table columns"*

This node ensures that the table columns follow a specific order.

- Input "Input" of node #126 receives data from output "Output" of node #125
- The parameter item #1 of "Manual order" (manual-order) is set to **"AssayType"**
- The parameter item #2 of "Manual order" (manual-order) is set to **"Experiment"**
- The parameter item #3 of "Manual order" (manual-order) is set to **"Sample"**
- The parameter item #4 of "Manual order" (manual-order) is set to **"PixelSize"**
- The parameter item #5 of "Manual order" (manual-order) is set to **"Threshold"**

###### *Node #127 "Remove table column"*

This node removes the “PixelSize” table column.

- Input "Input" of node #127 receives data from output "Output" of node #126
- The parameter "Filters" (filters) is set to **`value IN ARRAY("t", "PixelSize")`**

###### *Node #128 "Merge table rows"*

This node creates a large all-in-one table.

- Input "Input" of node #128 receives data from output "Output" of node #127
- The parameter "Grouping method" in category "Merging iteration step generation" (jipipe:data-batch-generation/column-matching) is set to **"MergeAll"**

###### *Node #129 "Rename table column"*

This node renames table columns to the desired final name.

- Input "Input" of node #129 receives data from output "Output" of node #127
- The parameter item #1 of "Renaming entries" (renaming-entries) is set to **source-column = `value IN ARRAY("AUC", "FoG")`, new-name = `current_name + TO_INTEGER(NUM(#Threshold) * 100)`**
- The parameter item #2 of "Renaming entries" (renaming-entries) is set to **source-column = `"RAD_mm"`, new-name = `"RAD" + TO_INTEGER(NUM(#Threshold) * 100) + "_mm"`**
- The parameter item #3 of "Renaming entries" (renaming-entries) is set to **source-column = `"RAD_px"`, new-name = `"RAD" + TO_INTEGER(NUM(#Threshold) * 100) + "_px"`**

###### *Node #130 "Export table"*

This node exports the incoming table as a CSV file.

- Input "Input" of node #130 receives data from output "Output" of node #128
- The parameter "File path" (file-path) is set to **`PATH_COMBINE(project_data_dir.tmp_dir, "results", "statistics")`**

###### *Node #131 "Export table"*

- Input "Input" of node #131 receives data from output "Output" of node #128
- The parameter "File format" (file-format) is set to **"XLSX"**
- The parameter "File path" (file-path) is set to **`PATH_COMBINE(project_data_dir.tmp_dir, "results", "statistics")`**

###### *Node #132 "Remove table column"*

This node removes all remaining internal thresholding information from the results.

- Input "Input" of node #132 receives data from output "Output" of node #129
- The parameter "Filters" (filters) is set to **`value IN ARRAY("Threshold", "threshold")`**

###### *Node #133 "Merge table rows"*

This node creates a large all-in-one table per threshold.

- Input "Input" of node #133 receives data from output "Output" of node #132
- The parameter "Grouping method" in category "Merging iteration step generation" (jipipe:data-batch-generation/column-matching) is set to **"Custom"**
- The parameter "Custom grouping columns" in category "Merging iteration step generation" (jipipe:data-batch-generation/custom-matched-columns-expression) is set to **"#Threshold"**

###### *Node #134 "Merge table columns (supplement)"*

This node adds columns from one table into another. The algorithm uses specific reference columns to align the data.

- Input "Target" of node #134 receives data from output "Output" of node #123
- Input "Source" of node #134 receives data from output "Output" of node #133
- The parameter "Column filter" (column-filter) is set to **`value != "PixelSize"`**
- The parameter "Reference columns" (reference-columns) is set to **`value IN ARRAY("Experiment", "AssayType", "Sample")`**
- The parameter "Grouping method" in category "Merging iteration step generation" (jipipe:data-batch-generation/column-matching) is set to **"MergeAll"**

###### *Node #135 "Export table"*

This node exports the incoming table as a CSV file.

- Input "Input" of node #135 receives data from output "Output" of node #134
- The parameter "File path" (file-path) is set to **`PATH_COMBINE(project_data_dir.tmp_dir, "results", "statistics_aio")`**

###### *Node #136 "Export table"*

This node exports the incoming table as an Excel file.

- Input "Input" of node #136 receives data from output "Output" of node #134
- The parameter "File format" (file-format) is set to **"XLSX"**

- The parameter "File path" (file-path) is set to **`PATH_COMBINE(project_data_dir.tmp_dir, "results", "statistics_aio")`**

### 3 MIC detection algorithm

The *J-AST* MIC detector algorithm (**Fig. 6a**) begins with basic corrective operations, including the normalization of the image size to ensure a resolution of 600 DPI, as well as a set of operations that rotate the image so that the strip is vertical. The operations are required for a successful application of an optical character recognition (OCR) using *Tesseract* (https://github.com/tesseract-ocr/tesseract) on the strip. Due to the variety in imaging conditions multiple background subtraction parameters in a configurable range from 20 to 64 pixels with a step size of four are tested. For each candidate the OCR yields readouts of detected text together with their location, which is filtered by a confidence threshold of 90%. The best candidate is chosen by applying a Smith-Waterman alignment algorithm (match score 1, gap score 0, mismatch score -9999) where the detected sequence is aligned against the user-provided reference sequence of concentrations shown on the strip. The high mismatch and low gap score ensure that the algorithm can handle long gaps of missing detections without starting to misalign correct results. If at least two points are available, the *J-AST* Etest pipeline then is able to reconstruct the per-concentration pixel locations which allows it to segment the image into regions assigned to each individual value. Additional points are used to improve the estimation of the tick spacing. The resulting vertical segmentation is combined with a horizontal partitioning into cells (**Fig. 6b**) that have a width of 0.3 mm. The area captured by cells extends up to 30 mm from the strip edge for performance reasons. Each cell is associated with a concentration value and the mean intensity of all marked pixels. *J-AST* then extracts the lawn intensity by calculating the median intensity of the outer-most cells of the three lowest concentrations. The ZOI background intensity is determined by calculating the fifth percentile of all cell intensity values. The next step involves the rough estimation of the ZOI boundary distance per concentration. This is done by finding the minimum distance from the strip edge where a 30% increase in intensity compared to the background is detected. To filter out artifacts, distances below 0.5 mm are excluded and it is ensured that the boundary distance monotonically increases with concentration. The maximum of all boundary distances is used as a classifier between narrow and wide ZOIs where if the value is higher than 7.5 mm, the ZOI shape is assumed to be wide. For narrow ZOI images a corridor $d$ of 0.3 mm is used for the concentration $c$ to intensity mapping $I(c, d)$, where the size for wide ZOIs is set to 3 mm. This is required as with narrow ZOIs the corridor needs to capture the sharp inhibition zone, while for wider ZOIs the algorithm can benefit from the increased robustness to handle sparse colonies. We normalize the intensities by applying $I_{norm}(c, d) = \frac{I(c,d)-I_{bgr}(d)}{I_{max}(d)-I_{bgr}(d)}$ where the background $I_{bgr}(d)$ is calculated as mean intensity of the by default three highest concentrations within the corridor while the maximum growth $I_{max}(d)$ is the mean of the three lowest concentration intensities. The MIC in the standard configuration is then detected as the highest concentration where $I_{norm}(d)$ is at least 25.5%. The resulting concentration value is then rounded to the closest tick in log space. Our optional detection for assays without a valid MIC then applies a set of three rules: The first rule calculates a contrast $C(d) = \frac{I_{max}(d)-I_{bgr}(d)}{(I_{max}(d)+I_{bgr}(d))/2}$ for both the narrow (0.3 mm) and wide (3 mm) corridors and checks if both are over 44%, thus excluding images without a ZOI as only lawn intensities are measured. The second rule checks if the maximum ZOI boundary distance is above 1.5 mm, which removes cases where no proper ZOI boundary can be found. The third rule calculates the coefficient of variation $CV$ of the intensities associated to the by default three highest concentrations, where a $CV > 15\%$ is an indicator of growth inside the ZOI. If any of the three rules apply the algorithm returns with a MIC of N/A.

The following steps are applied by our MIC detection algorithm. An overview can be found in **Fig 6**. The structure is explained in **Supplementary Figure 36**.

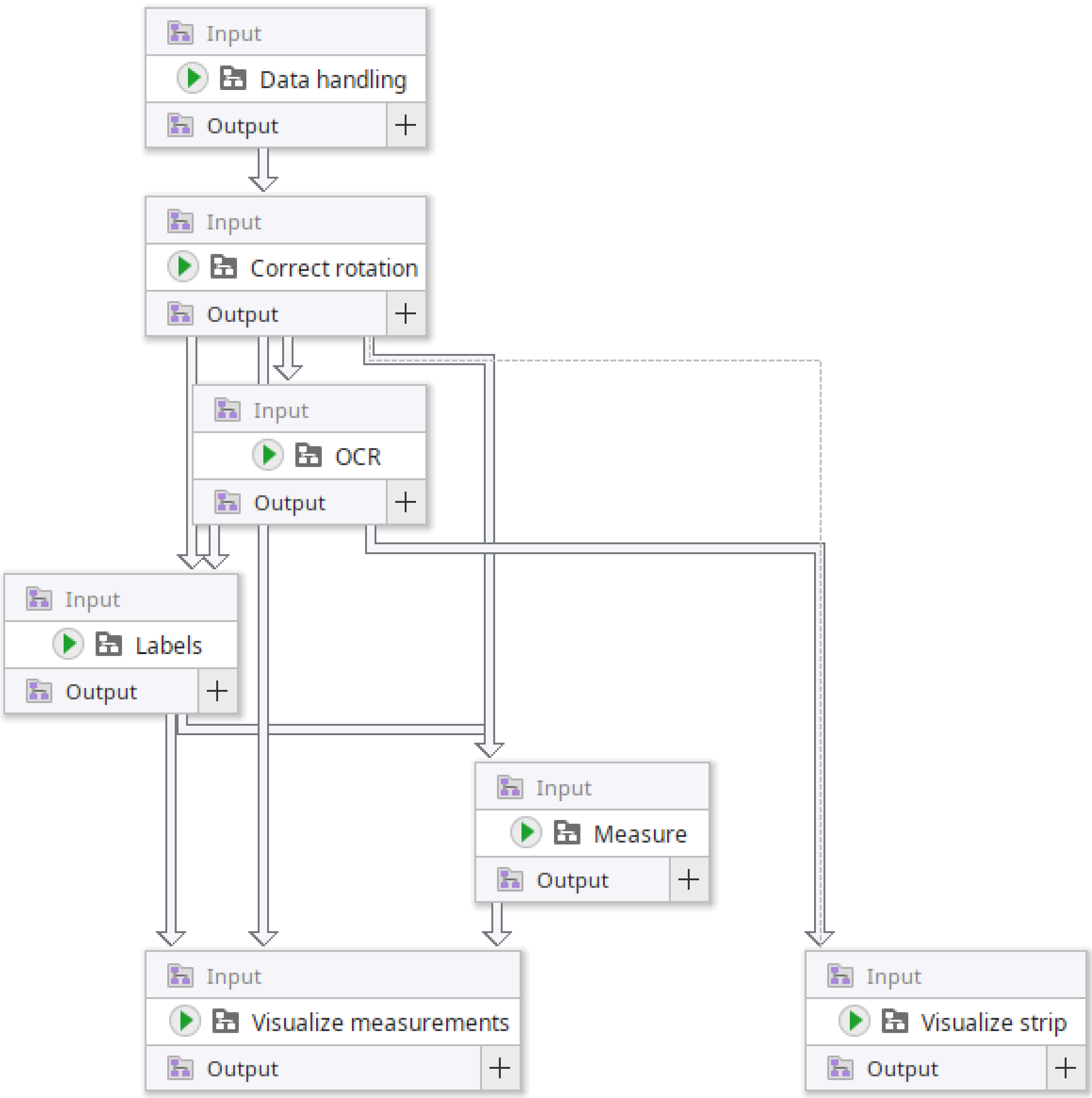

**Supplementary Figure 36:** Compartment structure of the MIC detection algorithm.

#### Project-wide parameters

The following parameters are attached to the project:

| Key | Name | Default value |
|---|---|---|
| targetDPI | Target DPI | 600 |
| confidenceThreshold | Confidence threshold | 90.0 |

| | | |
|---|---|---|
| stripSafetyDistanceMillimeters | Safety distance from strip border (mm) | 0.5 |
| stripMeasurementDistanceMillimeters | Measure from strip border (mm) | 5.0 |
| handleInvertedImages | Handle inverted images | false |
| bgrSubtractionStartRadius | Background subtraction start radius | 20 |
| bgrSubtractionEndRadius | Background subtraction end radius | 64 |
| bgrSubtractionRadiusIncrement | Background subtraction radius increment | 4 |

#### Compartment C1 “Data handling”

This compartment is responsible for collecting and organizing the input data. A screenshot can be found in **Supplementary Figure 37**.

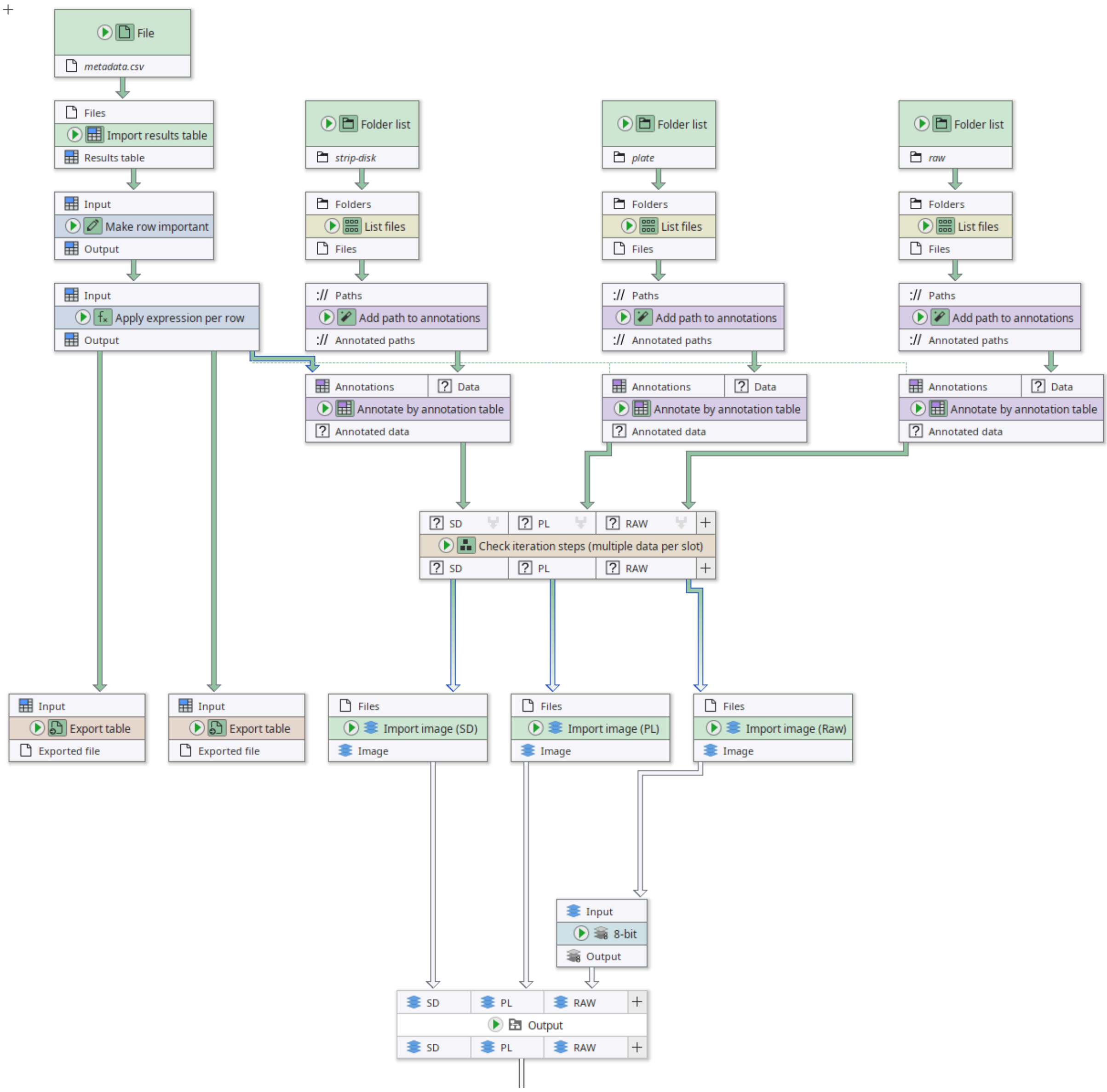

**Supplementary Figure 37:** “Data handling” compartment of the MIC analysis pipeline.

##### *Node #1 "Folder list"*

- The parameter item #1 of "Folder paths" (folder-paths) is set to **"strip-disk"**

##### *Node #2 "File"*

- The parameter "File name" (file-name) is set to **"metadata.csv"**

##### *Node #3 "Folder list"*

- The parameter item #1 of "Folder paths" (folder-paths) is set to **"plate"**

##### *Node #4 "Folder list"*

- The parameter item #1 of "Folder paths" (folder-paths) is set to **"raw"**

##### *Node #5 "List files"*

- Input "Folders" of node #5 receives data from output "Folder paths" of node #1
- The parameter "Keep file if ..." (filters) is set to **STRING_MATCHES_GLOB(name, "*.png")**

*Node #6 "Import results table"*

- Input "Files" of node #6 receives data from output "Filename" of node #2

*Node #7 "List files"*

- Input "Folders" of node #7 receives data from output "Folder paths" of node #3
- The parameter "Keep file if ..." (filters) is set to **STRING_MATCHES_GLOB(name, "*.png")**

*Node #8 "List files"*

- Input "Folders" of node #8 receives data from output "Folder paths" of node #4
- The parameter "Keep file if ..." (filters) is set to **STRING_MATCHES_GLOB(name, "*.png")**

*Node #9 "Add path to annotations"*

- Input "Paths" of node #9 receives data from output "Files" of node #5
- The parameter "Generated annotation" (generated-annotation) is set to **"#ImageId"**

*Node #10 "Make row important"*

- Input "Input" of node #10 receives data from output "Results table" of node #6
- The parameter item #1 of "Renaming entries" (renaming-entries) is set to **source-column = "GroupRow", new-name = "#GroupRow"**

*Node #11 "Add path to annotations"*

- Input "Paths" of node #11 receives data from output "Files" of node #7
- The parameter "Generated annotation" (generated-annotation) is set to **"#ImageId"**

*Node #12 "Add path to annotations"*

- Input "Paths" of node #12 receives data from output "Files" of node #8
- The parameter "Generated annotation" (generated-annotation) is set to **"#ImageId"**

*Node #13 "Apply expression per row"*

- Input "Input" of node #13 receives data from output "Output" of node #10
- The parameter item #1 of "Generated values" (entries) is set to **column-name = "MinGroupColumn", value = MIN(all.GroupColumn)**

*Node #14 "Annotate by annotation table"*

- Input "Annotations" of node #14 receives data from output "Output" of node #13
- Input "Data" of node #14 receives data from output "Annotated paths" of node #9

*Node #15 "Annotate by annotation table"*

- Input "Annotations" of node #15 receives data from output "Output" of node #13
- Input "Data" of node #15 receives data from output "Annotated paths" of node #12

*Node #16 "Export table"*

- Input "Input" of node #16 receives data from output "Output" of node #13
- The parameter "File path" (file-path) is set to **PATH_COMBINE(project_data_dir.tmp_dir, "results", "metadata")**

***Node #17 "Annotate by annotation table"***

- Input "Annotations" of node #17 receives data from output "Output" of node #13
- Input "Data" of node #17 receives data from output "Annotated paths" of node #11

***Node #18 "Export table"***

- Input "Input" of node #18 receives data from output "Output" of node #13
- The parameter "File format" (file-format) is set to **"XLSX"**
- The parameter "File path" (file-path) is set to **PATH_COMBINE(project_data_dir.tmp_dir, "results", "metadata")**

***Node #19 "Check iteration steps (multiple data per slot)"***

- Input "SD" of node #19 receives data from output "Annotated data" of node #14
- Input "PL" of node #19 receives data from output "Annotated data" of node #17
- Input "RAW" of node #19 receives data from output "Annotated data" of node #15
- The parameter "Custom grouping columns" in category "Merging iteration step generation" (jipipe:data-batch-generation/custom-matched-columns-expression) is set to **value == "#GroupRow"**

***Node #20 "Import image (SD)" of type "Import image"***

- Input "Files" of node #20 receives data from output "SD" of node #19

***Node #21 "Import image (PL)" of type "Import image"***

- Input "Files" of node #21 receives data from output "PL" of node #19

***Node #22 "Import image (Raw)" of type "Import image"***

- Input "Files" of node #22 receives data from output "RAW" of node #19
- The parameter "Remove LUT" (remove-lut) is set to **true**

***Node #23 "8-bit" of type "Convert image to 8-bit"***

- Input "Input" of node #23 receives data from output "Image" of node #22

***Node #24 "Output"***

- Input "SD" of node #24 receives data from output "Image" of node #20
- Input "PL" of node #24 receives data from output "Image" of node #21
- Input "RAW" of node #24 receives data from output "Output" of node #23
- The parameter "jipipe:compartment:output-slot-name" (jipipe:compartment:output-slot-name) is set to **"Output"**

#### Compartment C2 “Correct rotation”

This compartment applies DPI-aware upscaling to the input images and rotates the image so that the strip is perfectly vertical. A screenshot can be found in **Supplementary Figure 38**.

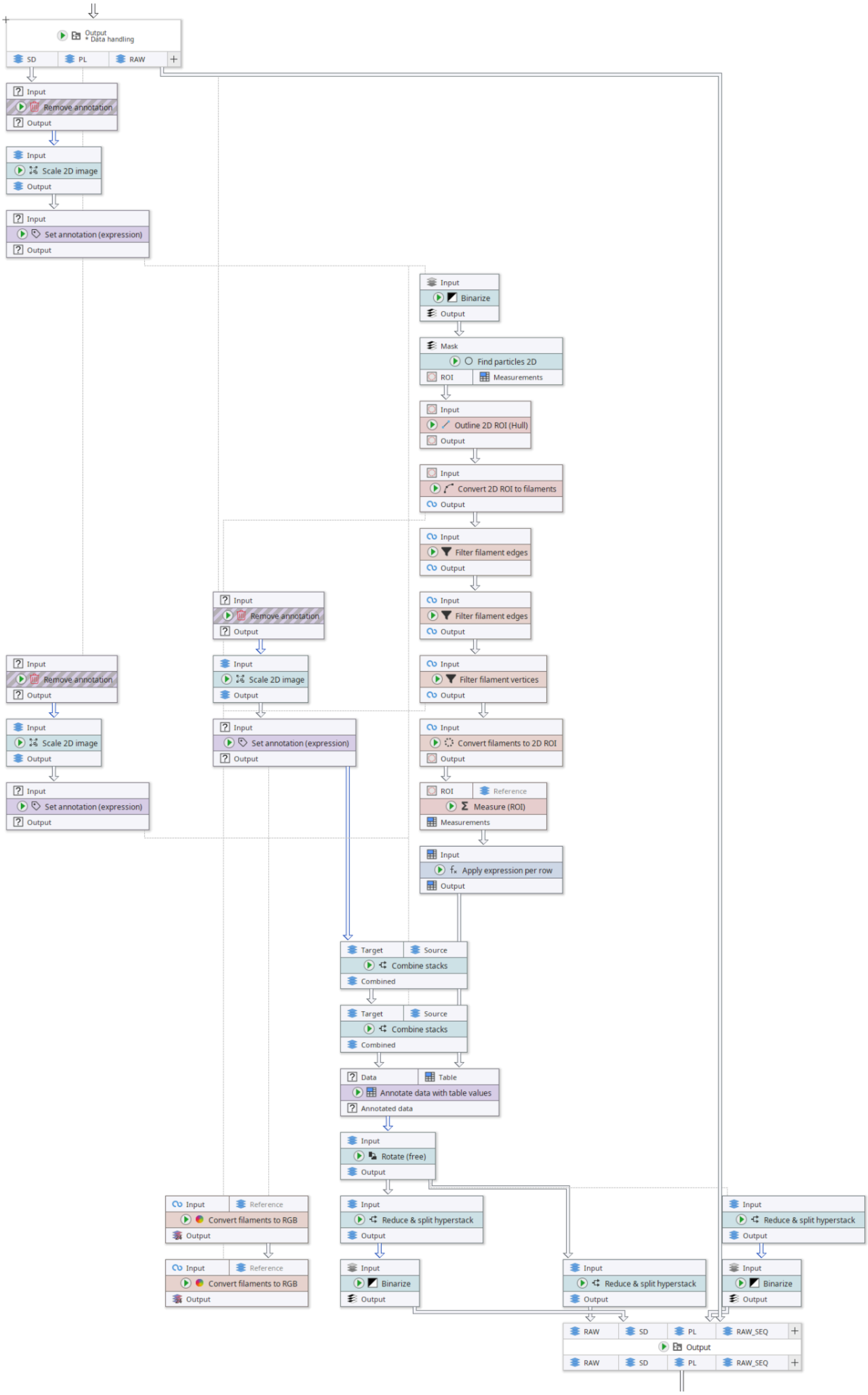

**Supplementary Figure 38:** “Correct rotation” compartment of the MIC analysis pipeline.

*Node #24 "Output" of type "Compartment output"*

- Input "SD" of node #24 receives data from output "Image" of node #20
- Input "PL" of node #24 receives data from output "Image" of node #21
- Input "RAW" of node #24 receives data from output "Output" of node #23
- The parameter "jipipe:compartment:output-slot-name" (jipipe:compartment:output-slot-name) is set to **"Output"**

*Node #25 "Remove annotation"*

- Input "Input" of node #25 receives data from output "SD" of node #24
- The parameter "Pass through" (jipipe:algorithm:pass-through) is set to **true**
- The parameter "Removed annotations" (annotation-type) is set to **key == "StripSequence"**

*Node #26 "Remove annotation"*

- Input "Input" of node #26 receives data from output "PL" of node #24
- The parameter "Pass through" (jipipe:algorithm:pass-through) is set to **true**
- The parameter "Removed annotations" (annotation-type) is set to **key == "StripSequence"**

*Node #27 "Remove annotation"*

- Input "Input" of node #27 receives data from output "RAW" of node #24
- The parameter "Pass through" (jipipe:algorithm:pass-through) is set to **true**
- The parameter "Removed annotations" (annotation-type) is set to **key == "StripSequence"**

*Node #28 "Scale 2D image"*

- Input "Input" of node #28 receives data from output "Output" of node #25
- The parameter "Y axis" (y-axis) is set to **height * ( _global.targetDPI / (25.4 / PixelSize))**
- The parameter "X axis" (x-axis) is set to **width * ( _global.targetDPI / (25.4 / PixelSize))**

*Node #29 "Scale 2D image"*

- Input "Input" of node #29 receives data from output "Output" of node #26
- The parameter "Y axis" (y-axis) is set to **height * ( _global.targetDPI / (25.4 / PixelSize))**
- The parameter "X axis" (x-axis) is set to **width * ( _global.targetDPI / (25.4 / PixelSize))**

*Node #30 "Scale 2D image"*

- Input "Input" of node #30 receives data from output "Output" of node #27
- The parameter "Y axis" (y-axis) is set to **height * ( _global.targetDPI / (25.4 / PixelSize))**
- The parameter "X axis" (x-axis) is set to **width * ( _global.targetDPI / (25.4 / PixelSize))**

*Node #31 "Set annotation (expression)"*

- Input "Input" of node #31 receives data from output "Output" of node #28
- The parameter "Annotation name" (annotation-name) is set to **"PixelSizeRescaled"**
- The parameter "Annotation value" (annotation-value) is set to **1 / ( _global.targetDPI * 1 / 25.4)**

*Node #32 "Set annotation (expression)"*

- Input "Input" of node #32 receives data from output "Output" of node #29
- The parameter "Annotation name" (annotation-name) is set to **"PixelSizeRescaled"**
- The parameter "Annotation value" (annotation-value) is set to **1 / ( _global.targetDPI * 1 / 25.4)**

***Node #33 "Set annotation (expression)"***

- Input "Input" of node #33 receives data from output "Output" of node #30
- The parameter "Annotation name" (annotation-name) is set to **"PixelSizeRescaled"**
- The parameter "Annotation value" (annotation-value) is set to **1 / ( _global.targetDPI * 1 / 25.4)**

***Node #34 "Binarize"***

- Input "Input" of node #34 receives data from output "Output" of node #31

***Node #35 "Combine stacks"***

- Input "Target" of node #35 receives data from output "Output" of node #33
- Input "Source" of node #35 receives data from output "Output" of node #31

***Node #36 "Find particles 2D"***

- Input "Mask" of node #36 receives data from output "Output" of node #34

***Node #37 "Combine stacks"***

- Input "Target" of node #37 receives data from output "Combined" of node #35
- Input "Source" of node #37 receives data from output "Output" of node #32

***Node #38 "Outline 2D ROI (Hull)"***

- Input "Input" of node #38 receives data from output "ROI" of node #36
- The parameter "Outline method" (outline) is set to **"MinimumBoundingRectangle"**

***Node #39 "Convert 2D ROI to filaments"***

- Input "Input" of node #39 receives data from output "Output" of node #38

***Node #40 "Convert filaments to RGB"***

- Input "Input" of node #40 receives data from output "Output" of node #39
- Input "Reference" of node #40 receives data from output "Output" of node #33
- The parameter "Override edge thickness" in category "Filament drawing settings" (filaments-drawer/override-edge-thickness) is set to **5**

***Node #41 "Filter filament edges"***

- Input "Input" of node #41 receives data from output "Output" of node #39
- The parameter "Only keep edge if" (filter) is set to **length > AVG(all.length)**

***Node #42 "Filter filament edges"***

- Input "Input" of node #42 receives data from output "Output" of node #41
- The parameter "Only keep edge if" (filter) is set to **uuid == FIRST_OF(all.uuid)**

***Node #43 "Filter filament vertices"***

- Input "Input" of node #43 receives data from output "Output" of node #42
- The parameter "Only keep vertex if" (filter) is set to **degree > 0**

*Node #44 "Convert filaments to RGB"*

- Input "Input" of node #44 receives data from output "Output" of node #43
- Input "Reference" of node #44 receives data from output "Output" of node #40
- The parameter "Override edge color" in category "Filament drawing settings" (filaments-drawer/override-edge-color) is set to **"#00FF00"**
- The parameter "Override edge thickness" in category "Filament drawing settings" (filaments-drawer/override-edge-thickness) is set to **5**

*Node #45 "Convert filaments to 2D ROI"*

- Input "Input" of node #45 receives data from output "Output" of node #43
- The parameter "With vertices" (with-vertices) is set to **false**

*Node #46 "Measure (ROI)" of type "Extract 2D ROI statistics"*

- Input "ROI" of node #46 receives data from output "Output" of node #45
- The parameter "Extracted measurements" (measurements) is set to **{"values":["FeretDiameter"],"collapsed":false}**

*Node #47 "Apply expression per row"*

- Input "Input" of node #47 receives data from output "Measurements" of node #46
- The parameter item #1 of "Generated values" (entries) is set to **column-name = "Angle2", value = IF_ELSE(Angle < 0, Angle + 180, Angle)**
- The parameter item #2 of "Generated values" (entries) is set to **column-name = "rotation", value = Angle2 - 90**

*Node #48 "Annotate data with table values"*

- Input "Data" of node #48 receives data from output "Combined" of node #37
- Input "Table" of node #48 receives data from output "Output" of node #47
- The parameter item #1 of "Generated annotations" (generated-annotations) is set to **name = "rotation_angle", value = FIRST_OF(rotation)**

*Node #49 "Rotate (free)" of type "Rotate 2D image (free)"*

- Input "Input" of node #49 receives data from output "Annotated data" of node #48
- The parameter "Angle (in degrees)" (angle) is set to **45.0**
- The parameter item #1 of "Overridden parameters" in category "Adaptive parameters" (jipipe:adaptive-parameters/overridden-parameters) is set to **(NUM(rotation_angle), "angle")**

*Node #50 "Reduce & split hyperstack"*

- Input "Input" of node #50 receives data from output "Output" of node #49

*Node #51 "Reduce & split hyperstack"*

- Input "Input" of node #51 receives data from output "Output" of node #49
- The parameter "Indices (Z)" (indices-z) is set to **2**

*Node #52 "Reduce & split hyperstack"*

- Input "Input" of node #52 receives data from output "Output" of node #49

- The parameter "Indices (Z)" (indices-z) is set to **1**

*Node #53 "Binarize"*

- Input "Input" of node #53 receives data from output "Output" of node #51

*Node #54 "Binarize"*

- Input "Input" of node #54 receives data from output "Output" of node #52

*Node #55 "Output" of type "Compartment output"*

- Input "RAW" of node #55 receives data from output "Output" of node #50
- Input "SD" of node #55 receives data from output "Output" of node #54
- Input "PL" of node #55 receives data from output "Output" of node #53
- Input "RAW_SEQ" of node #55 receives data from output "RAW" of node #24
- The parameter "jipipe:compartment:output-slot-name" (jipipe:compartment:output-slot-name) is set to **"Output"**

#### Compartment C3 “OCR”

This compartment is responsible for optical character recognition (OCR) and segmenting the image into per-concentration measurement labels. A screenshot can be seen in **Supplementary Figure 39**.

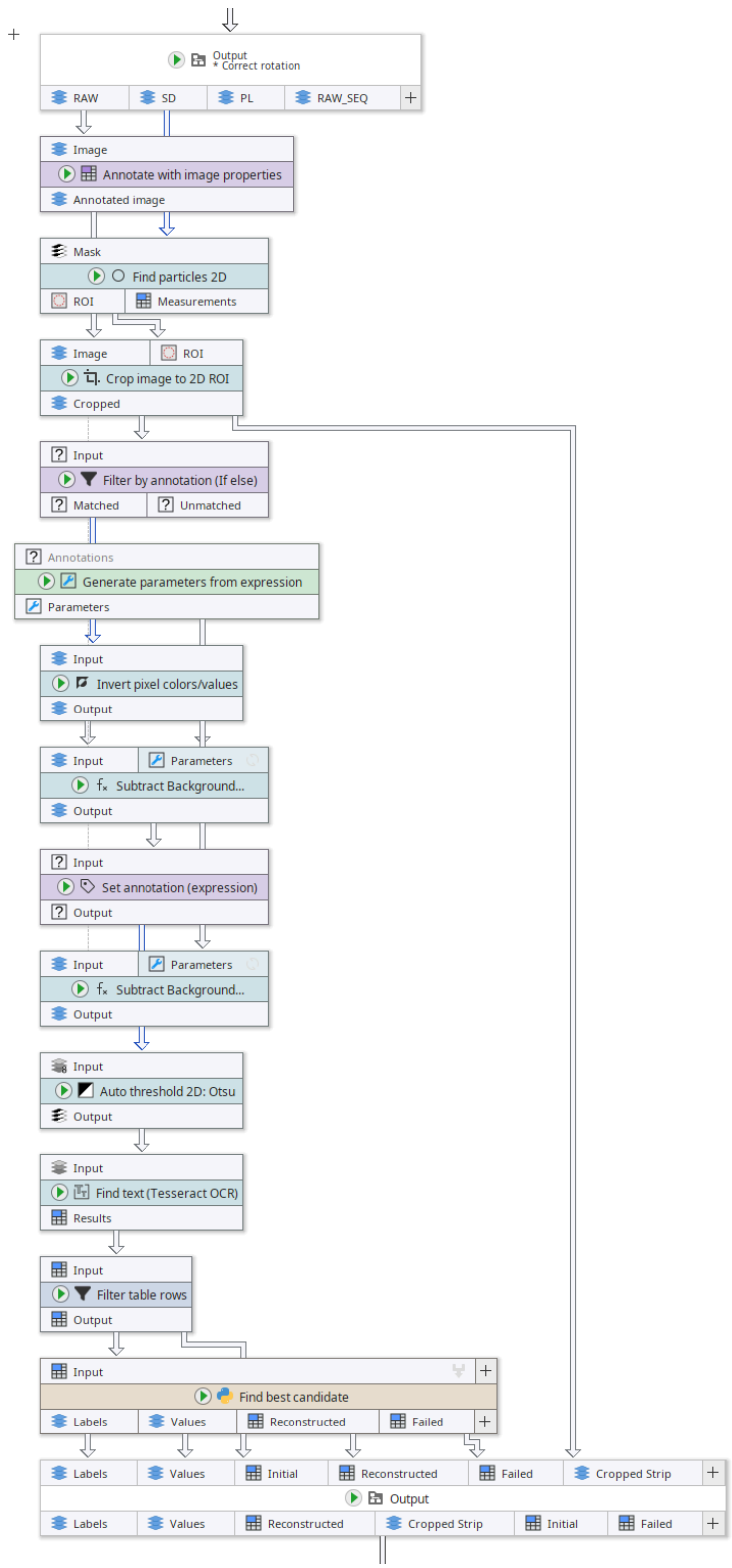

**Supplementary Figure 39:** “OCR” compartment of the MIC analysis pipeline.

##### *Node #56 "Generate parameters from expression" of type "Generate parameters from expression"*

- The parameter item #1 of "Generated parameter columns" (columns) is set to **values = MAKE_SEQUENCE(_global.bgrSubtractionStartRadius, _global.bgrSubtractionEndRadius + 1, _global.bgrSubtractionRadiusIncrement), type = "int", values-are-json = false, key = "radius"**

##### *Node #55 "Output" of type "Compartment output"*

- Input "RAW" of node #55 receives data from output "Output" of node #50
- Input "SD" of node #55 receives data from output "Output" of node #54
- Input "PL" of node #55 receives data from output "Output" of node #53
- Input "RAW_SEQ" of node #55 receives data from output "RAW" of node #24
- The parameter "jipipe:compartment:output-slot-name" (jipipe:compartment:output-slot-name) is set to **"Output"**

***Node #57 "Annotate with image properties" of type "Annotate with image properties"***

- Input "Image" of node #57 receives data from output "RAW" of node #55
- The parameter "Annotate with image height" (height-annotation) is set to **"ih"**
- The parameter "Annotate with image width" (width-annotation) is set to **"iw"**

***Node #58 "Find particles 2D" of type "Find particles 2D"***

- Input "Mask" of node #58 receives data from output "SD" of node #55

***Node #59 "Crop image to 2D ROI" of type "Crop image to 2D ROI"***

- Input "Image" of node #59 receives data from output "Annotated image" of node #57
- Input "ROI" of node #59 receives data from output "ROI" of node #58
- The parameter "Crop channel plane" (crop-c) is set to **false**
- The parameter "Crop Z plane" (crop-z) is set to **false**
- The parameter "Annotate with Y location" (annotation-y) is set to **"cy"**
- The parameter "Annotate with X location" (annotation-x) is set to **"cx"**
- The parameter "Crop frame plane" (crop-t) is set to **false**

***Node #60 "Subtract Background..." of type "Find or subtract background 2D"***

- Input "Input" of node #60 receives data from output "Cropped" of node #59
- Input "{Parameters}" of node #60 receives data from output "Parameters" of node #56
- The parameter "Multiple parameters" in category "Multi-parameter settings" (jipipe:parameter-slot-algorithm/has-parameter-slot) is set to **true**

***Node #61 "Filter by annotation (If else)" of type "Filter by annotation (If else)"***

- Input "Input" of node #61 receives data from output "Cropped" of node #59
- The parameter "Only match data if" (filter) is set to **_global.handleInvertedImages**

***Node #62 "Invert pixel colors/values" of type "Invert pixel colors/values"***

- Input "Input" of node #62 receives data from output "Matched" of node #61

***Node #63 "Subtract Background..." of type "Find or subtract background 2D"***

- Input "Input" of node #63 receives data from output "Output" of node #62
- Input "{Parameters}" of node #63 receives data from output "Parameters" of node #56
- The parameter "Multiple parameters" in category "Multi-parameter settings" (jipipe:parameter-slot-algorithm/has-parameter-slot) is set to **true**

***Node #64 "Set annotation (expression)" of type "Set annotation (expression)"***

- Input "Input" of node #64 receives data from output "Output" of node #63
- The parameter "Annotation name" (annotation-name) is set to **"Radius"**
- The parameter "Annotation value" (annotation-value) is set to **Radius + "_inv"**

***Node #65 "Auto threshold 2D: Otsu" of type "Auto threshold 2D"***

- Input "Input" of node #65 receives data from output "Output" of node #64
- Input "Input" of node #65 receives data from output "Output" of node #60
- The parameter "Method" (method) is set to **"Otsu"**

*Node #66 "Find text (Tesseract OCR)" of type "Find text (Tesseract OCR)"*

- Input "Input" of node #66 receives data from output "Output" of node #65
- The parameter "Specify recognized characters" (override-char-allow-list) is set to **"0123456789-.,"**
- The parameter "Override Tesseract OCR environment" (override-environment) is set to **[Disabled]**
- The parameter "Page segmentation method" (page-segmentation-method) is set to **"PSM6"**

*Node #67 "Filter table rows" of type "Filter table rows"*

- Input "Input" of node #67 receives data from output "Results" of node #66
- The parameter "Filters" (filters) is set to **conf >= _global.confidenceThreshold**

*Node #68 "Find best candidate" of type "Jython script (merging)"*

- Input "Input" of node #68 receives data from output "Output" of node #67

The script evaluates the incoming candidates and aligns them against the known concentration sequence using a Smith-Waterman alignment. The candidate with the highest score and with at least two sequence items is selected. Then the value mapping and measurement labels are generated directly within the script.

The following code is contained within the script:

```
from org.hkijena.jipipe.plugins.tables.datatypes import ResultsTableData
from org.hkijena.jipipe.plugins.imagejdatatypes.datatypes import
ImagePlusData
from ij.process import FloatProcessor
from ij import ImagePlus
import json

def mean(x):
    return 1.0 * sum(x) / len(x)

def num(x):
    x = x.replace(",", ".")
    try:
        return float(x)
    except:
        return None
    return None

def smith_waterman_current_match_indices(reference, current,
                                         match_score=1,
                                         mismatch_penalty=-9999,
                                         gap_penalty=0,
                                         eps=1e-6):
```

```
    """
    Local alignment (Smith–Waterman) between `reference` and `current`.
    Returns indices in `current` that are matched (diagonal steps) in the
best local alignment.

    Scoring:
      match: +match_score
      mismatch: +mismatch_penalty (not allowed - very negative value)
      gap (in either seq): +gap_penalty  (negative)
    """
    n = len(reference)
    m = len(current)

    # H: score matrix; P: traceback pointers
    # P values: 0=stop, 1=diag, 2=up, 3=left
    H = [[0] * (m + 1) for _ in range(n + 1)]
    P = [[0] * (m + 1) for _ in range(n + 1)]

    def is_match(a, b):
        if b is None or a is None:
            return False
        try:
            return abs(float(a) - float(b)) <= eps
        except:
            return False

    best_score = 0
    best_i = 0
    best_j = 0

    for i in range(1, n + 1):
        ra = reference[i - 1]
        for j in range(1, m + 1):
            cb = current[j - 1]

            diag = H[i - 1][j - 1] + \
                (match_score if is_match(ra, cb) else mismatch_penalty)
            up = H[i - 1][j] + gap_penalty      # gap in current
            left = H[i][j - 1] + gap_penalty      # gap in reference

            # local alignment => scores can't go below 0
            score = 0
            ptr = 0
            if diag > score:
                score = diag
                ptr = 1
            if up > score:
                score = up
                ptr = 2
            if left > score:
                score = left
                ptr = 3

            H[i][j] = score
            P[i][j] = ptr
```

```
            if score > best_score:
                best_score = score
                best_i = i
                best_j = j

    # Traceback from best cell until stop (0)
    i, j = best_i, best_j
    matched_current_indices = []
    matched_ref_indices = []

    while i > 0 and j > 0 and P[i][j] != 0:
        ptr = P[i][j]
        if ptr == 1:  # diag
            # only record if it was an actual match (not a mismatch)
            if is_match(reference[i - 1], current[j - 1]):
                matched_current_indices.append(j - 1)
                matched_ref_indices.append(i - 1)
            i -= 1
            j -= 1
        elif ptr == 2:  # up
            i -= 1
        else:  # left
            j -= 1

    matched_current_indices.reverse()
    matched_ref_indices.reverse()
    return matched_current_indices, matched_ref_indices, best_score

# Slots
input_tables = input_slot_map["Input"]
output_labels = output_slot_map["Labels"]
output_values = output_slot_map["Values"]
output_reconstructed = output_slot_map["Reconstructed"]
output_failed = output_slot_map["Failed"]

# Get the row indices from the input slot
input_rows = [x for x in data_batch.getInputRows("Input")]

# Globals
ref_strip_sequence = json.loads(
    "[" + input_tables.getTextAnnotation(input_rows[0],
"StripSequence").getValue() + "]")
interpolation = "ceil"
imgw = int(input_tables.getTextAnnotation(input_rows[0], "iw").getValue())
imgh = int(input_tables.getTextAnnotation(input_rows[0], "ih").getValue())
shift_y = int(input_tables.getTextAnnotation(input_rows[0], "cy").getValue())

progress_info.log("")
progress_info.log("Ref sequence:")
progress_info.log(str(ref_strip_sequence))

best_match_indices, best_match_ref_indices, best_match_score = None, None, 0
best_strip_sequence, best_strip_locations = None, None
```

```
for row in input_rows:

    input_table = input_tables.getData(row, ResultsTableData, progress_info)
    current_strip_sequence = []
    current_strip_locations = []

    for i in range(input_table.getRowCount()):
        current_strip_sequence.append(input_table.getValueAsString(i,
"text"))
        current_strip_locations.append({"i": i, "x":
input_table.getValueAsDouble(
            i, "left"), "y": input_table.getValueAsDouble(i, "top"), "h":
input_table.getValueAsDouble(i, "height")})

    # The indices within current that match with with a reference strip
    match_indices, match_ref_indices, match_score =
smith_waterman_current_match_indices(
        reference=ref_strip_sequence, current=current_strip_sequence)

    if len(match_indices) >= 2 and match_score > best_match_score:

        progress_info.log("")
        progress_info.log("Found better match with score " +
str(match_score))

        best_match_indices = match_indices
        best_match_ref_indices = match_ref_indices
        best_match_score = match_score
        best_strip_sequence = current_strip_sequence
        best_strip_locations = current_strip_locations

        progress_info.log("")
        progress_info.log("Current strip:")
        progress_info.log(str([x for x in current_strip_sequence]))
        progress_info.log(str([num(x) for x in current_strip_sequence]))

        progress_info.log("")
        progress_info.log("Matches within current:")
        progress_info.log(str(match_indices))

        progress_info.log("")
        progress_info.log("Matches within ref:")
        progress_info.log(str(match_ref_indices))

        progress_info.log("--------")

if best_match_indices is not None:

    # Get the best info
    match_indices = best_match_indices
    match_ref_indices = best_match_ref_indices
    current_strip_locations = best_strip_locations
    current_strip_sequence = best_strip_sequence
```

```
    # Find the average y distance between ticks (assuming that tick distance
is equal; assume perfectly vertical)
    sum_avgdy = 0
    num_avgdy = 0

    for i in range(len(match_indices)):
        for j in range(i + 1, len(match_indices)):
            index1 = match_indices[i]
            index2 = match_indices[j]
            ref_index1 = match_ref_indices[i]
            ref_index2 = match_ref_indices[j]

            d = abs(ref_index1 - ref_index2)
            y1 = current_strip_locations[index1]["y"]
            y2 = current_strip_locations[index2]["y"]
            avgdy = abs(y2 - y1) / d

            sum_avgdy += avgdy
            num_avgdy += 1
            # progress_info.log(str(i) + "<>" + str(j) + " = " + str(avgdy))

    avgdy = 1.0 * sum_avgdy / num_avgdy

    progress_info.log("")
    progress_info.log("Avg distance between tick marks: " + str(avgdy) +
"px")

    # Find average height
    avgh = mean([x["h"] for x in current_strip_locations])

    progress_info.log("")
    progress_info.log("Avg distance letter height: " + str(avgh) + "px")

    # Recover the y locations of all reference strip entries
    ref_strip_y = [None for v in ref_strip_sequence]

    for i in range(len(match_indices)):
        index_in_current = match_indices[i]
        index_in_ref = match_ref_indices[i]
        pos = current_strip_locations[index_in_current]

        ref_strip_y[index_in_ref] = pos["y"]

    progress_info.log("")
    progress_info.log("Initial y locations:")
    progress_info.log(str(ref_strip_sequence))
    progress_info.log(str(ref_strip_y))

    orig_ref_strip_y = [x for x in ref_strip_y]

    for i in range(len(ref_strip_y)):
        if ref_strip_y[i] is None:

            # Find the closest detected tick (bias towards earlier strip refs
if same score)
```

```
            di = 99999
            y = None

            for j in range(len(ref_strip_y)):
                oy = orig_ref_strip_y[j]
                di2 = i - j
                if i != j and abs(di2) < abs(di) and oy is not None:
                    di = di2
                    y = oy

            assert y is not None

            ref_strip_y[i] = y + di * avgdy

    progress_info.log("")
    progress_info.log("Initial+Updated y locations:")
    progress_info.log(str(ref_strip_sequence))
    progress_info.log(str(orig_ref_strip_y))
    progress_info.log(str(ref_strip_y))

    # Correct y by letter height
    for i in range(len(ref_strip_y)):
        ref_strip_y[i] += avgh / 2

    # Output locations
    reconstructed = ResultsTableData()
    for i in range(len(ref_strip_sequence)):
        reconstructed.addAndModifyRow().set("v", ref_strip_sequence[i]).set(
            "y_cropped", ref_strip_y[i]).set("y", ref_strip_y[i] + shift_y)
    data_batch.addOutputData(output_reconstructed,
                             reconstructed, progress_info)

    # Create labels and values by interpolating within pixel space
    labels_ip = FloatProcessor(1, imgh)
    values_ip = FloatProcessor(1, imgh)

    miny = int(ref_strip_y[0] + shift_y)
    maxy = int(ref_strip_y[-1] + shift_y)

    if maxy < imgh:
        progress_info.log("Created ip w=" + str(imgw) + " h=" + str(imgh) +
                          " miny=" + str(miny) + " maxy=" + str(maxy) + "
shifty=" + str(shift_y))

        if interpolation == "linear":
            # Linear interpolation between the two points

            # Generate labels
            for i in range(maxy - miny + 1):
                labels_ip.setf(0, miny + i, i + 1)

            # Generate values
            i0 = 0
            i1 = 1
```

```
            for i in range(maxy - miny + 1):
                y0 = ref_strip_y[i0] + shift_y
                y1 = ref_strip_y[i1] + shift_y
                v0 = ref_strip_sequence[i0]
                v1 = ref_strip_sequence[i1]
                y = miny + i

                p = min(1, max(0, 1.0 * (y - y0) / (y1 - y0)))
                v = v0 + p * (v1 - v0)

                # progress_info.log(str(y) + " IN " + str(y0) + ", " +
str(y1) + " -> p=" + str(p) + " -> v=" + str(v))

                values_ip.setf(0, y, v)

                if (y + 1) >= y1 and (i1 + 1) < len(ref_strip_y):
                    i0 += 1
                    i1 += 1
        elif interpolation == "ceil":
            # Assigns label to the higher value
            # "the "proper" way to read the MIC if the ZOI edge is between
two ticks is to take the higher concentration. e.g. if the ZOI is between 1
and 1.5 like in the example you attached, the MIC should be recorded as 1.5."

            # Generate values and labels
            i0 = 0
            i1 = 1
            lbl = 0
            lbl_v0 = float("nan")

            for i in range(maxy - miny + 1):
                y0 = ref_strip_y[i0] + shift_y
                y1 = ref_strip_y[i1] + shift_y
                v0 = ref_strip_sequence[i0]
                v1 = ref_strip_sequence[i1]
                y = miny + i

                v = v0  # Always the previous value (higher)

                if v != lbl_v0:
                    lbl += 1
                    lbl_v0 = v

                values_ip.setf(0, y, v)
                labels_ip.setf(0, y, lbl)

                if (y + 1) >= y1 and (i1 + 1) < len(ref_strip_y):
                    i0 += 1
                    i1 += 1
        else:
            raise ValueError("Unsupported interpolation")

        labels_ip = labels_ip.resize(imgw, imgh)
        values_ip = values_ip.resize(imgw, imgh)
```

```
        # Output everything
        data_batch.addOutputData(output_labels, ImagePlusData(
            ImagePlus("labels", labels_ip)), progress_info)
        data_batch.addOutputData(output_values, ImagePlusData(
            ImagePlus("values", values_ip)), progress_info)

    else:
        progress_info.log("Prediction is wrong! (maxy condition) ip w=" +
str(imgw) + " h=" + str(
            imgh) + " miny=" + str(miny) + " maxy=" + str(maxy) + " shifty="
+ str(shift_y))
        data_batch.addOutputData(output_failed, input_table, progress_info)
else:
    progress_info.log(
        "Unable to match detected sequence with expected sequence")
    data_batch.addOutputData(output_failed, input_table, progress_info)
```

###### *Node #69 "Output" of type "Compartment output"*

- Input "Labels" of node #69 receives data from output "Labels" of node #68
- Input "Values" of node #69 receives data from output "Values" of node #68
- Input "Initial" of node #69 receives data from output "Output" of node #67
- Input "Reconstructed" of node #69 receives data from output "Reconstructed" of node #68
- Input "Failed" of node #69 receives data from output "Failed" of node #68
- Input "Cropped Strip" of node #69 receives data from output "Cropped" of node #59
- The parameter "jipipe:compartment:output-slot-name" (jipipe:compartment:output-slot-name) is set to **"Output"**

#### Compartment C4 “Labels”

This compartment is responsible for creating the final labels for the measurement process, adding a safety margin around the strip, and removing the areas outside the plate area. A Screenshot is provided in **Supplementary Figure 40**.

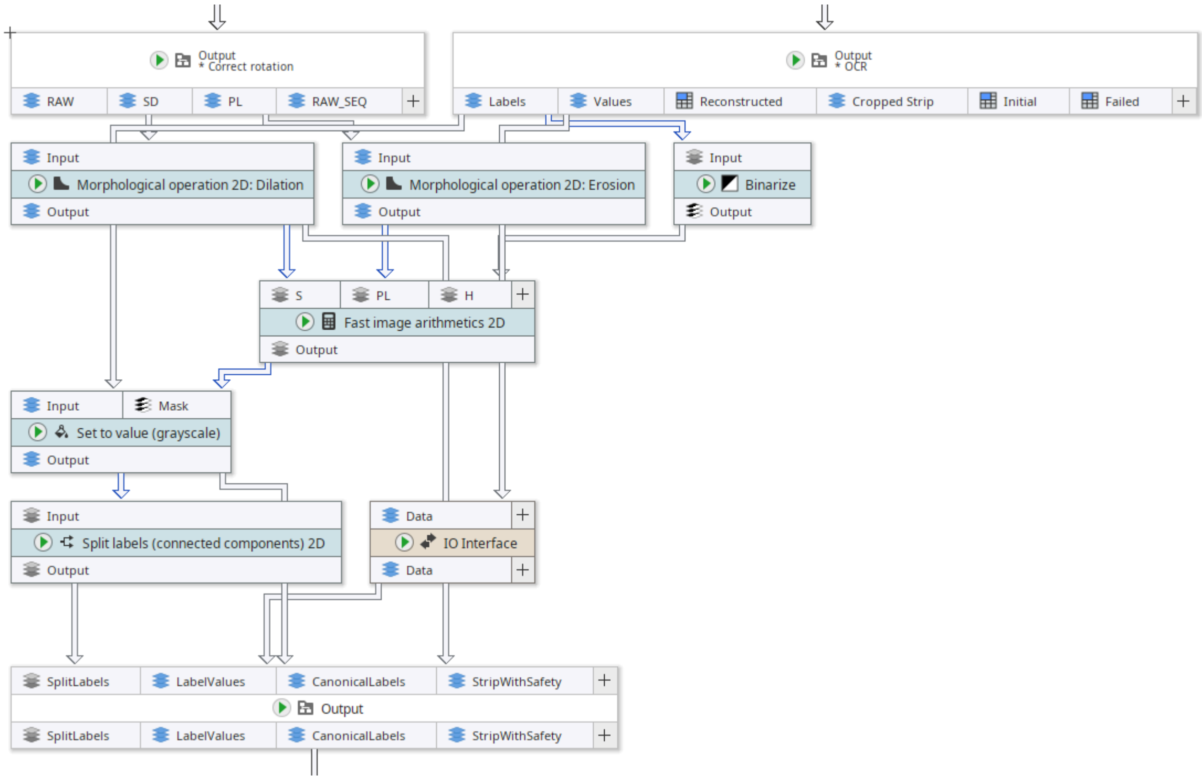

**Supplementary Figure 40:** "Labels" compartment of the MIC analysis pipeline.

##### *Node #55 "Output" of type "Compartment output"*

- Input "RAW" of node #55 receives data from output "Output" of node #50
- Input "SD" of node #55 receives data from output "Output" of node #54
- Input "PL" of node #55 receives data from output "Output" of node #53
- Input "RAW_SEQ" of node #55 receives data from output "RAW" of node #24
- The parameter "jipipe:compartment:output-slot-name" (jipipe:compartment:output-slot-name) is set to **"Output"**

##### *Node #70 "Morphological operation 2D: Dilation"*

- Input "Input" of node #70 receives data from output "SD" of node #55
- The parameter "Structure element" (element) is set to **"SQUARE"**
- The parameter item #1 of "Overridden parameters" in category "Adaptive parameters" (jipipe:adaptive-parameters/overridden-parameters) is set to **(_global.stripSafetyDistanceMillimeters * 2 / NUM(PixelSizeRescaled) * 0.5, "radius")**

##### *Node #71 "Morphological operation 2D: Erosion"*

- Input "Input" of node #71 receives data from output "PL" of node #55
- The parameter "Operation" (operation) is set to **"EROSION"**
- The parameter item #1 of "Overridden parameters" in category "Adaptive parameters" (jipipe:adaptive-parameters/overridden-parameters) is set to **(5 / NUM(PixelSizeRescaled), "radius")**

##### *Node #69 "Output" of type "Compartment output"*

- Input "Labels" of node #69 receives data from output "Labels" of node #68
- Input "Values" of node #69 receives data from output "Values" of node #68
- Input "Initial" of node #69 receives data from output "Output" of node #67

- Input "Reconstructed" of node #69 receives data from output "Reconstructed" of node #68
- Input "Failed" of node #69 receives data from output "Failed" of node #68
- Input "Cropped Strip" of node #69 receives data from output "Cropped" of node #59
- The parameter "jipipe:compartment:output-slot-name" (jipipe:compartment:output-slot-name) is set to **"Output"**

***Node #72 "Binarize" of type "Binarize"***

- Input "Input" of node #72 receives data from output "Labels" of node #69

***Node #73 "IO Interface"***

- Input "Data" of node #73 receives data from output "Values" of node #69

***Node #74 "Fast image arithmetics 2D"***

- Input "S" of node #74 receives data from output "Output" of node #70
- Input "PL" of node #74 receives data from output "Output" of node #71
- Input "H" of node #74 receives data from output "Output" of node #72
- The parameter "Expression" (expression) is set to **MAX(0, MIN(PL, H) - S)**
- The parameter "Skip incomplete data sets" in category "Input management" (jipipe:data-batch-generation/skip-incomplete) is set to **true**

***Node #75 "Set to value (grayscale)"***

- Input "Input" of node #75 receives data from output "Labels" of node #69
- Input "Mask" of node #75 receives data from output "Output" of node #74
- The parameter "Only apply to ..." (roi:target-area) is set to **"OutsideMask"**

***Node #76 "Split labels (connected components) 2D"***

- Input "Input" of node #76 receives data from output "Output" of node #75

***Node #77 "Output" of type "Compartment output"***

- Input "SplitLabels" of node #77 receives data from output "Output" of node #76
- Input "LabelValues" of node #77 receives data from output "Data" of node #73
- Input "CanonicalLabels" of node #77 receives data from output "Output" of node #75
- Input "StripWithSafety" of node #77 receives data from output "Output" of node #70
- The parameter "jipipe:compartment:output-slot-name" (jipipe:compartment:output-slot-name) is set to **"Output"**

#### Compartment C5 “Visualize strip”

This compartment is responsible for creating a visualization that allows the evaluation of the performance of the OCR compartment. A screenshot is provided in **Supplementary Figure 41**.

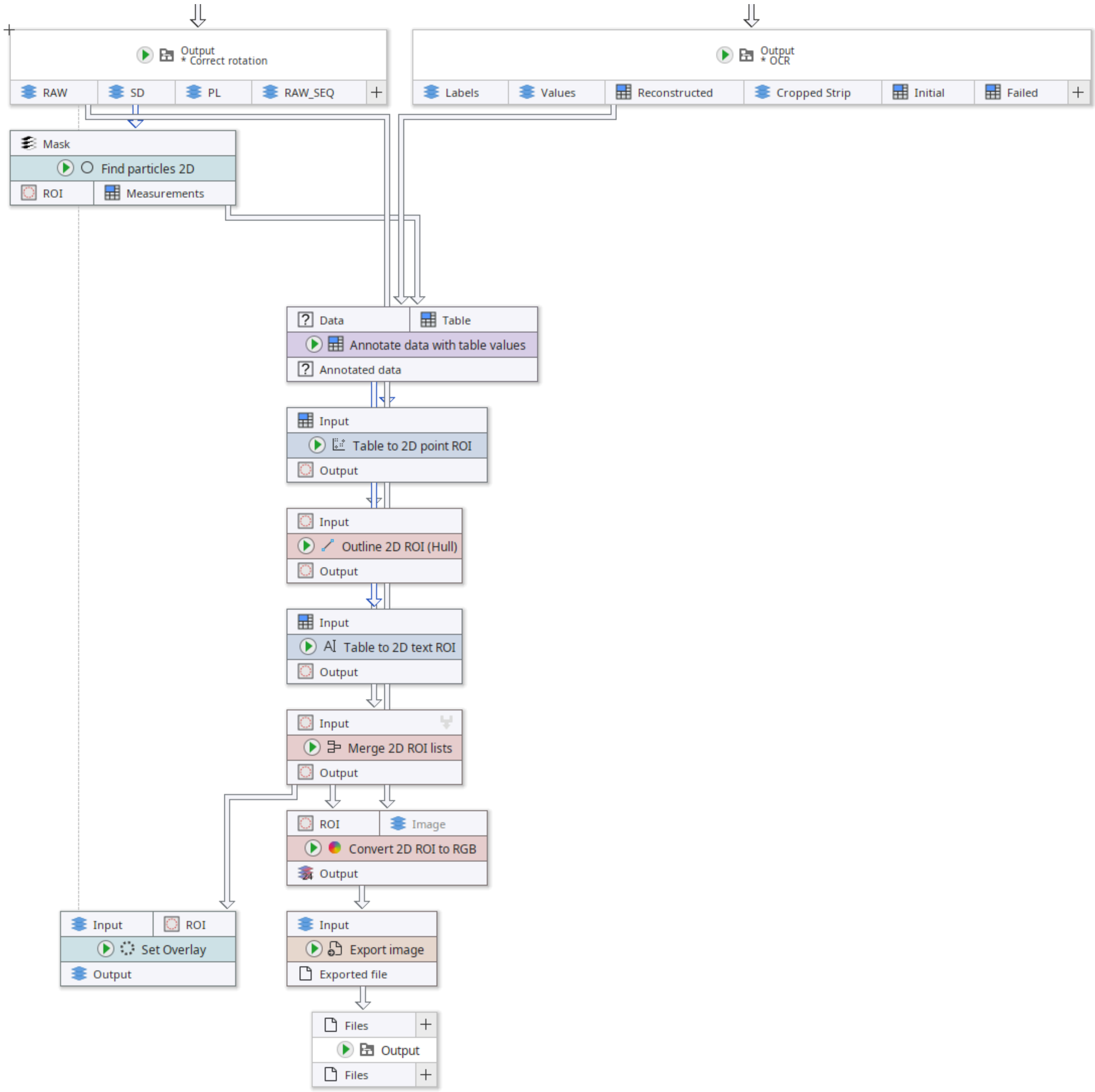

**Supplementary Figure 41:** “Visualize strip” compartment of the MIC analysis pipeline.

##### *Node #55 "Output" of type "Compartment output"*

- Input "RAW" of node #55 receives data from output "Output" of node #50
- Input "SD" of node #55 receives data from output "Output" of node #54
- Input "PL" of node #55 receives data from output "Output" of node #53
- Input "RAW_SEQ" of node #55 receives data from output "RAW" of node #24
- The parameter "jipipe:compartment:output-slot-name" (jipipe:compartment:output-slot-name) is set to **"Output"**

##### *Node #78 "Find particles 2D"*

- Input "Mask" of node #78 receives data from output "SD" of node #55
- The parameter "Extracted measurements" (measurements) is set to **{"values":["BoundingRectangle"],"collapsed":false}**

##### *Node #69 "Output" of type "Compartment output"*

- Input "Labels" of node #69 receives data from output "Labels" of node #68

- Input "Values" of node #69 receives data from output "Values" of node #68
- Input "Initial" of node #69 receives data from output "Output" of node #67
- Input "Reconstructed" of node #69 receives data from output "Reconstructed" of node #68
- Input "Failed" of node #69 receives data from output "Failed" of node #68
- Input "Cropped Strip" of node #69 receives data from output "Cropped" of node #59
- The parameter "jipipe:compartment:output-slot-name" (jipipe:compartment:output-slot-name) is set to **"Output"**

***Node #79 "Annotate data with table values"***

- Input "Data" of node #79 receives data from output "Reconstructed" of node #69
- Input "Table" of node #79 receives data from output "Measurements" of node #78
- The parameter item #1 of "Generated annotations" (generated-annotations) is set to **name = "sw", value = FIRST_OF(Width)**
- The parameter "Skip incomplete data sets" in category "Input management" (jipipe:data-batch-generation/skip-incomplete) is set to **true**

***Node #80 "Table to 2D text ROI"***

- Input "Input" of node #80 receives data from output "Annotated data" of node #79
- The parameter "Font size" (font-size) is set to **22**
- The parameter "Column 'Y'" (column-y) is set to **("ExistingColumn", "y")**
- The parameter "Column 'X'" (column-x) is set to **("Generate", NUM(cx) + NUM(sw) / 2)**
- The parameter "Background margin" (background-margin) is set to **{"left":{"expression":"2.0"},"top":{"expression":"2.0"},"right":{"expression":"2.0"},"bottom":{"expression":"2.0"}}**
- The parameter "Column 'Text'" (column-text) is set to **("ExistingColumn", "v")**

***Node #81 "Table to 2D point ROI"***

- Input "Input" of node #81 receives data from output "Annotated data" of node #79
- The parameter "Column 'X'" (column-x1) is set to **("Generate", NUM(cx) + NUM(sw) / 2)**
- The parameter "ROI name" (column-name) is set to **("Generate", "v")**
- The parameter "Column 'Y'" (column-y1) is set to **("ExistingColumn", "y")**

***Node #82 "Outline 2D ROI (Hull)"***

- Input "Input" of node #82 receives data from output "Output" of node #81
- The parameter "Outline method" (outline) is set to **"BoundingRectangle"**

***Node #83 "Merge 2D ROI lists"***

- Input "Input" of node #83 receives data from output "Output" of node #80
- Input "Input" of node #83 receives data from output "Output" of node #82

***Node #84 "Set Overlay" of type "Set 2D overlay"***

- Input "Input" of node #84 receives data from output "RAW" of node #55
- Input "ROI" of node #84 receives data from output "Output" of node #83
- The parameter "Skip incomplete data sets" in category "Input management" (jipipe:data-batch-generation/skip-incomplete) is set to **true**

***Node #85 "Convert 2D ROI to RGB"***

- Input "ROI" of node #85 receives data from output "Output" of node #83
- Input "Image" of node #85 receives data from output "RAW" of node #55

- The parameter "Override fill color" (override-fill-color) is set to **"#FF0000"**
- The parameter "Override line width" (override-line-width) is set to **5.0**
- The parameter "Override line color" (override-line-color) is set to **"#FF0000"**
- The parameter "Skip incomplete data sets" in category "Input management" (jipipe:data-batch-generation/skip-incomplete) is set to **true**

***Node #86 "Export image"***

- Input "Input" of node #86 receives data from output "Output" of node #85
- The parameter "File path" (file-path) is set to **PATH_COMBINE(project_data_dir.tmp_dir, "results", "predicted_reference_points", #ImageId + "_" + #AssayType + "_" + #Experiment + "_" + #Sample + "_" + #TimePoint)**

***Node #87 "Output" of type "Compartment output"***

- Input "Files" of node #87 receives data from output "Exported file" of node #86
- The parameter "jipipe:compartment:output-slot-name" (jipipe:compartment:output-slot-name) is set to **"Output"**

#### Compartment C6 “Measure”

This compartment applies the script for measuring the MIC and exports results in the form of a table. A screenshot is provided in **Supplementary Figure 42**.

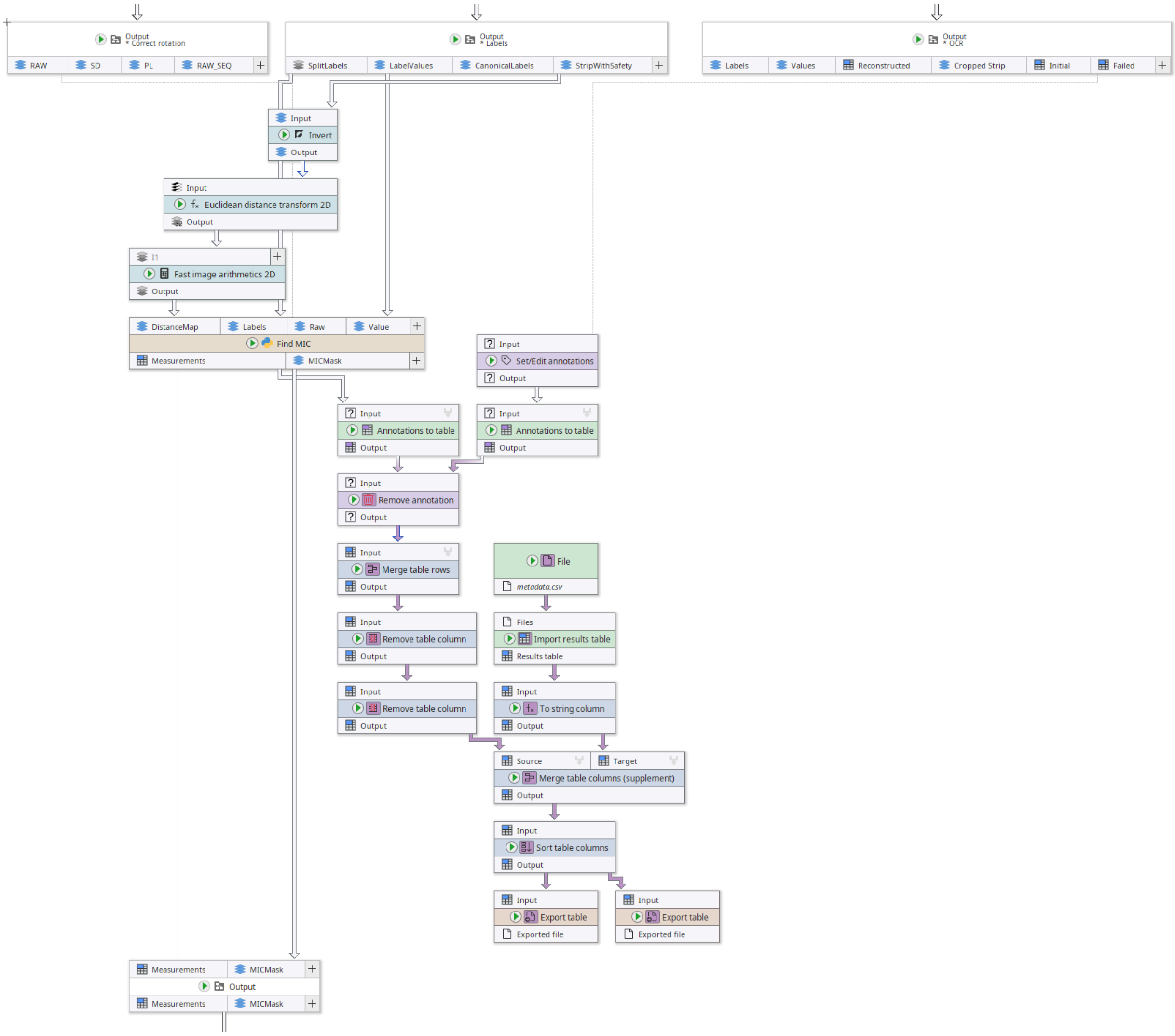

**Supplementary Figure 42:** “Measure” compartment of the MIC analysis pipeline.

##### *Node #88 "File"*

- The parameter "File name" (file-name) is set to **"metadata.csv"**

##### *Node #89 "Import results table"*

- Input "Files" of node #89 receives data from output "Filename" of node #88

##### *Node #90 "To string column"*

- Input "Input" of node #90 receives data from output "Results table" of node #89

##### *Node #55 "Output" of type "Compartment output"*

- Input "RAW" of node #55 receives data from output "Output" of node #50
- Input "SD" of node #55 receives data from output "Output" of node #54
- Input "PL" of node #55 receives data from output "Output" of node #53
- Input "RAW_SEQ" of node #55 receives data from output "RAW" of node #24
- The parameter "jipipe:compartment:output-slot-name" (jipipe:compartment:output-slot-name) is set to **"Output"**

***Node #69 "Output" of type "Compartment output"***

- Input "Labels" of node #69 receives data from output "Labels" of node #68
- Input "Values" of node #69 receives data from output "Values" of node #68
- Input "Initial" of node #69 receives data from output "Output" of node #67
- Input "Reconstructed" of node #69 receives data from output "Reconstructed" of node #68
- Input "Failed" of node #69 receives data from output "Failed" of node #68
- Input "Cropped Strip" of node #69 receives data from output "Cropped" of node #59
- The parameter "jipipe:compartment:output-slot-name" (jipipe:compartment:output-slot-name) is set to **"Output"**

***Node #91 "Set/Edit annotations"***

- Input "Input" of node #91 receives data from output "Failed" of node #69
- The parameter item #1 of "Annotations" (generated-annotation) is set to **("OCR_FAILURE", "MIC")**

***Node #92 "Annotations to table" of type "Convert to annotation table"***

- Input "Input" of node #92 receives data from output "Output" of node #91

***Node #77 "Output" of type "Compartment output"***

- Input "SplitLabels" of node #77 receives data from output "Output" of node #76
- Input "LabelValues" of node #77 receives data from output "Data" of node #73
- Input "CanonicalLabels" of node #77 receives data from output "Output" of node #75
- Input "StripWithSafety" of node #77 receives data from output "Output" of node #70
- The parameter "jipipe:compartment:output-slot-name" (jipipe:compartment:output-slot-name) is set to **"Output"**

***Node #93 "Invert" of type "Invert pixel colors/values"***

- Input "Input" of node #93 receives data from output "StripWithSafety" of node #77

***Node #94 "Euclidean distance transform 2D"***

- Input "Input" of node #94 receives data from output "Output" of node #93

***Node #95 "Fast image arithmetics 2D"***

- Input "I1" of node #95 receives data from output "Output" of node #94
- The parameter "Expression" (expression) is set to **I1 * PixelSizeRescaled**

***Node #96 "Find MIC" of type "Jython script (iterating)"***

- Input "DistanceMap" of node #96 receives data from output "Output" of node #95
- Input "Labels" of node #96 receives data from output "SplitLabels" of node #77
- Input "Raw" of node #96 receives data from output "RAW" of node #55
- Input "Value" of node #96 receives data from output "LabelValues" of node #77
- The parameter "Enable inverse detection" in category "Script parameters" (script-parameters/enableInverseDetection) is set to **false**

- The parameter "Inverse detection variation threshold" in category "Script parameters" (script-parameters/inverseDetectionVariationThreshold) is set to **0.05**
- The parameter "Inverse detection min concentrations" in category "Script parameters" (script-parameters/inverseDetectionMinConcentrations) is set to **3.0**
- The parameter "Ring width (mm)" in category "Script parameters" (script-parameters/ringWidthMillimeters) is set to **0.3**
- The parameter "Ring width max (mm)" in category "Script parameters" (script-parameters/ringWidthMaxMillimeters) is set to **30.0**
- The parameter "ZOI boundary detection lawn fraction" in category "Script parameters" (script-parameters/zoiBoundaryDetectionLawnFraction) is set to **0.3**
- The parameter "ZOI boundary detection min inhibition threshold (mm)" in category "Script parameters" (script-parameters/zoiBoundaryDetectionMinInhibitionThresholdMillimeters) is set to **0.5**
- The parameter "Corridor width wide (mm)" in category "Script parameters" (script-parameters/corridorWidthWideMillimeters) is set to **3.0**
- The parameter "Corridor width narrow (mm)" in category "Script parameters" (script-parameters/corridorWidthNarrowMillimeters) is set to **0.3**
- The parameter "ZOI boundary wide threshold (mm)" in category "Script parameters" (script-parameters/zoiBoundaryWideThresholdMillimeters) is set to **7.2**
- The parameter "Enable NA detection" in category "Script parameters" (script-parameters/enableNADetection) is set to **true**
- The parameter "Scale factor" in category "Script parameters" (script-parameters/scaleFactor) is set to **1.0**
- The parameter "NA detection contrast threshold" in category "Script parameters" (script-parameters/naDetectionContrastThreshold) is set to **0.44**
- The parameter "NA detection close ZOI boundary threshold (mm)" in category "Script parameters" (script-parameters/naDetectionCloseZOIBoundaryThresholdMillimeters) is set to **1.5**
- The parameter "NA detection max coefficient of variation at high" in category "Script parameters" (script-parameters/naDetectionMaxCoefficientOfVariationAtHigh) is set to **0.15**
- The parameter "MIC threshold N reference" in category "Script parameters" (script-parameters/micThresholdNReference) is set to **3**
- The parameter "MIC threshold" in category "Script parameters" (script-parameters/micThreshold) is set to **0.255**

The script executes the following Jython script that handles the core aspects of the MIC detection, including extracting of measurements, N/A detection, and calculation of the final MIC value.

```
"""
Input Slots:
- "DistanceMap" (ImageData) -- Pre-computed distance map (distance from strip
edge in mm)
- "Labels" (ImageData) -- Split canonical label image (each label = separate
zone)
- "Raw" (ImageData) -- Raw intensity image
- "Value" (ImageData) -- Concentration value image (pixel value = antibiotic
concentration)
```

```
Output Slots:
- "Measurements" (ResultsTableData) -- Per-concentration measurements for the
selected corridor
- "MICMask" (ImageData) -- Binary image highlighting the label(s) associated
with the predicted MIC

Text Annotations (Input):
- "StripSequence" -- Strip concentration sequence (e.g. "256;128;64;...")
- PIXEL_SIZE_ANNOTATION_NAME -- Physical pixel size in mm

Text Annotations (Output):
- "MIC" -- Predicted MIC string (e.g. "0.5") or "N/A"
- "is_na" -- "true" or "false"
- "na_reason" -- Reason string if N/A, else ""
- "is_narrow" -- "true" or "false"
- "corridor_width_mm" -- Selected corridor width in mm
- "d_high" -- Max ZOI boundary distance in mm
- "mic_threshold" -- MIC threshold value used
"""

from org.hkijena.jipipe.api.data import JIPipeData
from org.hkijena.jipipe.api.annotation import JIPipeTextAnnotation,
JIPipeTextAnnotationMergeMode
from org.hkijena.jipipe.plugins.tables.datatypes import ResultsTableData
from org.hkijena.jipipe.plugins.imagejdatatypes.datatypes import
ImagePlusData
from ij import ImagePlus
from ij.process import FloatProcessor
import math

#
============================================================================
# Configuration Constants
#
============================================================================

# --------------------------------------------------------------------------
# Concentration matching
# --------------------------------------------------------------------------
CONC_MATCH_LOG_TOLERANCE = 0.1       # Max log-space difference for
concentration matching (we have exact values)

# --------------------------------------------------------------------------
# Inverted contrast detection
# --------------------------------------------------------------------------
ENABLE_INVERSE_DETECTION = enableInverseDetection        # Set to False to
skip inverse contrast detection (default false)
```

```
CONTRAST_VARIATION_THRESHOLD = inverseDetectionVariationThreshold   # Min
relative intensity variation to determine contrast (default 0.05)
CONTRAST_MIN_CONCENTRATIONS = inverseDetectionMinConcentrations       # Min
unique concentrations needed for contrast detection (default 3)

# ---------------------------------------------------------------------------
# Ring matrix extraction (for wide/narrow classification)
# ---------------------------------------------------------------------------
RING_WIDTH_MM = ringWidthMillimeters                  # Width of each ring
bin in mm (default 0.3)
EXTENDED_MAX_RING_WIDTH_MM = ringWidthMaxMillimeters    # Maximum distance
for ring matrix (default 30)

# ---------------------------------------------------------------------------
# ZOI boundary detection
# ---------------------------------------------------------------------------
LAWN_FRACTION = zoiBoundaryDetectionLawnFraction                   # Fraction
of lawn intensity for ZOI boundary threshold (default 0.3)
MIN_INHIBITION_THRESHOLD_MM =
zoiBoundaryDetectionMinInhibitionThresholdMillimeters    # Minimum boundary
distance to consider valid (default 0.5)

# ---------------------------------------------------------------------------
# Wide/narrow classification
# ---------------------------------------------------------------------------
D_HIGH_THRESHOLD = zoiBoundaryWideThresholdMillimeters               # d_high
threshold: <= this -> narrow, > this -> wide (default 7.2)
CORRIDOR_WIDTH_WIDE_MM = corridorWidthWideMillimeters         # Corridor
width for wide ZOI (default 3)
CORRIDOR_WIDTH_NARROW_MM = corridorWidthNarrowMillimeters       # Corridor
width for narrow ZOI (default 0.3)

# ---------------------------------------------------------------------------
# N/A detection thresholds (fitting-free, 3 OR-combined rules)
# ---------------------------------------------------------------------------
NA_MIN_GROWTH_RANGE_RATIO_THRESHOLD = naDetectionContrastThreshold  # Rule 1:
min(grr_wide, grr_narrow) — low growth at either corridor (default 0.44)
NA_MAX_BOUNDARY_MM_THRESHOLD =
naDetectionCloseZOIBoundaryThresholdMillimeters         # Rule 2: max ZOI
boundary distance (d_high) (default 1.50)
NA_HI_CONC_CV_THRESHOLD = naDetectionMaxCoefficientOfVariationAtHigh
# Rule 3: CV of top-3 highest-conc measurements (growth inside ZOI) (default
0.15)
ENABLE_NA_DETECTION = enableNADetection  # Set to False to disable N/A
detection and always report MIC (default true)

# ---------------------------------------------------------------------------
# MIC threshold settings
```

```
# ---------------------------------------------------------------------------
N_REFERENCE = micThresholdNReference                      # Number of
reference concentrations for background/growth (default 3)
DEFAULT_MIC_THRESHOLD = micThreshold         # Default MIC threshold (default
0.255)

# The annotation that contains the pixel size
PIXEL_SIZE_ANNOTATION_NAME = "PixelSizeRescaled" # In JIPipe workflow this is
the correct annotation

# ---------------------------------------------------------------------------
# Image scaling
# ---------------------------------------------------------------------------
IMAGE_SCALE_FACTOR = scaleFactor    # Scale factor for image resizing (1.0 =
no scaling, 0.5 = half size)
IMAGE_INTERPOLATION = 1     # Default interpolation method:
0=NEAREST_NEIGHBOR, 1=BILINEAR, 2=BICUBIC

#
============================================================================
# Utility Functions
#
============================================================================

def is_valid_number(v):
    """Check if a value is a valid finite number."""
    try:
        f = float(v)
    except Exception:
        return False
    if math.isnan(f) or math.isinf(f):
        return False
    return True

def applyScaling(ip, interpolation=1):
    """Apply image scaling if IMAGE_SCALE_FACTOR != 1.0.

    Args:
        ip: ImageProcessor to scale
        interpolation: Interpolation method (0=NEAREST_NEIGHBOR, 1=BILINEAR,
2=BICUBIC)

    Returns:
```

```
        Scaled ImageProcessor if IMAGE_SCALE_FACTOR != 1.0, otherwise the
original ip
    """
    if IMAGE_SCALE_FACTOR == 1.0:
        return ip
    w = ip.getWidth()
    h = ip.getHeight()
    new_w = int(w * IMAGE_SCALE_FACTOR)
    new_h = int(h * IMAGE_SCALE_FACTOR)
    ip.setInterpolationMethod(interpolation)
    return ip.resize(new_w, new_h, True)

def parse_strip_sequence(strip_sequence_text):
    """Parse strip sequence text into a list of positive concentration
values.

    Handles both comma and semicolon separators.
    """
    values = []
    if strip_sequence_text is None:
        return values
    text = str(strip_sequence_text)
    if ";" in text:
        parts = text.split(";")
    else:
        parts = text.split(",")
    i = 0
    while i < len(parts):
        t = parts[i].strip()
        if len(t) > 0:
            try:
                v = float(t)
                if v > 0:
                    values.append(v)
            except (ValueError, OverflowError):
                pass
        i += 1
    return values

def parse_strip_sequence_pairs(strip_sequence_text):
    """Parse strip sequence text into list of (numeric_value, original_token)
pairs.

    Preserves original token formatting for MIC string output.
    """
    pairs = []
```

```
    if strip_sequence_text is None:
        return pairs
    text = str(strip_sequence_text)
    if ";" in text:
        parts = text.split(";")
    else:
        parts = text.split(",")
    i = 0
    while i < len(parts):
        token = parts[i]
        t = token.strip()
        if len(t) > 0:
            try:
                v = float(t)
                if v > 0 and not math.isnan(v) and not math.isinf(v):
                    pairs.append((v, t))
            except (ValueError, OverflowError):
                pass
        i += 1
    return pairs

#
=============================================================================
# Step 2: Inverted Contrast Detection
#
=============================================================================

def detect_inverted_contrast(ip_raw, ip_value, ip_labels, w, h):
    """Detect if the image has inverted contrast (higher concentration ->
higher intensity).

    For E-test strips, higher concentration should correspond to LOWER
intensity
    (more inhibition = less growth = darker area). If the Pearson correlation
    between log(concentration) and mean intensity is positive, contrast is
inverted.

    Returns True if contrast is inverted, False otherwise.
    """
    # Collect intensity values per unique concentration for labeled pixels
    conc_intensity_map = {}  # concentration -> list of intensities

    y = 0
    while y < h:
        x = 0
        while x < w:
```

```
            label_val = ip_labels.getf(x, y)
            if label_val > 0:
                conc_val = ip_value.getf(x, y)
                raw_val = ip_raw.getf(x, y)
                conc_rounded = round(conc_val, 6)
                if conc_rounded > 0:
                    if conc_rounded not in conc_intensity_map:
                        conc_intensity_map[conc_rounded] = []
                    conc_intensity_map[conc_rounded].append(raw_val)
            x += 1
        y += 1

    if len(conc_intensity_map) < CONTRAST_MIN_CONCENTRATIONS:
        return False

    # Compute mean intensity per concentration
    log_concs = []
    mean_intensities = []
    conc_keys = list(conc_intensity_map.keys())
    i = 0
    while i < len(conc_keys):
        conc = conc_keys[i]
        intensities = conc_intensity_map[conc]
        if len(intensities) == 0:
            i += 1
            continue
        mean_val = sum(intensities) / float(len(intensities))
        log_concs.append(math.log(float(conc)))
        mean_intensities.append(mean_val)
        i += 1

    if len(log_concs) < CONTRAST_MIN_CONCENTRATIONS:
        return False

    # Check if intensity variation is too small to reliably determine
contrast
    mean_intensity = sum(mean_intensities) / float(len(mean_intensities))
    intensity_range = max(mean_intensities) - min(mean_intensities)
    if mean_intensity > 0 and intensity_range / mean_intensity <
CONTRAST_VARIATION_THRESHOLD:
        return False

    # Compute Pearson correlation between log(concentration) and mean
intensity
    n = len(log_concs)
    sum_x = 0.0
    sum_y = 0.0
    sum_xy = 0.0
```

```
    sum_x2 = 0.0
    sum_y2 = 0.0
    i = 0
    while i < n:
        sum_x += log_concs[i]
        sum_y += mean_intensities[i]
        sum_xy += log_concs[i] * mean_intensities[i]
        sum_x2 += log_concs[i] * log_concs[i]
        sum_y2 += mean_intensities[i] * mean_intensities[i]
        i += 1

    denom_x = n * sum_x2 - sum_x * sum_x
    denom_y = n * sum_y2 - sum_y * sum_y
    if denom_x <= 0 or denom_y <= 0:
        return False

    correlation = (n * sum_xy - sum_x * sum_y) / math.sqrt(denom_x * denom_y)

    return correlation > 0

#
============================================================================
# Step 3b: Strip Edge Detection & Concentration Row Assignment
#
============================================================================

def find_strip_edges(ip_labels, w, h):
    """Find left/right edges of the strip gap per row.

    For each row, finds the largest gap between labeled columns, which
    represents the strip gap between left and right sides.

    Returns (strip_left, strip_right) lists of length h.
    Elements are None for rows where no strip gap was found.
    """
    strip_left = [None] * h
    strip_right = [None] * h

    y = 0
    while y < h:
        # Collect labeled columns for this row
        labeled_cols = []
        x = 0
        while x < w:
            if ip_labels.getf(x, y) > 0:
                labeled_cols.append(x)
```

```python
            x += 1

        if len(labeled_cols) >= 2:
            # Find largest gap between consecutive labeled columns
            best_gap_idx = -1
            best_gap_size = 0
            i = 0
            while i < len(labeled_cols) - 1:
                gap = labeled_cols[i + 1] - labeled_cols[i]
                if gap > 1 and gap > best_gap_size:
                    best_gap_size = gap
                    best_gap_idx = i
                i += 1

            if best_gap_idx >= 0:
                strip_left[y] = labeled_cols[best_gap_idx]
                strip_right[y] = labeled_cols[best_gap_idx + 1]

        y += 1

    return strip_left, strip_right

def compute_conc_row_centers(ip_value, conc_values, w, h,
                             tol_log=CONC_MATCH_LOG_TOLERANCE):
    """Precompute median row for each concentration value using log-space
matching.

    Returns dict: {conc_value: median_row_px}
    """
    # Precompute log values for valid concentrations
    log_concs = []
    valid_concs = []
    for cv in conc_values:
        if cv <= 0:
            continue
        try:
            log_cv = math.log(cv)
            log_concs.append(log_cv)
            valid_concs.append(cv)
        except (ValueError, OverflowError):
            continue

    if not valid_concs:
        return {}

    # Collect rows for each concentration
    conc_rows = {}
```

```
    for cv in valid_concs:
        conc_rows[cv] = []

    y = 0
    while y < h:
        x = 0
        while x < w:
            val = ip_value.getf(x, y)
            if val > 0:
                log_val = math.log(max(val, 1e-30))
                for i in range(len(valid_concs)):
                    if abs(log_val - log_concs[i]) < tol_log:
                        conc_rows[valid_concs[i]].append(y)
            x += 1
        y += 1

    # Compute median row for each concentration
    result = {}
    for cv in valid_concs:
        rows = conc_rows[cv]
        if len(rows) > 0:
            rows_sorted = sorted(rows)
            mid = len(rows_sorted) // 2
            if len(rows_sorted) % 2 == 0:
                median_row = (rows_sorted[mid - 1] + rows_sorted[mid]) / 2.0
            else:
                median_row = float(rows_sorted[mid])
            result[cv] = median_row

    return result

def build_row_to_conc(conc_row_centers, h):
    """Build a row-to-concentration lookup using nearest center row.

    For each row, assigns the concentration whose center row is closest.
    This is the Jython equivalent of numpy's searchsorted approach.

    Returns a list of length h, where each element is the concentration
    for that row, or 0.0 if no centers are available.
    """
    if not conc_row_centers:
        return [0.0] * h

    # Sort centers by row
    sorted_centers = sorted(
        [(row, cv) for cv, row in conc_row_centers.items()],
        key=lambda x: x[0],
```

```
    )
    center_rows = [r for r, _ in sorted_centers]
    center_concs = [cv for _, cv in sorted_centers]
    n_centers = len(center_rows)

    row_to_conc = [0.0] * h

    y = 0
    while y < h:
        # Binary search for insertion position
        lo = 0
        hi = n_centers
        while lo < hi:
            mid = (lo + hi) // 2
            if center_rows[mid] < y:
                lo = mid + 1
            else:
                hi = mid
        pos = lo

        # Clip to [1, n_centers-1] (matches numpy clip(insert_pos, 1, len-1))
        if pos < 1:
            pos = 1
        if pos > n_centers - 1:
            pos = n_centers - 1

        # Compare distances to left and right centers
        dist_left = abs(y - center_rows[pos - 1])
        dist_right = abs(y - center_rows[pos])

        if dist_left <= dist_right:
            row_to_conc[y] = center_concs[pos - 1]
        else:
            row_to_conc[y] = center_concs[pos]

        y += 1

    return row_to_conc

#
==========================================================================
# Step 4: Wide/Narrow Classification (Ring Matrix Approach)
#
==========================================================================

def extract_ring_matrix(pixel_data, strip_sequence_values,
```

```
                        max_width_mm=EXTENDED_MAX_RING_WIDTH_MM,
                        ring_width_mm=RING_WIDTH_MM):
    """Extract ring intensity matrix from pixel data using pure Jython dicts.

    Bins pixels by distance from strip edge per concentration, producing
    a 2D matrix (ring x concentration -> mean intensity).

    Args:
        pixel_data: List of (distance_mm, raw_intensity, concentration)
tuples
        strip_sequence_values: List of strip concentration values
        max_width_mm: Maximum distance to consider for rings
        ring_width_mm: Width of each ring bin in mm

    Returns:
        (ring_matrix, ring_centers, ring_boundaries, conc_values)
        ring_matrix: dict of (ring_idx, conc_idx) -> mean intensity
        ring_centers: list of ring center distances in mm
        ring_boundaries: list of (inner_mm, outer_mm) tuples
        conc_values: list of concentration values (sorted descending)
    """
    n_rings = int(round(max_width_mm / ring_width_mm))

    ring_boundaries = []
    ring_centers = []
    i = 0
    while i < n_rings:
        inner = round(i * ring_width_mm, 2)
        outer = round((i + 1) * ring_width_mm, 2)
        ring_boundaries.append((inner, outer))
        ring_centers.append(round((inner + outer) / 2.0, 2))
        i += 1

    if len(pixel_data) == 0:
        return {}, ring_centers, ring_boundaries, []

    # Build positive strip sequence for log-space matching
    sseq_pos = []
    i = 0
    while i < len(strip_sequence_values):
        sv = strip_sequence_values[i]
        if sv > 0:
            sseq_pos.append(sv)
        i += 1

    if len(sseq_pos) == 0:
        return {}, ring_centers, ring_boundaries, []
```

```
    sseq_log = []
    i = 0
    while i < len(sseq_pos):
        sseq_log.append(math.log(sseq_pos[i]))
        i += 1

    # Concentration values sorted descending (matching simplified_mic.py
convention)
    conc_values = sorted(strip_sequence_values, reverse=True)
    conc_to_col = {}
    i = 0
    while i < len(conc_values):
        conc_to_col[conc_values[i]] = i
        i += 1

    # Accumulate sums and counts per (ring_idx, conc_col)
    ring_sums = {}   # (ring_idx, col_idx) -> sum of intensities
    ring_counts = {} # (ring_idx, col_idx) -> count of pixels

    i = 0
    while i < len(pixel_data):
        dist, raw_val, conc = pixel_data[i]

        # Compute ring index
        if dist < 0:
            i += 1
            continue
        ring_idx = int(dist / ring_width_mm)
        if ring_idx < 0 or ring_idx >= n_rings:
            i += 1
            continue

        # Match concentration to strip sequence value in log space
        if conc <= 0:
            i += 1
            continue

        try:
            conc_log = math.log(float(conc))
        except (ValueError, OverflowError):
            i += 1
            continue

        best_j = -1
        best_diff = float("inf")
        j = 0
        while j < len(sseq_log):
            diff = abs(conc_log - sseq_log[j])
```

```python
            if diff < best_diff:
                best_diff = diff
                best_j = j
            j += 1

        if best_j < 0 or best_diff >= CONC_MATCH_LOG_TOLERANCE:
            i += 1
            continue

        matched_sv = sseq_pos[best_j]
        if matched_sv not in conc_to_col:
            i += 1
            continue

        col_idx = conc_to_col[matched_sv]
        key = (ring_idx, col_idx)

        if key not in ring_sums:
            ring_sums[key] = 0.0
            ring_counts[key] = 0
        ring_sums[key] += raw_val
        ring_counts[key] += 1

        i += 1

    # Compute mean intensities
    ring_matrix = {}
    keys = list(ring_sums.keys())
    i = 0
    while i < len(keys):
        key = keys[i]
        if ring_counts[key] > 0:
            ring_matrix[key] = ring_sums[key] / float(ring_counts[key])
        i += 1

    return ring_matrix, ring_centers, ring_boundaries, conc_values

def _get_ring_value(ring_matrix, ring_idx, col_idx):
    """Get value from ring matrix dict, returns None if not present."""
    key = (ring_idx, col_idx)
    if key in ring_matrix:
        return ring_matrix[key]
    return None

def compute_global_lawn_intensity(ring_matrix, ring_centers, conc_values,
n_lowest=3):
```

```
    """Estimate global lawn intensity from outermost rings of lowest
concentrations.

    The lowest concentrations have the largest ZOI, so their outermost rings
    represent the lawn (uninhibited growth) intensity.

    Returns median of the outermost valid ring values from the n_lowest
concentrations.
    """
    n_rings = len(ring_centers)
    if n_rings == 0 or len(conc_values) == 0:
        return None

    lawn_intensities = []
    # conc_values is sorted descending, so first n_lowest are the highest
concentrations
    n_cols_to_use = min(n_lowest, len(conc_values))
    col_idx = 0
    while col_idx < n_cols_to_use:
        # Scan from outermost ring inward to find the outermost valid value
        ring_idx = n_rings - 1
        while ring_idx >= 0:
            val = _get_ring_value(ring_matrix, ring_idx, col_idx)
            if val is not None:
                lawn_intensities.append(val)
                break
            ring_idx -= 1
        col_idx += 1

    if len(lawn_intensities) == 0:
        return None

    # Return median (pure Python implementation)
    lawn_intensities.sort()
    n = len(lawn_intensities)
    if n % 2 == 1:
        return lawn_intensities[n // 2]
    else:
        return (lawn_intensities[n // 2 - 1] + lawn_intensities[n // 2]) /
2.0

def compute_global_zoi_min(ring_matrix, conc_values, percentile=5.0):
    """Estimate global ZOI minimum intensity from all ring data.

    Uses the given percentile of all valid ring matrix values.
    Implements linear interpolation matching NumPy's percentile() behavior.
    """
```

```
    # Collect all valid values from ring matrix
    all_values = []
    keys = list(ring_matrix.keys())
    i = 0
    while i < len(keys):
        val = ring_matrix[keys[i]]
        if val is not None:
            all_values.append(val)
        i += 1

    if len(all_values) == 0:
        return None

    all_values.sort()
    n = len(all_values)

    # NumPy-style linear interpolation
    virtual_idx = percentile / 100.0 * (n - 1)
    lower_idx = int(virtual_idx)
    upper_idx = lower_idx + 1
    if upper_idx >= n:
        return all_values[-1]
    fraction = virtual_idx - lower_idx
    return all_values[lower_idx] + fraction * (all_values[upper_idx] -
all_values[lower_idx])

def detect_zoi_boundary_normalized_threshold(ring_matrix, ring_centers,
conc_values,
                                              lawn_intensity,
zoi_min_global=None,
                                              lawn_fraction=LAWN_FRACTION):
    """Find ZOI boundary per concentration using normalized threshold.

    threshold = zoi_min + lawn_fraction * (lawn_intensity - zoi_min)

    Scans from innermost ring outward; the first ring whose intensity
    is >= threshold marks the ZOI boundary for that concentration.

    Returns dict: {concentration_value: boundary_distance_mm}
    """
    n_rings = len(ring_centers)
    if n_rings == 0 or lawn_intensity is None or lawn_intensity <= 0:
        return {}

    boundaries = {}

    col_idx = 0
```

```python
    while col_idx < len(conc_values):
        conc = conc_values[col_idx]

        # Get column values for this concentration
        col_values = []
        ring_idx = 0
        while ring_idx < n_rings:
            val = _get_ring_value(ring_matrix, ring_idx, col_idx)
            col_values.append(val)
            ring_idx += 1

        # Check if any valid values exist
        has_valid = False
        j = 0
        while j < len(col_values):
            if col_values[j] is not None:
                has_valid = True
                break
            j += 1

        if not has_valid:
            col_idx += 1
            continue

        # Determine zoi_min for this concentration
        if zoi_min_global is not None:
            zoi_min = zoi_min_global
        else:
            # Use minimum of valid values in this column
            valid_vals = []
            j = 0
            while j < len(col_values):
                if col_values[j] is not None:
                    valid_vals.append(col_values[j])
                j += 1
            if len(valid_vals) == 0:
                col_idx += 1
                continue
            zoi_min = min(valid_vals)

        # Check innermost intensity for relative drop
        innermost_intensity = col_values[0]  # may be None
        if innermost_intensity is None:
            # Find first valid value from innermost
            j = 0
            while j < len(col_values):
                if col_values[j] is not None:
                    innermost_intensity = col_values[j]
```

```
                break
            j += 1

        if innermost_intensity is not None:
            dynamic_range = lawn_intensity - zoi_min
            if dynamic_range > 0:
                relative_drop = (lawn_intensity - innermost_intensity) /
dynamic_range
                if relative_drop < 0.1:
                    col_idx += 1
                    continue

        # Compute threshold
        threshold_val = zoi_min + lawn_fraction * (lawn_intensity - zoi_min)

        # Scan from innermost ring outward, find first ring above threshold
        boundary_ring = None
        ring_idx = 0
        while ring_idx < n_rings:
            val = col_values[ring_idx]
            if val is not None and val >= threshold_val:
                boundary_ring = ring_idx
                break
            ring_idx += 1

        if boundary_ring is not None:
            boundaries[conc] = ring_centers[boundary_ring]

        col_idx += 1

    return boundaries

def enforce_monotonicity(detectable, strip_sequence_values):
    """Enforce that boundary distances are non-decreasing with concentration.

    As concentration increases, the ZOI boundary distance should not
decrease.
    """
    if len(detectable) < 2:
        return detectable

    sorted_concs = sorted(detectable.keys())

    running_max = 0.0
    i = 0
    while i < len(sorted_concs):
        c = sorted_concs[i]
```

```
        if detectable[c] < running_max:
            detectable[c] = running_max
        else:
            running_max = detectable[c]
        i += 1

    return detectable

def derive_key_concentrations_simple(boundary_distances,
strip_sequence_values):
    """Derive c_high, d_high from boundary distances.

    c_high is the concentration with the maximum boundary distance.
    d_high is that maximum boundary distance.

    Returns (c_high, d_high).
    """
    if not boundary_distances:
        return None, None

    c_high = max(boundary_distances, key=boundary_distances.get)
    d_high = boundary_distances[c_high]

    return c_high, d_high

def classify_wide_narrow(pixel_data, strip_sequence_values):
    """Classify image as wide or narrow ZOI and select corridor width.

    Algorithm:
    1. Extract ring matrix from pixel data
    2. Compute global lawn intensity and ZOI minimum
    3. Detect ZOI boundary per concentration
    4. Filter boundaries < MIN_INHIBITION_THRESHOLD_MM
    5. Enforce monotonicity
    6. Derive d_high = max(boundary_distances)
    7. Classify: d_high <= D_HIGH_THRESHOLD -> narrow, else -> wide

    Returns:
        (is_narrow, d_high, corridor_width_mm, c_high)
    """
    # Extract ring matrix
    ring_matrix, ring_centers, ring_boundaries, conc_values = \
        extract_ring_matrix(pixel_data, strip_sequence_values)

    if len(conc_values) == 0 or len(ring_centers) == 0:
        return False, None, CORRIDOR_WIDTH_WIDE_MM, None
```

```
    # Compute lawn intensity and ZOI minimum
    lawn_intensity = compute_global_lawn_intensity(ring_matrix, ring_centers,
conc_values)
    zoi_min = compute_global_zoi_min(ring_matrix, conc_values)

    if lawn_intensity is None or zoi_min is None:
        return False, None, CORRIDOR_WIDTH_WIDE_MM, None

    # Detect ZOI boundaries
    boundaries = detect_zoi_boundary_normalized_threshold(
        ring_matrix, ring_centers, conc_values, lawn_intensity, zoi_min)

    # Filter boundaries below minimum threshold
    filtered = {}
    keys = list(boundaries.keys())
    i = 0
    while i < len(keys):
        c = keys[i]
        d = boundaries[c]
        if d >= MIN_INHIBITION_THRESHOLD_MM:
            filtered[c] = d
        i += 1

    # Enforce monotonicity
    filtered = enforce_monotonicity(filtered, strip_sequence_values)

    # Derive key concentrations
    c_high, d_high = derive_key_concentrations_simple(filtered,
strip_sequence_values)

    if d_high is None:
        return False, None, CORRIDOR_WIDTH_WIDE_MM, None

    # Classify
    is_narrow = d_high <= D_HIGH_THRESHOLD
    corridor_width = CORRIDOR_WIDTH_NARROW_MM if is_narrow else
CORRIDOR_WIDTH_WIDE_MM

    return is_narrow, d_high, corridor_width, c_high

#
============================================================================
# Step 5: Corridor Measurement Extraction
#
============================================================================
```

```
def extract_measurements_for_corridor(pixel_data, corridor_width_mm,
strip_sequence_values):
    """Extract intensity measurements for a given corridor width.

    For each concentration in strip_sequence_values (sorted descending):
    - Select pixels where conc matches (log-space tolerance) AND 0 < dist <=
corridor_width_mm
    - Compute mean, min, max intensity and pixel count

    Args:
        pixel_data: List of (distance_mm, raw_intensity, concentration)
tuples
        corridor_width_mm: Maximum distance from strip edge in mm
        strip_sequence_values: List of strip concentration values

    Returns:
        List of measurement dicts sorted by s_value descending, each with:
        s_value, log_s_value, mean, min, max, n_pixels
    """
    # Filter pixels within corridor width and group by concentration
    conc_intensities = {}  # concentration -> list of intensities

    i = 0
    while i < len(pixel_data):
        dist, raw_val, conc = pixel_data[i]
        if dist > 0 and dist <= corridor_width_mm:
            if conc not in conc_intensities:
                conc_intensities[conc] = []
            conc_intensities[conc].append(raw_val)
        i += 1

    if len(conc_intensities) == 0:
        return []

    # Match each unique concentration to nearest strip sequence value in log
space.
    # 1:1 invariant: each concentration maps to exactly one strip value, so
no
    # many-to-one merging is needed — each strip value's pixels come from one
    # concentration. Log-space matching is still needed because
floating-point
    # concentrations in the Value image may not exactly equal strip values.
    conc_to_strip = {}
    conc_keys = list(conc_intensities.keys())
    i = 0
    while i < len(conc_keys):
        uc = conc_keys[i]
```

```python
        best_sv = None
        best_diff = float("inf")
        j = 0
        while j < len(strip_sequence_values):
            sv = strip_sequence_values[j]
            if sv > 0:
                try:
                    diff = abs(math.log(float(uc)) - math.log(sv))
                except (ValueError, OverflowError):
                    diff = float("inf")
                if diff < best_diff:
                    best_diff = diff
                    best_sv = sv
            j += 1
        if best_sv is not None and best_diff < CONC_MATCH_LOG_TOLERANCE:
            conc_to_strip[uc] = best_sv
        i += 1

    if len(conc_to_strip) == 0:
        return []

    # Compute statistics per strip value (1:1 mapping — no merging needed)
    measurements = []
    i = 0
    while i < len(conc_keys):
        uc = conc_keys[i]
        if uc in conc_to_strip:
            sv = conc_to_strip[uc]
            intensities = conc_intensities[uc]
            if len(intensities) > 0:
                mean_val = sum(intensities) / float(len(intensities))
                min_val = min(intensities)
                max_val = max(intensities)
                measurements.append({
                    "s_value": float(sv),
                    "log_s_value": math.log(float(sv)),
                    "mean": mean_val,
                    "min": min_val,
                    "max": max_val,
                    "n_pixels": len(intensities),
                })
        i += 1

    # Sort by concentration descending (high to low)
    measurements.sort(key=lambda m: m["s_value"], reverse=True)

    return measurements
```

```
#
==============================================================================
# Step 6: MIC Finding (Approach 3: Threshold-based)
#
==============================================================================

def find_nearest_strip_value(mic_raw, strip_sequence_values):
    """Find the nearest strip value to mic_raw in log space.

    Returns the nearest strip concentration value, or None if mic_raw is
invalid.
    """
    if mic_raw is None or mic_raw <= 0 or math.isnan(mic_raw) or
math.isinf(mic_raw):
        return None
    sorted_strips = sorted(strip_sequence_values)
    log_mic = math.log(mic_raw)
    best_sv = None
    best_diff = float("inf")
    i = 0
    while i < len(sorted_strips):
        sv = sorted_strips[i]
        if sv > 0:
            try:
                diff = abs(math.log(sv) - log_mic)
            except (ValueError, OverflowError):
                diff = float("inf")
            if diff < best_diff:
                best_diff = diff
                best_sv = sv
        i += 1
    return best_sv

def approach3_mic(measurements, threshold, strip_sequence_values):
    """Compute MIC using intensity threshold on raw measurements.

    Algorithm (approach3 -- mean-of-3 normalization):
    1. background = mean of 3 highest-concentration measurements
    2. max_growth = mean of 3 lowest-concentration measurements
    3. excess_growth = (mean - background) / (max_growth - background)
    4. Scan from high to low concentration, find first eg >= threshold
    5. Interpolate in log-space between adjacent concentrations
    6. Snap to nearest strip value

    Args:
```

```
        measurements: List of measurement dicts sorted by s_value DESCENDING
        threshold: MIC threshold (e.g. 0.18)
        strip_sequence_values: List of strip concentration values

    Returns:
        dict with keys:
            mic_snapped, mic_raw, excess_growths, crossing_idx,
            background, max_growth, growth_range_ratio, threshold_intensity
        or None if computation fails entirely
    """
    if len(measurements) < 4:
        return None

    # measurements sorted by s_value DESCENDING (high to low concentration)
    n_right = min(N_REFERENCE, len(measurements))
    n_left = min(N_REFERENCE, len(measurements))

    background = sum(m["mean"] for m in measurements[:n_right]) / n_right
    max_growth = sum(m["mean"] for m in measurements[-n_left:]) / n_left

    growth_range = max_growth - background
    growth_range_ratio = growth_range / ((max_growth + background) / 2) if
(max_growth + background) > 0 else 0.0
    if growth_range <= 0:
        return None

    # Compute excess growth for each concentration
    excess_growths = []
    i = 0
    while i < len(measurements):
        eg = (measurements[i]["mean"] - background) / growth_range
        excess_growths.append(eg)
        i += 1

    # Threshold intensity = background + threshold * growth_range
    threshold_intensity = background + threshold * growth_range

    # Scan from high conc (index 0) to low conc (last index)
    # Find first index where excess_growth >= threshold
    crossing_idx = None
    i = 0
    while i < len(excess_growths):
        if excess_growths[i] >= threshold:
            crossing_idx = i
            break
        i += 1

    mic_raw = None
```

```
    if crossing_idx is None:
        # No concentration exceeds threshold -> MIC at lowest concentration
        mic_raw = measurements[-1]["s_value"]
    elif crossing_idx == 0:
        # Even highest concentration exceeds threshold -> MIC at highest
concentration
        mic_raw = measurements[0]["s_value"]
    else:
        # Interpolate between crossing_idx-1 and crossing_idx in log-space
        eg_prev = excess_growths[crossing_idx - 1]
        eg_curr = excess_growths[crossing_idx]
        if abs(eg_curr - eg_prev) < 1e-15:
            log_mic = measurements[crossing_idx]["log_s_value"]
        else:
            t = (threshold - eg_prev) / (eg_curr - eg_prev)
            log_mic = (measurements[crossing_idx - 1]["log_s_value"] +
                       t * (measurements[crossing_idx]["log_s_value"] -
                            measurements[crossing_idx - 1]["log_s_value"]))
        try:
            mic_raw = math.exp(log_mic)
        except (OverflowError, ValueError):
            return None

    if mic_raw is None or mic_raw <= 0 or math.isnan(mic_raw) or
math.isinf(mic_raw):
        return None

    # Snap to nearest strip value in log-space
    mic_snapped = find_nearest_strip_value(mic_raw, strip_sequence_values)

    return {
        "mic_snapped": mic_snapped,
        "mic_raw": mic_raw,
        "excess_growths": excess_growths,
        "crossing_idx": crossing_idx,
        "background": background,
        "max_growth": max_growth,
        "growth_range_ratio": growth_range_ratio,
        "threshold_intensity": threshold_intensity,
    }

#
============================================================================
# Step 7: N/A Detection (Threshold-based, Fitting-free)
#
============================================================================
```

```
def compute_hi_conc_cv(measurements, n_reference=N_REFERENCE):
    """Compute coefficient of variation of the top-N highest-concentration
measurements.

    A high CV at the highest concentrations indicates variable growth inside
the
    ZOI -- some areas show growth while others show inhibition. This is a
strong
    signal that the MIC reading is unreliable and should be reported as N/A.

    Args:
        measurements: List of measurement dicts sorted by s_value DESCENDING
                      (as returned by extract_measurements_for_corridor)
        n_reference: Number of highest-concentration measurements to use

    Returns:
        CV value (float) or None if computation fails
    """
    n = min(n_reference, len(measurements))
    if n < 2:
        return None

    hi_means = []
    i = 0
    while i < n:
        hi_means.append(measurements[i]["mean"])
        i += 1

    mean_val = sum(hi_means) / n
    if mean_val <= 0:
        return None

    variance = 0.0
    i = 0
    while i < n:
        diff = hi_means[i] - mean_val
        variance += diff * diff
        i += 1
    variance /= n

    return math.sqrt(variance) / mean_val

def detect_na_threshold(growth_range_ratio, max_boundary_mm,
                        ref_growth_range_ratio, hi_conc_cv=None):
    """Threshold-based N/A detection (no logistic fitting).
```

```
    Uses 3 OR-combined rules:
    1. min(growth_range_ratio, ref_growth_range_ratio) < 0.44 -> low growth
variation
       at either corridor width (merges former separate Rules 1 & 3)
    2. max_boundary_mm < 1.50 -> very small/no ZOI boundary
    3. hi_conc_cv > 0.15 -> variable growth inside ZOI (growth at high
concentrations)

    Args:
        growth_range_ratio: from approach3_mic() on the wide corridor
        max_boundary_mm: d_high from classify_wide_narrow() (max ZOI boundary
distance).
                         None if no boundaries detected (treated as 0.0 ->
N/A).
        ref_growth_range_ratio: from approach3_mic() on the narrow corridor
        hi_conc_cv: coefficient of variation of top-3 highest-concentration
measurements
                    from the wide corridor. None if not computable.

    Returns (is_na, reason_string).
    """
    reasons = []

    # Rule 1: low growth variation at either corridor width
    min_grr = min(growth_range_ratio, ref_growth_range_ratio)
    if min_grr < NA_MIN_GROWTH_RANGE_RATIO_THRESHOLD:
        reasons.append(
            "low_growth_ratio=" + str(round(min_grr, 3)) +
            "<" + str(NA_MIN_GROWTH_RANGE_RATIO_THRESHOLD) +
            "(grr=" + str(round(growth_range_ratio, 3)) +
            ",ref=" + str(round(ref_growth_range_ratio, 3)) + ")")

    # Rule 2: very small/no ZOI boundary
    # Handle d_high = None: no boundaries detected -> treat as 0.0 (triggers
N/A)
    boundary_val = max_boundary_mm if max_boundary_mm is not None else 0.0
    if boundary_val < NA_MAX_BOUNDARY_MM_THRESHOLD:
        reasons.append(
            "small_zoi=" + str(round(boundary_val, 2)) +
            "mm<" + str(NA_MAX_BOUNDARY_MM_THRESHOLD) + "mm")

    # Rule 3: high CV of top-3 highest-concentration measurements -> growth
inside ZOI
    if hi_conc_cv is not None and hi_conc_cv > NA_HI_CONC_CV_THRESHOLD:
        reasons.append(
            "hi_conc_cv=" + str(round(hi_conc_cv, 3)) +
            ">" + str(NA_HI_CONC_CV_THRESHOLD))
```

```
    is_na = len(reasons) > 0
    reason = "; ".join(reasons) if reasons else "not_na"

    return is_na, reason

#
==========================================================================
# Step 8: MIC String Formatting
#
==========================================================================

def format_mic_string(mic_value, strip_sequence_text):
    """Format MIC value using original strip sequence token.

    Since mic_value is already snapped to a strip sequence value earlier in
    the algorithm (by find_nearest_strip_value), a direct dict lookup
suffices.
    A float-tolerance fallback handles floating-point edge cases.
    """
    if mic_value is None or mic_value <= 0:
        return "N/A"

    pairs = parse_strip_sequence_pairs(strip_sequence_text)
    if not pairs:
        if mic_value == int(mic_value) and abs(mic_value) < 1e15:
            return str(int(mic_value))
        return str(mic_value)

    # Direct lookup: mic_value should exactly match one of the pair values
    value_to_token = {v: token for v, token in pairs}
    if mic_value in value_to_token:
        return value_to_token[mic_value]

    # Fallback: float comparison with tolerance for floating-point edge cases
    for v, token in pairs:
        if abs(v - mic_value) < 1e-6:
            return token

    return str(mic_value)

#
==========================================================================
# Main Data Batch Processing
#
==========================================================================
```

```
input_slot_dist = input_slot_map["DistanceMap"]
input_slot_labels = input_slot_map["Labels"]
input_slot_raw = input_slot_map["Raw"]
input_slot_value = input_slot_map["Value"]
output_slot_measurements = output_slot_map["Measurements"]
output_slot_mask = output_slot_map["MICMask"]

try:
    # ------------------------------------------------------------------
    # Step 1: Read text annotations & load images
    # ------------------------------------------------------------------
    strip_sequence_text = ""
    try:
        strip_sequence_text =
data_batch.getMergedTextAnnotation("StripSequence").getValue()
    except Exception:
        pass

    pixel_size_mm = -1
    try:
        ps_text =
data_batch.getMergedTextAnnotation(PIXEL_SIZE_ANNOTATION_NAME).getValue()
        pixel_size_mm = float(ps_text)
        pixel_size_mm = pixel_size_mm / IMAGE_SCALE_FACTOR
    except Exception:
        pass
    if pixel_size_mm <= 0:
        progress_info.log("ERROR: " + PIXEL_SIZE_ANNOTATION_NAME + "
annotation is missing or invalid. "
                          + "Pixel size must be provided via JIPipe
annotation.")
        raise ValueError(PIXEL_SIZE_ANNOTATION_NAME + " annotation is missing
or invalid. "
                         + "Using a default pixel size would produce
incorrect results.")

    progress_info.log("Parsing strip sequence")
    strip_sequence_values = parse_strip_sequence(strip_sequence_text)

    # Load images via ImageJ API
    ip_dist = data_batch.getInputData(input_slot_dist, ImagePlusData,
progress_info).getImage().getProcessor()
    ip_labels = data_batch.getInputData(input_slot_labels, ImagePlusData,
progress_info).getImage().getProcessor()
    ip_raw = data_batch.getInputData(input_slot_raw, ImagePlusData,
progress_info).getImage().getProcessor()
```

```
    ip_value = data_batch.getInputData(input_slot_value, ImagePlusData,
progress_info).getImage().getProcessor()

    # Apply image scaling for enhanced performance
    ip_dist = applyScaling(ip_dist, 1)    # BILINEAR - continuous distance
values
    ip_labels = applyScaling(ip_labels, 0) # NEAREST_NEIGHBOR - discrete
label IDs
    ip_raw = applyScaling(ip_raw, 1)       # BILINEAR - continuous intensity
values
    ip_value = applyScaling(ip_value, 0)   # NEAREST_NEIGHBOR - discrete
concentration values

    w = ip_labels.getWidth()
    h = ip_labels.getHeight()

    # ----------------------------------------------------------------
    # Step 2: Detect inverted contrast
    # ----------------------------------------------------------------
    if ENABLE_INVERSE_DETECTION:
        inverted = detect_inverted_contrast(ip_raw, ip_value, ip_labels, w,
h)
    else:
        progress_info.log("Inverse contrast detection is disabled")
        inverted = False

    # Determine max raw value for potential inversion
    raw_max = 0.0
    if inverted:
        try:
            raw_max = float(ip_raw.getStats().max)
        except Exception:
            # Fallback: compute max manually
            raw_max = 0.0
            y = 0
            while y < h:
                x = 0
                while x < w:
                    v = ip_raw.getf(x, y)
                    if v > raw_max:
                        raw_max = v
                    x += 1
                y += 1

    # ----------------------------------------------------------------
    # Step 3: Collect pixel data (extended — includes growth/lawn pixels)
    # Pre-collect all (distance, intensity, concentration) tuples.
    # Distance comes from the DistanceMap input (not computed internally).
```

```
    # Includes ALL pixels where dist > 0 (off the strip), not just labeled.
    # Unlabeled pixels get their concentration from the nearest row center.
    # ---------------------------------------------------------------
    progress_info.log("Computing strip edges and concentration row centers")
    strip_left, strip_right = find_strip_edges(ip_labels, w, h)
    conc_row_centers = compute_conc_row_centers(ip_value,
strip_sequence_values, w, h)
    row_to_conc = build_row_to_conc(conc_row_centers, h)

    # Build valid row mask: only rows where strip edges were found
    valid_row = [False] * h
    y = 0
    while y < h:
        if strip_left[y] is not None and strip_right[y] is not None:
            valid_row[y] = True
        y += 1

    n_valid_rows = 0
    y = 0
    while y < h:
        if valid_row[y]:
            n_valid_rows += 1
        y += 1
    progress_info.log("Valid rows (with strip edges): " + str(n_valid_rows) +
"/" + str(h))
    progress_info.log("Concentration row centers: " +
str(len(conc_row_centers)))

    progress_info.log("Collecting [distance, intensity, concentration] tuples
(extended)")
    pixel_data = []  # list of (distance_mm, raw_intensity, concentration)
    max_dist_mm = EXTENDED_MAX_RING_WIDTH_MM

    y = 0
    while y < h:
        if not valid_row[y]:
            y += 1
            continue
        x = 0
        while x < w:
            dist = ip_dist.getf(x, y)
            if dist > 0 and dist <= max_dist_mm:
                raw_val = ip_raw.getf(x, y)
                # Skip background (black) pixels outside plate boundary
                if raw_val <= 0:
                    x += 1
                    continue
                if inverted:
```

```
                    raw_val = raw_max - raw_val
                # Concentration: labeled pixels use value_f, unlabeled use
row_to_conc
                label_val = ip_labels.getf(x, y)
                if label_val > 0:
                    conc_val = ip_value.getf(x, y)
                    conc_rounded = round(conc_val, 6)
                else:
                    conc_rounded = round(row_to_conc[y], 6)
                if conc_rounded > 0:
                    pixel_data.append((dist, raw_val, conc_rounded))
            x += 1
        y += 1

    progress_info.log("Pixels collected (extended): " + str(len(pixel_data)))
    progress_info.log("Inverted contrast: " + str(inverted))

    # ----------------------------------------------------------------
    # Step 4: Wide/Narrow Classification
    # ----------------------------------------------------------------
    is_narrow, d_high, corridor_width_mm, c_high = classify_wide_narrow(
        pixel_data, strip_sequence_values)

    classification_str = "NARROW" if is_narrow else "WIDE"
    progress_info.log("Classification: " + classification_str +
                      ", d_high=" + str(d_high) +
                      ", corridor_width=" + str(corridor_width_mm) + "mm")

    # ----------------------------------------------------------------
    # Step 5: Extract corridor measurements
    # Always extract at both wide and narrow corridors for N/A detection.
    # The selected corridor is used for MIC computation.
    # ----------------------------------------------------------------
    measurements_wide = extract_measurements_for_corridor(
        pixel_data, CORRIDOR_WIDTH_WIDE_MM, strip_sequence_values)

    measurements_narrow = extract_measurements_for_corridor(
        pixel_data, CORRIDOR_WIDTH_NARROW_MM, strip_sequence_values)

    # Select measurements based on classification
    if is_narrow:
        measurements = measurements_narrow
    else:
        measurements = measurements_wide

    # Free pixel data memory (no longer needed after measurement extraction)
    pixel_data = None
```

```
    progress_info.log("Measurements at " + str(corridor_width_mm) + "mm: " +
                      str(len(measurements)) + " concentrations")
    progress_info.log("Measurements at " + str(CORRIDOR_WIDTH_WIDE_MM) + "mm:
" +
                      str(len(measurements_wide)) + " concentrations")
    progress_info.log("Measurements at " + str(CORRIDOR_WIDTH_NARROW_MM) +
"mm: " +
                      str(len(measurements_narrow)) + " concentrations")

    # ----------------------------------------------------------------
    # Early check: too few measurements for reliable MIC finding
    # ----------------------------------------------------------------
    if len(measurements) < 4 and ENABLE_NA_DETECTION:
        is_na = True
        na_reason = "too_few_measurements ({})".format(len(measurements))
        mic_result = None
        mic_snapped = None
        mic_raw = None
        growth_range_ratio = 0.0
        progress_info.log("Too few measurements (" + str(len(measurements)) +
"), skipping MIC finding")
    else:
        # ----------------------------------------------------------------
        # Step 6: Find MIC using approach3 (threshold-based)
        # ----------------------------------------------------------------
        mic_result = approach3_mic(measurements, DEFAULT_MIC_THRESHOLD,
strip_sequence_values)

        if mic_result is not None:
            growth_range_ratio = mic_result["growth_range_ratio"]
            mic_snapped = mic_result["mic_snapped"]
            mic_raw = mic_result["mic_raw"]
            progress_info.log("MIC: raw=" + str(round(mic_raw, 4)) +
                              ", snapped=" + str(mic_snapped) +
                              ", growth_range_ratio=" +
str(round(growth_range_ratio, 3)))
        else:
            growth_range_ratio = 0.0
            mic_snapped = None
            mic_raw = None
            progress_info.log("MIC: approach3 failed (insufficient data or no
growth range)")

        # ----------------------------------------------------------------
        # Step 7: N/A Detection (3 OR-combined threshold rules)
        # Need growth_range_ratio from both wide and narrow corridors.
        # ----------------------------------------------------------------
```

```
        # growth_range_ratio from wide corridor (N/A rule 1)
        if not is_narrow and mic_result is not None:
            # Already computed for wide images
            growth_range_ratio_wide = growth_range_ratio
        else:
            # Need to compute for narrow images (or when mic_result failed)
            mic_result_wide = approach3_mic(measurements_wide,
DEFAULT_MIC_THRESHOLD, strip_sequence_values)
            growth_range_ratio_wide = mic_result_wide["growth_range_ratio"]
if mic_result_wide is not None else 0.0

        # growth_range_ratio from narrow corridor (N/A rule 3)
        if is_narrow and mic_result is not None:
            # Already computed for narrow images
            growth_range_ratio_narrow = growth_range_ratio
        else:
            # Need to compute for wide images (or when mic_result failed)
            mic_result_narrow = approach3_mic(measurements_narrow,
DEFAULT_MIC_THRESHOLD, strip_sequence_values)
            growth_range_ratio_narrow =
mic_result_narrow["growth_range_ratio"] if mic_result_narrow is not None else
0.0

        # Rule 3: CV of top-3 highest-concentration measurements (growth
inside ZOI)
        hi_conc_cv = compute_hi_conc_cv(measurements_wide)

        if ENABLE_NA_DETECTION:
            is_na, na_reason = detect_na_threshold(
                growth_range_ratio_wide, d_high, growth_range_ratio_narrow,
hi_conc_cv)
        else:
            is_na = False
            na_reason = "na_detection_disabled"

        min_grr = min(growth_range_ratio_wide, growth_range_ratio_narrow)
        cv_str = str(round(hi_conc_cv, 3)) if hi_conc_cv is not None else
"N/A"
        progress_info.log("N/A detection: is_na=" + str(is_na) +
                          ", min_grr=" + str(round(min_grr, 3)) +
                          "(wide=" + str(round(growth_range_ratio_wide, 3)) +
                          ",narrow=" + str(round(growth_range_ratio_narrow,
3)) + ")" +
                          ", d_high=" + str(d_high) +
                          ", hi_conc_cv=" + cv_str +
                          ", reason=" + str(na_reason))

    # ------------------------------------------------------------------
```

```
    # Step 8: Format MIC string
    # ----------------------------------------------------------------
    if is_na or mic_snapped is None:
        mic_string = "N/A"
        mic_value_for_mask = None
    else:
        mic_string = format_mic_string(mic_snapped, strip_sequence_text)
        mic_value_for_mask = mic_snapped

    # ----------------------------------------------------------------
    # Step 9: Create MICMask image
    # Binary mask: pixels belonging to label(s) whose concentration matches
    # the MIC strip value are set to 255, all others are 0.
    # If MIC is "N/A", output an all-black image.
    # ----------------------------------------------------------------
    mask_ip = FloatProcessor(w, h)  # starts with all 0.0

    if mic_string != "N/A" and mic_value_for_mask is not None:
        # Single pass: build label→concentration map on first encounter,
        # set mask immediately. 1:1 invariant: each label ID maps to exactly
        # one concentration value, so we can cache and reuse the
concentration
        # without needing a two-pass scan.
        label_conc = {}
        y = 0
        while y < h:
            x = 0
            while x < w:
                label_id = int(ip_labels.getf(x, y))
                if label_id > 0:
                    if label_id not in label_conc:
                        label_conc[label_id] = ip_value.getf(x, y)
                    if abs(label_conc[label_id] - mic_value_for_mask) < 1e-6:
                        mask_ip.setf(x, y, 255.0)
                x += 1
            y += 1

    mask_img = ImagePlus("MICMask", mask_ip)

    # ----------------------------------------------------------------
    # Step 10: Build output tables
    # ----------------------------------------------------------------

    # Measurements table: one row per concentration for the selected corridor
    measurements_table = ResultsTableData()
    i = 0
    while i < len(measurements):
        m = measurements[i]
```

```
        row = measurements_table.addAndModifyRow()
        row.set("s_value", m["s_value"])
        row.set("log_s_value", m["log_s_value"])
        row.set("mean", m["mean"])
        row.set("min", m["min"])
        row.set("max", m["max"])
        row.set("n_pixels", m["n_pixels"])
        row.set("corridor_width_mm", corridor_width_mm)
        # Add excess growth if available
        if mic_result is not None and i < len(mic_result["excess_growths"]):
            row.set("excess_growth", mic_result["excess_growths"][i])
        else:
            row.set("excess_growth", 0.0)
        i += 1

    # Free intermediate data
    measurements_wide = None
    measurements_narrow = None

    # -----------------------------------------------------------------
    # Step 11: Output with text annotations
    # -----------------------------------------------------------------
    d_high_str = str(round(d_high, 2)) if d_high is not None else ""

    annotations = [
        JIPipeTextAnnotation("MIC", str(mic_string)),
        JIPipeTextAnnotation("is_na", "true" if is_na else "false"),
        JIPipeTextAnnotation("na_reason", str(na_reason) if na_reason else
""),
        JIPipeTextAnnotation("is_narrow", "true" if is_narrow else "false"),
        JIPipeTextAnnotation("corridor_width_mm", str(corridor_width_mm)),
        JIPipeTextAnnotation("d_high", d_high_str),
        JIPipeTextAnnotation("mic_threshold", str(DEFAULT_MIC_THRESHOLD)),
        JIPipeTextAnnotation("na_detection_enabled", "true" if
ENABLE_NA_DETECTION else "false"),
    ]

    data_batch.addOutputData(
        output_slot_measurements,
        measurements_table,
        annotations,
        JIPipeTextAnnotationMergeMode.OverwriteExisting,
        progress_info
    )

    data_batch.addOutputData(
        output_slot_mask,
        ImagePlusData(mask_img),
```

```
        annotations,
        JIPipeTextAnnotationMergeMode.OverwriteExisting,
        progress_info
    )

    progress_info.log(
        "MIC=" + str(mic_string) +
        ", is_na=" + str(is_na) +
        ", is_narrow=" + str(is_narrow) +
        ", corridor_width=" + str(corridor_width_mm) + "mm" +
        ", d_high=" + d_high_str
    )

except Exception as e:
    # Error handling: output empty tables with N/A annotations
    progress_info.log("ERROR in find_mic_node_v4: " + str(e))

    measurements_table = ResultsTableData()

    error_annotations = [
        JIPipeTextAnnotation("MIC", "N/A"),
        JIPipeTextAnnotation("is_na", "true"),
        JIPipeTextAnnotation("na_reason", "error: " + str(e)),
        JIPipeTextAnnotation("is_narrow", ""),
        JIPipeTextAnnotation("corridor_width_mm", ""),
        JIPipeTextAnnotation("d_high", ""),
        JIPipeTextAnnotation("mic_threshold", str(DEFAULT_MIC_THRESHOLD)),
    ]

    try:
        data_batch.addOutputData(
            output_slot_measurements,
            measurements_table,
            error_annotations,
            JIPipeTextAnnotationMergeMode.OverwriteExisting,
            progress_info
        )
    except Exception:
        pass

    try:
        error_mask_ip = FloatProcessor(1, 1)
        error_mask_img = ImagePlus("MICMask", error_mask_ip)
        data_batch.addOutputData(
            output_slot_mask,
            ImagePlusData(error_mask_img),
            error_annotations,
            JIPipeTextAnnotationMergeMode.OverwriteExisting,
```

```
            progress_info
        )
    except Exception:
        pass
```

##### *Node #97 "Annotations to table" of type "Convert to annotation table"*

- Input "Input" of node #97 receives data from output "Measurements" of node #96

##### *Node #98 "Output" of type "Compartment output"*

- Input "Measurements" of node #98 receives data from output "Measurements" of node #96
- Input "MICMask" of node #98 receives data from output "MICMask" of node #96
- The parameter "jipipe:compartment:output-slot-name" (jipipe:compartment:output-slot-name) is set to **"Output"**

##### *Node #99 "Remove annotation"*

- Input "Input" of node #99 receives data from output "Output" of node #97
- Input "Input" of node #99 receives data from output "Output" of node #92
- The parameter "Removed annotations" (annotation-type) is set to **true**

##### *Node #100 "Merge table rows"*

- Input "Input" of node #100 receives data from output "Output" of node #99
- The parameter "Grouping method" in category "Merging iteration step generation" (jipipe:data-batch-generation/column-matching) is set to **"MergeAll"**

##### *Node #101 "Remove table column"*

- Input "Input" of node #101 receives data from output "Output" of node #100
- The parameter "Filters" (filters) is set to **NOT STRING_STARTS_WITH(value, "#") AND value != "MIC" AND value != "StripSequence"**

##### *Node #102 "Remove table column"*

- Input "Input" of node #102 receives data from output "Output" of node #101
- The parameter "Filters" (filters) is set to **value == "#GroupRow"**

##### *Node #103 "Merge table columns (supplement)"*

- Input "Source" of node #103 receives data from output "Output" of node #102
- Input "Target" of node #103 receives data from output "Output" of node #90
- The parameter "Column filter" (column-filter) is set to **value IN ARRAY("MIC")**
- The parameter "Reference columns" (reference-columns) is set to **value == "#ImageId"**

##### *Node #104 "Sort table columns"*

- Input "Input" of node #104 receives data from output "Output" of node #103
- The parameter item #1 of "Manual order" (manual-order) is set to **"FileName"**
- The parameter item #2 of "Manual order" (manual-order) is set to **"#ImageId"**
- The parameter item #3 of "Manual order" (manual-order) is set to **"#AssayType"**
- The parameter item #4 of "Manual order" (manual-order) is set to **"#Experiment"**
- The parameter item #5 of "Manual order" (manual-order) is set to **"#Sample"**
- The parameter item #6 of "Manual order" (manual-order) is set to **"#TimePoint"**
- The parameter item #7 of "Manual order" (manual-order) is set to **"StripSequence"**
- The parameter item #8 of "Manual order" (manual-order) is set to **"PixelSize"**
- The parameter item #9 of "Manual order" (manual-order) is set to **"GroupColumn"**
- The parameter item #10 of "Manual order" (manual-order) is set to **"GroupRow"**
- The parameter item #11 of "Manual order" (manual-order) is set to **"ExpectedMIC"**
- The parameter item #12 of "Manual order" (manual-order) is set to **"MIC"**

###### *Node #105 "Export table"*

- Input "Input" of node #105 receives data from output "Output" of node #104
- The parameter "File path" (file-path) is set to **PATH_COMBINE(project_data_dir.tmp_dir, "results", "mic_results")**

###### *Node #106 "Export table"*

- Input "Input" of node #106 receives data from output "Output" of node #104
- The parameter "File format" (file-format) is set to **"XLSX"**
- The parameter "File path" (file-path) is set to **PATH_COMBINE(project_data_dir.tmp_dir, "results", "mic_results")**

#### Compartment C7 “Visualize measurements”

This compartment visualizes the measurements and exports them as image files. A screenshot is provided in **Supplementary Figure 43**.

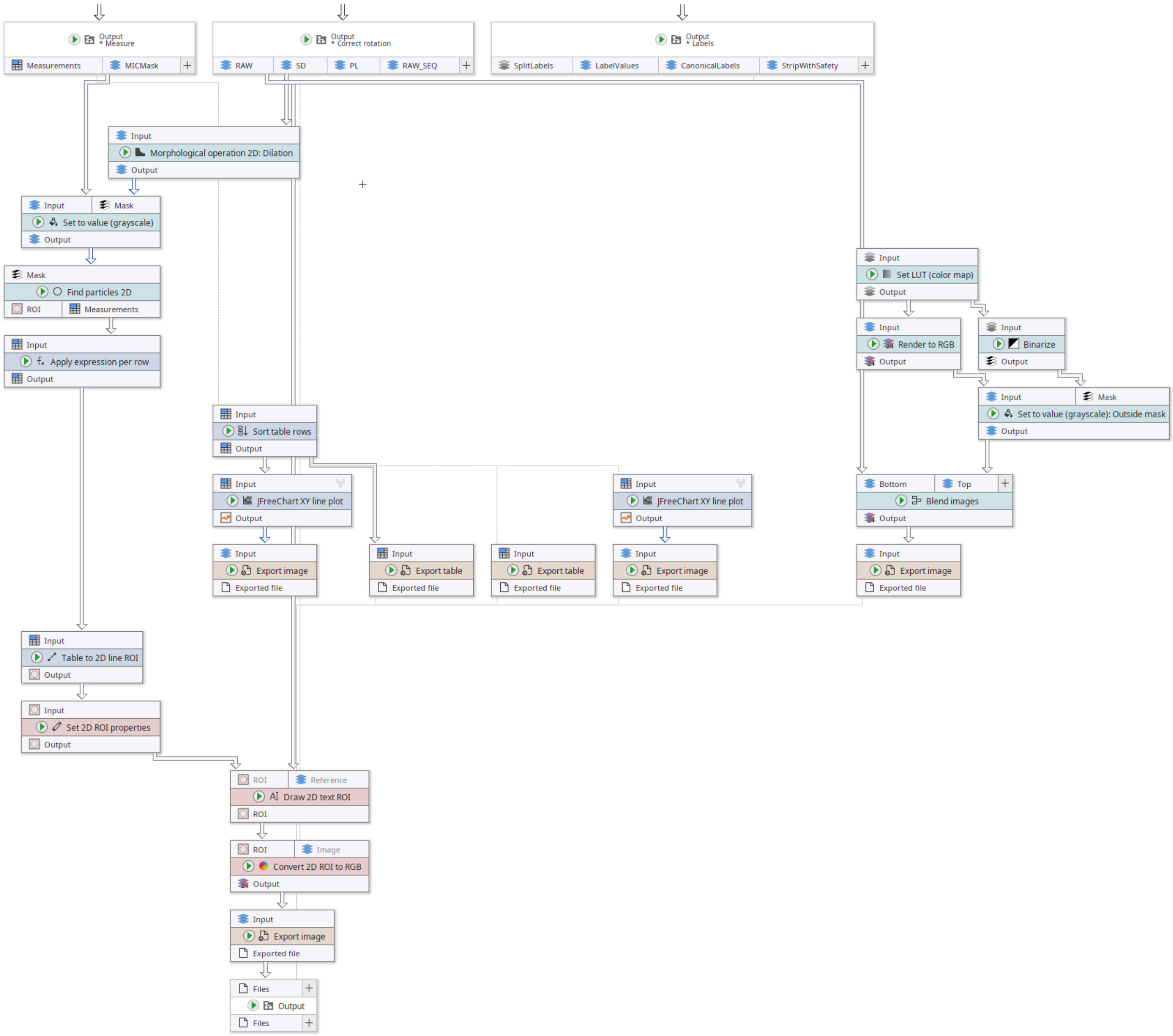

**Supplementary Figure 43:** “Visualize measurements” compartment of the MIC analysis pipeline.

##### *Node #55 "Output" of type "Compartment output"*

- Input "RAW" of node #55 receives data from output "Output" of node #50
- Input "SD" of node #55 receives data from output "Output" of node #54
- Input "PL" of node #55 receives data from output "Output" of node #53
- Input "RAW_SEQ" of node #55 receives data from output "RAW" of node #24
- The parameter "jipipe:compartment:output-slot-name" (jipipe:compartment:output-slot-name) is set to **"Output"**

##### *Node #107 "Morphological operation 2D: Dilation" of type "Morphological operation 2D"*

- Input "Input" of node #107 receives data from output "SD" of node #55
- The parameter "Structure element" (element) is set to **"SQUARE"**
- The parameter item #1 of "Overridden parameters" in category "Adaptive parameters" (jipipe:adaptive-parameters/overridden-parameters) is set to **(3 / NUM(PixelSizeRescaled), "radius")**

##### *Node #77 "Output" of type "Compartment output"*

- Input "SplitLabels" of node #77 receives data from output "Output" of node #76
- Input "LabelValues" of node #77 receives data from output "Data" of node #73
- Input "CanonicalLabels" of node #77 receives data from output "Output" of node #75
- Input "StripWithSafety" of node #77 receives data from output "Output" of node #70
- The parameter "jipipe:compartment:output-slot-name" (jipipe:compartment:output-slot-name) is set to **"Output"**

*Node #108 "Set LUT (color map)"*

- Input "Input" of node #108 receives data from output "CanonicalLabels" of node #77
- The parameter "Color map" (color-map) is set to **"glasbey"**

*Node #109 "Binarize"*

- Input "Input" of node #109 receives data from output "Output" of node #108

*Node #110 "Render to RGB"*

- Input "Input" of node #110 receives data from output "Output" of node #108

*Node #111 "Set to value (grayscale): Outside mask"*

- Input "Input" of node #111 receives data from output "Output" of node #110
- Input "Mask" of node #111 receives data from output "Output" of node #109
- The parameter "Only apply to ..." (roi:target-area) is set to **"OutsideMask"**

*Node #112 "Blend images"*

- Input "Bottom" of node #112 receives data from output "RAW" of node #55
- Input "Top" of node #112 receives data from output "Output" of node #111
- The parameter "Skip incomplete data sets" in category "Input management" (jipipe:data-batch-generation/skip-incomplete) is set to **true**
- The parameter "Bottom" in category "Layers" (layers/Bottom) is set to **{"opacity":1.0}**
- The parameter "Top" in category "Layers" (layers/Top) is set to **{"opacity":1.0}**

*Node #113 "Export image"*

- Input "Input" of node #113 receives data from output "Output" of node #112
- The parameter "File path" (file-path) is set to **PATH_COMBINE(project_data_dir.tmp_dir, "results", "measurement_area", #ImageId + "_" + #AssayType + "_" + #Experiment + "_" + #Sample + "_" + #TimePoint)**

*Node #98 "Output" of type "Compartment output"*

- Input "Measurements" of node #98 receives data from output "Measurements" of node #96
- Input "MICMask" of node #98 receives data from output "MICMask" of node #96
- The parameter "jipipe:compartment:output-slot-name" (jipipe:compartment:output-slot-name) is set to **"Output"**

*Node #114 "Sort table rows"*

- Input "Input" of node #114 receives data from output "Measurements" of node #98
- The parameter item #1 of "Filters" (sort-order) is set to **("s_value", "Ascending")**

*Node #115 "Set to value (grayscale)"*

- Input "Input" of node #115 receives data from output "MICMask" of node #98
- Input "Mask" of node #115 receives data from output "Output" of node #107
- The parameter "Only apply to ..." (roi:target-area) is set to **"OutsideMask"**

*Node #116 "Export table"*

- Input "Input" of node #116 receives data from output "Output" of node #114
- The parameter "File path" (file-path) is set to **PATH_COMBINE(project_data_dir.tmp_dir, "results", "raw_values", #ImageId + "_" + #AssayType + "_" + #Experiment + "_" + #Sample + "_" + #TimePoint)**

*Node #117 "JFreeChart XY line plot"*

- Input "Input" of node #117 receives data from output "Output" of node #114
- The parameter "X" in category "Input columns" (input-columns/X) is set to **("ExistingColumn", "s_value")**
- The parameter "Y" in category "Input columns" (input-columns/Y) is set to **("ExistingColumn", "mean")**
- The parameter "X axis label" in category "Plot parameters" (plot-parameters/x-axis-label) is set to **"E-Test strip value"**
- The parameter "Y axis label" in category "Plot parameters" (plot-parameters/y-axis-label) is set to **"Mean intensity"**

*Node #118 "Export table"*

- Input "Input" of node #118 receives data from output "Output" of node #114
- The parameter "File format" (file-format) is set to **"XLSX"**
- The parameter "File path" (file-path) is set to **PATH_COMBINE(project_data_dir.tmp_dir, "results", "raw_values", #ImageId + "_" + #AssayType + "_" + #Experiment + "_" + #Sample + "_" + #TimePoint)**

*Node #119 "JFreeChart XY line plot"*

- Input "Input" of node #119 receives data from output "Output" of node #114
- The parameter "X" in category "Input columns" (input-columns/X) is set to **("ExistingColumn", "log_s_value")**
- The parameter "Y" in category "Input columns" (input-columns/Y) is set to **("ExistingColumn", "mean")**
- The parameter "X axis label" in category "Plot parameters" (plot-parameters/x-axis-label) is set to **"log(E-Test strip value)"**
- The parameter "Y axis label" in category "Plot parameters" (plot-parameters/y-axis-label) is set to **"Mean intensity"**

*Node #120 "Find particles 2D"*

- Input "Mask" of node #120 receives data from output "Output" of node #115

*Node #121 "Export image"*

- Input "Input" of node #121 receives data from output "Output" of node #117
- The parameter "File path" (file-path) is set to **PATH_COMBINE(project_data_dir.tmp_dir, "results", "plots", #ImageId + "_" + #AssayType + "_" + #Experiment + "_" + #Sample + "_" + #TimePoint)**

*Node #122 "Export image"*

- Input "Input" of node #122 receives data from output "Output" of node #119
- The parameter "File path" (file-path) is set to **PATH_COMBINE(project_data_dir.tmp_dir, "results", "plots_log", #ImageId + "_" + #AssayType + "_" + #Experiment + "_" + #Sample + "_" + #TimePoint)**

*Node #123 "Apply expression per row"*

- Input "Input" of node #123 receives data from output "Measurements" of node #120
- The parameter item #1 of "Generated values" (entries) is set to **column-name = "X1", value = BX**
- The parameter item #2 of "Generated values" (entries) is set to **column-name = "Y1", value = BY**
- The parameter item #3 of "Generated values" (entries) is set to **column-name = "X2", value = BX + Width**
- The parameter item #4 of "Generated values" (entries) is set to **column-name = "Y2", value = BY**

*Node #124 "Table to 2D line ROI"*

- Input "Input" of node #124 receives data from output "Output" of node #123

*Node #125 "Set 2D ROI properties"*

- Input "Input" of node #125 receives data from output "Output" of node #124
- The parameter "Line width" (line-width) is set to **5.0**
- The parameter "Line color" (line-color) is set to **"#FF3333"**

*Node #126 "Draw 2D text ROI"*

- Input "ROI" of node #126 receives data from output "Output" of node #125
- Input "Reference" of node #126 receives data from output "RAW" of node #55
- The parameter "Background margin" (background-margin) is set to **{"left":{"expression":"15"},"top":{"expression":"15"},"right":{"expression":"15"},"bottom":{"expression":"15"}}**
- The parameter "Font size" (font-size) is set to **50**
- The parameter "Background color" (background-color) is set to **"#000000"**
- The parameter "Location" (location) is set to **{"left":{"expression":"0"},"top":{"expression":"15"},"right":{"expression":"15"},"bottom":{"expression":"0"},"anchor":"TopRight"}**
- The parameter "Text" (text) is set to **IF_ELSE(HAS_VARIABLE("ExpectedMIC"), "MIC = " + GET_VARIABLE("MIC", "OCR_FAILURE") + new_line + "Expected MIC = " + GET_VARIABLE("ExpectedMIC", "?"), "MIC = " + GET_VARIABLE("MIC", "OCR_FAILURE"))**

- The parameter "Skip incomplete data sets" in category "Input management" (jipipe:data-batch-generation/skip-incomplete) is set to **true**

***Node #127 "Convert 2D ROI to RGB"***

- Input "ROI" of node #127 receives data from output "ROI" of node #126
- Input "Image" of node #127 receives data from output "RAW" of node #55
- The parameter "Override line width" (override-line-width) is set to **[Disabled]**
- The parameter "Override line color" (override-line-color) is set to **[Disabled]**
- The parameter "Skip incomplete data sets" in category "Input management" (jipipe:data-batch-generation/skip-incomplete) is set to **true**

***Node #128 "Export image"***

- Input "Input" of node #128 receives data from output "Output" of node #127
- The parameter "File path" (file-path) is set to **PATH_COMBINE(project_data_dir.tmp_dir, "results", "mic", #ImageId + "_" + #AssayType + "_" + #Experiment + "_" + #Sample + "_" + #TimePoint)**

***Node #129 "Output" of type "Compartment output"***

- Input "Files" of node #129 receives data from output "Exported file" of node #128
- Input "Files" of node #129 receives data from output "Exported file" of node #118
- Input "Files" of node #129 receives data from output "Exported file" of node #122
- Input "Files" of node #129 receives data from output "Exported file" of node #116
- Input "Files" of node #129 receives data from output "Exported file" of node #113
- The parameter "jipipe:compartment:output-slot-name" (jipipe:compartment:output-slot-name) is set to **"Output"**

### 4 J-AST application

| **Feature** | ***diskImageR*** | ***J-AST*** |
|---|---|---|
| **Distribution** | *Windows* application (*R*, *ImageJ*/*Fiji*) | Multi-platform web application and desktop software (*Spring*, *Electron*, *JIPipe*) |
| **Plate and disk detection** | Automatic | Automatic and manual |
| **Supported assay types** | DDAs | DDAs and Etests |
| **(Meta)data management** | No | Yes, automatic and manual |
| **Results management** | No | Yes |

**Supplementary Table 1**: Comparison of *diskImageR* and *J-AST* features.

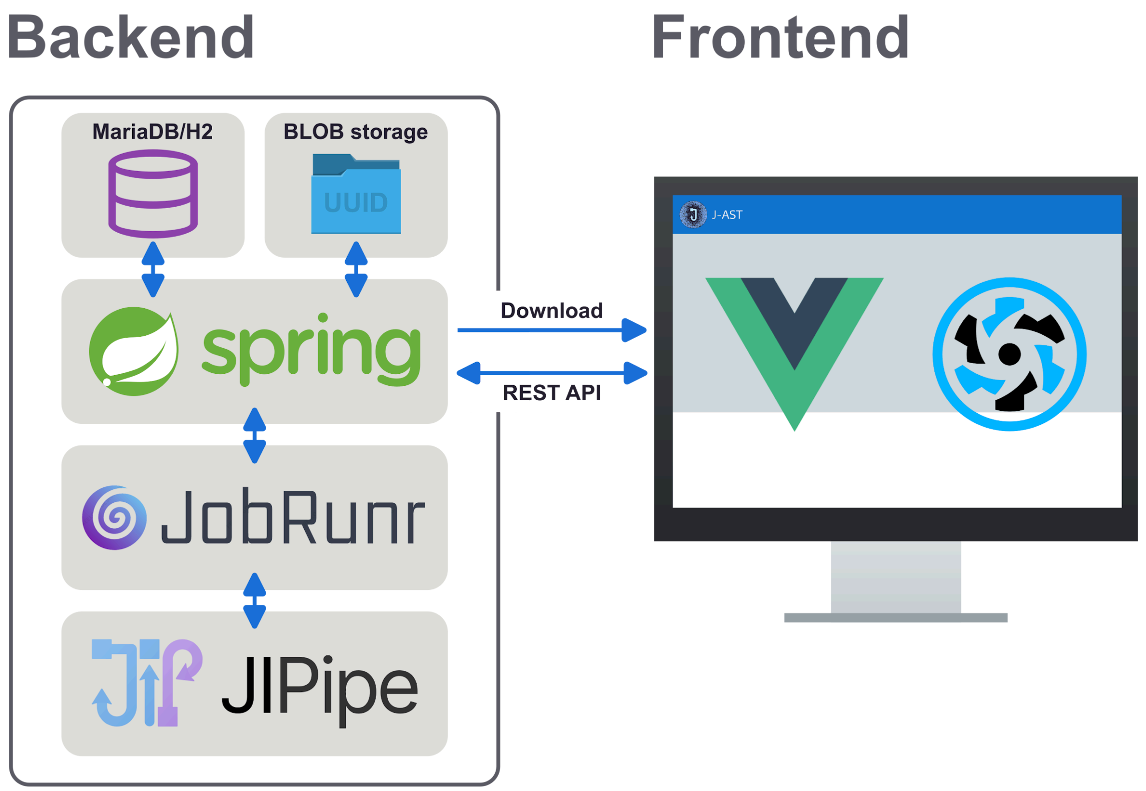

**Supplementary Figure 44**: Architecture of the *J-AST* platform. The tool is divided into a frontend and a backend component. The web browser or Electron downloads the frontend web application built on *Vue* and *Quasar* from the *Spring* server, which communicates with the backend's API via REST. The backend manages relational data in *MariaDB* or *H2*, and larger files inside a dedicated BLOB storage. Image processing tasks are scheduled via *JobRunr*, which runs the appropriate *JIPipe* pipelines for the selected operation.

#### 4.1 Backend

| Library | Version | Authors |
|---|---|---|
| JIPipe | 5.3.0 | Gerst, R., Cseresnyés, Z., & Figge, M. T. (2023). JIPipe: visual batch processing for ImageJ. *nature methods*, *20*(2), 168-169. |
| Spring Boot | 3.3.12 | Broadcom Inc. |
| Spring Boot Starter Data JPA | 3.3.12 | Broadcom Inc. |
| Spring Boot Starter Security | 3.3.12 | Broadcom Inc. |
| Spring Boot Starter Web | 3.3.12 | Broadcom Inc. |
| Spring Boot Configuration Processor | 3.3.12 | Broadcom Inc. |
| Spring Boot Starter Validation | 3.3.12 | Broadcom Inc. |
| Spring Boot Dev Tools | 3.3.12 | Broadcom Inc. |

| | | |
|---|---|---|
| H2 | 2.2.224 | Thomas Müller |
| MariaDB Java Client | 3.3.4 | MariaDB Foundation |
| JobRunR Spring Boot 3 Starter | 7.4.0 | JobRunr by Rosoco BV |
| Guava | 33.0.0-jre | Alphabet Inc. |
| Apache Commons Text | 1.11.0 | The Apache Software Foundation |
| Apache Commons IO | 2.14.0 | The Apache Software Foundation |
| Apache Commons Lang3 | 3.12.0 | The Apache Software Foundation |
| Apache Commons Compress | 1.26.0 | The Apache Software Foundation |
| Apache Commons Exec | 1.3 | The Apache Software Foundation |
| Apache Commons CSV | 1.10.0 | The Apache Software Foundation |
| JGraphT Core | 1.5.1 | Barak Naveh et al. |
| JNA Platform | 5.13.0 | Timothy Wall, Matthias Bläsing et al. |
| Java JWT | 0.12.6 | Les Hazlewood et al. |
| Java JWT Jackson | 0.12.6 | Les Hazlewood et al. |
| Reflections | 0.10.2 | ronmamo (GitHub) |
| Hypersistence Utils Hibernate 63 | 3.9.0 | Vlad Mihalcea |
| jsoup | 1.20.1 | Jonathan Hedley |

**Supplementary Table 2**: Dependencies of the backend component.

#### 4.2 Frontend

| Library | Version | Authors |
|---|---|---|
| Node | 20.19.0 | OpenJS Foundation |
| NPM | 6.13.4 | Microsoft Corporation |
| Electron Builder | 24.13.3 | Mike Maietta et al. |
| Vite | 1.9.0 | VoidZero Inc. & Vite Contributors |
| Electron | 36.2.1 | OpenJS Foundation |
| Typescript | 5.5.4 | Microsoft Corporation |
| Quasar | 2.16.0 | PULSARDEV SRL, Razvan Stoenescu |

| | | |
|---|---|---|
| Quasar Extras | 1.16.4 | PULSARDEV SRL, Razvan Stoenescu |
| VueUse | 11.2.0 | Anthony Fu and VueUse contributors |
| Axios | 1.2.1 | axios community https://github.com/axios |
| Class Transformer | 0.5.1 | TypeStack community https://github.com/typestack |
| Email Validator | 2.0.4 | Manish Saraan |
| JSZip | 3.10.1 | Stuart Knightley |
| JWT-Decode | 4.0.0 | Okta, Inc. |
| Konva | 9.3.16 | konva community https://github.com/konvajs |
| Papaparse | 5.5.1 | Matt Holt |
| Pinia | 2.0.11 | Eduardo San Martin Morote |
| Q-Floodfill | 1.3.1 | Pavel Kukov |
| Vue | 3.4.18 | Evan You et al. |
| Vue-Konva | 3.1.2 | konva community https://github.com/konvajs |
| Vue-Router | 4.0.12 | Evan You et al. |
| YAML | 2.8.0 | Eemeli Aro |

**Supplementary Table 3**: Dependencies of the frontend component.

### 5 Benchmarking setup and evaluation datasets

| Strain name | Source |
|---|---|
| SC5314 | Gillum, A. M. et al. Isolation of the Candida albicans gene for orotidine-5'-phosphate decarboxylase by complementation of S. cerevisiae ura3 and Escherichia coli pyrF mutations. Mol. Gen. Genet. 1984. 198:179-182. |
| 12C<br>19F<br>GC75<br>L26<br>P34048<br>P37005<br>P37037<br>P57072<br>P60002<br>P75016<br>P75063<br>P76055<br>P78042<br>P78048<br>P87 | Hirakawa, M. et al. Genetic and phenotypic intra-species variation in Candida albicans. Genome Res. 2015. 25: 413-425 |
| 81/064<br>AM2003/0089<br>AM2003/0165<br>AM2004/0028<br>AM2005/0377<br>FC22<br>FJ9<br>HUN96<br>IHEM16945<br>J981315<br>J981318<br>L1086<br>RV4688<br>T101<br>YSU751 | MacCallum, D. et al. Property differences among the four major Candida albicans strain clades. Eukaryot. Cell. 2009. 8: 373-387 |

| | |
|---|---|
| OKP90 | Blignaut, E. et al. Ca3 fingerprinting of Candida albicans isolates from human immunodeficiency virus-positive and healthy individuals reveals a new clade in South Africa. J. Clin. Microbiol. 2002. 30:826-836 |

**Supplementary Table 4**: Strains used in this study.

### 6 MIC detection results

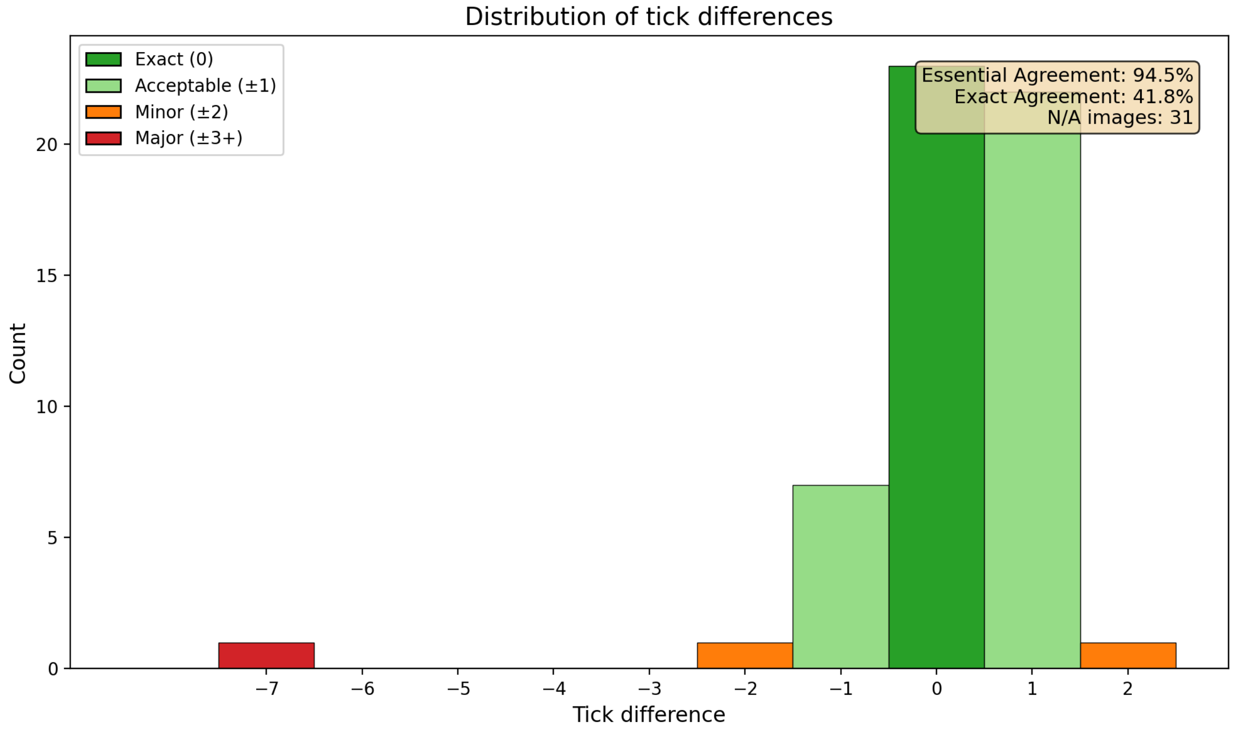

**Supplementary Figure 45:** MIC detection performance. 94.5% of resulting MIC results are within the acceptable range.

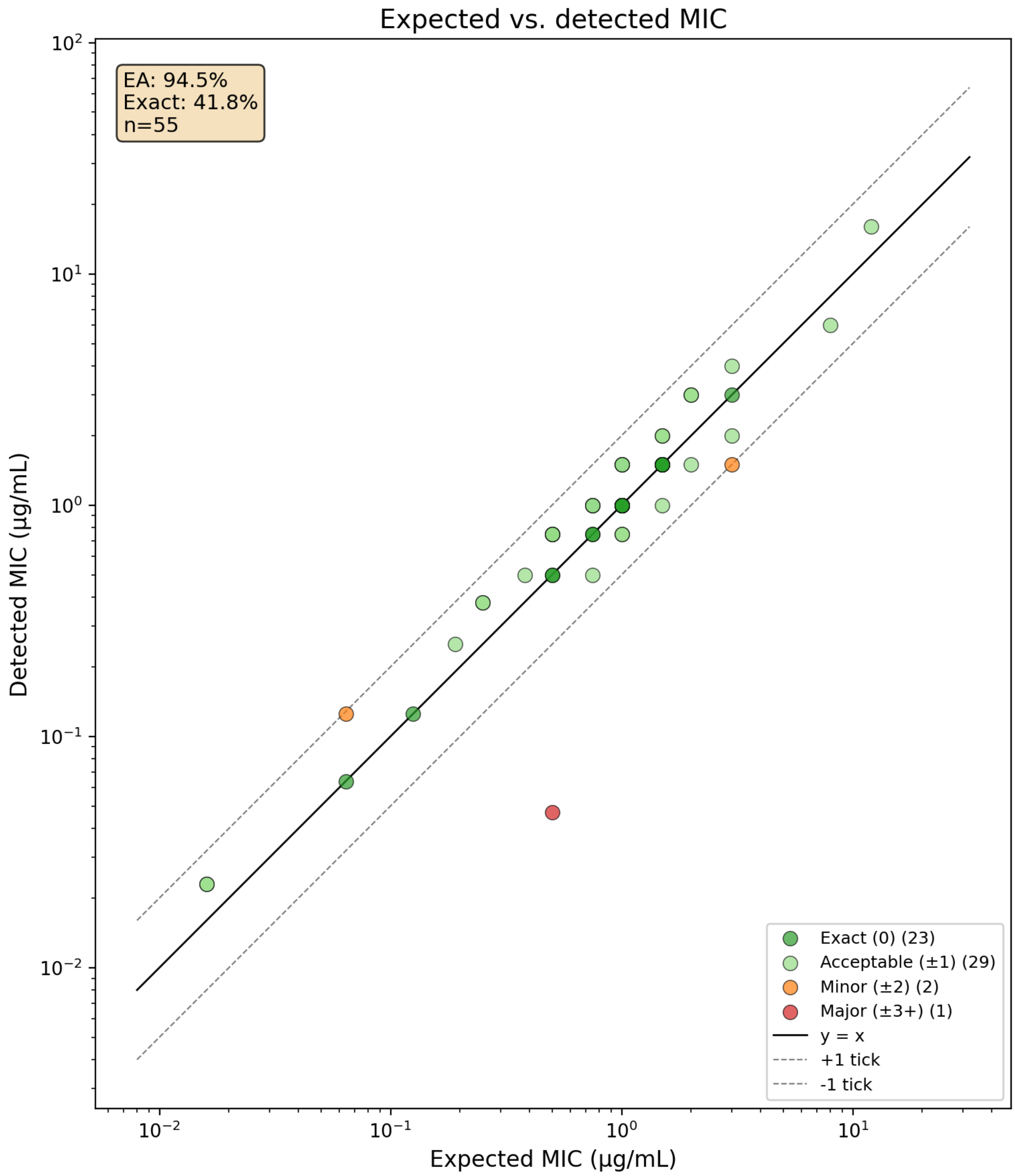

**Supplementary Figure 46:** MIC detection performance.

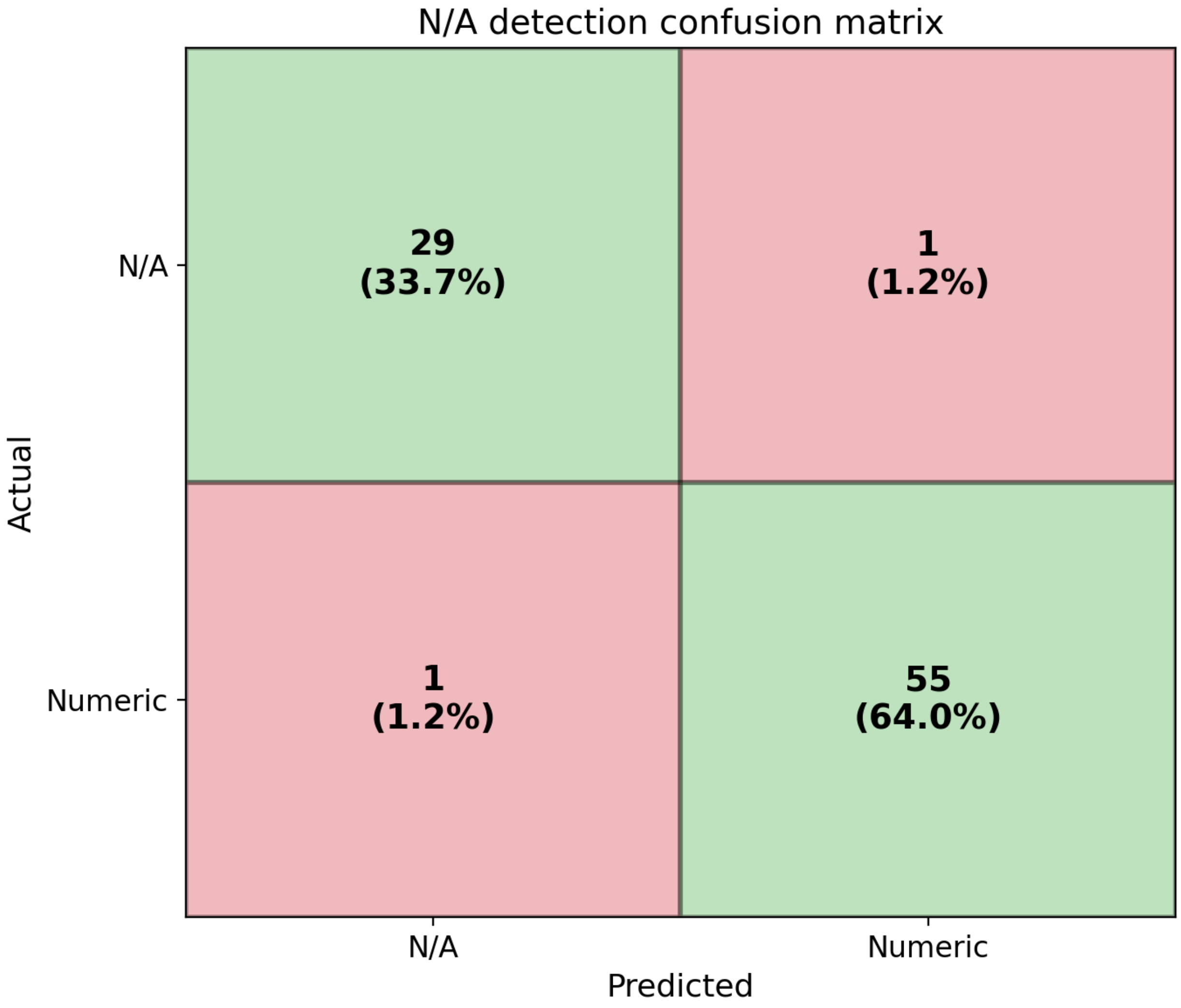

**Supplementary Figure 47:** Performance of the detector algorithm for images without MIC.

| FileName | OCR | Expected MIC | Detected MIC | Tick difference | NA classifier result |
|---|---|---|---|---|---|
| 10271.png | Successful | 0.75 | N/A | | FP |
| 10274.png | Successful | N/A | N/A | | TP |
| 10310.png | Successful | 1.5 | 2 | 1 | TN |
| 10302.png | Successful | 0.5 | 0.5 | 0 | TN |
| 10305.png | Successful | 0.016 | 0.023 | 1 | TN |
| 10323.png | Successful | 1.5 | 1 | -1 | TN |
| 10387.png | Successful | 1 | 1 | 0 | TN |
| 10318.png | Successful | 1 | 1 | 0 | TN |

| | | | | | |
|---|---|---|---|---|---|
| 10344.png | Successful | 1 | 1 | 0 | TN |
| 10270.png | Successful | 1 | 0.75 | -1 | TN |
| 10329.png | Successful | N/A | N/A | | TP |
| 10276.png | Successful | N/A | N/A | | TP |
| 10321.png | Successful | N/A | N/A | | TP |
| 10311.png | Successful | 1 | 0.75 | -1 | TN |
| 10293.png | Successful | 0.19 | 0.25 | 1 | TN |
| 10397.png | Successful | N/A | N/A | | TP |
| 10366.png | Successful | 3 | 1.5 | -2 | TN |
| 10333.png | Successful | 1 | 1 | 0 | TN |
| 10346.png | Successful | 2 | 3 | 1 | TN |
| 10388.png | Successful | 0.5 | 0.5 | 0 | TN |
| 10294.png | Successful | 0.75 | 1 | 1 | TN |
| 10261.png | Successful | 0.5 | 0.047 | -7 | TN |
| 10395.png | Successful | N/A | N/A | | TP |
| 10386.png | Successful | 1.5 | 1.5 | 0 | TN |
| 10292.png | Successful | 0.25 | 0.38 | 1 | TN |
| 10301.png | Successful | 1 | 1.5 | 1 | TN |
| 10336.png | Successful | N/A | N/A | | TP |
| 10348.png | Successful | 1 | 1 | 0 | TN |
| 10322.png | Successful | 1 | 1 | 0 | TN |
| 10325.png | Successful | 1.5 | 1.5 | 0 | TN |
| 10306.png | Successful | 0.5 | 0.75 | 1 | TN |
| 10390.png | Successful | 2 | 3 | 1 | TN |
| 10389.png | Successful | N/A | N/A | | TP |
| 10347.png | Successful | 0.75 | 1 | 1 | TN |
| 10328.png | Successful | 1 | 1 | 0 | TN |
| 10281.png | Successful | 0.5 | 0.75 | 1 | TN |
| 10255.png | Successful | N/A | N/A | | TP |
| 10331.png | Successful | N/A | N/A | | TP |
| 10324.png | Successful | 1.5 | 1.5 | 0 | TN |

| | | | | | |
|---|---|---|---|---|---|
| 10257.png | Successful | N/A | N/A | | TP |
| 10385.png | Successful | N/A | N/A | | TP |
| 10384.png | Successful | 0.75 | 1 | 1 | TN |
| 10273.png | Successful | N/A | N/A | | TP |
| 10287.png | Successful | 0.064 | 0.125 | 2 | TN |
| 10300.png | Successful | 0.125 | 0.125 | 0 | TN |
| 10335.png | Successful | 3 | 3 | 0 | TN |
| 10369.png | Successful | 1 | 1.5 | 1 | TN |
| 10308.png | Successful | 1 | 1.5 | 1 | TN |
| 10256.png | Successful | 0.064 | 0.064 | 0 | TN |
| 10337.png | Successful | 3 | 4 | 1 | TN |
| 10282.png | Successful | N/A | N/A | | TP |
| 10382.png | Successful | N/A | N/A | | TP |
| 10262.png | Successful | N/A | N/A | | TP |
| 10340.png | Successful | N/A | N/A | | TP |
| 10297.png | Successful | 0.38 | 0.5 | 1 | TN |
| 10264.png | Successful | 12 | 16 | 1 | TN |
| 10291.png | Successful | 0.5 | 0.75 | 1 | TN |
| 10332.png | Successful | 1.5 | 1.5 | 0 | TN |
| 10317.png | Successful | 3 | 2 | -1 | TN |
| 10312.png | Successful | 1 | 1 | 0 | TN |
| 10398.png | Successful | N/A | N/A | | TP |
| 10363.png | Successful | 1.5 | 1.5 | 0 | TN |
| 10339.png | Successful | N/A | N/A | | TP |
| 10334.png | Successful | 1 | 1 | 0 | TN |
| 10393.png | Successful | N/A | N/A | | TP |
| 10279.png | Successful | 0.25 | 0.38 | 1 | TN |
| 10290.png | Successful | N/A | N/A | | TP |
| 10272.png | Successful | N/A | N/A | | TP |
| 10351.png | Successful | N/A | 0.5 | | FN |
| 10309.png | Successful | 8 | 6 | -1 | TN |

| | | | | | |
|---|---|---|---|---|---|
| 10268.png | Successful | 0.016 | 0.023 | 1 | TN |
| 10343.png | Successful | N/A | N/A | | TP |
| 10283.png | Successful | N/A | N/A | | TP |
| 10326.png | Successful | 0.75 | 0.75 | 0 | TN |
| 10296.png | Successful | N/A | N/A | | TP |
| 10371.png | Successful | 1.5 | 2 | 1 | TN |
| 10379.png | Successful | 2 | 1.5 | -1 | TN |
| 10253.png | Successful | 0.75 | 0.75 | 0 | TN |
| 10355.png | Failed | N/A | N/A | | TP |
| 10350.png | Successful | 0.5 | 0.75 | 1 | TN |
| 10361.png | Successful | 1.5 | 1.5 | 0 | TN |
| 10258.png | Successful | N/A | N/A | | TP |
| 10314.png | Successful | N/A | N/A | | TP |
| 10354.png | Successful | N/A | N/A | | TP |
| 10252.png | Successful | 1 | 1 | 0 | TN |
| 10357.png | Successful | 0.75 | 0.5 | -1 | TN |

**Supplementary Table 5**: All results of the MIC detection.
